# Cryo-EM structure of enteric adenovirus HAdV-F41 highlights structural divergence among human adenoviruses

**DOI:** 10.1101/2020.07.01.177519

**Authors:** Marta Pérez-Illana, Marta Martínez, Gabriela N. Condezo, Mercedes Hernando-Pérez, Casandra Mangroo, Martha Brown, Roberto Marabini, Carmen San Martín

## Abstract

Enteric adenoviruses are one of the main causes of viral gastroenteritis in the world. To carry out a successful infection, the virions must withstand the harsh conditions found in the gut. This requirement suggests that capsid stability must be different from that of other adenoviruses. We have determined the structure of a human enteric adenovirus, HAdV-F41, at 4.0 Å resolution by single particle averaging cryo-electron microscopy, and compared it with that of other adenoviruses with respiratory (HAdV-C5) and ocular (HAdV-D26) tropisms. While the overall structures of hexon, penton base and internal minor coat proteins IIIa and VIII are conserved, we observe partially ordered elements reinforcing the vertex region, which suggests their role in enhancing the physicochemical capsid stability of HAdV-F41. Unexpectedly, we find an organization of the external minor coat protein IX different from all previously characterized human and non-human mastadenoviruses. Knowledge of the structure of enteric adenoviruses can provide a starting point for the design of vectors suitable for oral delivery or intestinal targeting.

## Main

Adenoviruses (AdVs) are pathogens which can be engineered to become therapeutic tools (Appaiahgari and Vrati, 2015; Baker *et al*., 2018). Over 100 types of human AdVs (HAdV) have been described so far (http://hadvwg.gmu.edu/; https://talk.ictvonline.org/), and grouped in seven species (HAdV-A to G). HAdVs cause acute infections in the respiratory and gastrointestinal tracts, as well as conjunctivitis. Most AdV infections are subclinical, but they can cause significant morbidity and even mortality in immunocompromised patients (Lion, 2014). There are currently no vaccines for use in the general population or approved antiviral therapies for AdV infections.

High resolution (3.2-3.5 Å) structures are available only for two HAdVs: HAdV-C5 and HAdV-D26 (Dai *et al*., 2017; Liu *et al*., 2010; Yu *et al*., 2017). The AdV capsid stands out among the non-enveloped viruses because of its large size (∼950 Å, 150 MDa), triangulation number (*pseudo* T = 25), and complex composition. The facets are formed by 240 trimeric hexons, while pentamers of penton base protein fill the vertices. Receptor-binding trimeric fibres of various lengths, depending on the virus type, project from the penton bases (Nicklin *et al*., 2005). Minor coat proteins IIIa and VIII on the inner capsid surface, and IX on the outer one, contribute to modulate the quasi-equivalent icosahedral interactions. The membrane lytic protein VI competes with genome condensing protein VII for binding to an internal cavity in hexons (Dai *et al*., 2017; Hernando-Perez *et al*., 2020).

HAdVs in species F are unusual in that they have a narrow enteric tropism. HAdV-F40 and HAdV-F41 are one of the three major causes of viral gastroenteritis in young children, together with rota-and noroviruses (Chhabra *et al*., 2013; Corcoran *et al*., 2014). They account for more than half of the AdVs identified in stools of immunocompetent symptomatic children (Brandt *et al*., 1985; Brown, 1990; Uhnoo *et al*., 1984). Differences in the capsid structure of HAdV-F, relative to HAdV-C, may confer stability to gastric conditions en route to the intestine *in vivo*. Indeed, infectivity assays after incubation in acidic conditions have shown that HAdV-F41 is stable, and even possibly activated, at low pH, in contrast to HAdV-C5 (Favier *et al*., 2004). The fact that HAdV-F viruses code for two different fibres has been related with their enteric tropism, although the exact relation is unclear (Favier *et al*., 2002; Kidd *et al*., 1993).

Here we use cryo-EM to obtain the structure of the HAdV-F41 virion, and compare it with the two other known HAdV structures. We interpret the structural differences in the context of the high physicochemical stability of the enteric AdV capsid.

## Results

### Physicochemical stability of HAdV-F41 virions

To evaluate the stability of HAdV-F41 virions, we assessed their infectivity after treatment with synthetic gastric, intestinal, or gastric followed by intestinal fluids. GFP fluorescence measured by flow cytometry was used to determine the number of infected cells in HEK293 cultures. Neither gastric nor intestinal conditions reduced infectivity. Interestingly, consecutive incubation in synthetic gastric and intestinal fluid resulted in a clear increase of GFP expression, indicating enhanced infectivity (**Fig. S1a**). That is, not only is HAdV-F41 resistant to acidic pH as previously shown (Favier *et al*., 2004), but it is also resistant to salt and proteases, and even activated by the combined action of several of these factors.

We also examined the physical stability of HAdV-F41 particles in comparison to HAdV-C5 using extrinsic fluorescence of the DNA intercalating agent YOYO-1 to characterize capsid disruption as a function of temperature. The fluorescence emission of YOYO-1 increased with temperature for both specimens, as genomes became exposed to the solvent (**Fig. S1b)**. While the half-transition temperature (T_0.5_) estimated for HAdV-C5 was 47°C (Hernando-Perez *et al*., 2020), for HAdV-F41 the maximum rate of DNA exposure happened at near 51°C. This result indicates that HAdV-F41 particles are thermally more stable than HAdV-C5.

### Overall HAdV-F41 structure

The cryo-EM map of HAdV-F41 at 4.0 Å resolution (**Fig. 1 and S2, Table S1**) shows the general characteristics common to all previous AdV structures: a ∼950 Å diameter *pseudo* T=25 icosahedral particle with 12 trimeric hexon capsomers per facet, plus the pentameric pentons at the vertices. The N-terminus of protein IX lays on the outer capsid surface, in the valleys formed by hexon trimers (**Fig. 1b top**). Minor coat proteins IIIa and VIII, plus small fragments of VI and core protein VII, can be observed on the inner surface of the icosahedral shell, oriented towards the virion core (**Fig. 1b bottom**). Due to their flexibility and symmetry mismatch with the penton base, fibres cannot be resolved by cryo-EM when imposing icosahedral symmetry as done here.

**Figure 1.**
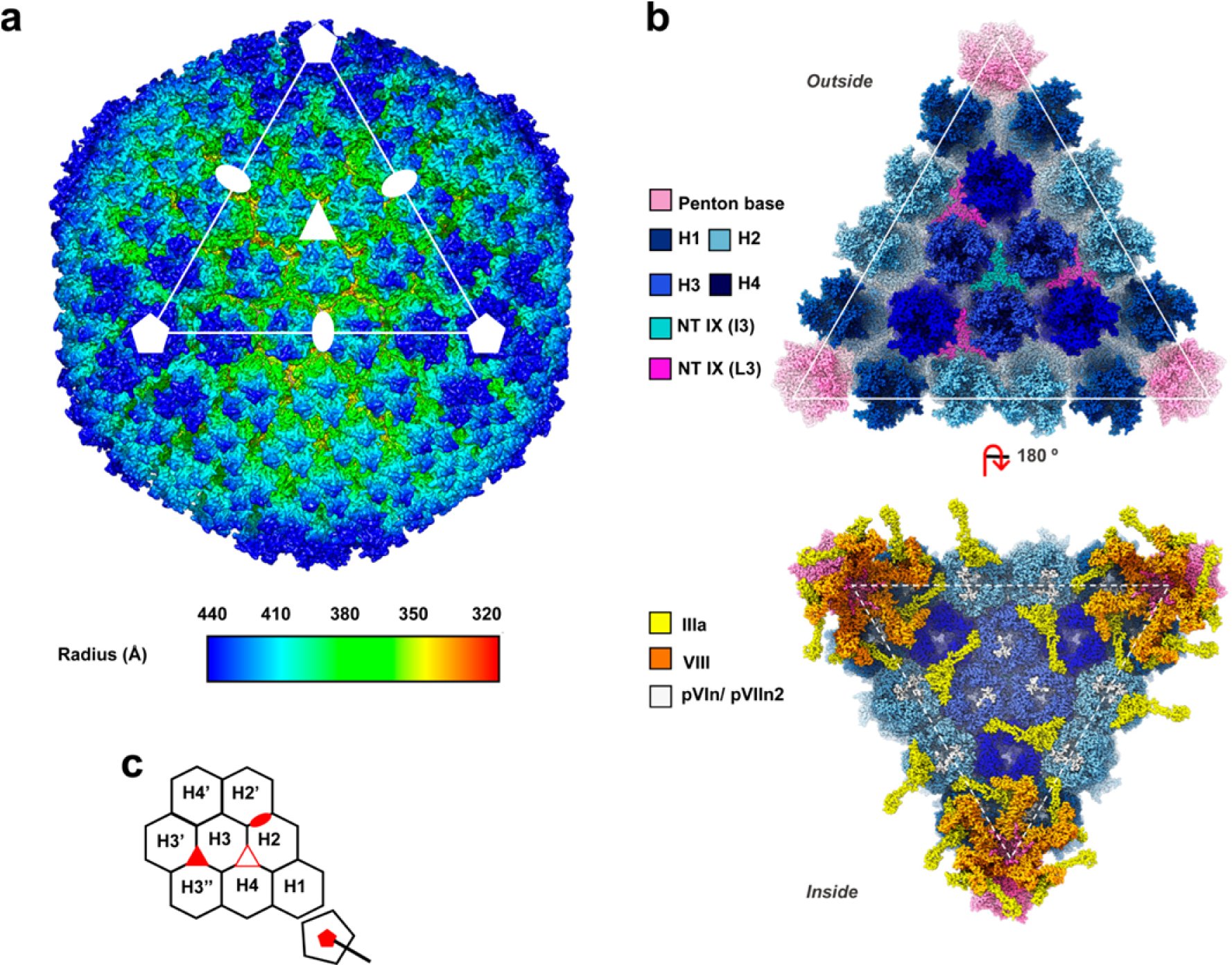
The cryo-EM structure of HAdV-F41 at 4.0 Å. **(a)** Radially coloured surface rendering of the HAdV-F41 map viewed down the 2-fold icosahedral axis. One facet is identified by a white triangle. Icosahedral symmetry axes are indicated with white symbols: 5-fold (pentagon), 3-fold (triangle) and 2-fold (oval). A Gaussian filter has been applied to the map for a clearer appreciation of the main features. **(b)** The molecular models of the major and minor coat proteins traced in the icosahedral HAdV-F41 facet. Top: view from outside the capsid. Bottom: view inside the capsid. Protein colours as indicated by the legend at the left hand side. The four hexon trimers in the AU are numbered H1-H4. **(c)** Cartoon depicting the icosahedral AU and surrounding hexons, as seen from outside the capsid. The icosahedral symmetry axes are indicated with red filled symbols. A hollow red triangle indicates the L3 axis between hexons 2, 3 and 4.

Except for proteins VI and VIII, the HAdV-F41 structural polypeptides are shorter than their HAdV-C5 counterparts (**Table S2**). Sequence identity varies between 52 and 78%. The predicted isoelectric points for hexon and penton base suggest a more basic capsid surface for HAdV-F41 (Favier *et al*., 2004), except for the contribution of the external cementing protein IX, which is remarkably more acidic (**Table S2**). On the inner capsid surface, proteins IIIa, VI and VIII are more acidic in the enteric AdV.

### Major coat protein: hexon

The architecture of the HAdV-F41 hexon is very similar to that of HAdV-C5 (Dai *et al*., 2017) (**Table S3, Fig. 2a**). The double jelly roll in each monomer that forms the trimer pseudo-hexagonal base is highly conserved. As expected (Crawford-Miksza and Schnurr, 1996), the main differences between the structures are in the loops located in the hexon towers, the so-called hyper variable regions (HVRs) which constitute serotype-specific epitopes exposed on the virion surface (**Fig. 2a-b, and Table S4**). As in HAdV-C5, HVRs in HAdV-F41 are flexible (**Fig. S3**), but we were able to fully trace all of them except HVR4 (**Fig. 2a-b, Table S5**). The 32-residue acidic loop in HVR1, unique to HAdVs in species C (Ebner *et al*., 2005), has not been traced in any of the available HAdV-C hexon structures. We were able to completely trace HVR1 in HAdV-F41, probably because its shorter length limits flexibility (**Fig. 2b, Table S4**). Lack of the long acidic region reflects on the absence of a large negatively charged patch in the hexon towers of HAdV-F41 (**Table S4**), and could have an influence in the interplay with host factors under the extreme pH conditions in the gastrointestinal tract. However, the acidic stretch is also absent in HAdV-D26, which has ocular tropism and where HVR1 is also wholly traced (Yu *et al*., 2017). Residue Tyr784 also has RMSD > 2 Å between HAdV-C5 and HAdV-F41, and could be involved in the different interaction with protein IX (see IX section below). Besides the HVRs, other regions already described for HAdV-C5 and D26 also display variability among the twelve hexon monomers in the icosahedral asymmetric unit (AU) in HAdV-F41 (**Fig. S3**).

**Figure 2.**
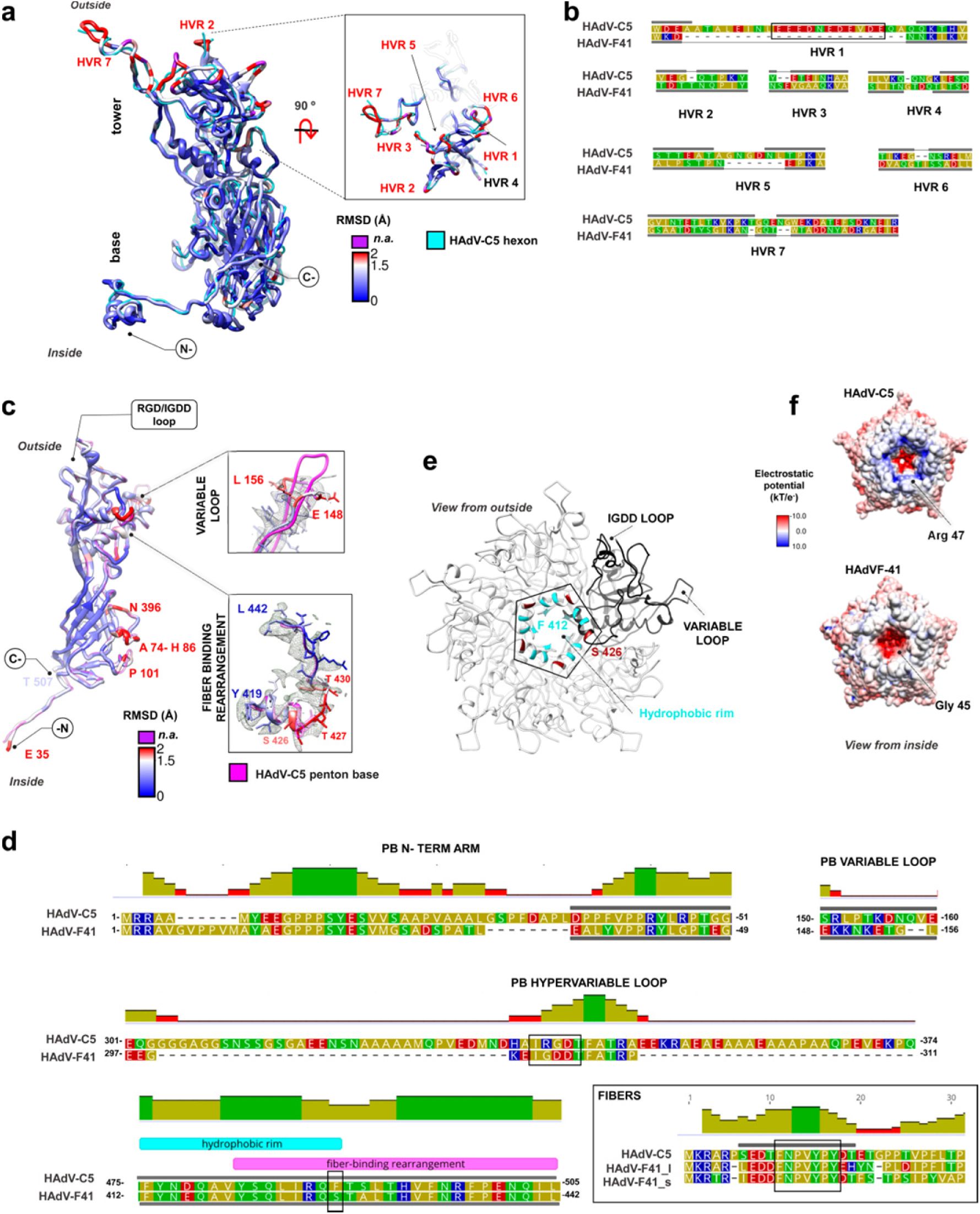
Hexon and penton base structures. **(a)** Superposition of the HAdV-C5 hexon monomer (PDB ID: 6B1T Chain A, cyan) with that of HAdV-F41, coloured by RMSD from blue to red as indicated. Residues exceeding the Chimera RMSD cut-off, or where RMSD calculation is not possible because they are not traced in HAdV-C5, are depicted in purple. The six (out of seven) HVRs traced in the enteric virus are labelled in red, and HVR4 is labelled in black. The **base** and **tower** regions of the capsomer, as well as the orientation with respect to the capsid (**outside/inside**) are indicated. The tower region is shown in the right hand side inset as seen from outside the capsid. **(b)** Sequence alignment of hexon HVRs from HAdV-C5 and HAdV-F41. Grey bars indicate traced regions. The HAdV-C5 acidic region is highlighted by a black rectangle. Amino acids are coloured by polarity (yellow: non-polar; green: polar uncharged; red: polar acidic and blue: polar basic. **(c)** Superposition of the HAdV-C5 penton base monomer (PDB ID: 6B1T Chain M, pink) and HAdV-F41 coloured by RMSD from blue to red and purple as in (a). At the right hand side, panels show the variable loop (top) and fibre binding rearrangement (bottom) regions with the HAdV-F41 map in grey mesh. **(d)** Sequence alignments focusing on the penton base (PB) N-terminal arm, variable and hypervariable loops and hydrophobic rim, and on the fibre N-terminal peptide, as indicated. Black rectangles highlight the RGD/IGDD sequence motifs, the conserved fibre peptide, and the non-conserved Ser426 in the hydrophobic rim. Grey horizontal bars indicate regions traced in the structure. For fibre, the bar indicates traced residues in the HAdV-C5 fibre (PDB ID: 3IZO); suffixes **_l** and **_s** refer to the long and short fibers in HAdV-F41. The histogram indicates the mean pairwise identity over all pairs in the column (green: 100% identity, olive: at least 30% and under 100% identity, red: below 30% identity). **(e)** The penton base pentamer is shown as viewed from outside the capsid, with one monomer in black and the rest in white ribbon. Hydrophobic residues conserved with HAdV-C5 in the fibre binding ring are coloured cyan (Phe412, Tyr413, Tyr419 and Leu422) or maroon (non-conserved, Ser426). **(f)** Penton base pentamer in HAdV-C5 and HAdV-F41 coloured by surface electrostatic potential and viewed from inside the virion, showing the differences in charge in the inner penton cavity.

### Vertex capsomer: penton base

The architecture of the HAdV-F41 penton base protein is very similar to that of HAdV-C5 (Dai *et al*., 2017) (**Table S3, Fig. 2c)**. As in previously solved structures (Liu *et al*., 2010; Yu *et al*., 2017), the first 34 amino acids could not be traced (**Table S5, Fig. 2d**). This region is either flexible, or buried within the noisy, non-icosahedral core density. However, weak density suggests its location in HAdV-F41 (see below, section on additional internal densities). The protein is 63 amino acids shorter than in HAdV-C5 (**Table S2**). This length difference corresponds mainly to a hypervariable loop at the periphery of the pentamer, which in HAdVC-5 bears the integrin-binding RGD sequence motif (**Fig. 2c-d**). All HAdVs bear this RGD motif, except HAdV-F40 and F41, which instead have RGAD and IGDD (Albinsson and Kidd, 1999), and HAdV-D60, which has a deletion of the loop (Robinson *et al*., 2013). The RGD loop is highly flexible, precluding its tracing in any of the available structures. In HAdV-F41 the IGDD-containing region is 57 residues shorter than the HAdV-C5 RGD loop, but could not be traced either (**Table S5, Fig. 2c-d**), indicating that, despite being much shorter, it is also flexible. Lack of the RGD sequence is not a determinant for enteric tropism, as this motif is present in other enteric AdVs such as HAdV-A31 (UniProtKB entry: D0Z5S7_ADE31) and HAdV-G52 (A0MK51_9ADEN). In spite of lacking the RGD motif, it has recently been shown that HAdV-F41 can bind to laminin-binding integrins (Rajan *et al*., 2018). The variable loop (Glu148-Leu156), one of the least conserved regions in the penton base sequences and likewise exposed at the periphery of the pentamer, has also a different conformation in HAdV-F41 (**Fig. 2c-d)**. The role of this loop is unknown, but it has been proposed as a site to be engineered for gene therapy (Zubieta *et al*., 2005).

The N-terminal peptides of AdV fibres (**Fig. 2d**) bind to the groove formed at the interface between monomers in the penton base pentamer. Both penton base and fibre residues involved in this interaction are conserved (Zubieta *et al*., 2005) and we do not observe structural differences in this region. However, we notice a possible difference in the interaction between the start of the fibre shaft and a ring of hydrophobic residues at the centre of the pentamer (Liu *et al*., 2011; Zubieta *et al*., 2005). In this region, fibre sequences are not conserved (**Fig. 2d**). In the penton base of HAdV-F41, the hydrophobic residues forming the ring are conserved, except for HAdV-C5 Phe489 which is instead a polar residue (Ser426) in HAdV-F41 (**Fig. 2d-e**). Next to Ser426, Thr427-Thr430 have a large RMSD from the equivalent residues in HAdV-C5 (**Fig. 2c**). This region forms part of a larger stretch (Tyr419-Leu442) which undergoes a conformational rearrangement upon fibre binding (Zubieta *et al*., 2005). These observations suggest a different fibre-penton binding mode in HAdV-F41, with less hydrophobic and more hydrogen bridging interactions than in HAdV-C5. We also find some differences between HAdV-C5 and HAdV-F41 in the interactions between penton base monomers, and between penton base and the surrounding peripentonal hexons, which are described in **Fig. S4**. Finally, at the pentamer cavity oriented towards the viral core, Arg47 residues in HAdV-C5 form a positively charged ring absent in HAdV-F41, which has Gly45 instead (**Fig. 2f**). This charge variation suggests a different interaction between penton base and the viral genome, or its packaging machinery.

### Proteins IIIa and VIII

Minor coat proteins IIIa and VIII line the HAdV-F41 internal capsid surface. Five copies of protein IIIa are located beneath each vertex, bridging the penton and the peripentonal hexons (**Fig. 1b**), and presumably interacting with packaging proteins during assembly (Condezo *et al*., 2015; Ma and Hearing, 2011). The extent of protein IIIa traced and its architecture are very similar to those of HAdV-C5 (**Tables S3, S5**), with the domain organization previously described (**Fig. 3a-b**) (Liu *et al*., 2010). HAdV-F41 protein IIIa has an eight amino acid insertion at the N-terminus with predicted α-helical structure (residues 2-9, **Fig. 3c**), which could form extra intramolecular interactions in the peripentonal region and contribute to make the vertex region more stable. However, we did not observe interpretable density for this insertion, or for the appendage domain (APD) traced in HAdV-D26 (Yu *et al*., 2017) (**Fig. 3c**). The largest difference with HAdV-C5 corresponds to a kink between the 4^th^ and 5^th^ turns in the helix connecting the GOS-glue to the VIII-binding domain **(Fig. 3a-b)**. This kink may cause differences in the interactions between neighbouring molecules beneath the vertex (see next section).

**Figure 3.**
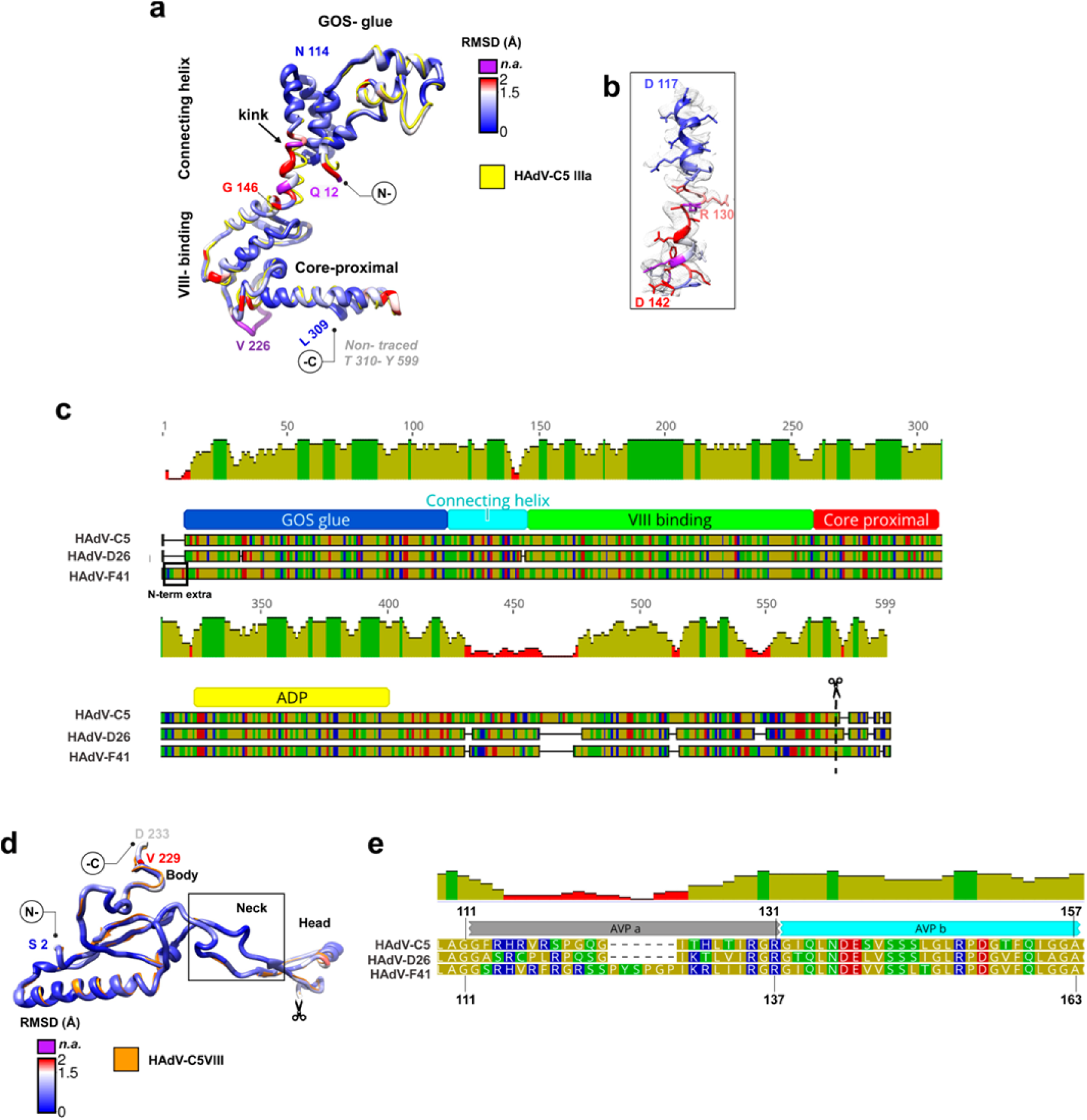
Proteins IIIa and VIII. **(a)** Superposition of HAdV-C5 IIIa (PDB ID: 6B1T, Chain N) in yellow and HAdV-F41 IIIa coloured by RMSD as in Fig. 2. The **GOS-glue, connecting helix, VIII-binding** and **core proximal** domains are indicated. **(b)** Detail of the connecting helix to show its fit into the cryo-EM map. **(c)** Schematics showing the alignment of protein IIIa sequences in HAdV-C5, D26 and F41. Traced domains are indicated below, in different colours. **APD** is the appendage domain traced in HAdV-D26, but not in the other two viruses. The N-terminal extension in HAdV-F41 (**N-term extra**) and maturation cleavage site (**scissors**) are also indicated. **(d)** Superposition of HAdV-C5 protein VIII (PDB ID: 6B1T Chain O) in orange and HAdV-F41 VIII coloured by RMSD as above. The **body, neck** and **head** domains are indicated, as well as the gap corresponding to the peptide cleaved during maturation (**scissors**). **(e)** Sequence alignment showing the two central peptides of protein VIII cleaved by AVP (**AVPa, AVPb**) in HAdV-C5, D26 and F41. Amino acid and mean pairwise identity histogram colour schemes are the same as in Fig. 2.

Two copies of protein VIII are present in each AU of the AdV icosahedral shell. One of them interacts with protein IIIa and contributes to stabilize the vertex, while the second one helps keep the non-peripentonal hexons together in the central plate of the facet (**Fig. 1b**). HAdV-F41 protein VIII is highly conserved compared to HAdV-C5, both in sequence and structure (**Tables S2-S3 and Fig. 3d**). Only the central region of the protein, cleaved by the maturation protease (AVP) (Mangel and San Martín, 2014), presents sequence divergence (**Fig. 3e**, and see next section).

### Proteins VI and VII, and additional internal elements

Analysis of density unoccupied by the molecules traced so far in the HAdV-F41 cryo-EM map (remnant density, RD) revealed several features of interest (**Fig. 4a**). At the rim of the central cavity of the hexon trimers, we observe discontinuous weak density (RD1) that, by similarity with HAdV-C5, corresponds to peptides pVIn and pVIIn_2_, cleaved from precursor proteins pVI and pVII during maturation (Dai *et al*., 2017; Hernando-Perez *et al*., 2020). Since side chains are poorly defined in these regions due to lack of order and low occupancy, we have only tentatively traced four copies of pVIn and two of pVIIn_2_, out of the twelve possible equivalent sites in the AU (**Fig. 1b, Fig. 4b and Table S5**) (Hernando-Perez *et al*., 2020). Protein VI is longer in HAdV-F41 than in HAdV-C5 (**Table S2**) and bears a particular epitope common to enteric AdVs (species A and F) in its central domain (residues 114-125) (Grydsuk *et al*., 1996). However, we do not observe any interpretable density that could inform about either extra interactions between the longer VI chain and hexon, or the central region traced in HAdV-C5 (Dai *et al*., 2017).

**Figure 4.**
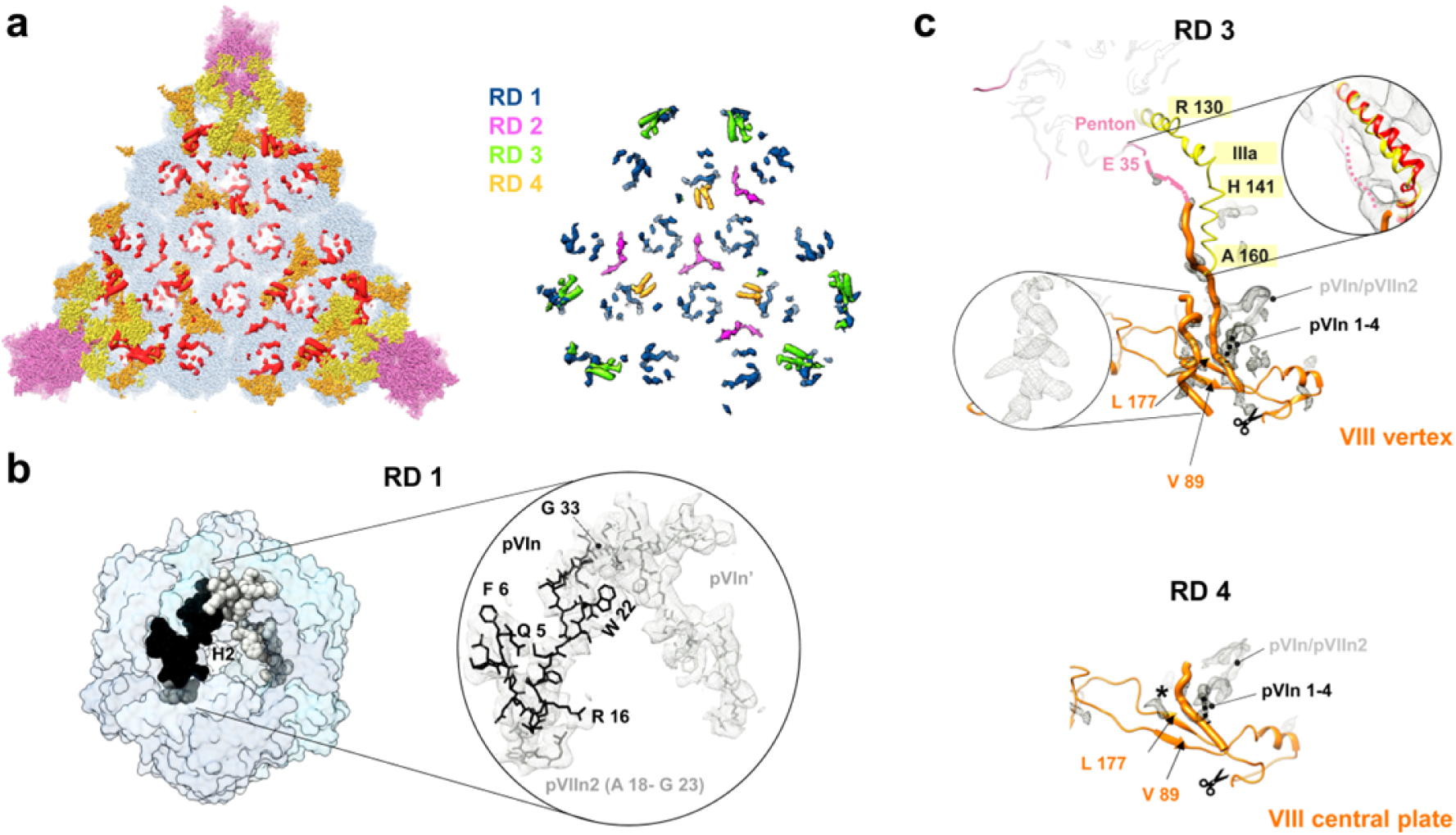
Additional internal components of the HAdV-F41 capsid. **(a)** The left hand side panel shows the remnant densities in one facet (red), not occupied by traced proteins hexon (light blue), penton (light pink), IIIa (yellow) or VIII (orange). In the right hand side panel, components of the remnant map have been classified as RD1-4 (see text) and coloured as indicated. Densities correspond to the unsharpened map contoured at 1.5s. **(b)** N-terminal peptides of proteins VI (**pVIn**, two copies) and VII (**pVIIn**_**2**_) tentatively traced in RD1, inside the hexon cavity. The surface of one hexon trimer is depicted as seen from inside the capsid. **(c)** Proposed interpretation of remnant densities RD3 and RD4. Except for the inset at the right hand side, remnant density in the sharpened map is shown, contoured at 2.5s. Top, a view of the vertex region seen from inside the capsid showing the location of RD3. Bottom, view from an equivalent point of view of the protein VIII copy located beneath the central plate of hexons. The traced part of the two copies of protein VIII (VIII vertex and VIII central plate) is shown as a thin ribbon, and the poly-Ala peptides modelled in the unassigned densities proposed to correspond to the central peptide of VIII are in orange thick worms. The inset at the left shows the possibly α-helical shape of one of the remnant densities in RD3. This density is much weaker in RD4, where we have placed a black star to indicate its position. Scissors indicate the AVP cleavage site in protein VIII. Peptide pVIn is represented as a grey ribbon, with the possible location of its four untraced N-terminal residues as a black dotted line. In the RD3 panel, the IIIa connecting helix is in yellow, the first residue traced in the penton base protein (E35) is colored pink, and a dotted pink line indicates the possible trajectory of the untraced 34 residues. In the framed zoom at the right hand side, the IIIa connecting helix is shown overlapped with the HAdV-C5 helix in red, and the unsharpened map contoured at 1s is shown as a grey mesh, to emphasize low resolution density proposed to correspond to the penton base N-terminal arm.

L-shaped densities at the icosahedral 3-fold (I3) axis and the local 3-fold (L3) axis between hexons 2, 3 and 4 (**Fig. 4a**, RD2) were also observed in HAdV-D26, but their assignment is uncertain (Yu *et al*., 2017). In HAdV-F41, we observe two additional RDs that were not reported in previous studies (Dai *et al*., 2017; Yu *et al*., 2017). RD3 and RD4 are located near the two independent copies of protein VIII and have similar shapes, but RD3 (located beneath the penton region) has stronger density. Both RD3 and RD4 are near the gap in protein VIII left by AVP cleavage during maturation (**Fig. 3d-e**, and **Fig. 4a, c**). In RD3, we could model two poly-Ala peptides. One of them could correspond to a 21 residue α-helix, while the other one is an extended 23 residue peptide (**Fig. 4c, top**). These lengths correlate well with those of the pVIII peptides cleaved by AVP (**Fig. 3e**). In HAdV-F41, one of the cleaved peptides is 6 residues longer than in HAdV-C5 and HAdV-D26, has low sequence conservation (**Fig. 3e**), and secondary structure predictions suggest that, unlike in HAdV-C5, it may have some propensity to form an α-helix, although with low confidence (**Fig. S5**). In RD4, the density is weaker and only a 10 residue extended peptide could be modelled (**Fig. 4c, bottom**). We hypothesize that at least part of RD3 and RD4 correspond to these excised peptides of protein VIII, which in HAdV-F41 would be ordered and reinforcing the network of contacts on the inner capsid surface (particularly beneath the vertex), therefore contributing to a higher capsid stability. RD3 and RD4 also connect to density corresponding to pVIn, and could account for its untraced first four residues (**Table S5** and **Fig. 4c, black dotted lines**). Fragmented density in RD3, which is not present in RD4, seems to connect to the first traced residue in penton base, Glu35, and runs parallel to the IIIa connecting helix. We propose that this density corresponds to the untraced N-terminus of penton base (**Fig. 4c, pink dotted line**), that would interact with the kinked connecting helix in IIIa, a contact which would not happen in HAdV-C5, where the helix is not kinked (**Fig. 3a)** and the sequence of the penton base N-terminal sequence differs (**Fig. 2d)**.

Determining the identity of weak, discontinuous density patches in AdV is problematic, as there are many virion components that do not follow icosahedral symmetry (packaging proteins, protein VI, half of protein IIIa, cleaved peptides, core proteins), making the combinatorial problem of sequence assignment unsolvable (Condezo *et al*., 2015; Hernando-Perez *et al*., 2020; San Martín, 2012). Nevertheless, our interpretation suggests that the central peptides of pVIII, the N-terminus of penton base, and the kinked IIIa helix would collaborate to strengthen the HAdV-F41 vertex region in a manner different to that previously observed in other HAdVs.

### External cementing network: protein IX

Previous studies (Liu *et al*., 2010; Yu *et al*., 2017) have shown that each of the twelve monomers of protein IX in a facet of the AdV capsid presents an extended conformation and is composed of three domains. The N-terminal domains of three IX molecules associate in a triskelion-shaped feature. One of the four triskelions in each facet occupies the valley between hexons at the I3 axis, while the other three lay at the L3 axes formed by hexons 2, 3 and 4 in the AU (**Fig. 1b and Fig. 5a**). The central domain of protein IX is highly flexible, and has also been termed “rope” domain. In HAdV-C5 and D26, the rope domains of three conformationally unique IX molecules, each one originating in a different triskelion, crawl around the hexons on the surface of the icosahedral facet until they reach the edge. There, the C-terminal domains of these three molecules join a fourth one coming from the neighbouring facet to form a coiled coil with three parallel and one anti-parallel α-helices. There are two of these four-helix bundles per facet edge **(Fig. 5a)**. In non-human mastadenoviruses, the rope domain is shorter (**Fig. 5b**), and the C-terminal domains of IX form coiled coils with only three parallel α-helices located directly on top of their N-terminal triskelions (four such bundles per facet), as exemplified in the bovine adenovirus BAdV-3 structure (**Fig. 5a**) (Cheng *et al*., 2014; Hackenbrack *et al*., 2016; Reddy, 2017; Schoehn *et al*., 2008). Of all HAdV-F41 structural proteins, protein IX is the least similar to its HAdV-C5 counterpart (**Table S2**). We find that this protein also displays a conformation different from all previously reported mastadenovirus structures.

**Figure 5.**
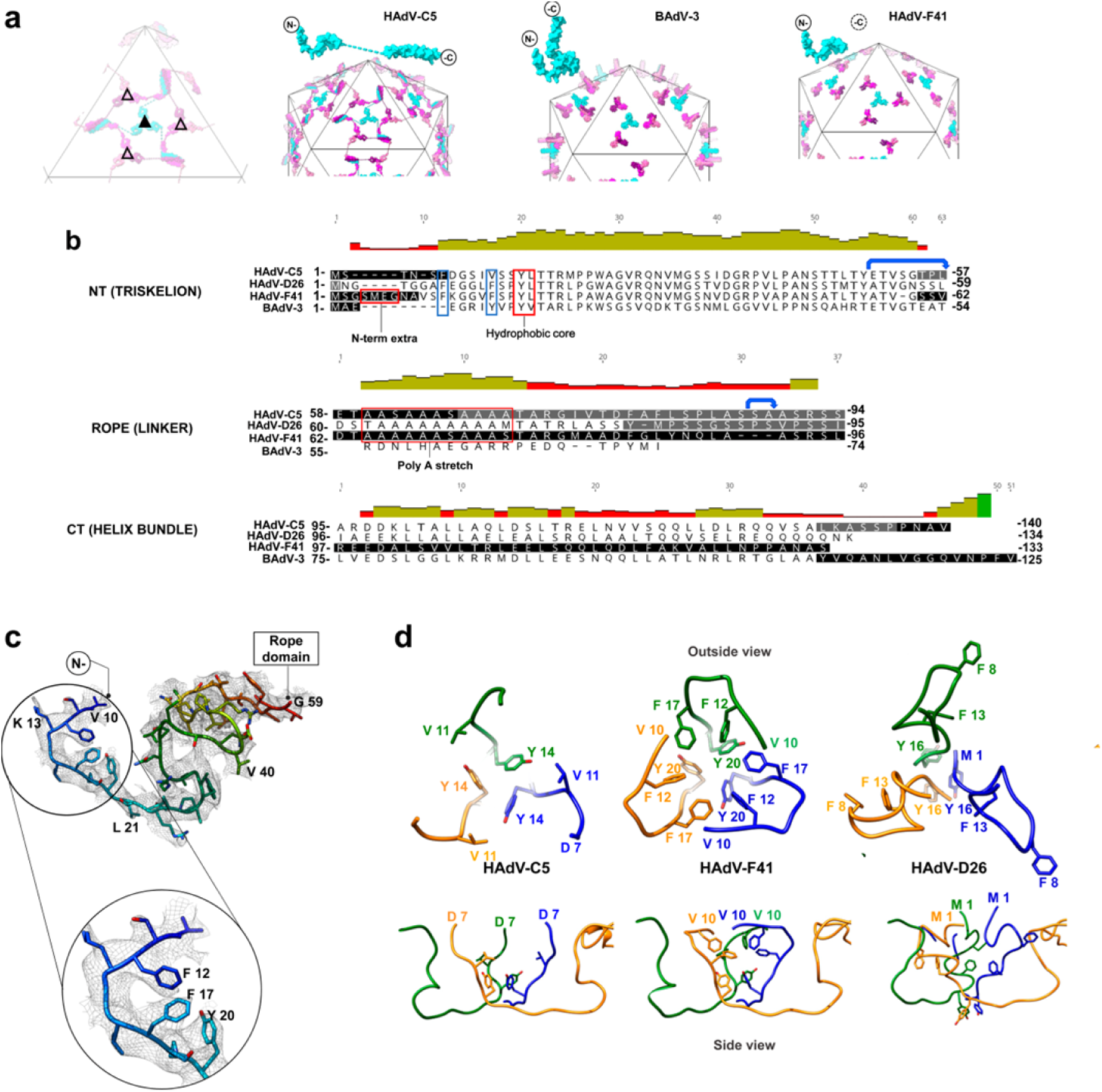
Protein IX triskelion structure. **(a)** Schematics showing the organization of protein IX in HAdV-C5, BAdV-3 and HAdV-F41, where only the N-terminal domain is ordered. Triskelions located at the I3 symmetry axis (filled triangle) are depicted in cyan, and those located at the L3 axis (empty triangle) in several shades of pink. The rope domains of HAdV-C5 (dashed lines indicating non-modelled residues) crawl around the hexons on the surface until they reach the edge, where they form 4-helix bundles. In contrast, BAdV-3 rope domains fold back and form 3-helix bundles directly on top of their triskelions. One IX monomer is depicted in cyan on top of each schematic, with the N and C termini indicated. **(b)** Structure-based sequence alignment of protein IX in HAdV-C5, HAdV-D26, HAdV-F41 and BAdV-3. Regions spanning the amino terminal (**NT**), **rope** and carboxy terminal (**CT**) domains are indicated. Black text corresponds to residues modelled in the different structures. White text on black background corresponds to regions not modelled. On grey background, regions modelled, but not in all independent copies of IX. The N-terminal extra residues in HAdV-F41, the triskelion hydrophobic core and the poly-Ala stretch (absent in BAdV-3) are indicated with red boxes. Blue boxes highlight the two Phe residues discussed in the text. The two flexible bends are indicated with blue arrows. **(c)** N-terminal domain of HAdV-F41 protein IX rainbow coloured from blue (N-term) to red (last traced residue), with the density map in grey mesh, and a zoom into the hydrophobic residues discussed in the text. **(d)** Comparison of the triskelions in HAdV-C5 (PDB ID: 6B1T), HAdV-F41 and HAdV-D26 (PDB ID: 5TX1). Top row: triskelions as seen from outside the capsid. Bottom: a view rotated by 90°, so that the hexon shell surface would be located at the bottom. Only the first residue traced for each virus is labelled in this view.

We have traced residues Val10 to Gly59 (**Fig. 5c**), which form the triskelion in a manner very similar to that of HAdV-C5. As previously reported (Liu *et al*., 2010), the triskelion is underpinned by a core of hydrophobic residues, Tyr20-Leu21 in HAdV-F41 (**Fig. 5b, d**). The first residues traced fold over the triskelion centre. Notably, this arrangement implies that Phe12 and Phe17 add two tiers of hydrophobic interactions to the triskelion core (**Fig. 5d**). In HAdV-C5, one of the two phenylalanines is absent (Val11, **Fig. 5b**), and the region containing the other (Phe6) could not be traced, implying lack of icosahedral order (**Fig. 5d)**. Both phenylalanine residues (Phe8 and Phe13, **Fig. 5b**) are modelled in HAdV-D26 protein IX, but in a different conformation, oriented outwards from the triskelion core (**Fig. 5d**).

Protein IX in HAdV-F41 has a five-residue insertion at the N-terminus when compared to HAdV-C5 (four-residue compared to HAdV-D26). The N-terminal domain is even shorter in BAdV-3 (**Fig. 5b**). We hypothesize that, in HAdV-F41, the region containing the two phenylalanine residues is more ordered than in HAdV-C5, and in a different conformation from HAdV-D26, due to the presence of these extra residues. Weak density capping the triskelion at the I3 axis could account for the presence of the longer N-terminus (**Fig. 6a-b**). Both the dense network of hydrophobic residues at the triskelion core and the presence of extra residues at the N-terminus of IX would reinforce the intermolecular interactions within the triskelions, likely contributing to stabilize the protein IX trimer. It has previously been shown that the triskelion is sufficient to provide capsid thermostability in HAdV-C5 (Vellinga *et al*., 2005). More stable triskelions may enhance the physicochemical stability of HAdV-F41.

**Figure 6.**
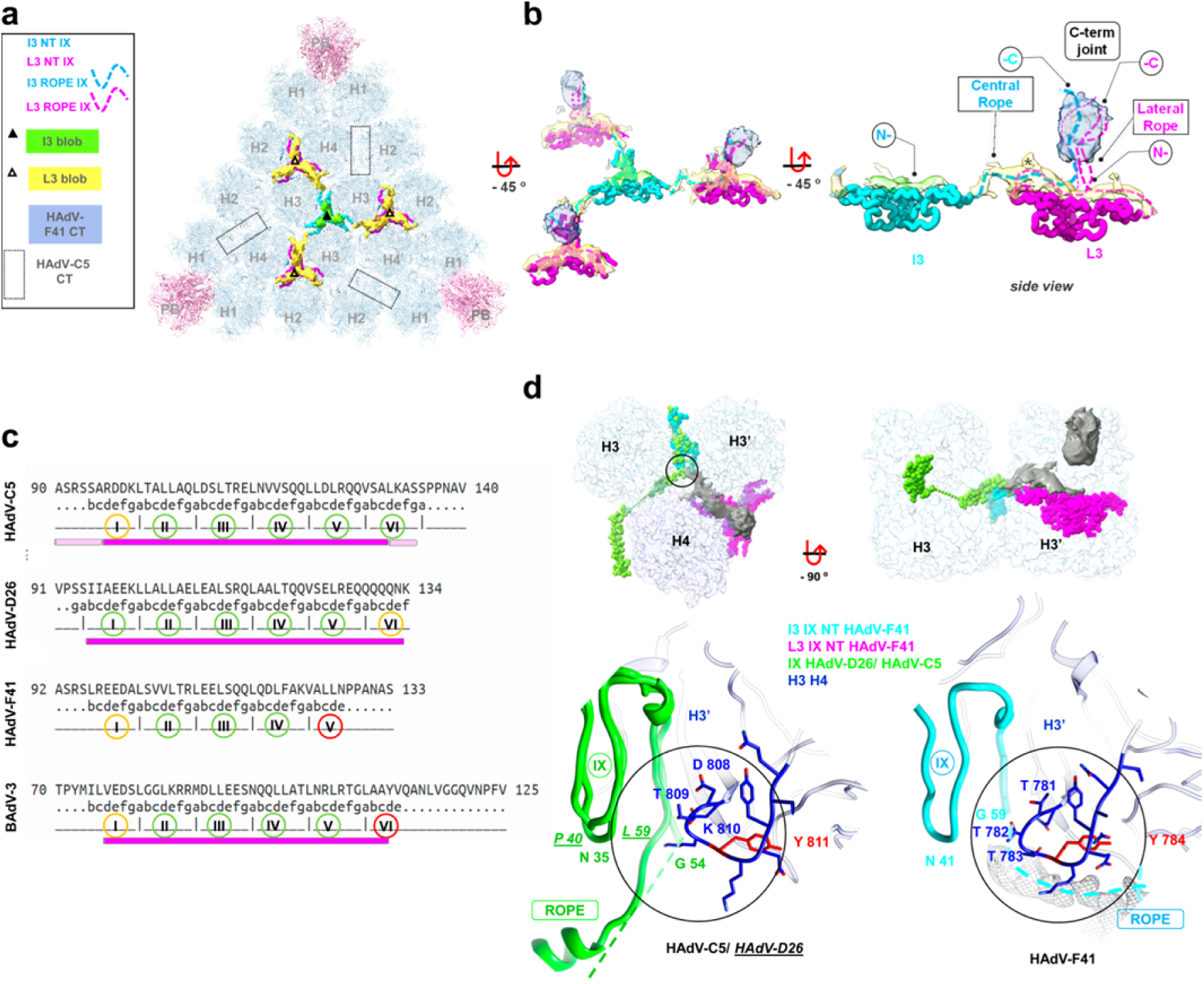
Organization of protein IX in the HAdV-F41 capsid. **(a)** Remnant density map as seen from outside the capsid. Penton (**PB**) and hexon (**H1-H4**) models in one facet are shown in pale colours. The modelled triskelions are shown in cyan (triskelion at the I3 symmetry axis, **I3 NT**) and magenta (triskelions at the L3 symmetry axes, **L3 NT**). Remnant unassigned density corresponding to the unsharpened map contoured at 0.2s is coloured green and yellow. The location of helix bundles in HAdV-C5 is indicated with dotted rectangles. **(b)** Interpretation of the remnant map. Apart from the elements shown in (a), the density protruding at the L3 axes is shown in blue, and the proposed path for the rope domains corresponding to each triskelion is in cyan and magenta dashed lines. Note that the green density would only harbour the I3 extra N-terminal residues, whereas the yellow density would accommodate both the extra N-term of the L3 triskelions and all rope domains. The star indicates where the I3 molecule would join the L3 one. **(c)** Heptad repeat prediction for the C-terminal domain of protein IX in HAdV-C5, HAdV-D26, HAdV-F41, and BAdV-3. For each virus, the first line shows the protein sequence, and the second shows the predicted coiled-coil regions whose amino acids are classified as a heptad pattern (abcdefg). Absence of coiled-coil is denoted as a dot. In the third line, heptads are numbered with roman numbers within a green (complete repeat), yellow (almost complete) or red (incomplete) circle. The fourth row represents α-helices traced in the structures, in magenta if the region is modelled for all copies of IX in the AU, and in light pink if it is not. Note that HAdV-F41 predicted coiled-coil region is the shortest, with only three complete repeats. **(d)** The rope domain of the IX molecules forming the I3 triskelion follows different paths in HAdV-C5/D26 and HAdV-F41. Top: one of the IX monomers forming the HAdV-D26 triskelion at the I3 axis is shown in green; the equivalent monomer in HAdV-F41 is in cyan; and the HAdV-F41 L3 triskelion is shown in magenta. Surrounding hexons (H3, H3’ and H4) are depicted in transparent blue, and the HAdV-F41 remnant map is in grey surface. The view in the left hand side panel is as seen from outside the capsid. For clarity, H4 is not depicted in the right hand side view. Bottom: zoom in on the region where the triskelion ends and the rope domain turns. The structures of IX in HAdV-C5 and HAdV-D26 are overlapped and depicted in green. HAdV-D26 residue labels are <u>underlined</u>. The rope domains in HAdV-C5 and HAdV-F41 are not traced, but are represented as dashed lines. Remnant density of HAdV-F41 corresponding to its rope domain is depicted as a grey mesh. The hexon (H3) amino acids with RMSD above 2 Å are depicted in red. Note the threonine triplet in the case of HAdV-F41 (781-783), absent in HAdV-C5 and D26.

The flexibility of the rope domain makes it difficult to observe: only one of the four copies per AU has been traced for HAdV-D26 or HAdV-C5 (only partially for the latter) (Dai *et al*., 2017; Yu *et al*., 2017). We do not observe density with enough definition to trace any of the rope domains in HAdV-F41. Surprisingly, in our map there is no density corresponding to the C-terminal 4-helix bundle at the icosahedron edges, although secondary structure prediction indicates a helical propensity. Instead, we observe blurry density protruding in a radial orientation between the towers of hexons 2, 3 and 4, on top of the triskelions at the L3 axis (**Fig. 6b** and **S6**). The presence of this protrusion would suggest that in HAdV-F1 the protein IX C-terminal domains could be arranged in a similar way to the shorter protein IX in BAdV-3, forming a three-helix bundle directly adjacent to each triskelion (**Fig. 5a**). However, in our HAdV-F41 map there is no equivalent weak density near the triskelion at the 3-fold icosahedral axis (**Fig. 6b** and **S6**), indicating that the organization of IX is not the same as in BAdV-3. Focused classification did not yield any subset of particles with a protrusion at the I3 axis (**Fig. S7**). Therefore, the arrangement of the C-terminal domain of HAdV-F41 protein IX does not seem to follow either the typical human or the non-human AdV architectures previously observed.

In both human and non-human AdVs with solved structures, heptad repeats are involved in the formation of protein IX C-terminal helix bundles (Cheng *et al*., 2014; Liu *et al*., 2010; Yu *et al*., 2017). While HAdV-C5 and D26 have five complete heptad repeats and BAdV-3 has four, prediction algorithms find only 3 complete repeats in the C-terminal sequence of HAdV-F41 protein IX (**Fig. 6c**). The shorter series of heptad repeats hinders the formation of a stable, ordered helix bundle in HAdV-F41 (**Fig. S8**). An intriguing question is why the helix bundle, even if disordered, would form at the L3 axis, and not at the facet edge as in the other HAdVs. Sequence alignment indicates that the rope domain in HAdV-F41 is 3 residues shorter than in HAdV-C5 and D26 (**Fig. 5b**). This difference in length, together with the shorter heptad repeat region, may be responsible for hindering interaction of the three parallel α-helices originating from one facet with the fourth one, coming in anti-parallel orientation from the adjacent facet. This consideration would explain why the four-helix bundle is not formed at the facet edges, but the question why there is no density at the centre of the facet that could correspond to a disordered three-helix bundle remains.

A remnant map showing weak density on the capsid surface provides a possible explanation for this unusual arrangement. At low threshold, we observe elongated density connecting one arm of the triskelions at the L3 axes with the triskelion at the centre of each facet (**Fig. 6a-b**). Extra density at the other two arms connects only the end of the modelled region to the centre of the same triskelion (**Fig. 6b**). We interpret that the elongated density corresponds to the rope domain of the central triskelion, which as in HAdV-C5 runs on the capsid surface towards the facet periphery. At the local-3fold triskelions however, lack of partner molecules with which to form a stable helix bundle at the icosahedron edges results in the rope domain turning back on itself. In this situation, the C-terminal regions of the three IX copies at the L3 triskelion would be joined by one copy coming from the facet centre. The flexibility and length of the rope domains, together with the extra N-terminal disordered residues capping all triskelions, and defective heptad repeats, would preclude formation of a stable, ordered helix bundle, producing weak density only at the peripheral triskelions in each facet (**Fig. 6b**).

According to our interpretation, in HAdV-F41 the rope domain would make a counter clockwise turn at the exit of the central triskelion, instead of clockwise as in HAdV-C5 and D26 (**Fig. 6d**). It has been proposed that protein IX in HAdV has two flexible bends that facilitate its coupling to the hexon contours (**Fig. 5b**) (Reddy, 2017). The different path followed by IX in our HAdV-F41 model may be determined by differences in the hexon residues interacting with IX near the first bend. In particular, Tyr784 in hexon 3, located near the end of the triskelion (residues 56-59), is one of the few residues presenting high RMSD when comparing the HAdV-F41 and HAdV-C5 hexon structures (**Fig. 2a** and **Fig. 6d**). HAdV-C5 also has a tyrosine in this position, but the upstream sequence is very different in both viruses, with a conspicuous threonine triplet in HAdV-F41 (residues 781-783); and protein IX has a one amino acid deletion at this bend (**Fig. 5b**). All these changes may result in different interactions causing protein IX to bend in a different direction in both viruses.

## Discussion

The narrow enteric tropism of HAdV species F is not well understood. Infectivity analyses have shown that, unlike HAdV-C5, these viruses are not inactivated by acidic environments (Favier *et al*., 2004), and a main player conferring this resistance to low pH seems to be the short fibre, one of the differentiating structural features of HAdV-F (Rodriguez *et al*., 2013). Here we show that HAdV-F41 capsids are more thermostable than HAdV-C5, requiring higher temperatures to open up and expose its genome to the solvent; and that infectivity is stable, even activated, in simulated gastric and intestinal conditions. We have solved the structure of the HAdV-F41 capsid at near atomic resolution, and analysed it looking for possible determinants of its enhanced physicochemical stability.

The exact molecular basis of virus capsid stability is difficult to unravel, and is still a subject of intense investigation even for so-called simple viruses (Mateu, 2013). We describe some intermolecular contacts that could have a different nature in HAdV-F41 and HAdV-C5, but it is difficult to conclude if these differences will have an effect in capsid stability, due to the large complexity and sheer number of interactions present in AdV virions. However, we do find partially ordered features that suggest collaboration between the N-terminus of penton base, protein IIIa, pVIn and the cleaved peptides of VIII in stabilizing the vertex region. On the outer capsid surface, the protein IX triskelion has a stronger hydrophobic core than in HAdV-C5, while the rest of the protein is disordered, notably lacking the four-helix bundle at its C-terminus. It has previously been observed that the four-helix bundle in HAdV-C5 is easily disturbed, becoming disordered when protein IX was modified by a C-terminal fusion (Marsh *et al*., 2006) or antibody labelling (Fabry *et al*., 2009). It is possible that the lack of well-defined density for the C-terminal region of IX in our HAdV-F41 map is simply caused by partial disruption accidentally occurring during sample preparation. However, in studies where the HAdV-C5 helix bundle was not clearly visible, no weak density was observed in other capsid locations. Differences in sequence and weak densities in the map support the presence of an organization for protein IX in HAdV-F41 different from all described for other AdVs. Our observations also imply that the formation of a well ordered helix bundle is not required to enhance capsid stability, reinforcing the idea that the triskelion is the crucial part for the cementing action of the protein (Vellinga *et al*., 2005). The more exposed location of the protein IX C-terminus in HAdV-F41 makes it an interesting locale for exogenous peptide insertion for retargeting, epitope display or other biotechnological purposes (Matteson *et al*., 2018).

## Methods

### Virus production

E1-deleted HAdV-F41-EGFP virus (kindly provided by Douglas Brough, GenVec/Precigen) was propagated in 2V6.11 cells (ATCC JHU67) with E4 orf6 protein induced with 1μg/ml ponasterone A 24 hours prior to infection. Cells were infected at an input multiplicity of infection (MOI) <0.1 fluorescent units (FU) per cell, and cultures were incubated until cytopathic effect (CPE) was complete or almost complete but before cells showed signs of disintegration. Flasks were hit sharply to dislodge infected cells from the surface and cells were harvested by centrifugation at 1600 x *g* for 10 minutes, washed with PBS, then re-suspended in a small volume of serum-free medium. Any cells remaining attached to the flask were re-fed with a 1:1 mixture of fresh medium and clarified medium from the infected flasks. If uninfected cells had become confluent, they were sub-cultured to allow for virus spread. The process was continued until CPE was complete, typically about 1-2 weeks post infection (p.i.). Progeny virus was released by five cycles of freezing-thawing and the cell lysate was clarified by centrifugation at 1600 x *g* for 10 minutes. Viruses were used in experiments at passage level 3.

Virus was purified by two cycles of caesium chloride (CsCl) density gradient centrifugation. Clarified cell lysate (∼4.5ml) was underlaid with 4 ml CsCl (1.2 g/ml in 50mM Tris-HCl, pH 8.1) and 2 ml CsCl (1.4g/ml in 50mM Tris-HCl, pH 8.1). The gradient was centrifuged in a SW41 rotor at 120 000 x g at 4°C for 1 hour. Virions were collected from the interface of each step gradient and diluted in 50mM Tris-HCl to a density <1.2g/ml, as determined by measuring refractive index, and layered on top of a preformed gradient (1.2-1.4 g/ml) that was prepared using the Gradient Master (BioComp). The gradient was centrifuged in a SW41 rotor at 120,000 x *g* at 4°C for 2 hours. The viral band of mature virus particles was recovered and dialyzed at 4°C against 3 changes of storage buffer (50mM Tris-HCl, pH 8.1, 150mM NaCl, 10mM MgCl_2_ and 10% glycerol) with a buffer change every 45 minutes. Optical density was measured with a NanoDrop® ND-1000 Spectrophotometer (v3.3) and virion concentration was calculated using the equation: 1A_260_ = 1.1 × 10^12^ particles/ml (Maizel *et al*., 1968). Infectivity was determined by endpoint dilution assays with HEK 293 cells, in duplicate, in 60-well Terasaki plates as previously described (Brown, 1985). Assays were scored for the presence of green cells (expressing EGFP), and titres, calculated by the Reed and Muench method (Reed and Muench, 1938), were expressed as fluorescent units (FU)/ml. Purified HAdV-F41 particles from two different preparations were pooled and concentrated by centrifugation in a CsCl gradient and stored in 20mM HEPES pH 7.8, 150 mM NaCl, 10% glycerol at −80°C.

HAdV-C5 used as a control for extrinsic fluorescence experiments was the E1-E3 deleted, GFP expressing Ad5/attP vector which is structurally wildtype (Alba *et al*., 2007), propagated at 37°C in HEK293 cells, harvested at 36 h post-infection and purified as described (Hernando-Perez *et al*., 2020).

### Infectivity assays under simulated gastrointestinal conditions

HEK293 cells were seeded in 6-well plates at a density of 5×10^5^ cells/well one day prior to infection. HAdV-41-EGFP preparations (∼2 × 10^11^ particles/ml, 5 × 10^6^ FU/ml) were dialyzed against serum-free medium and frozen for a short period. Virus (50 µl) was then mixed 1:1 with 2x synthetic gastric fluid (SG) according to the US Pharmacopoeia at 0.01N HCl (4.0 g NaCl and 6.4 g pepsin (Sigma) dissolved in 14 ml of 1N HCl, made up to 1 liter with ddH_2_O, pH 1.2) and/or 2x synthetic intestinal fluid (SI −13.6 g of monobasic potassium phosphate in 250 ml ddH_2_O, 77 ml 0.4N NaOH, 500 ml ddH_2_O, then 20.0g pancreatin (Sigma), adjusted to pH 6.8, then diluted with water to 1 liter). Virus suspension was incubated for 1 hour at 37°C. For sequential treatments, virus was treated with synthetic gastric fluid for 1 hour at 37°C, then mixed 1:1 with synthetic intestinal fluid and incubated for an additional hour. To neutralize pH and proteases, complete MEM containing 10% FBS was added to final volume of 0.3 ml. HEK293 cells were infected with treated or mock treated virus for 1 hour at 37°C. Inoculum was removed, cells were harvested 19-21 hours post infection, and fixed with 2% PFA for flow cytometry. GFP expression was analysed with an LSR Fortessa (BD Biosciences) and data was analysed using FACSDiva (BD Biosciences) gating out doublets. 10 000 cells were counted per well. Each experiment was done in duplicate.

### Extrinsic fluorescence thermostability assays

The data for HAdV-C5 have been reported previously (Hernando-Perez *et al*., 2020). The data for HAdV-F41 were collected in the same time period, as part of a large study comparing the thermostability of different AdV specimens. HAdV-C5 and F41 samples (5×10^9^ virus particles in a final volume of 800 µl of buffer) were incubated overnight at 4 °C in 8 mM Na_2_HPO_4_, 2 mM KH_2_PO_4_, 150 mM NaCl, and 0.1 mM EDTA, pH 7.4. Fluorescence emission spectra from virus samples mixed with YOYO-1 were obtained every 2 °C along a temperature range from 20 to 70 °C in a Horiba FluoroLog spectrophotometer. The dye was excited at 490 nm and maximal emission intensity was achieved at 509 nm. Raw spectra were corrected by subtraction of the YOYO-1 spectrum in buffer at each tested temperature. A total of six (HAdV-C5) and four (HAdV-F41) independent experiments were considered for each averaged curve. The half transition temperatures (T_0.5_) were calculated from the fitting of the averaged fluorescence intensity curve as a function of temperature to a Boltzmann sigmoid equation as described (Hernando-Perez *et al*., 2020).

### Cryo-EM sample preparation and data collection

Pooled and concentrated HAdV-F41 samples (as described above) were dialyzed against 50 mM Tris-HCl pH 7.8, 150 mM NaCl, 10mM MgCl_2_ for 1 hour at 4°C and further concentrated by spinning in a Microcon YM-100 device for 15 min at 4°C, for a final estimated concentration of 3.1×10^12^ vp/ml. Samples were deposited in glow discharged, Quantifoil R2/4 300 mesh Cu/Rh grids and vitrified in liquid ethane after manual blotting in a Leica CPC device. Cryo-EM images (**Table S1**) were recorded at the CEITEC facility (Brno, Czech Republic) using a 300 kV Titan Krios microscope equipped with a Falcon II detector, with a total dose of 42 e-/Å^2^ distributed over 25 frames, at nominal pixel size 1.38 Å and defocus range between −1 and −2.5 µm.

### Cryo-EM data analysis

All image processing and 3D reconstruction tasks were performed within the Scipion framework (de la Rosa-Trevin *et al*., 2016). Frames 2-24 of each movie were aligned using whole-image motion correction implemented in Xmipp, followed by correction of local movements using Optical Flow (Abrishami *et al*., 2015). The contrast transfer function (CTF) was estimated using CTFFIND4 (Rohou and Grigorieff, 2015). Particles were semi-automatically picked from micrographs corrected for the phase oscillations of the CTF (phase-flipped), extracted into 780×780 pixel boxes, normalized and downsampled by a factor of 2, using Xmipp (de la Rosa-Trevin *et al*., 2013). All 2D and 3D classifications and refinements were performed using RELION (Scheres, 2012). 2D classification was used to discard low quality particles, and run for 25 iterations, with 50 classes, angular sampling 5 and regularization parameter T = 2. Classification in 3D was run for 40 iterations, with 3 classes, starting with an angular sampling of 3.7 degrees and sequentially decreasing to 0.5, and regularization parameter T = 4. Icosahedral symmetry was imposed throughout the refinement process. The initial reference for 3D classification was a non-human adenovirus cryo-EM map (unpublished), low-pass filtered to 60 Å resolution. The class yielding the best resolution was individually refined using the original 780 px boxed particles and the map obtained during the 3D classification as a reference, producing a final map at 4.0 Å resolution, as estimated according to the gold-standard FSC = 0.143 criterion implemented in RELION auto-refine and postprocess routines (Chen *et al*., 2013). A global B-factor was estimated after dividing the map Fourier transform by the modulation transfer function (MTF) of the Falcon II detector. The actual sampling for the map was estimated by comparison with a HAdV-C5 model (PDB ID 6B1T) (Dai *et al*., 2017) in UCSF Chimera (Pettersen *et al*., 2004), yielding a value of 1.36 Å/px.

Focused classification was carried out following the general procedures described in (Ilca *et al*., 2015), using RELION as implemented in Scipion (Abrishami *et al*., 2020). First, a spherical mask excluding the region of interest was applied to the 3D map obtained from the refinement with icosahedral symmetry enforced. The masked map was projected in the directions determined from the icosahedral refinement, and the projections subtracted from the corresponding experimental images. Then, the regions containing the feature of interest were located by projecting their position in the 3D map, extracted from the subtracted images in small boxes (100×100 pixels) and subjected to 3D classification without orientation change or symmetry enforcement.

### Model building and analysis

Interpretation of the HAdV-F41 3D map was performed using the molecular modelling workflow based on sequence homology in Scipion (Martinez *et al*., 2020). To facilitate chain tracing, we used MonoRes (Vilas *et al*., 2018) and Localdeblur (Ramirez-Aportela *et al*., 2020) to sharpen the map according to the local resolution. The initial model for each polypeptide chain was predicted with Modeller (Webb and Sali, 2016), using as input template the structure of the respective HAdV-C5 homolog chain (PDB ID 6B1T). UCSF Chimera (Pettersen *et al*., 2004) was used to perform a rigid fitting of each chain initial model into the sharpened map. Next, the fitted model of each chain was refined using Coot (Emsley *et al*., 2010) and Phenix *real space refine* (Afonine *et al*., 2018). Validation metrics to assess the quality of the atomic structure were computed with the Phenix *comprehensive validation (cryo-EM)* algorithm. Once we generated the whole structure of the AU, Chimera *findclash* (integrated in the Scipion protocol *chimera-contacts*) was executed to identify possible contacts between pairs of chains contained in the AU or between a chain in the AU and a chain from a neighbouring AU (**Supplementary file S1**). The default parameters, cut-off (−0.4) and allowance (0.0), were used to report as possible bonds all pair of atoms whose distance is no higher than the sum of the corresponding van der Waals radii plus 0.4 Å. To identify additional unmodeled densities, we generated remnant maps with Chimera in two alternative manners: (1) subtracting from the cryo-EM map a map generated with *molmap* around the modelled atoms (resolution 2.0 gridSpacing 1.36); or (2) masking off from the initial map a region within a 3, 4 or 8 Å radius from the modelled atoms (*vol zone invert true*).

Protein sequence alignments were performed with Clustal W2 (http://www.clustal.org/clustal2/) integrated in Geneious version 2019.0 created by Biomatters (https://www.geneious.com) (Larkin *et al*., 2007). For secondary structure prediction we used Jpred4 Server Jnet version 2.3.1 (http://www.compbio.dundee.ac.uk/jpred) (Drozdetskiy *et al*., 2015). The coiled-coil prediction programs Multicoil and Marcoil implemented in Waggawagga server (https://waggawagga.motorprotein.de/) were used to predict heptad repeats and the probability to form coiled coils (Delorenzi and Speed, 2002; Simm *et al*., 2015; Wolf *et al*., 1997). Molecular graphics and analyses including molecule superposition, RMSD calculation, map rendering, map normalization, coloring and measuring surface blobs and surface electrostatic potential rendering were performed with UCSF Chimera and ChimeraX (Goddard *et al*., 2018; Pettersen *et al*., 2004). UCSF Chimera *Hide dust* was used for clarity when composing figures representing density maps.

### Database deposition

The HAdV-F41 cryo-EM map and model are deposited at the Electron Microscopy Data Bank (EMDB, http://www.ebi.ac.uk/pdbe/emdb) and the Protein Data Bank (PDB, http://www.ebi.uk/pdbe) with accession numbers EMD-10768 and 6YBA, respectively. The validation report is included as **Supplementary file S2**.

## Acknowledgements

This work was supported by grants PID2019-104098GB-I00 and BFU2016-74868-P, co-funded by the Spanish State Research Agency and the European Regional Development Fund, as well as BFU2013-41249-P and BIO2015-68990-REDT (the Spanish Adenovirus Network, AdenoNet) from the Spanish Ministry of Economy, Industry and Competitiveness to C.S.M. Work in M.B’s. lab was supported by grant 194562-08 from the Natural Sciences and Engineering Research Council of Canada. M.H.-P. is a recipient of a Juan de la Cierva postdoctoral contract funded by the Spanish State Research Agency. M.P.-I. holds a predoctoral contract from La Caixa Foundation (ID 100010434), under agreement LCF/BQ/SO16/52270032).

We acknowledge Jiri Novacek and Miroslav Peterek at the CEITEC facility (Brno, Czech Republic) for cryo-EM data collection. Access to CEITEC was supported by iNEXT, project number 653706, funded by the Horizon 2020 programme of the European Union. CEITEC Cryo-electron Microscopy and Tomography core facility is supported by MEYS CR (LM2018127). The initial seed for HAdV-F41-EGFP was a gift from Douglas Brough (GenVec/Precigen). We thank Mark J. van Raaij (CNB-CSIC) for careful reading and insightful comments on the manuscript, Margarita Menéndez (IQFR-CSIC), Milagros Castellanos and Luis A. Campos (CNB-CSIC) for advice with fluorescence measurements and analyses, Mara I. Laguna (CNB-CSIC) for expert technical help and José M. Carazo (CNB-CSIC) for constant support.

## Author contributions

M.B., R.M. and C.S.M. designed research. M. P. -I., M. M., G.N.C, M.H.-P., C.M., M.B., R.M. and C.S.M. performed research. M. P. -I., M. M., M.H.-P., C.M., M.B., R.M. and C.S.M. analysed data. M.M. and R.M. contributed analysis tools. M.P.-I. and C.S.M. wrote the paper, with input from all other authors.

## List of supplementary material

Supplementary figures S1 to S8
Supplementary tables S1 to S5
Supplementary files S1 and S2
Supplementary references

