## Supplementary material for "Cryo-EM structure of enteric adenovirus HAdV-F41 highlights structural divergence among human adenoviruses"

[3]Department of Molecular Genetics. University of Toronto. Toronto, Ontario (Canada)

[4]Escuela Politécnica Superior. Universidad Autónoma de Madrid. Madrid (Spain)

**Supplementary Figures**

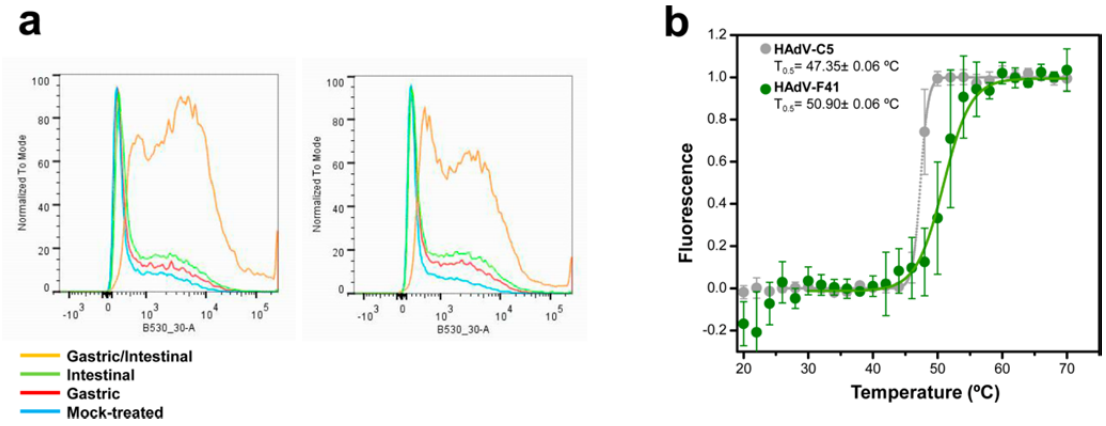


**Figure S1. Physicochemical stability of HAdV-F41 virions.**

**(a)** Representative flow cytometry histograms of HAdV-F41 exposed to gastric, intestinal, or consecutive gastric and intestinal conditions. Mock= serum free medium. Five experiments were performed in duplicate. The increased number of GFP-expressing cells after sequential gastric then intestinal treatment corresponds to a 6.4-fold increase in infectivity, based on the ratio of adsorbed MOI after infection with treated virions to adsorbed MOI after infection with untreated virions. Adsorbed MOI was determined from the proportion of GFP-expressing cells using cumulative Poisson probabilities.

**(b)** Extrinsic fluorescence curve showing DNA exposure to solvent upon temperature increase for HAdV-C5 (grey) and HAdV-F41 (green) particles. Average values and error bars corresponding to the standard deviation of a total of six (HAdV-C5) and four (HAdV-F41) independent experiments are plotted. Dotted lines correspond to the fit according to a Boltzmann sigmoid function.

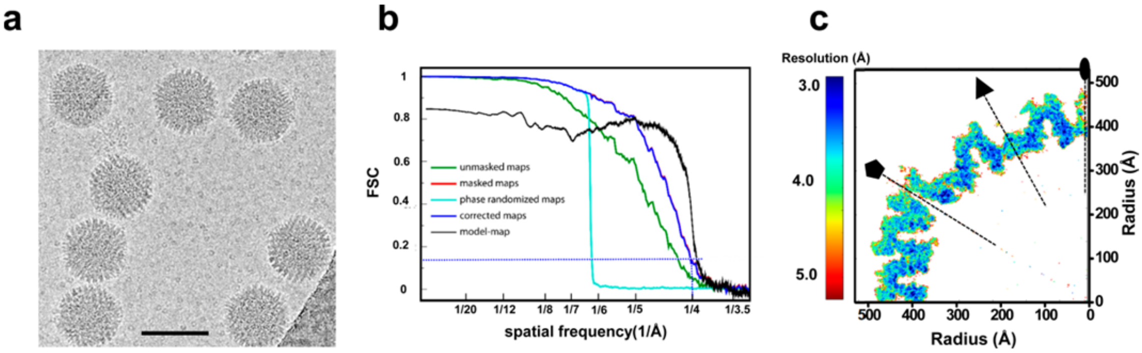


**Figure S2. Cryo-EM data and resolution.**

**(a)** Representative area of a cryo-EM motion-corrected micrograph. The bar represents 100 nm.

**(b)** Fourier Shell Correlation (FSC) curves, as provided by RELION *postprocess*. The FSC= 0.143 threshold criterion indicates an average map resolution of 4.0 Å. The black curve is the model-map FSC calculated by *Phenix*.

**(c)** Central slice of the HAdV-F41 density map viewed along a 2-fold icosahedral axis, coloured according to local resolution as calculated with ResMap (Kucukelbir *et al.*, 2014). The icosahedral 2-fold (oval), 3-fold (triangle), and 5-fold (pentagon) axes are indicated.

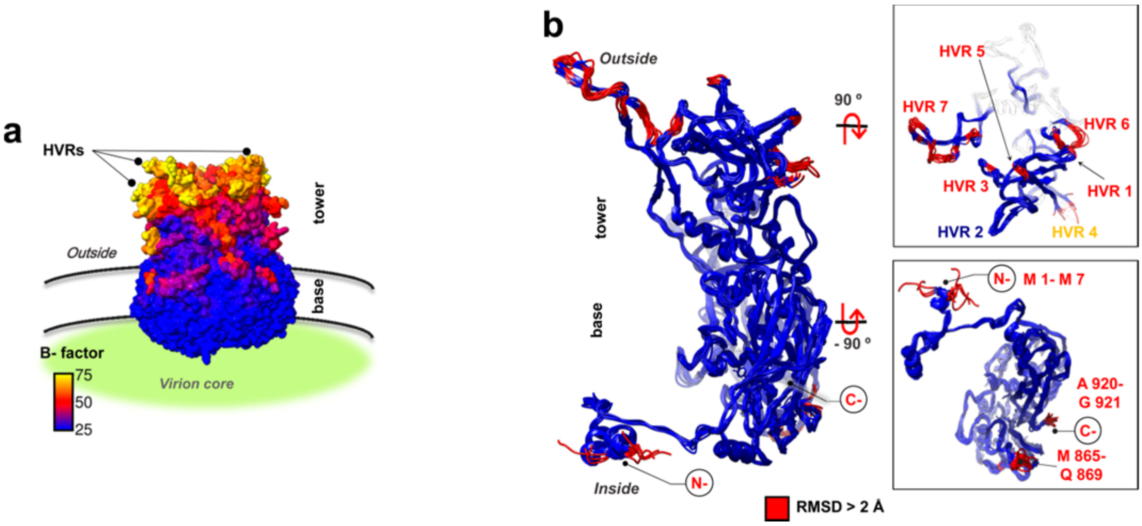


**Figure S3. Mobility among the twelve hexon monomers in the AU.**

**(a)** Hexon trimer coloured by B-factor as estimated by Phenix *real space refine*, emphasizing the hexon regions with lower agreement between map and model.

**(b)** Superposition of the twelve hexon monomers in the AU of HAdV-F41, coloured by RMSD as indicated. At the right hand side, panels show the tower (top) and base (bottom) regions as seen from outside and inside the capsid, respectively. All HVRs except HVR2 display variability among hexon monomers, consistent with their flexibility. As in HAdV-C5 and D26 (Liu *et al.*, 2010; Yu *et al.*, 2017), the N- (Met1-Met7) and C-termini (Ala920-Gly921 followed by four more disordered residues), as well as Met865-Asn869 at the innermost region of the pseudohexagonal base, are also flexible regions that work as molecular switches to fulfil non-equivalent interactions between hexon monomers and different neighbouring partners **(Supplementary file S1)**.

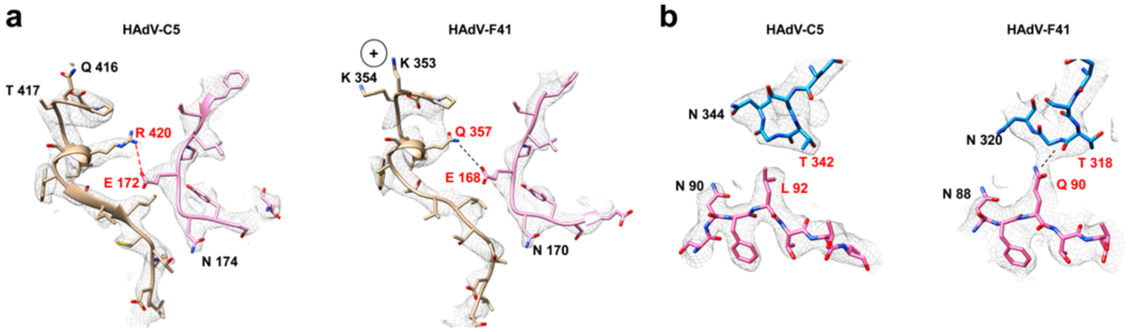


**Figure S4. Differences in penton-penton and penton-hexon contacts between HAdV-F41 and HAdV-C5.**

**(a)** Penton-penton interactions. A possible salt bridge formed by Arg420 and Glu172 between two penton subunits in HAdV-C5 (Zubieta *et al.*, 2005) cannot occur in HAdV-F41, where the arginine residue is substituted by Gln357 which could make a hydrogen bond with Glu168. Density maps are shown in grey mesh. The two penton base subunits are coloured pink and tan. Dashed lines indicate possible interactions (red: salt bridge, black: hydrogen bond).

**(b)** Intermolecular interactions between penton and hexon. In HAdV-F41, Gln90 in penton base (pink) could act as a hydrogen donor, and the O of the Thr318 C-α backbone in hexon (blue) as a hydrogen receptor. In HAdV-C5 (PDB ID: 6B1T, map EMD-7034), Leu92 in penton base is a hydrophobic amino acid that cannot contribute to a hydrogen bond. The potential hydrogen bond between penton base Gln90 and hexon Thr318 would be infrequent in the AdV family. A BLASTP search using HAdV-F41 penton base as query against the NCBI nucleotide collection (nr/nt) database indicated that the conserved amino acid at the Gln90 position for the majority of the available penton base sequences is Leu, which is not a hydrogen donor. The only exceptions were an enteric AdV infecting dolphins (YP_009704126) (Malmberg *et al.*, 2017) and a squirrel AdV (96-29/KOR, ALE33730) which have a Gln like HAdV-F41, and fowl adenovirus A (NP_043882) with Arg, also a hydrogen donor amino acid. This observation suggests that, despite having a conserved architecture, the penton-hexon interaction could be a tropism determinant. However, HAdV-G52 and A31 have enteric tropism and carry the conserved Leu.

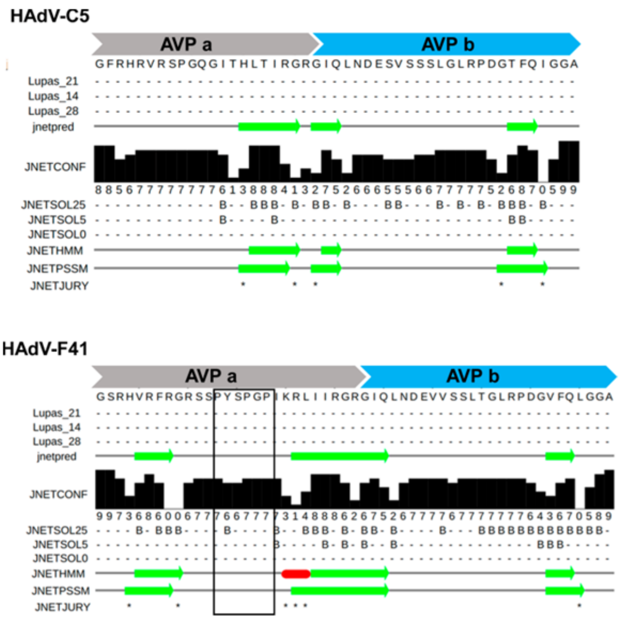


**Figure S5. Secondary structure prediction for the protein VIII peptides cleaved by AVP in HAdV-C5 and HAdV-F41.** The Jpred4 output is shown, with red rectangles indicating α-helices, and green arrows for β-sheets. The two peptides cleaved by AVP are denoted AVPa and AVPb, as in Fig. 3e. The black box highlights the position of the insertion in HAdV-F41 peptide AVPa, which has low sequence similarity with HAdV-C5. Only JNETHMM predicts an α-helix in this peptide for HAdV-F41. No helices are predicted for HAdV-C5.

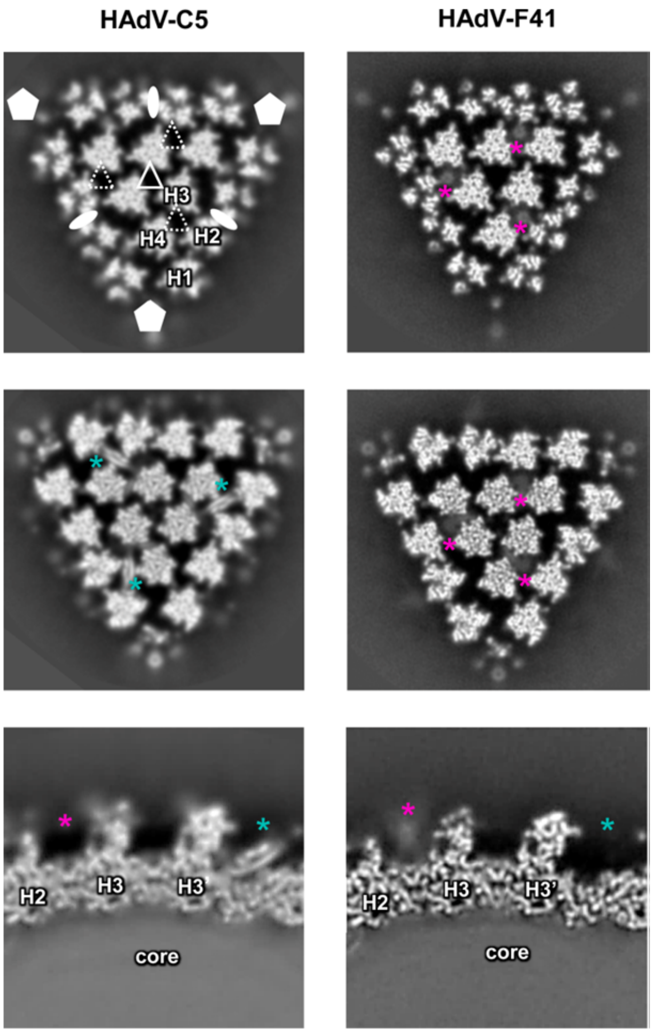


**Figure S6. Comparison between sections in HAdV-C5 and HAdV-F41 cryo-EM maps.** Top two rows: sections at two different heights along the 3-fold icosahedral axis. Bottom row: sections across the capsid. Higher density is shown in white. The HAdV-C5 map is EMD-4448, unsharpened (Hernando-Pérez *et al.*, 2020; Martín-González *et al.*, 2019). Hexons are numbered H1-H4. The white filled pentagons and ovals indicate the 5-fold and 2-fold icosahedral symmetry axes. The white, hollow triangle indicates the icosahedral 3-fold (I3) symmetry axis, and the white broken triangle indicates the local 3-fold (L3) symmetry axis between hexons 2, 3 and 4. Maps are filtered at a similar resolution (6.1 Å). Turquoise stars indicate the position of the protein IX C-terminal helix bundle in HAdV-C5. Pink stars indicate the position of weak protrusions at the local, but not at the icosahedral, 3-fold axes in HAdV-F41.

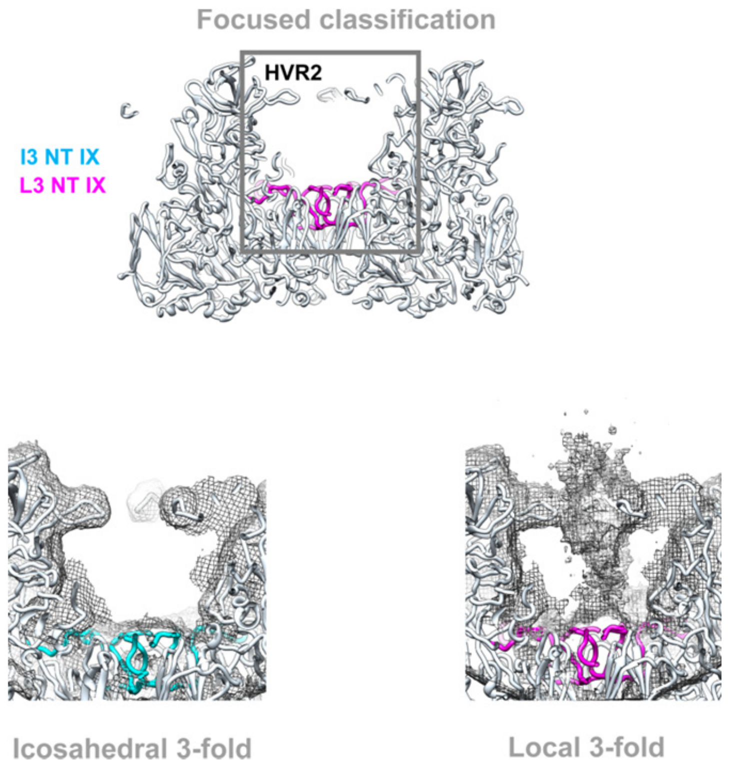


**Figure S7. Focused classification centred at the icosahedral and local 3-fold symmetry axes.** Hexons are shown in white, and the protein IX triskelions in cyan (at the I3 axis) and magenta (at the L3 axis). Density reconstructed in the area where the classification focused is shown as a grey mesh. Top: only the traced models are shown, with a rectangle indicating the area considered for focused classification. Hexon hypervariable region 2 is labelled **HVR2**. Bottom left: there was no density protrusion in any of the classes at the I3 axis. Bottom right: there was density protruding in all classes at the L3 axes. Classes only varied in the strength of the protrusion. None of them showed ordered density that could correspond to a helix bundle. The protrusion connects to the capsid surface formed by protein IX triskelion. Only one class is shown as an example for each of the two 3-fold axes, with the maps contoured at the same level.

**HAdV-C5**

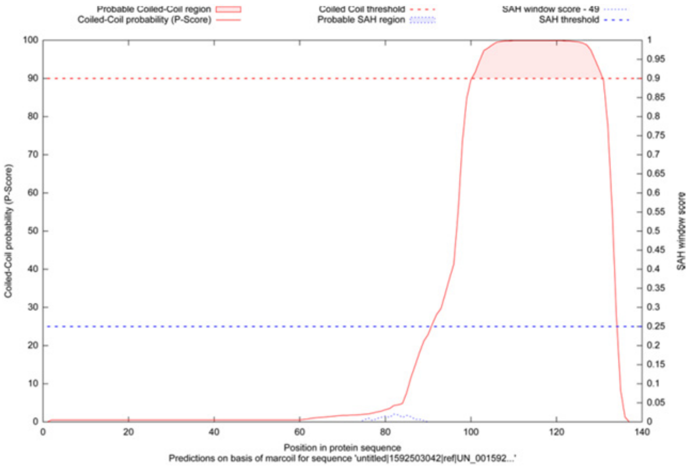


**HAdV-D26**

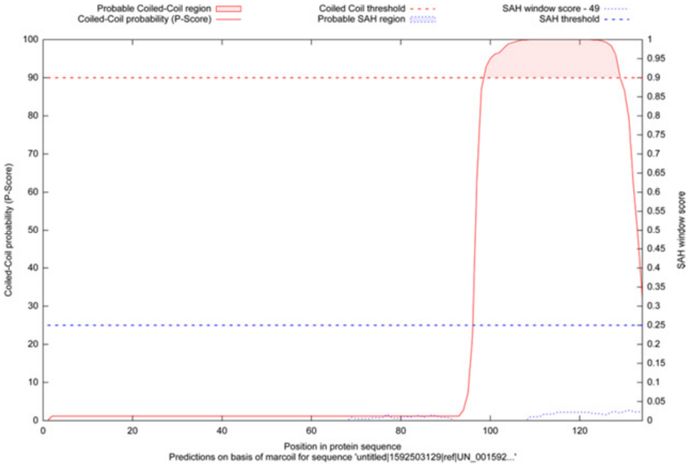


**HAdV-F41**

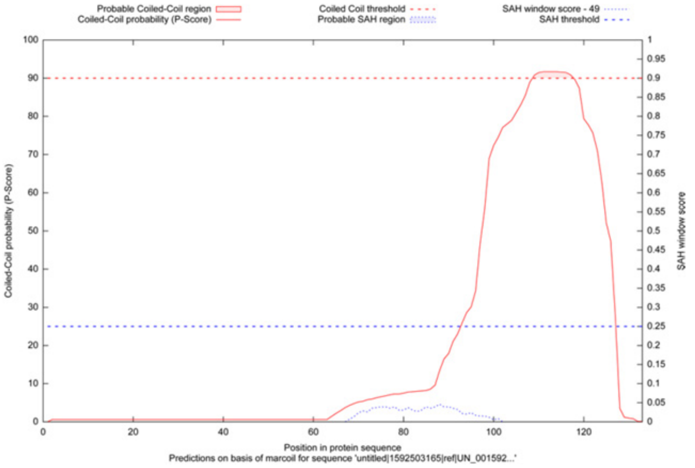


**Figure S8. Probability to form a coiled coil for protein IX in HAdV-C5, D26 or F41.** The plots show the probability calculated with Marcoil (red curves) depending on the sequence position (horizontal axis), as implemented in the Waggawagga Coiled-coil and Single-Alpha-Helix prediction for protein sequences (https://waggawagga.motorprotein.de/). While the prediction for a C-terminal coiled coil is clear in HAdV-C5 and D26, the curve barely surpasses the threshold (dotted red line) in HAdV-F1.

### Supplementary Tables

**Table S1. Cryo-EM data collection, refinement, modelling and validation statistics.**

| **Data collection** | |
|---|---|
| Microscope | Titan Krios |
| Camera | Falcon II |
| Voltage | 300 kV |
| Magnification | 50,000 |
| Nominal pixel size | 1.38 Å |
| Cumulative electron dose | 42 e/Å$^2$ |
| Exposure time | 1.5 s |
| Number of frames | 25 |
| Defocus range | -1 to -2.5 µm |
| Micrographs collected | 5,566 |
| Acquisition software | EPU |
| **Image processing** | |
| Preprocessing software | XMIPP, Scipion |
| Frame alignment software | Xmipp optical flow , Scipion |
| CTF estimation software | CTFFIND4, Scipion |
| Particle picking software | XMIPP, Scipion |
| Micrographs used | 4,867 |
| Particles selected | 13,534 |
| **Reconstruction** | |
| Software | RELION, Scipion |
| Particles included | 9926 |
| Symmetry imposed | Icosahedral |
| Rotational accuracy | 0.05 degrees |
| Translational accuracy | 0.15 pixels |
| B-factor applied | -219.759 Å$^2$ |
| Final resolution (gold standard FSC=0.143) | 4.0 |
| **Model building, refinement and validation** | |
| Experimental pixel size | 1.36 Å |
| Number of modeled residues | 12,388 |
| Number of chains | 26 |
| Software | MonoRes, LocalDeblur, Scipion/ Xmipp, Chimera, Coot, Phenix |
| Model to map correlation coefficient | 0.78 |
| R.M.S. deviations (bond length) | 0.008 Å |
| R.M.S. deviations (bond angle) | 1.465º |
| Ramachandran favored | 85.16% |
| Ramachandran outliers | 0.14% |
| Rotamer outliers | 0.60% |
| C-beta outliers | 0 |
| Molprobity score | 2.13 |
| Clashscore | 8.72 |

**Table S2. Comparison between HAdV-C5 and HAdV-F41 structural protein sequences and predicted isoelectric point.** The isoelectric point was calculated as per Isoelectric Point Calculator (http://isoelectric.org/index.html) (Kozlowski, 2016), using the following values for the amino acids: D=-3.9 E=-4.1 C=-8.5 Y=-10.1 H=6.5 K=10.8 R=12.5, as implemented in Geneious version 2019.0 (https://www.geneious.com). Protein sequence alignments were performed with Clustal W2 (http://www.clustal.org/clustal2/) (Larkin *et al.*, 2007), also integrated in Geneious. Pairwise % identity gives the average percent identity over the alignment. This is computed by looking at all pairs of amino acids at the same column and scoring a hit (one) when they are identical, divided by the total number of pairs.

| **Protein** | **Polypeptide length (amino acids)** | | **Isoelectric point (predicted)** | | **% sequence identity** |
|---|---|---|---|---|---|
| | **HAdV-C5** | **HAdV-F41** | **HAdV-C5** | **HAdV-F41** | |
| **Hexon** | 952 | 925 | 4.9 | 5.5 | 77 |
| **Penton** | 571 | 508 | 5.1 | 5.5 | 68 |
| **IIIa** | 585 | 579 | 5.7 | 5.1 | 69 |
| **VI** | 250 | 266 | 10.4 | 10.1 | 62 |
| **VII** | 198 | 183 | 12.4 | 12.3 | 64 |
| **VIII** | 227 | 233 | 9.2 | 8.5 | 78 |
| **IX** | 140 | 133 | 6.4 | 4.9 | 52 |

**Table S3. Comparison between HAdV-C5 and HAdV-F41 protein structures.** The root mean square deviation (RMSD) between the molecules traced in (Dai *et al.*, 2017) (PDB ID: 6B1T) and those traced in this work is shown, calculated either for all Cα atoms or only for those kept after pruning. Calculated with UCSF Chimera *matchmaker* with default settings (Pettersen *et al.*, 2004).

| Protein | RMSD (Å) | Cα atom pairs | RMSD pruned (Å) | Cα pruned atom pairs |
|---|---|---|---|---|
| **Hexon** | 1.1 | 904 | 0.8 | 853 |
| **Penton base** | 1.2 | 452 | 0.9 | 430 |
| **IIIa** | 1.3 | 288 | 0.9 | 266 |
| **VIII** | 0.7 | 180 | 0.7 | 178 |

**Table S4. Comparison between sequence features of HVRs in HAdV-C5 and HAdV-F41.** The limits of the HAdV-C5 HVRs were taken as (Crawford-Miksza and Schnurr, 1996). Protein sequence alignment, pairwise identity and isoelectric point calculation were performed as indicated in Table S2. The charge at pH 7.0 was calculated as described in http://isoelectric.org/index.html) (Kozlowski, 2016), with amino acid pKa values taken from (Lide and Haynes, 2009), and general pKa values for terminal amino and carboxy groups taken from (Berg *et al.*, 2015).

| **Region** | **% pairwise identity** | **Residue range** | | **Length (amino acids)** | | **Isoelectric point; Charge at pH 7** | |
|---|---|---|---|---|---|---|---|
| | | **HAdV-C5** | **HAdV-F41** | **HAdV-C5** | **HAdV-F41** | **HAdV-C5** | **HAdV-F41** |
| **HVR 1** | 8.8 % | 135-168 | 135-143 | 34 | 9 | 3.22; -13.98 | 10.37; +1.91 |
| **HVR 2** | 22.2 % | 188-195 | 164-172 | 8 | 9 | 2.09; -0.09 | 3.40; -1.09 |
| **HVR 3** | 9.1 % | 212-220 | 189-199 | 9 | 11 | 4.20; -1.99 | 6.09; -0.09 |
| **HVR 4** | 21.4 % | 249-261 | 228-241 | 13 | 14 | 9.18; +0.91 | 3.18; -2.09 |
| **HVR 5** | 11.8 % | 268-284 | 248-258 | 17 | 11 | 4.03; -1.09 | 6.09; -0.09 |
| **HVR 6** | 23.1 % | 305-315 | 279-291 | 11 | 13 | 6.24; -0.09 | 3.18; -2.09 |
| **HVR 7** | 30.6 %; | 419-453 | 395-426 | 35 | 32 | 4.84; -1.08 | 3.62; -4.09 |
| **Total** | 18.7 %; | | | 127 | 99 | 4.03; -16.86 | 3.85; -7 |

**Table S5. HAdV-F41 proteins traced in this model.**

| Protein (UniProt ID) | Length (amino acids) | Copy number in AU | Chain ID | Residues traced | Not traced | Number traced |
|---|---|---|---|---|---|---|
| **hexon (B2ZX09)** | 925 | 12 | A | 6-232; 237-921 | 1-5; 233-236; 922-925 | 912 |
| | | | B | 4-232; 238-921 | 1-3;233-237; 922-925 | 913 |
| | | | C | 2-231; 238-921 | 1; 232-237; 922-925 | 914 |
| | | | D | 6-230; 239-921 | 1-5; 231-238; 922-925 | 908 |
| | | | E | 2-234; 238-921 | 1; 235-237; 922-925 | 917 |
| | | | F | 6-231; 238-921 | 1-5; 232-237; 922-925 | 910 |
| | | | G | 6-231; 238-921 | 1-5; 232-237; 922-925 | 910 |
| | | | H | 1-231; 238-921 | 232-237; 922-925 | 915 |
| | | | I | 6-230; 239-921 | 1-5; 231-238; 922-925 | 908 |
| | | | J | 2-231; 238-921 | 1; 232-237; 922-925 | 914 |
| | | | K | 4-231; 238-921 | 1-3; 232-237; 922-925 | 912 |
| | | | L | 2-192; 195-230; 238-921 | 1; 193-194; 231-237; 922-925 | 911 |
| **penton base (Q9QAH8)** | 508 | 1 | M | 35-292; 315-508 | 1-34; 293-314; | 452 |
| **IIIa (Q67716)** | 579 | 1 | N | 12-309 | 1-11; 310-579 | 298 |
| **VIII (B5SNS9)** | 233 | 2 | O | 2-111; 164-233 | 1; 112-163 | 180 |
| | | | P | 2-111; 164-233 | 1; 112-163 | 180 |
| **IX (B5SNR3)** | 133 | 4 | Q | 10-59 | 1-9; 60-133 | 50 |
| | | | R | 10-59 | 1-9; 60-133 | 50 |
| | | | S | 10-59 | 1-9; 60-133 | 50 |
| | | | T | 10-59 | 1-9; 60-133 | 50 |
| **VI (B5SNS4)** | 266 | (undetermined) | U | 5-33 | 1-4; 34-266 | 29 |
| | | | V | 5-33 | 1-4; 34-266 | 29 |
| | | | Y | 5-33 | 1-4; 34-266 | 29 |
| | | | u | 5-33 | 1-4; 34-266 | 29 |
| **VII (B5SNS1)** | 183 | (undetermined) | W | 18-23 | 1-17; 24-183 | 6 |
| | | | w | 12-23 | 1-11; 24-183 | 12 |

**Supplementary files**

**File S1.** List of residues involved in interactions between traced molecules in the HAdV-F41 capsid, and comparison with HAdV-C5 (PDB: 6B1T). Interacting residues were found with UCSF Chimera *findclash* using the protocol *chimera-contacts* in Scipion (Martinez *et al.*, 2020). The first sheet summarizes the contacts between any two chains. The column labelled *#atoms* indicates the number of atoms involved in each contact in HAdV-F41 (black) or HAdV-C5 (red). Model numbers in columns *model_1* or *model_2* refer to different asymmetric units. No comparison has been performed for the small chains (fragments of proteins VI and VII), regardless of whether they had only been traced in both viruses (green shadowed cells) or only in HAdV-F41 (grey shadowed). The rest of the sheets include the name and position of HAdV-F41 and HAdV-C5 residues (black and red, respectively) involved in contacts between two chains. Chain identifiers are in light blue shaded cells. Sheet names refer to the particular interaction described. ST, TT and SS indicate the three kinds of interfaces between hexons (Liu *et al.*, 2010). S refers to the sides of the hexon trimer pseudo-hexagonal base composed by the two β-barrels in a single monomer; T refers to the faces composed by two β-barrels coming from two different hexon momoners. P represents the penton, and SP denotes the hexon-penton interface. Other interfaces are labeled as follows: IIIa monomers (IIIA_IIIA), IIIa-hexon (IIIA_H), IIIa-penton (IIIA_P), IIIa and VIII proteins (IIIA_VIIIO), VIII protein-hexon (VIIIO_H and VIIIP_H), IX protein (ixt1 and ixt2), and interactions between small chains and hexons in HAdV-F41 (smallest_H). Yellow shadowed cells highlight the regions that, according to the contact criteria (see Methods), are different between both viruses.

**File S2.** PDB validation report.
