## Supplementary file 2 for "Cryo-EM structure of enteric adenovirus HAdV-F41 highlights structural divergence among human adenoviruses"

### Full wwPDB EM Map/Model Validation Report (i)

Apr 13, 2020 – 06:10 PM BST

| | | |
|---|---|---|
| PDB ID | : | 6YBA |
| EMDB ID | : | EMD-10768 |
| Title | : | HAdV-F41 Capsid |
| Deposited on | : | 2020-03-16 |
| Resolution | : | 4.00 Å(reported) |

This is a Full wwPDB EM Map/Model Validation Report.

This report is produced by the wwPDB biocuration pipeline after annotation of the structure.

We welcome your comments at **
A user guide is available at
https://www.wwpdb.org/validation/2017/EMValidationReportHelp
with specific help available everywhere you see the (i) symbol.

CONFIDENTIAL VALIDATION REPORT

The following versions of software and data (see references (i)) were used in the production of this report:

| | | |
|---|---|---|
| EMDB validation analysis | : | 0.0.0.dev33 |
| MolProbity | : | 4.02b-467 |
| Percentile statistics | : | 20171227.v01 (using entries in the PDB archive December 27th 2017) |
| Ideal geometry (proteins) | : | Engh & Huber (2001) |
| Ideal geometry (DNA, RNA) | : | Parkinson et al. (1996) |
| Validation Pipeline (wwPDB-VP) | : | 2.10.1 |

### 1 Overall quality at a glance (i)

The following experimental techniques were used to determine the structure:
*ELECTRON MICROSCOPY*

The reported resolution of this entry is 4.00 Å.

Percentile scores (ranging between 0-100) for global validation metrics of the entry are shown in the following graphic. The table shows the number of entries on which the scores are based.

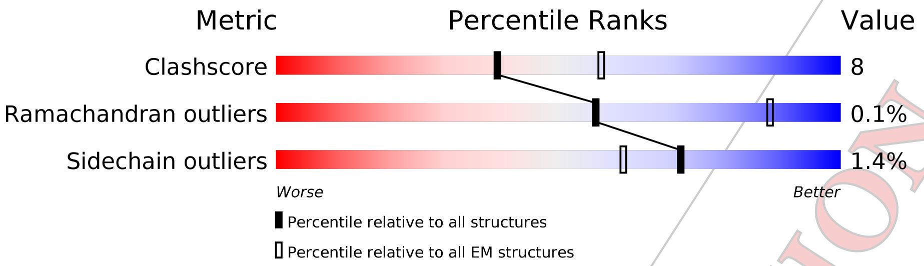


| Metric | Whole archive (#Entries) | EM structures (#Entries) |
|---|---|---|
| Clashscore | 136327 | 1886 |
| Ramachandran outliers | 132723 | 1663 |
| Sidechain outliers | 132532 | 1531 |

The table below summarises the geometric issues observed across the polymeric chains and their fit to the map. The red, orange, yellow and green segments on the bar indicate the fraction of residues that contain outliers for >=3, 2, 1 and 0 types of geometric quality criteria respectively. A grey segment represents the fraction of residues that are not modelled. The numeric value for each fraction is indicated below the corresponding segment, with a dot representing fractions <=5% The upper red bar (where present) indicates the fraction of residues that have poor fit to the EM map (all atom inclusion < 40%). The numeric value is given above the bar.

| Mol | Chain | Length | Quality of chain |
|---|---|---|---|
| 1 | A | 925 | 24% 72% 26% • |
| 1 | B | 925 | 25% 75% 23% • |
| 1 | C | 925 | 23% 72% 26% • |
| 1 | D | 925 | 18% 73% 25% • |
| 1 | E | 925 | 20% 72% 26% •• |
| 1 | F | 925 | 20% 72% 26% • |
| 1 | G | 925 | 18% 73% 25% • |
| 1 | H | 925 | 17% 75% 24% • |

*Continued on next page...*

CONFIDENTIAL VALIDATION REPORT

*Continued from previous page...*

| Mol | Chain | Length | Quality of chain |
|---|---|---|---|
| 1 | I | 925 | 18% / 73% 25% • |
| 1 | J | 925 | 21% / 74% 24% • |
| 1 | K | 925 | 20% / 75% 24% • |
| 1 | L | 925 | 20% / 75% 23% • |
| 2 | M | 508 | 33% / 71% 18% 11% |
| 3 | N | 579 | 21% / 44% 7% 49% |
| 4 | O | 233 | 13% / 58% 19% 23% |
| 4 | P | 233 | 11% / 62% 15% 23% |
| 5 | Q | 133 | 13% / 34% • 62% |
| 5 | R | 133 | 14% / 36% • 62% |
| 5 | S | 133 | 12% / 35% • 62% |
| 5 | T | 133 | 8% / 35% • 62% |
| 6 | U | 266 | 9% / 10% • 89% |
| 6 | V | 266 | 5% / 11% 89% |
| 6 | Y | 266 | • / 10% • 89% |
| 6 | u | 266 | 9% / 11% 89% |
| 7 | W | 183 | • / • • 97% |
| 7 | w | 183 | 5% / 7% 93% |

CONFIDENTIAL VALIDATION REPORT

#### 2 Entry composition (i)

There are 7 unique types of molecules in this entry. The entry contains 98037 atoms, of which 0 are hydrogens and 0 are deuteriums.

In the tables below, the AltConf column contains the number of residues with at least one atom in alternate conformation and the Trace column contains the number of residues modelled with at most 2 atoms.

- Molecule 1 is a protein called Hexon protein.

| Mol | Chain | Residues | Atoms | | | | | AltConf | Trace |
|---|---|---|---|---|---|---|---|---|---|
| | | | Total | C | N | O | S | | |
| 1 | A | 912 | 7244 | 4603 | 1226 | 1379 | 36 | 0 | 0 |
| 1 | B | 913 | 7250 | 4607 | 1227 | 1380 | 36 | 0 | 0 |
| 1 | C | 914 | 7254 | 4610 | 1227 | 1381 | 36 | 0 | 0 |
| 1 | D | 908 | 7214 | 4585 | 1221 | 1372 | 36 | 0 | 0 |
| 1 | E | 917 | 7273 | 4620 | 1231 | 1386 | 36 | 0 | 0 |
| 1 | F | 910 | 7229 | 4595 | 1223 | 1375 | 36 | 0 | 0 |
| 1 | G | 910 | 7229 | 4595 | 1223 | 1375 | 36 | 0 | 0 |
| 1 | H | 915 | 7262 | 4615 | 1228 | 1382 | 37 | 0 | 0 |
| 1 | I | 908 | 7214 | 4585 | 1221 | 1372 | 36 | 0 | 0 |
| 1 | J | 914 | 7254 | 4610 | 1227 | 1381 | 36 | 0 | 0 |
| 1 | K | 912 | 7242 | 4603 | 1225 | 1378 | 36 | 0 | 0 |
| 1 | L | 911 | 7238 | 4601 | 1224 | 1377 | 36 | 0 | 0 |

- Molecule 2 is a protein called Penton protein.

| Mol | Chain | Residues | Atoms | | | | | AltConf | Trace |
|---|---|---|---|---|---|---|---|---|---|
| | | | Total | C | N | O | S | | |
| 2 | M | 452 | 3609 | 2288 | 621 | 688 | 12 | 0 | 0 |

- Molecule 3 is a protein called Pre-hexon-linking protein IIIa.

CONFIDENTIAL VALIDATION REPORT

| Mol | Chain | Residues | Atoms | | | | | AltConf | Trace |
|---|---|---|---|---|---|---|---|---|---|
| 3 | N | 298 | Total 2315 | C 1447 | N 413 | O 451 | S 4 | 0 | 0 |

- Molecule 4 is a protein called Pre-hexon-linking protein VIII.

| Mol | Chain | Residues | Atoms | | | | | AltConf | Trace |
|---|---|---|---|---|---|---|---|---|---|
| 4 | O | 180 | Total 1379 | C 865 | N 236 | O 273 | S 5 | 0 | 0 |
| 4 | P | 180 | Total 1379 | C 865 | N 236 | O 273 | S 5 | 0 | 0 |

- Molecule 5 is a protein called Hexon-interlacing protein.

| Mol | Chain | Residues | Atoms | | | | | AltConf | Trace |
|---|---|---|---|---|---|---|---|---|---|
| 5 | Q | 50 | Total 365 | C 232 | N 65 | O 67 | S 1 | 0 | 0 |
| 5 | R | 50 | Total 365 | C 232 | N 65 | O 67 | S 1 | 0 | 0 |
| 5 | S | 50 | Total 365 | C 232 | N 65 | O 67 | S 1 | 0 | 0 |
| 5 | T | 50 | Total 365 | C 232 | N 65 | O 67 | S 1 | 0 | 0 |

- Molecule 6 is a protein called Pre-protein VI.

| Mol | Chain | Residues | Atoms | | | | | AltConf | Trace |
|---|---|---|---|---|---|---|---|---|---|
| 6 | U | 29 | Total 218 | C 135 | N 42 | O 40 | S 1 | 0 | 0 |
| 6 | V | 29 | Total 218 | C 135 | N 42 | O 40 | S 1 | 0 | 0 |
| 6 | Y | 29 | Total 218 | C 135 | N 42 | O 40 | S 1 | 0 | 0 |
| 6 | u | 29 | Total 218 | C 135 | N 42 | O 40 | S 1 | 0 | 0 |

- Molecule 7 is a protein called Pre-histone-like nucleoprotein.

| Mol | Chain | Residues | Atoms | | | | | AltConf | Trace |
|---|---|---|---|---|---|---|---|---|---|
| 7 | W | 6 | Total 37 | C 23 | N 6 | O 7 | S 1 | 0 | 0 |
| 7 | w | 12 | Total 83 | C 53 | N 15 | O 14 | S 1 | 0 | 0 |

CONFIDENTIAL VALIDATION REPORT

### 3 Residue-property plots (i)

These plots are drawn for all protein, RNA and DNA chains in the entry. The first graphic for a chain summarises the proportions of the various outlier classes displayed in the second graphic. The second graphic shows the sequence view annotated by issues in geometry and atom inclusion in map density. Residues are color-coded according to the number of geometric quality criteria for which they contain at least one outlier: green = 0, yellow = 1, orange = 2 and red = 3 or more. A red diamond above a residue indicates a poor fit to the EM map for this residue (all atom inclusion < 40%). Stretches of 2 or more consecutive residues without any outlier are shown as a green connector. Residues present in the sample, but not in the model, are shown in grey.

• Molecule 1: Hexon protein

Chain A:
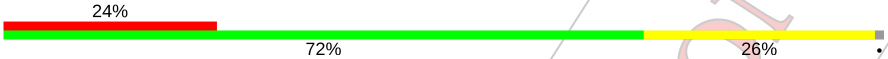


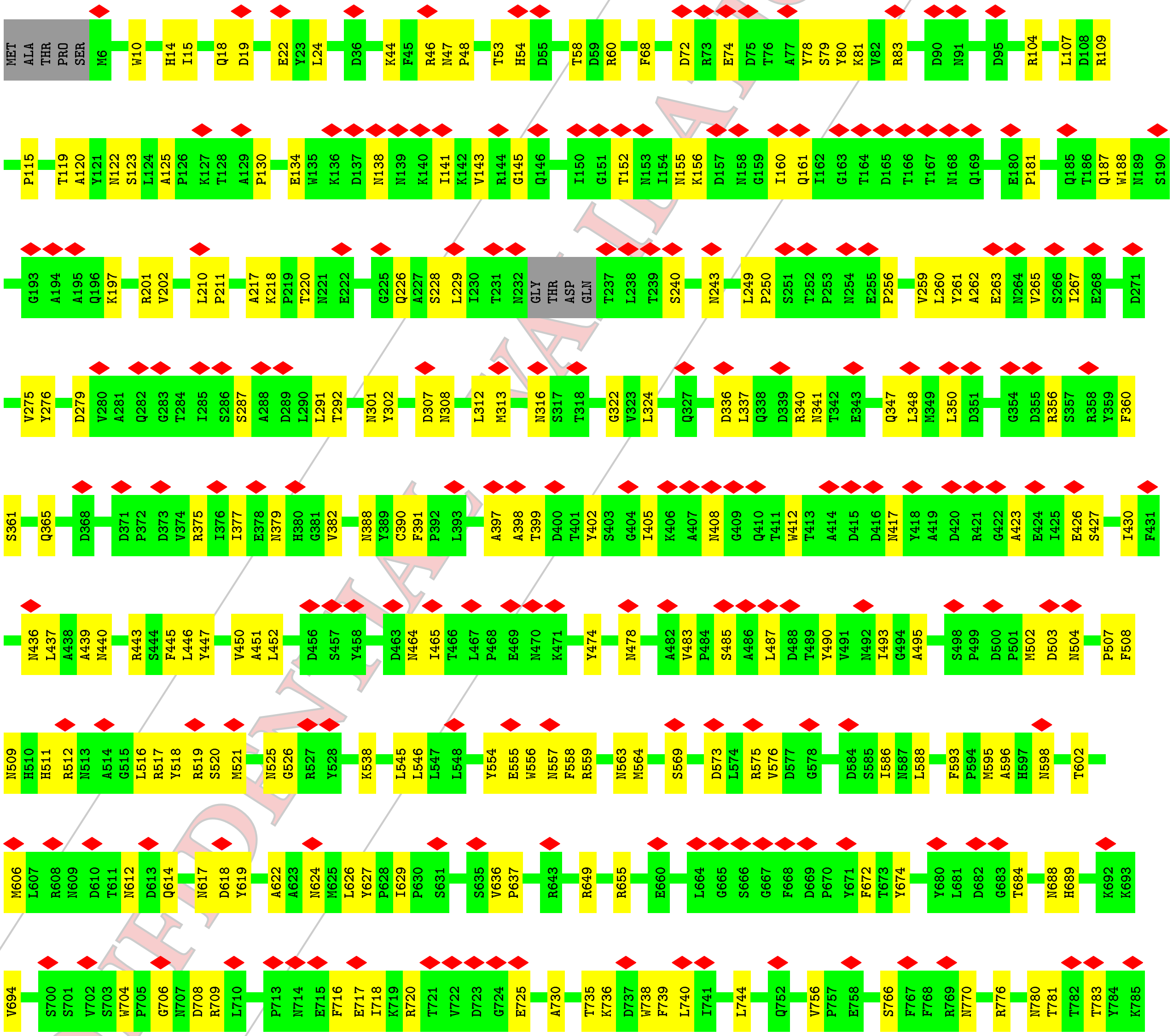


CONFIDENTIAL VALIDATION REPORT

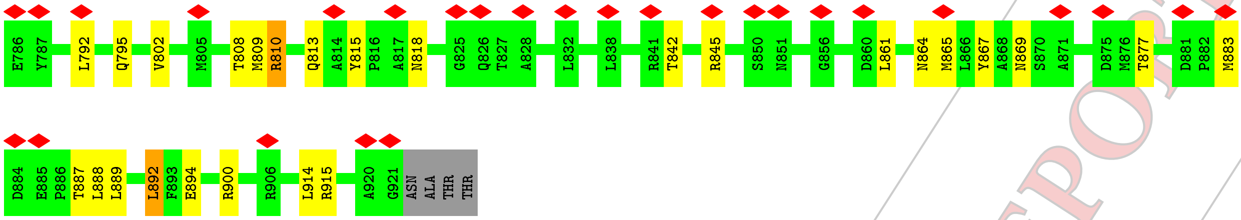

- Molecule 1: Hexon protein

Chain B:

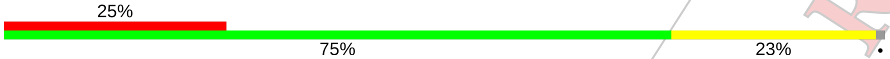

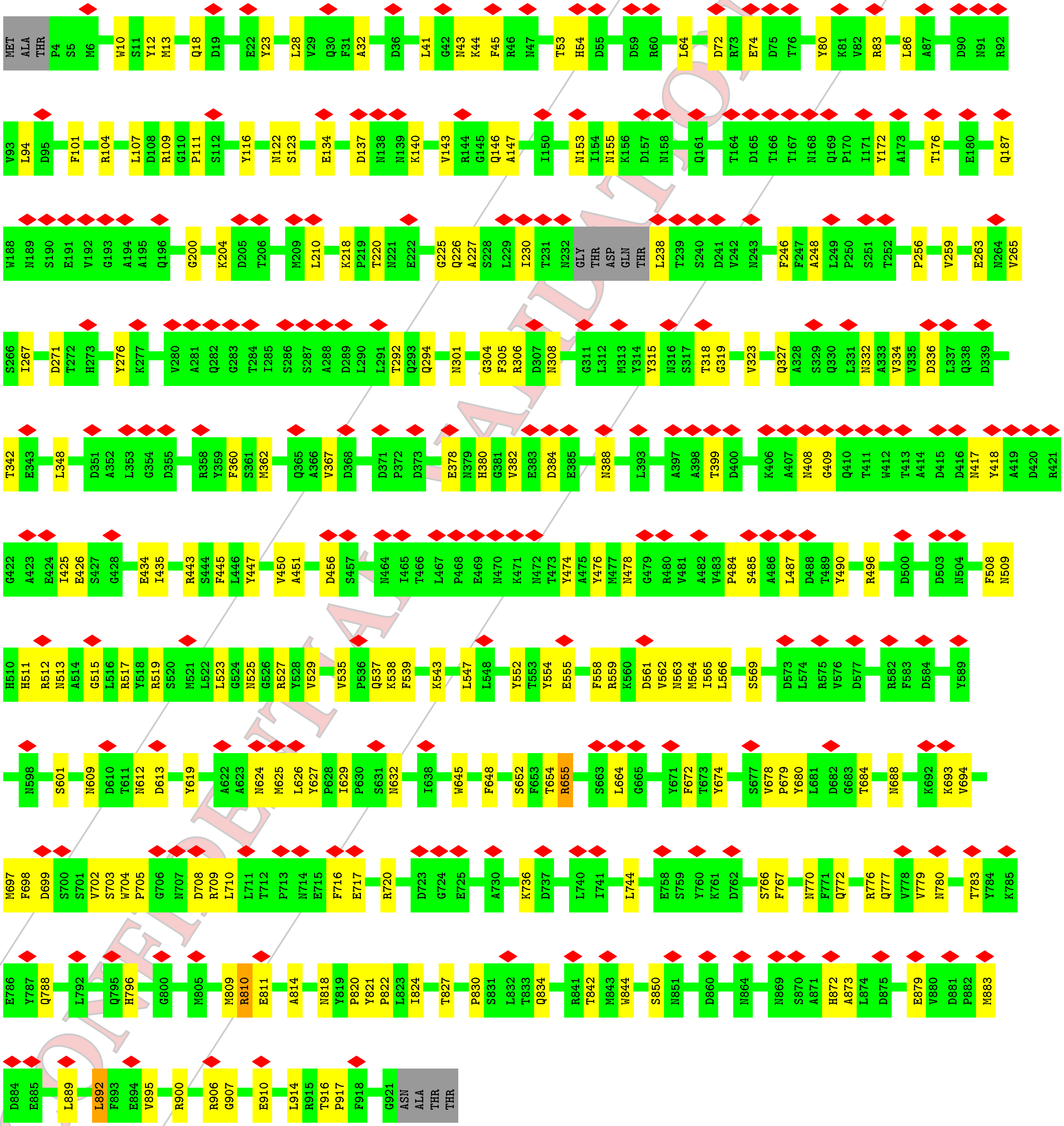

CONFIDENTIAL DRAFT VALIDATION REPORT

- Molecule 1: Hexon protein

Chain C:

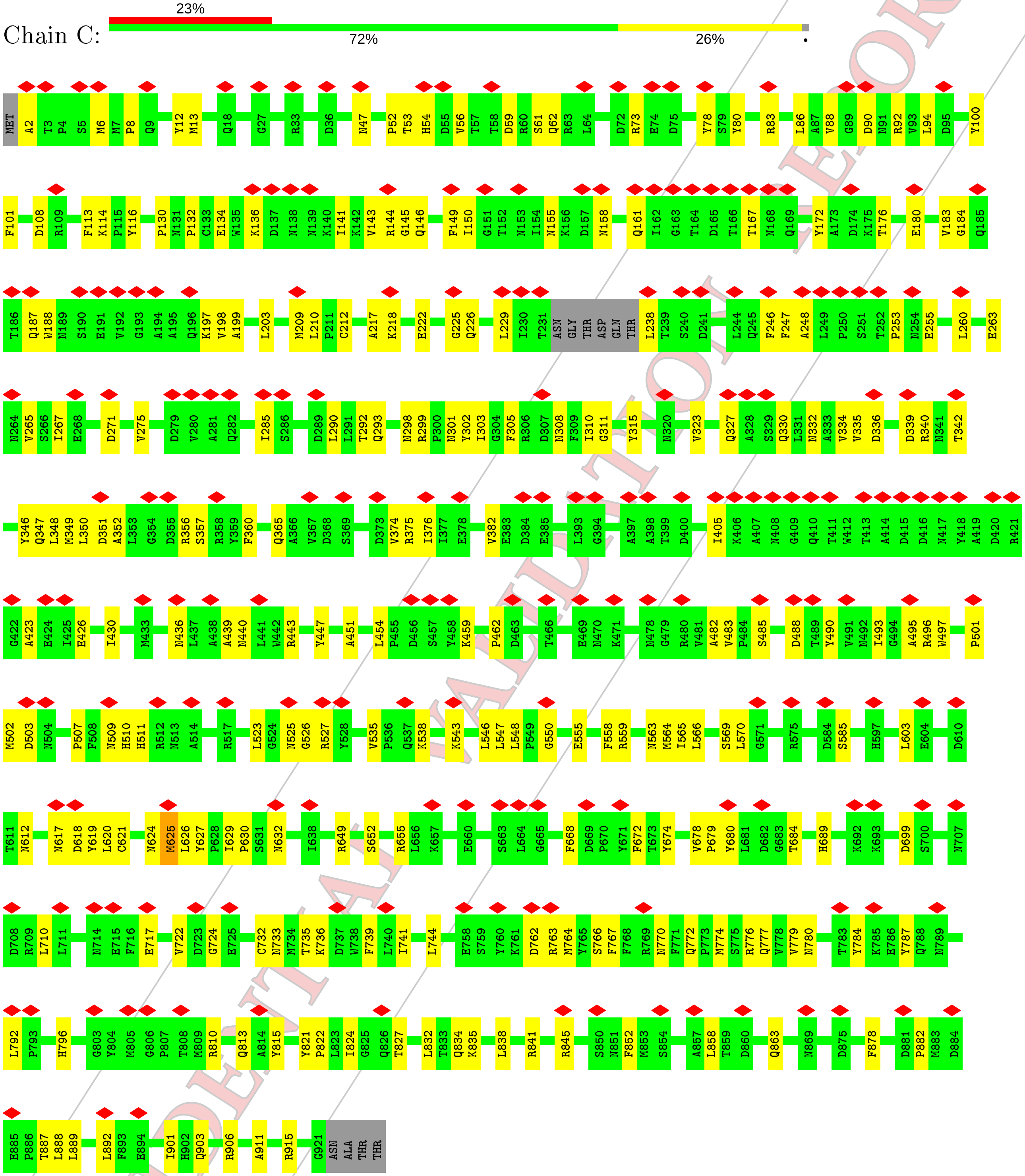


- Molecule 1: Hexon protein

Chain D:

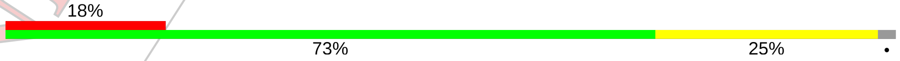


CONFIDENTIAL VALIDATION REPORT

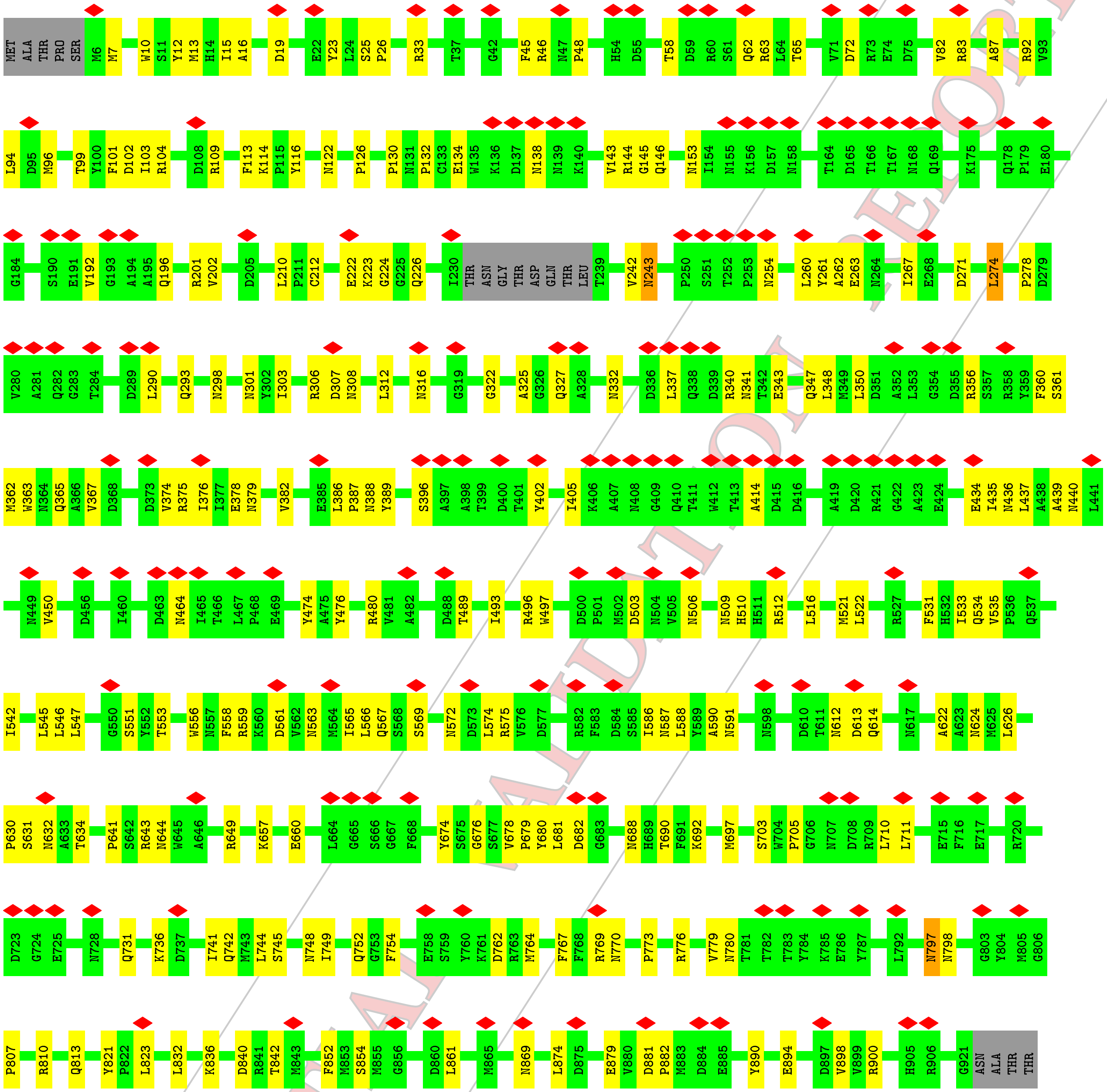

• Molecule 1: Hexon protein

Chain E:

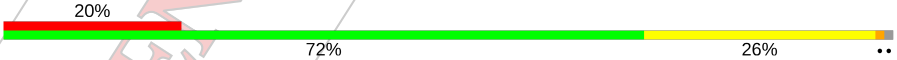


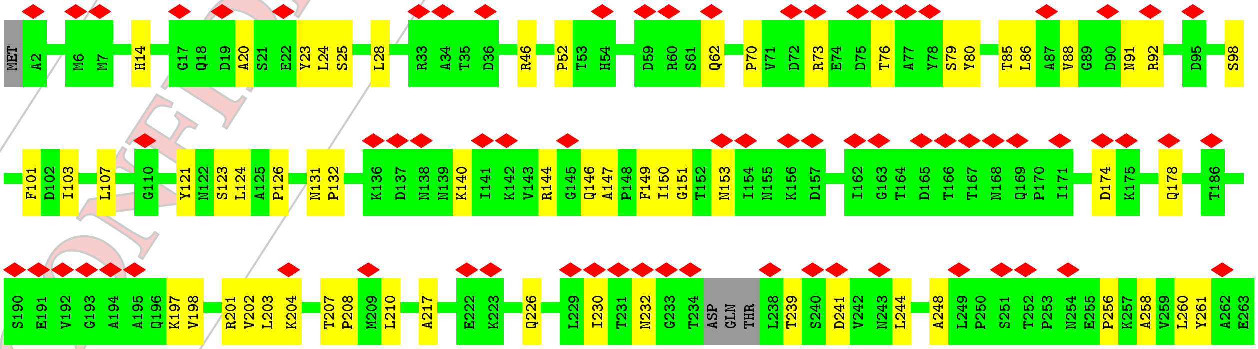

CONFIDENTIAL VALIDATION REPORT

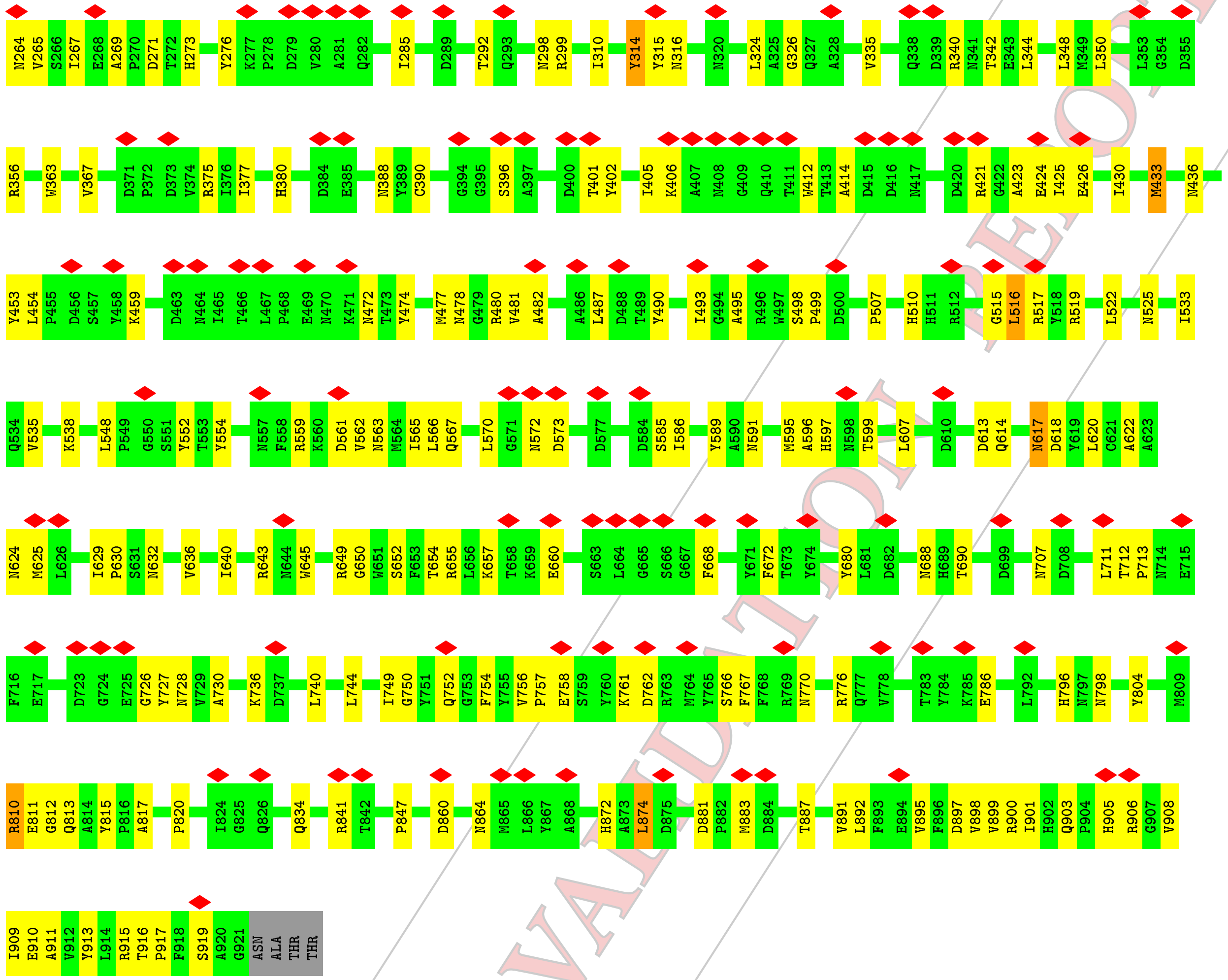


• Molecule 1: Hexon protein

Chain F:

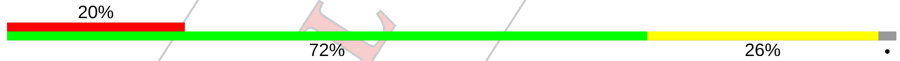


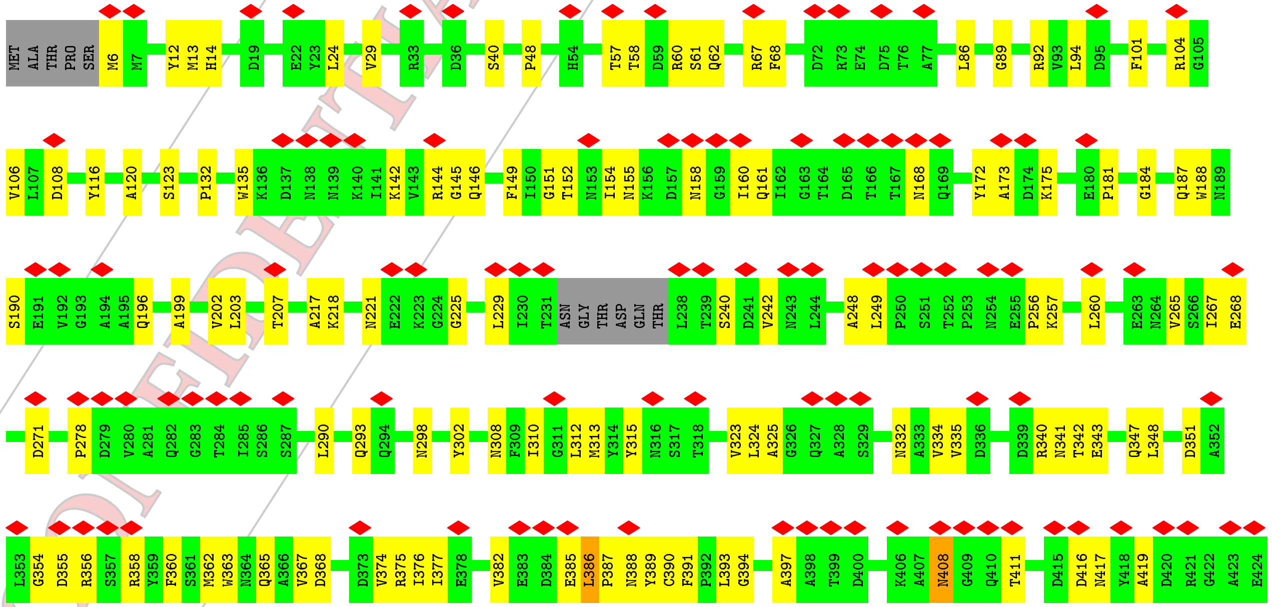


CONFIDENTIAL VALIDATION REPORT

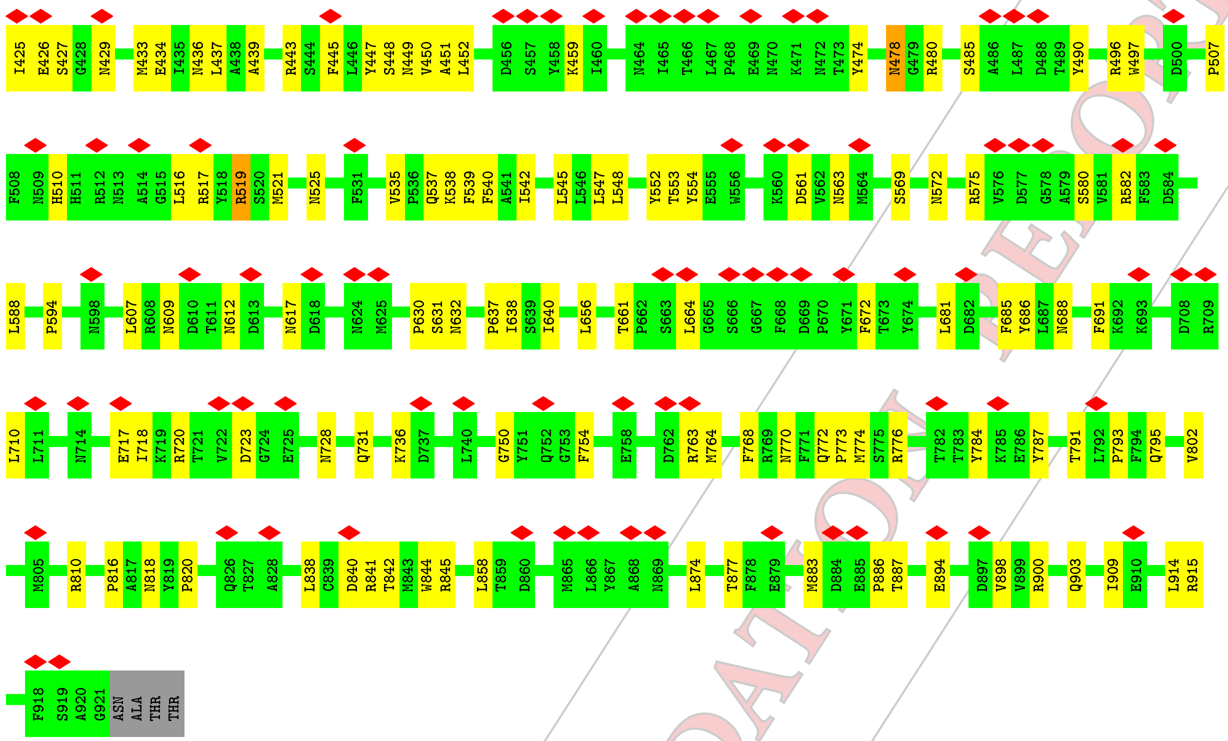


- Molecule 1: Hexon protein

Chain G:

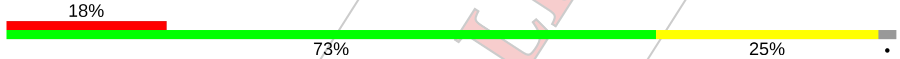


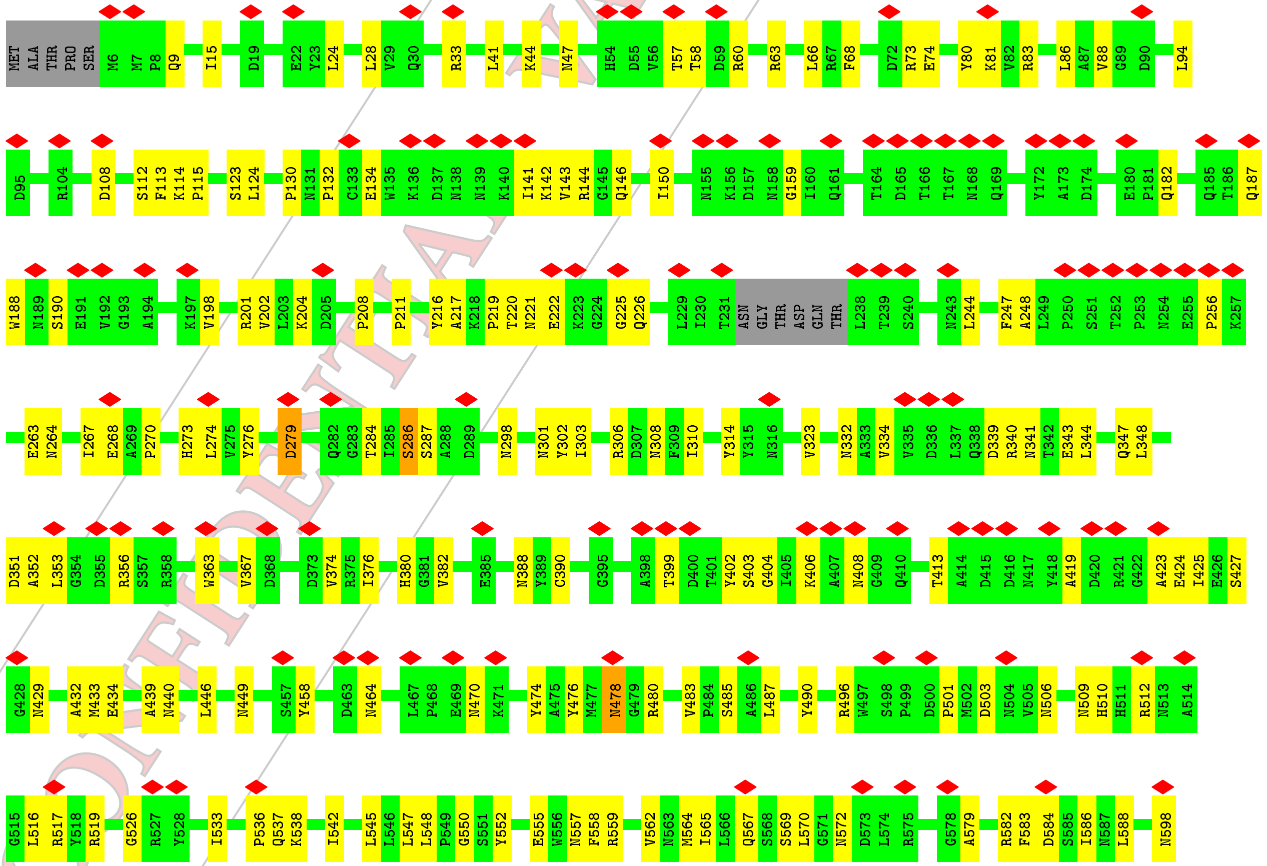


CONFIDENTIAL VALIDATION REPORT

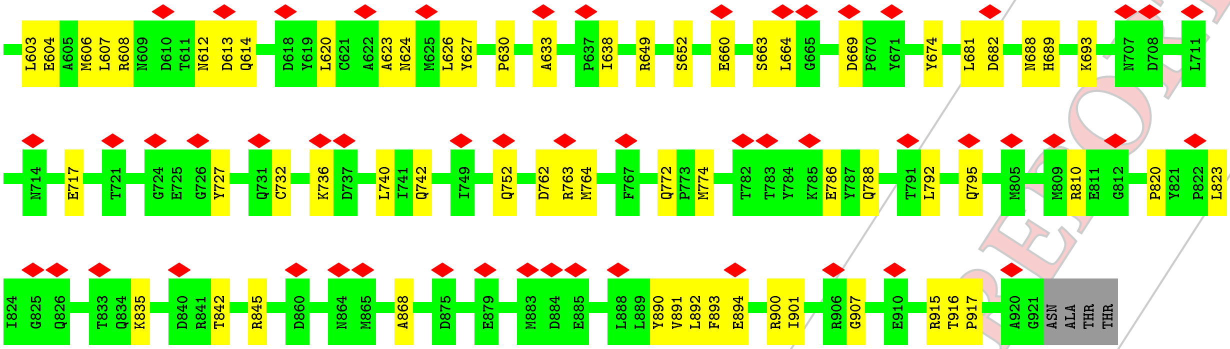


• Molecule 1: Hexon protein

Chain H:

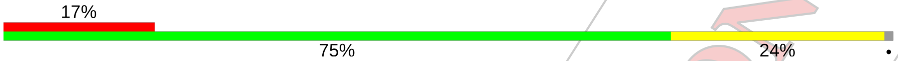


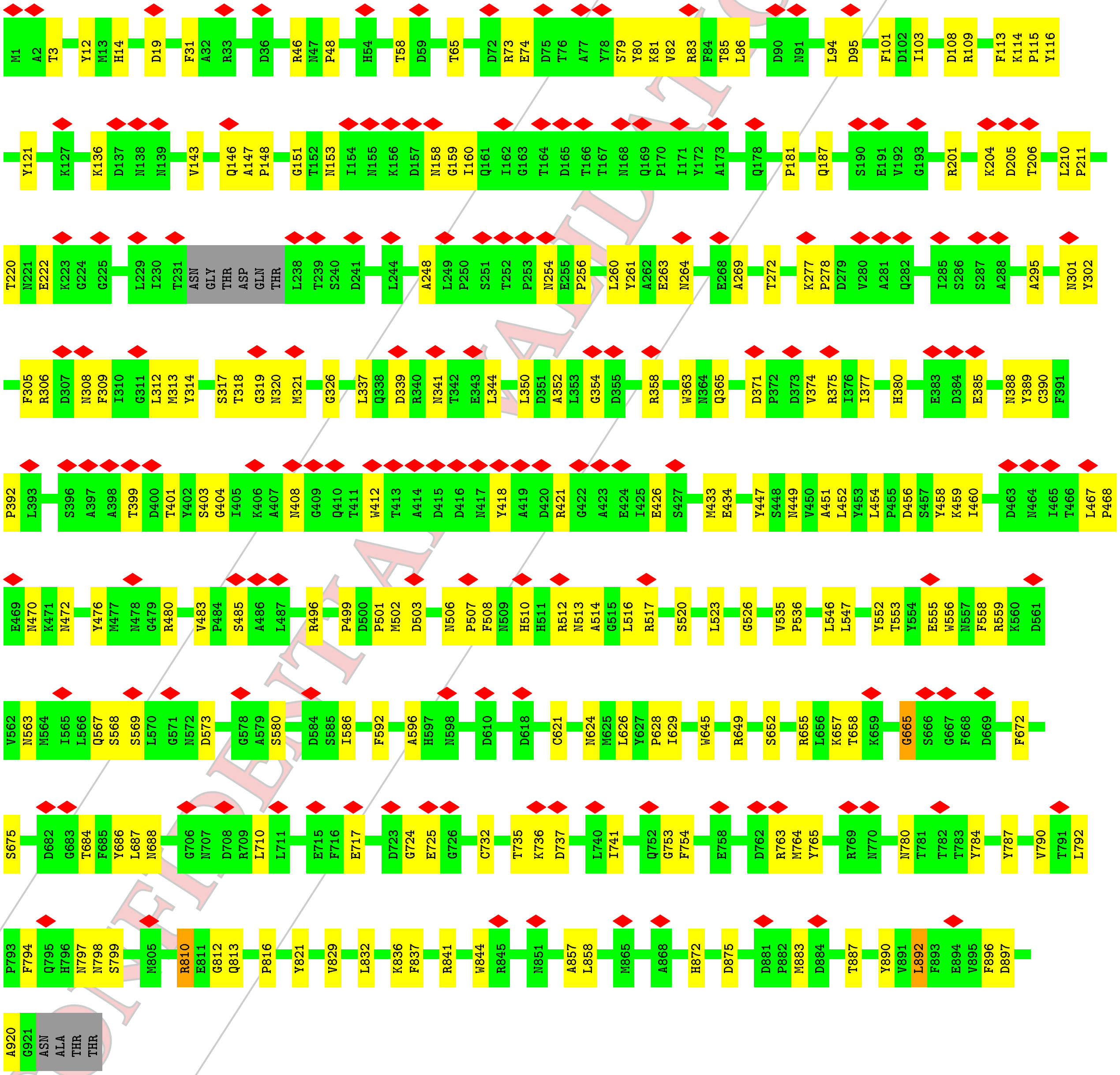

• Molecule 1: Hexon protein

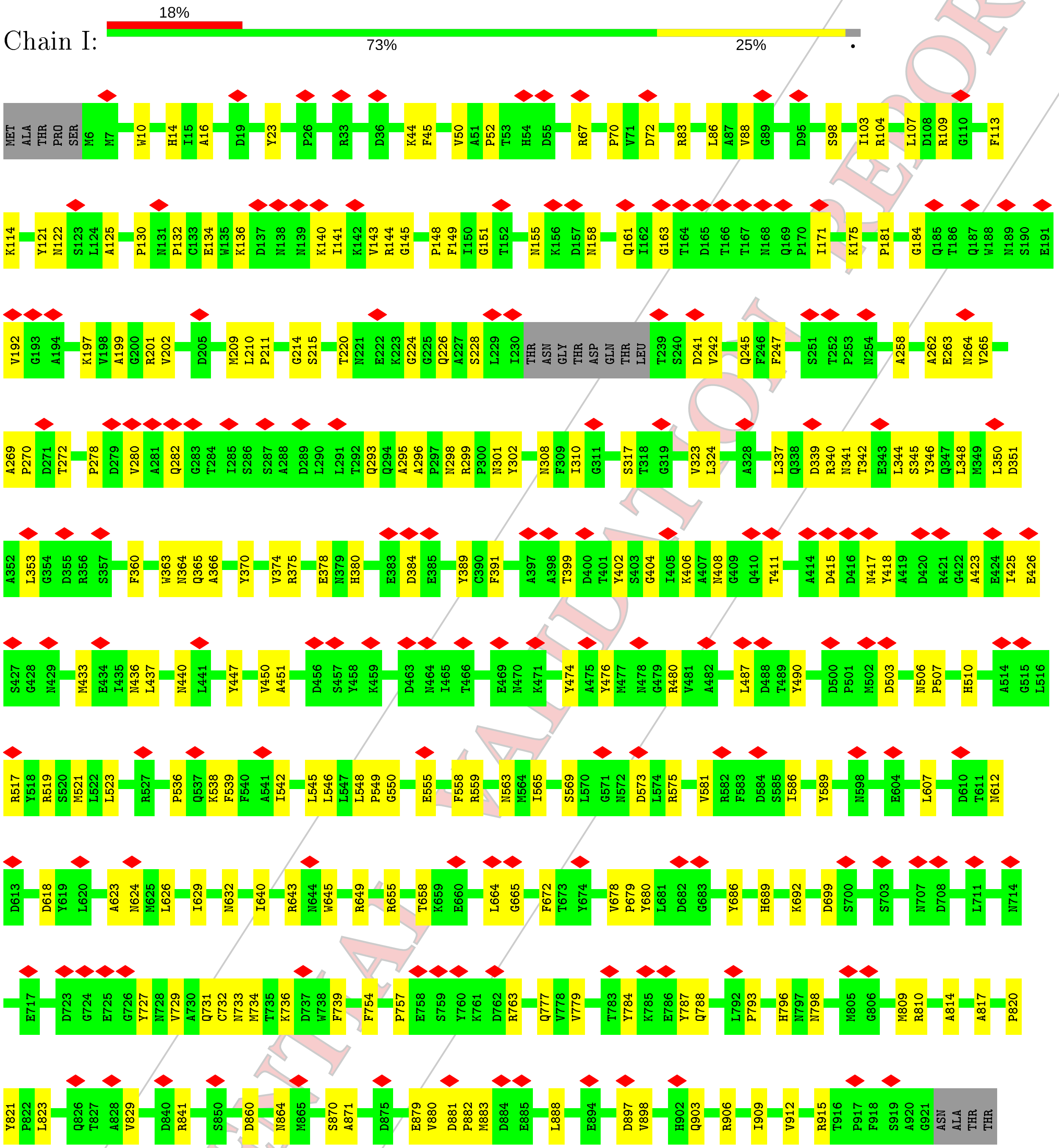


• Molecule 1: Hexon protein

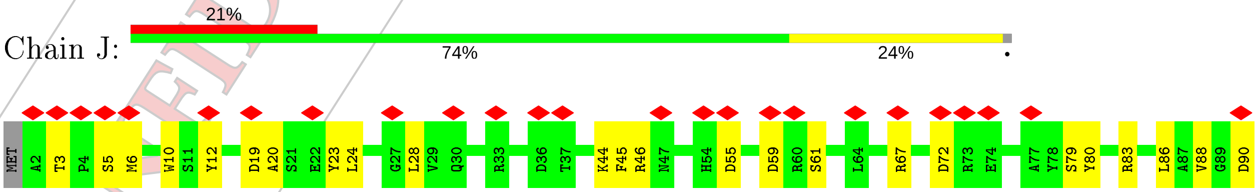


CONFIDENTIAL VALIDATION REPORT

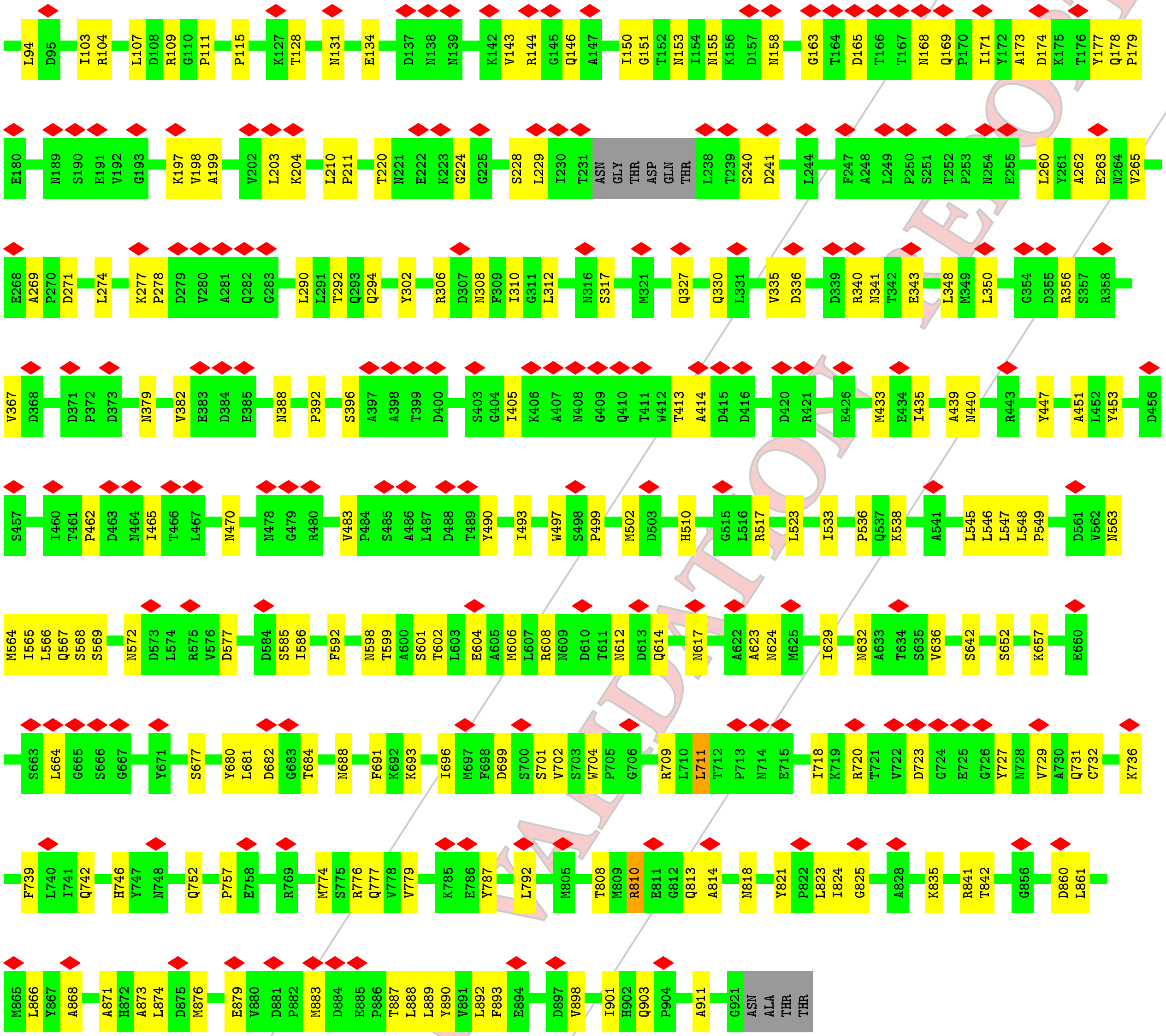


• Molecule 1: Hexon protein

Chain K:

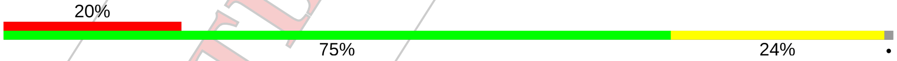


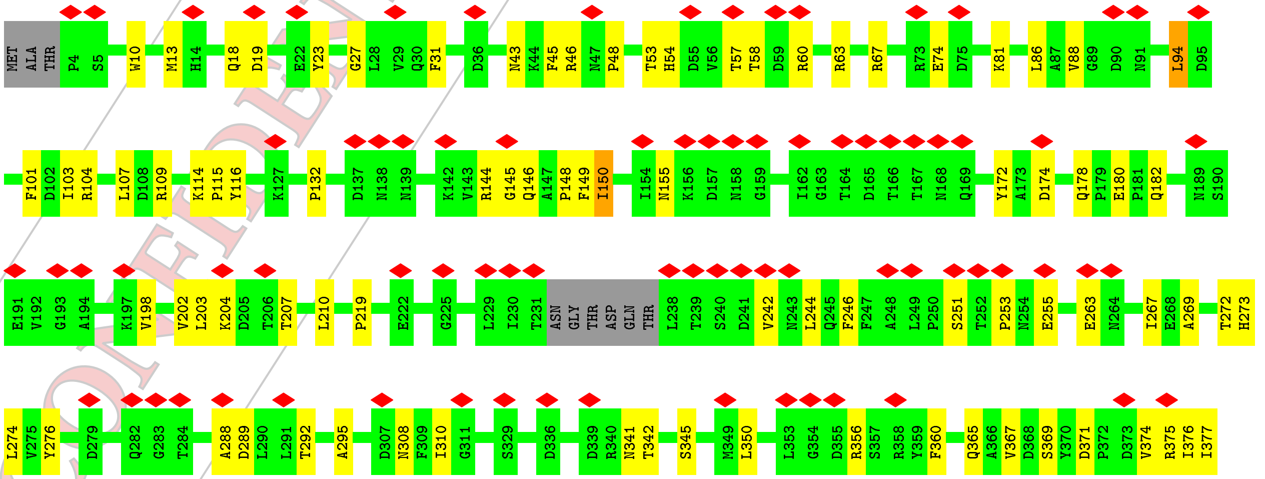


CONFIDENTIAL VALIDATION REPORT

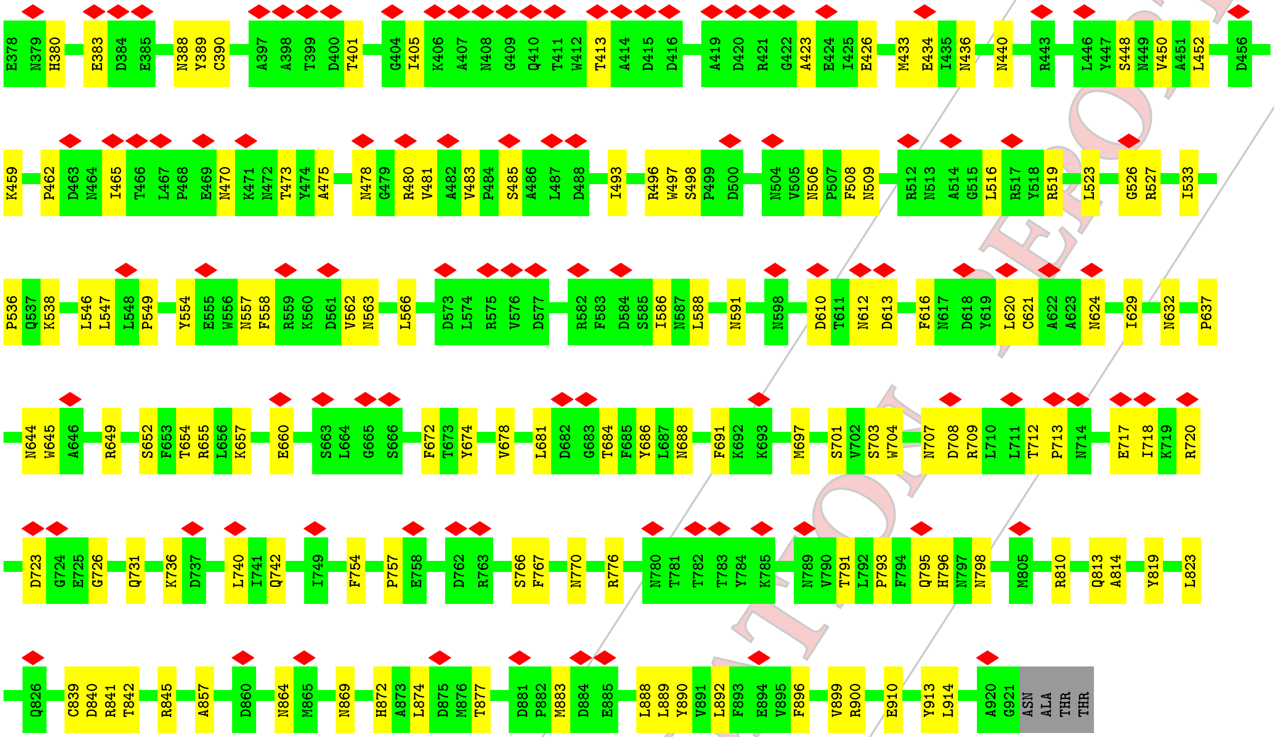


• Molecule 1: Hexon protein

Chain L:

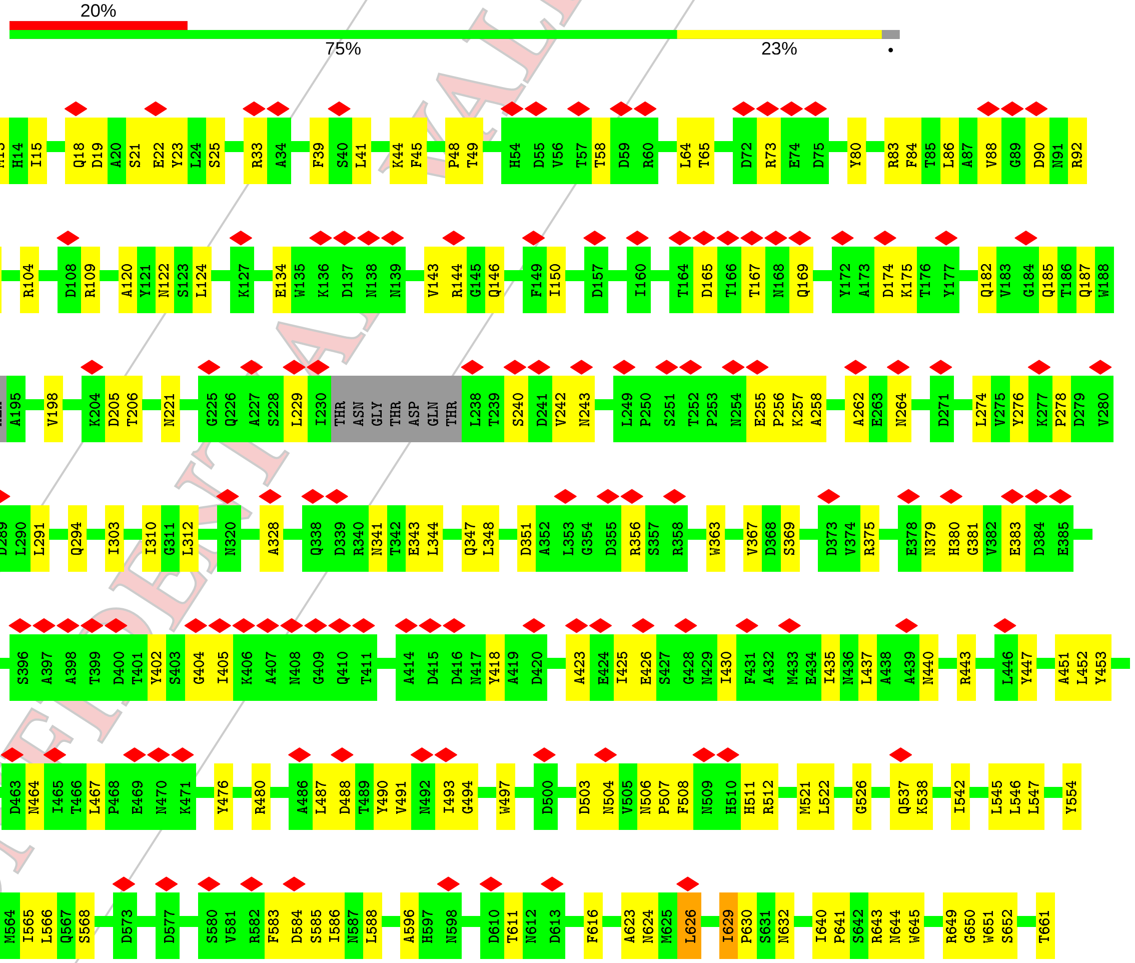

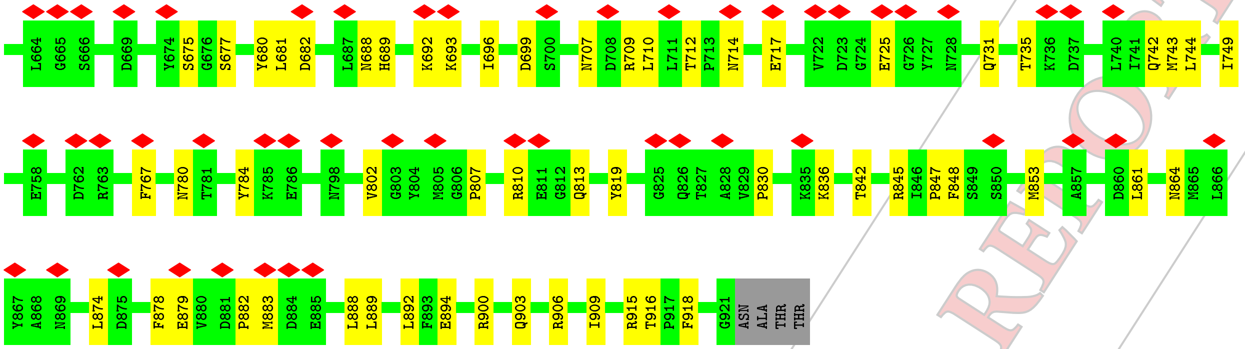


- Molecule 2: Penton protein

Chain M:

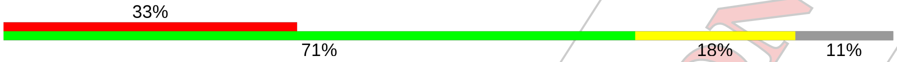


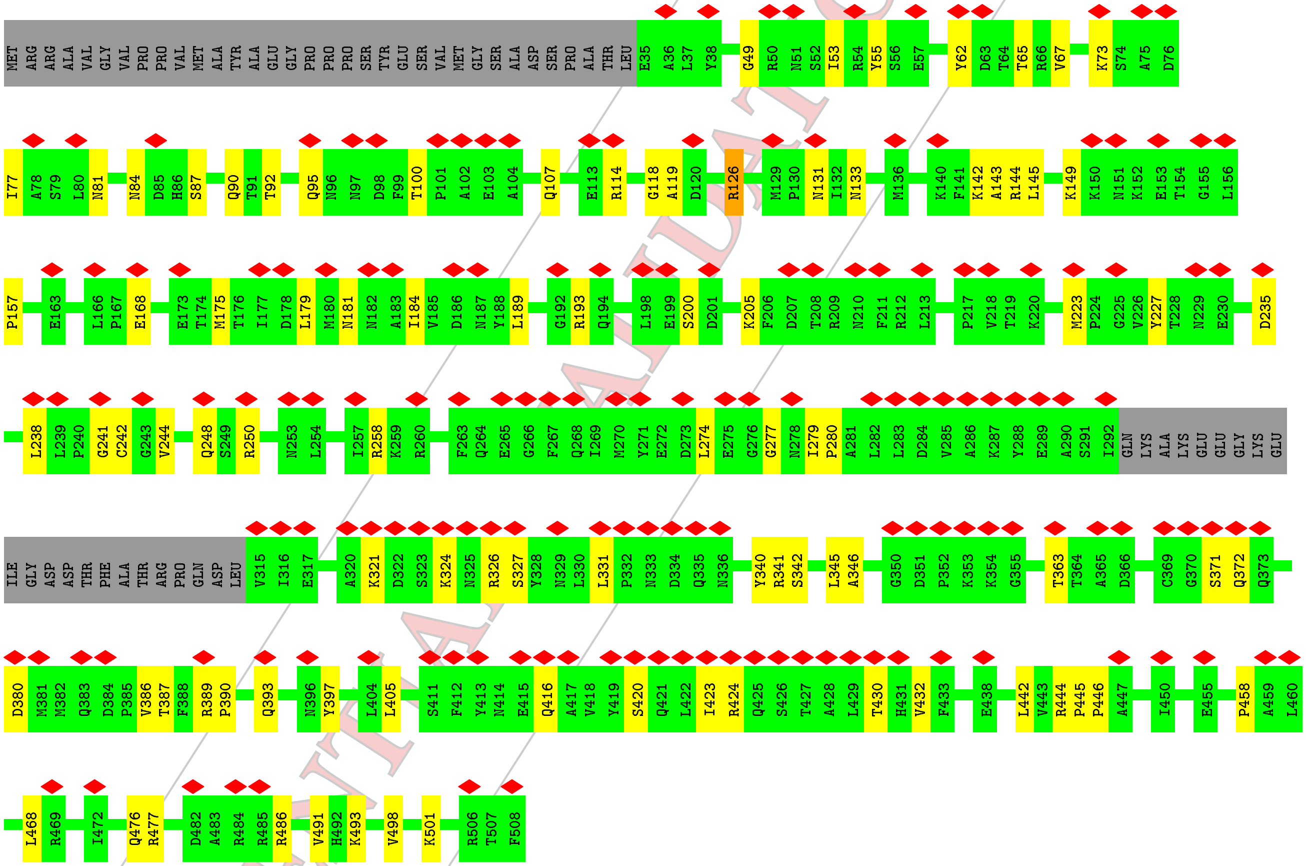


- Molecule 3: Pre-hexon-linking protein IIIa

Chain N:

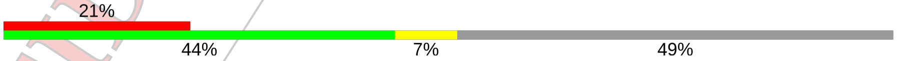


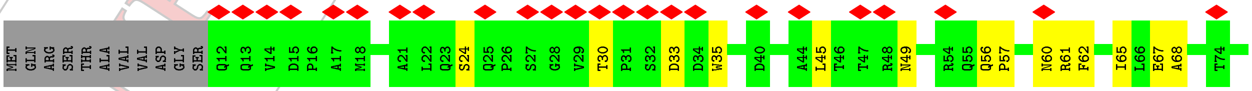


CONFIDENTIAL VALIDATION REPORT

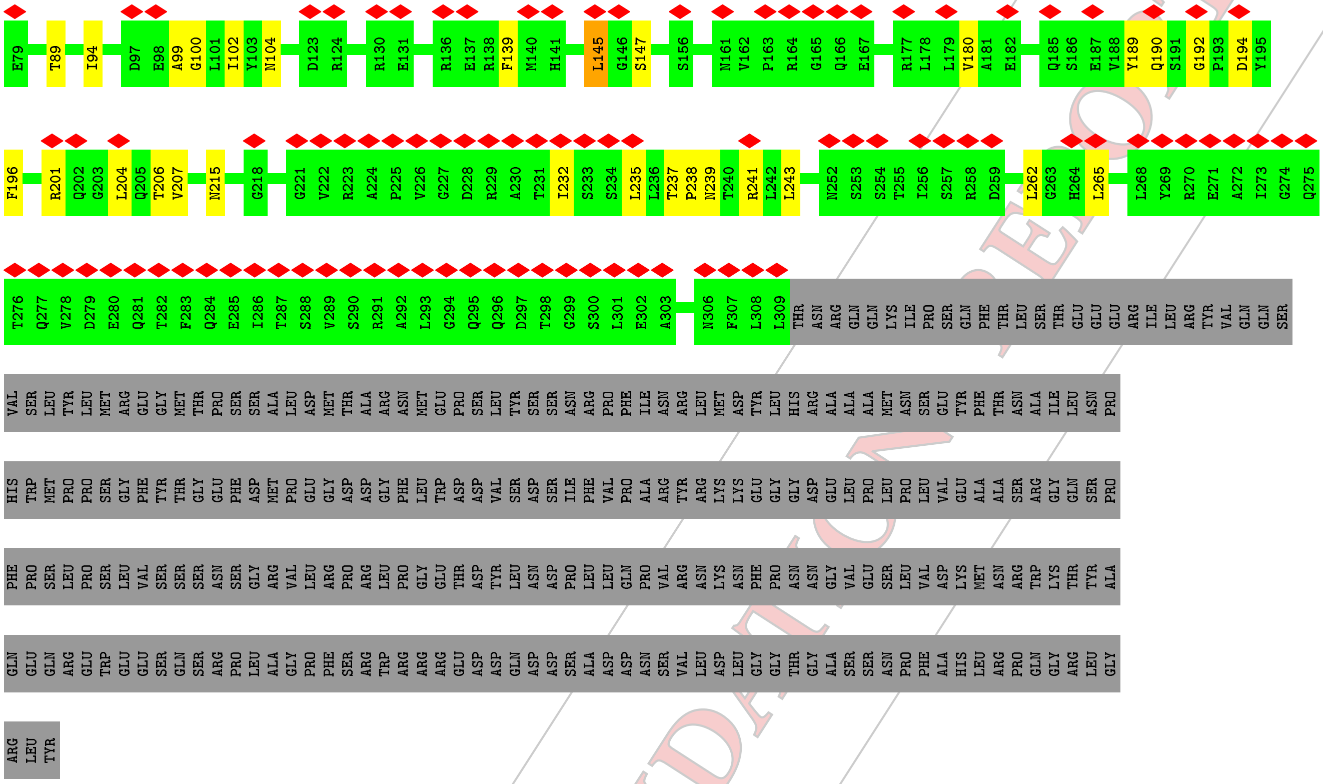


• Molecule 4: Pre-hexon-linking protein VIII

Chain O:

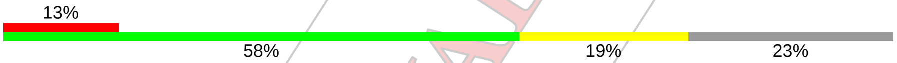


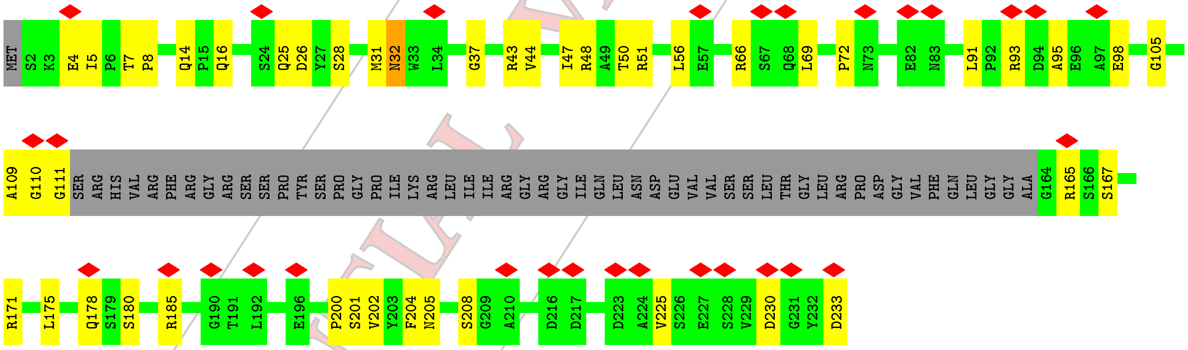


• Molecule 4: Pre-hexon-linking protein VIII

Chain P:

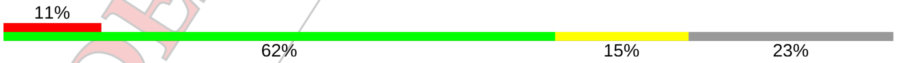


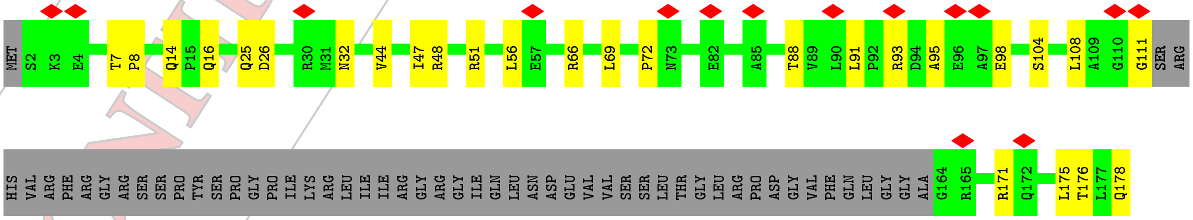


CONFIDENTIAL VALIDATION REPORT

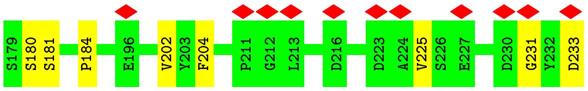


• Molecule 5: Hexon-interlacing protein

Chain Q:

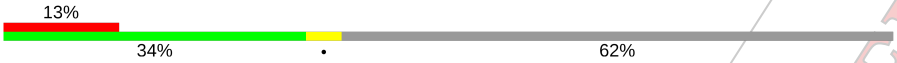


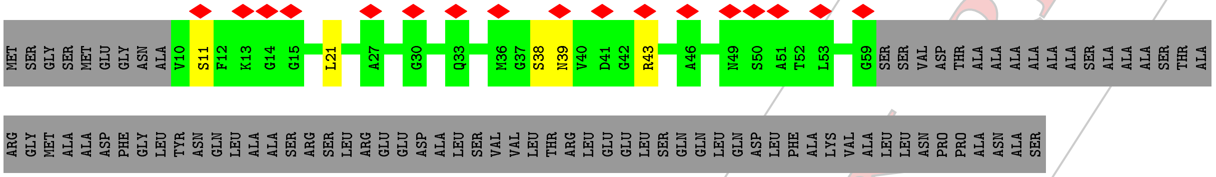


• Molecule 5: Hexon-interlacing protein

Chain R:

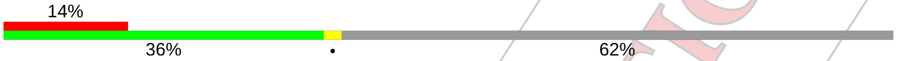


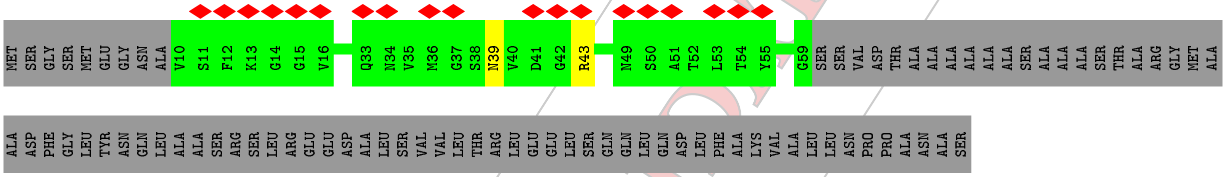


• Molecule 5: Hexon-interlacing protein

Chain S:

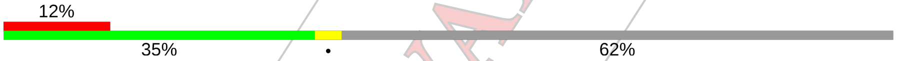


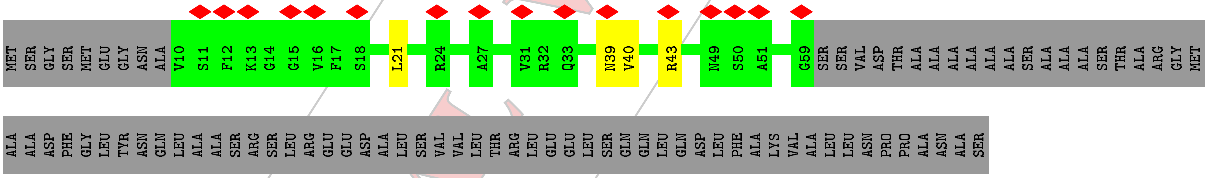


• Molecule 5: Hexon-interlacing protein

Chain T:

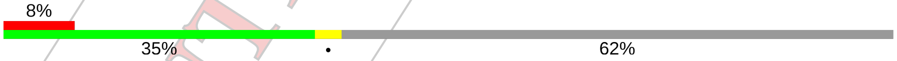


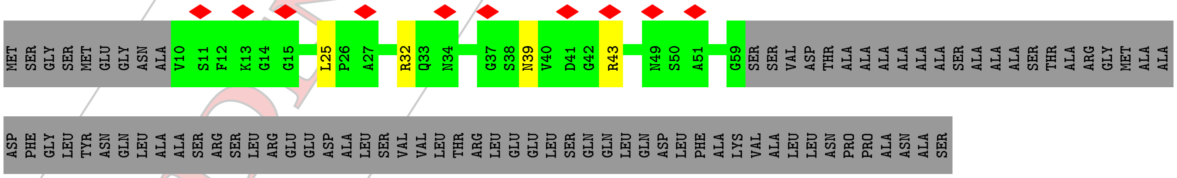


• Molecule 6: Pre-protein VI

Chain U:

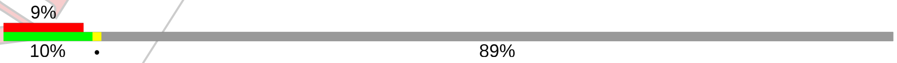


CONFIDENTIAL VALIDATION REPORT

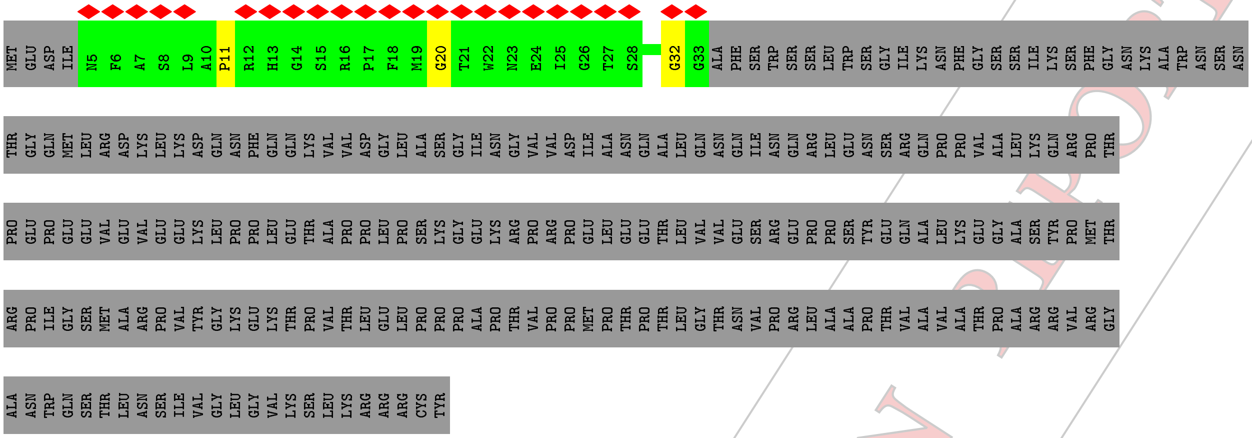


• Molecule 6: Pre-protein VI

Chain V:

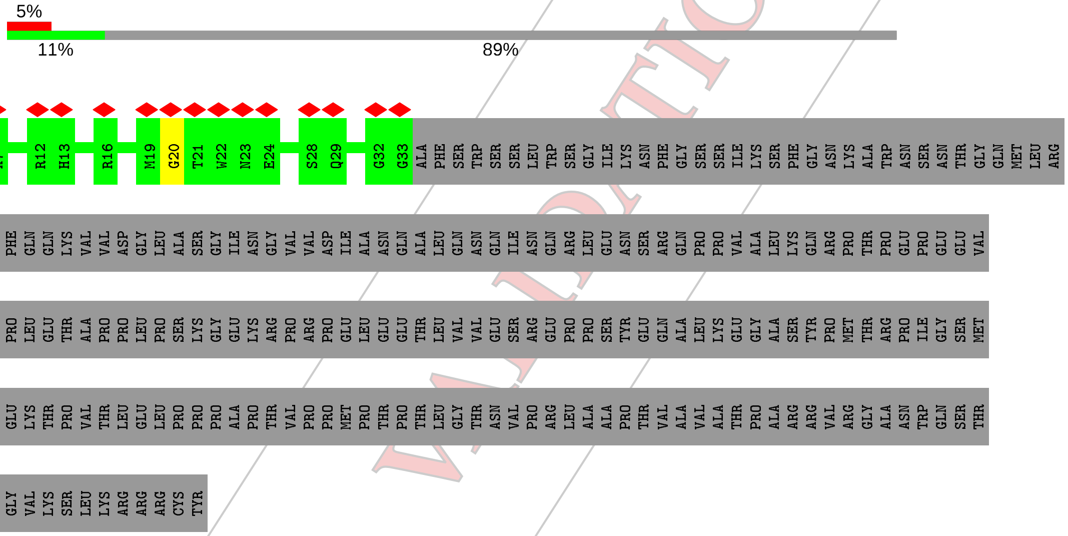


• Molecule 6: Pre-protein VI

Chain Y:

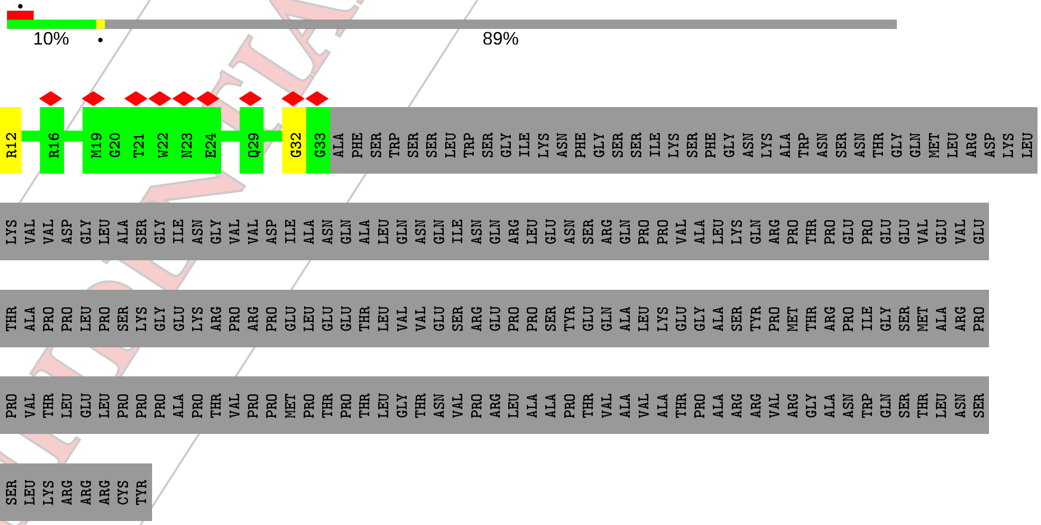


• Molecule 6: Pre-protein VI

CONFIDENTIAL VALIDATION REPORT

Chain u: 9% 11% 89%

MET GLU ASP ILE N5 F6 A7 S8 L9 A10 P11 R12 H13 G14 S15 R16 P17 F18 M19 G20 T21 W22 N23 E24 I25 G26 T27 S28 Q29 L30 N31 G32 G33 ALA PHE SER TRP SER SER LEU TRP SER GLY ILE LYS ASN PHE GLY SER SER ILE LYS SER PHE GLY ASN LYS ALA TRP ASN

SER ASN THR GLY GLN MET LEU ARG ASP LYS LEU LYS ASP GLN ASN PHE GLN GLN LYS VAL VAL ASP GLY LEU ALA SER GLY ILE ASN GLY VAL VAL ASP ILE ALA ASN GLN ALA LEU GLN ASN GLN ILE ASN GLN ARG LEU GLU ASN SER ARG GLN PRO PRO VAL ALA LEU LYS GLN ARG

PRO THR PRO GLU PRO GLU GLU VAL GLU VAL GLU GLU LYS LEU PRO PRO LEU GLU THR ALA PRO PRO LEU PRO SER LYS GLY GLU LYS ARG PRO ARG PRO GLU LEU GLU GLU THR LEU VAL VAL GLU SER ARG GLU PRO PRO SER TYR GLU GLN ALA LEU LYS GLU GLY ALA SER TYR PRO

MET THR ARG PRO ILE GLY SER MET ALA ARG PRO VAL TYR GLY LYS GLU LYS THR PRO VAL THR LEU GLU LEU PRO PRO PRO ALA PRO THR VAL PRO PRO MET PRO THR PRO THR LEU GLY THR ASN VAL PRO ARG LEU ALA ALA PRO THR VAL ALA VAL ALA THR PRO ALA ARG ARG VAL

ARG GLY ALA ASN TRP GLN SER THR LEU ASN SER ILE VAL GLY LEU GLY VAL LYS SER LEU LYS ARG ARG ARG CYS TYR

- Molecule 7: Pre-histone-like nucleoprotein

Chain W: 97%

MET SER ILE LEU ILE SER PRO ASP ASN ASN THR GLY TRP GLY LEU HIS SER A18 G19 M20 Y21 G22 G23 ALA LYS ARG ARG SER SER GLN HIS PRO VAL ARG VAL ARG GLY HIS TYR ARG ALA PRO TRP GLY ALA HIS THR ARG GLY ILE ILE SER GLY ARG THR THR VAL ASP ASP VAL

ILE ASP SER VAL VAL ALA ASP ALA ARG ARG TYR GLN ARG PRO THR SER THR VAL ASP SER VAL ILE ASP SER VAL VAL ALA ASP ALA ARG ARG TYR ALA GLN ARG LYS SER ARG LEU ARG ARG ARG ARG ARG ARG PRO THR THR ALA MET ILE ALA ALA ARG ALA VAL LEU ARG ARG ALA

ARG ARG ILE GLY ARG ARG ALA MET ARG ARG ALA ALA ALA ALA ALA SER ALA GLY ARG ALA ARG ARG GLN ALA ALA ARG GLN ALA ALA ALA ALA ILE ALA SER MET ALA GLN PRO ARG ARG GLY ASN VAL TYR TRP VAL ARG ASP ALA SER GLY VAL ARG VAL PRO VAL ARG THR ARG PRO

PRO ARG SER

- Molecule 7: Pre-histone-like nucleoprotein

Chain w: 5% 7% 93%

MET SER ILE LEU ILE SER PRO ASP ASN ASN THR G12 W13 G14 L15 H16 S17 A18 Y21 G22 G23 ALA LYS ARG ARG SER SER GLN HIS PRO VAL ARG VAL ARG GLY HIS TYR ARG ALA PRO TRP GLY ALA HIS THR ARG GLY ILE ILE SER GLY ARG THR THR VAL ASP ASP VAL ILE

ASP SER VAL VAL ALA ASP ALA ARG ARG TYR GLN ARG PRO THR SER THR VAL ASP SER VAL ILE ASP SER VAL VAL ALA ASP ALA ARG ARG TYR ALA GLN ARG LYS SER ARG LEU ARG ARG ARG ARG ARG ARG PRO THR THR ALA MET ILE ALA ALA ARG ALA VAL LEU ARG ARG ALA ARG

ARG ILE GLY ARG ARG ALA MET ARG ARG ALA ALA ALA ALA ALA SER ALA GLY ARG ALA ARG ARG GLN ALA ALA ARG GLN ALA ALA ALA ALA ILE ALA SER MET ALA GLN PRO ARG ARG GLY ASN VAL TYR TRP VAL ARG ASP ALA SER GLY VAL ARG VAL PRO VAL ARG THR ARG PRO PRO

ARG SER

CONFIDENTIAL VALIDATION REPORT

### 4 Experimental information (i)

| Property | Value | Source |
|---|---|---|
| EM reconstruction method | SINGLE PARTICLE | Depositor |
| Imposed symmetry | POINT, I | Depositor |
| Number of particles used | 9926 | Depositor |
| Resolution determination method | FSC 0.143 CUT-OFF | Depositor |
| CTF correction method | PHASE FLIPPING AND AMPLITUDE CORRECTION | Depositor |
| Microscope | FEI TITAN KRIOS | Depositor |
| Voltage (kV) | 300 | Depositor |
| Electron dose ($e^-/\text{Å}^2$) | 42 | Depositor |
| Minimum defocus (nm) | Not provided | Depositor |
| Maximum defocus (nm) | Not provided | Depositor |
| Magnification | Not provided | Depositor |
| Image detector | FEI FALCON III (4k x 4k) | Depositor |
| Maximum map value | 0.364 | Depositor |
| Minimum map value | -0.253 | Depositor |
| Average map value | 0.001 | Depositor |
| Map value standard deviation | 0.027 | Depositor |
| Recommended contour level | 0.07 | Depositor |
| Map size (Å) | 1060.8, 1060.8, 1060.8 | Depositor |
| Map dimensions | 780, 780, 780 | Depositor |
| Map angles (°) | 90.0, 90.0, 90.0 | Depositor |
| Pixel spacing (Å) | 1.36, 1.36, 1.36 | Depositor |

CONFIDENTIAL VALIDATION REPORT

### 5 Model quality (i)

#### 5.1 Standard geometry (i)

The Z score for a bond length (or angle) is the number of standard deviations the observed value is removed from the expected value. A bond length (or angle) with $|Z| > 5$ is considered an outlier worth inspection. RMSZ is the root-mean-square of all Z scores of the bond lengths (or angles).

| Mol | Chain | Bond lengths | | Bond angles | |
|---|---|---|---|---|---|
| | | RMSZ | #\|Z\| >5 | RMSZ | #\|Z\| >5 |
| 1 | A | 0.39 | 0/7444 | 0.68 | 5/10145 (0.0%) |
| 1 | B | 0.39 | 0/7451 | 0.70 | 4/10154 (0.0%) |
| 1 | C | 0.38 | 0/7455 | 0.70 | 8/10161 (0.1%) |
| 1 | D | 0.40 | 0/7414 | 0.71 | 9/10103 (0.1%) |
| 1 | E | 0.40 | 0/7474 | 0.69 | 11/10187 (0.1%) |
| 1 | F | 0.40 | 0/7429 | 0.69 | 7/10124 (0.1%) |
| 1 | G | 0.42 | 0/7429 | 0.71 | 9/10124 (0.1%) |
| 1 | H | 0.42 | 0/7463 | 0.72 | 7/10171 (0.1%) |
| 1 | I | 0.42 | 0/7414 | 0.70 | 2/10103 (0.0%) |
| 1 | J | 0.40 | 0/7455 | 0.71 | 5/10161 (0.0%) |
| 1 | K | 0.39 | 0/7443 | 0.70 | 10/10143 (0.1%) |
| 1 | L | 0.42 | 1/7438 (0.0%) | 0.70 | 10/10136 (0.1%) |
| 2 | M | 0.35 | 0/3695 | 0.58 | 0/5035 |
| 3 | N | 0.32 | 0/2353 | 0.59 | 2/3206 (0.1%) |
| 4 | O | 0.38 | 0/1417 | 0.54 | 0/1935 |
| 4 | P | 0.38 | 0/1417 | 0.54 | 0/1935 |
| 5 | Q | 0.39 | 0/374 | 0.66 | 0/512 |
| 5 | R | 0.39 | 0/374 | 0.66 | 0/512 |
| 5 | S | 0.39 | 0/374 | 0.66 | 0/512 |
| 5 | T | 0.39 | 0/374 | 0.66 | 0/512 |
| 6 | U | 0.31 | 0/224 | 0.54 | 0/302 |
| 6 | V | 0.31 | 0/224 | 0.54 | 0/302 |
| 6 | Y | 0.31 | 0/224 | 0.54 | 0/302 |
| 6 | u | 0.31 | 0/224 | 0.54 | 0/302 |
| 7 | W | 0.30 | 0/37 | 0.53 | 0/47 |
| 7 | w | 0.33 | 0/86 | 0.55 | 0/114 |
| All | All | 0.40 | 1/100706 (0.0%) | 0.69 | 89/137240 (0.1%) |

Chiral center outliers are detected by calculating the chiral volume of a chiral center and verifying if the center is modelled as a planar moiety or with the opposite hand.A planarity outlier is detected by checking planarity of atoms in a peptide group, atoms in a mainchain group or atoms of a sidechain that are expected to be planar.

CONFIDENTIAL VALIDATION REPORT

| Mol | Chain | #Chirality outliers | #Planarity outliers |
|---|---|---|---|
| 1 | A | 0 | 4 |
| 1 | B | 0 | 6 |
| 1 | C | 0 | 8 |
| 1 | D | 0 | 2 |
| 1 | E | 0 | 7 |
| 1 | F | 0 | 3 |
| 1 | G | 0 | 4 |
| 1 | H | 0 | 4 |
| 1 | I | 0 | 5 |
| 1 | J | 0 | 6 |
| 1 | K | 0 | 8 |
| 1 | L | 0 | 3 |
| 2 | M | 0 | 3 |
| All | All | 0 | 63 |

All (1) bond length outliers are listed below:

| Mol | Chain | Res | Type | Atoms | Z | Observed(Å) | Ideal(Å) |
|---|---|---|---|---|---|---|---|
| 1 | L | 547 | LEU | C-N | -5.50 | 1.21 | 1.34 |

All (89) bond angle outliers are listed below:

| Mol | Chain | Res | Type | Atoms | Z | Observed($^o$) | Ideal($^o$) |
|---|---|---|---|---|---|---|---|
| 1 | G | 274 | LEU | CA-CB-CG | 7.96 | 133.61 | 115.30 |
| 1 | J | 348 | LEU | CA-CB-CG | 7.79 | 133.21 | 115.30 |
| 1 | L | 274 | LEU | CA-CB-CG | 7.74 | 133.11 | 115.30 |
| 1 | C | 744 | LEU | CA-CB-CG | 7.71 | 133.04 | 115.30 |
| 1 | D | 274 | LEU | CA-CB-CG | 7.50 | 132.54 | 115.30 |
| 1 | C | 260 | LEU | CA-CB-CG | 7.44 | 132.41 | 115.30 |
| 1 | J | 260 | LEU | CA-CB-CG | 7.29 | 132.06 | 115.30 |
| 1 | C | 86 | LEU | CA-CB-CG | 7.06 | 131.54 | 115.30 |
| 1 | B | 86 | LEU | CA-CB-CG | 7.03 | 131.47 | 115.30 |
| 1 | A | 892 | LEU | CA-CB-CG | 6.74 | 130.80 | 115.30 |
| 1 | K | 276 | TYR | C-N-CA | -6.71 | 104.94 | 121.70 |
| 1 | G | 348 | LEU | CA-CB-CG | 6.56 | 130.39 | 115.30 |
| 1 | B | 744 | LEU | CA-CB-CG | 6.56 | 130.39 | 115.30 |
| 1 | F | 203 | LEU | CA-CB-CG | 6.55 | 130.35 | 115.30 |
| 1 | G | 353 | LEU | CA-CB-CG | 6.44 | 130.11 | 115.30 |
| 1 | I | 353 | LEU | CA-CB-CG | 6.37 | 129.94 | 115.30 |
| 1 | D | 823 | LEU | CA-CB-CG | 6.30 | 129.80 | 115.30 |
| 1 | D | 348 | LEU | CA-CB-CG | 6.28 | 129.74 | 115.30 |
| 1 | D | 516 | LEU | CA-CB-CG | 6.27 | 129.73 | 115.30 |
| 1 | F | 664 | LEU | CA-CB-CG | 6.18 | 129.52 | 115.30 |

*Continued on next page...*

CONFIDENTIAL VALIDATION REPORT

*Continued from previous page...*

| Mol | Chain | Res | Type | Atoms | Z | Observed(°) | Ideal(°) |
|---|---|---|---|---|---|---|---|
| 1 | K | 350 | LEU | CA-CB-CG | 6.15 | 129.45 | 115.30 |
| 1 | A | 348 | LEU | CA-CB-CG | 6.15 | 129.44 | 115.30 |
| 1 | J | 86 | LEU | CA-CB-CG | 6.13 | 129.41 | 115.30 |
| 1 | F | 312 | LEU | CA-CB-CG | 6.11 | 129.36 | 115.30 |
| 1 | H | 452 | LEU | CA-CB-CG | 6.11 | 129.35 | 115.30 |
| 1 | H | 86 | LEU | CA-CB-CG | 6.02 | 129.14 | 115.30 |
| 1 | E | 433 | MET | CB-CG-SD | 6.01 | 130.44 | 112.40 |
| 1 | F | 348 | LEU | CA-CB-CG | 6.01 | 129.11 | 115.30 |
| 1 | D | 874 | LEU | CA-CB-CG | 5.97 | 129.02 | 115.30 |
| 1 | C | 203 | LEU | CA-CB-CG | 5.96 | 129.00 | 115.30 |
| 1 | H | 626 | LEU | CA-CB-CG | 5.93 | 128.94 | 115.30 |
| 1 | K | 888 | LEU | CA-CB-CG | 5.92 | 128.92 | 115.30 |
| 1 | I | 86 | LEU | CA-CB-CG | 5.84 | 128.73 | 115.30 |
| 1 | C | 290 | LEU | CA-CB-CG | 5.79 | 128.62 | 115.30 |
| 3 | N | 145 | LEU | CA-CB-CG | 5.78 | 128.60 | 115.30 |
| 1 | K | 708 | ASP | CB-CG-OD1 | 5.78 | 123.50 | 118.30 |
| 1 | J | 203 | LEU | CA-CB-CG | 5.75 | 128.53 | 115.30 |
| 1 | H | 309 | PHE | CB-CG-CD1 | 5.67 | 124.77 | 120.80 |
| 1 | H | 181 | PRO | C-N-CA | 5.66 | 135.86 | 121.70 |
| 1 | B | 655 | ARG | NE-CZ-NH2 | -5.63 | 117.48 | 120.30 |
| 1 | L | 344 | LEU | CA-CB-CG | 5.62 | 128.23 | 115.30 |
| 1 | D | 744 | LEU | CA-CB-CG | 5.62 | 128.22 | 115.30 |
| 1 | G | 344 | LEU | CA-CB-CG | 5.62 | 128.22 | 115.30 |
| 1 | C | 710 | LEU | CA-CB-CG | 5.59 | 128.16 | 115.30 |
| 1 | E | 350 | LEU | CA-CB-CG | 5.58 | 128.13 | 115.30 |
| 1 | D | 290 | LEU | CA-CB-CG | 5.52 | 127.99 | 115.30 |
| 1 | F | 519 | ARG | NE-CZ-NH2 | -5.49 | 117.56 | 120.30 |
| 1 | G | 41 | LEU | CA-CB-CG | 5.46 | 127.85 | 115.30 |
| 1 | K | 740 | LEU | CA-CB-CG | 5.45 | 127.84 | 115.30 |
| 1 | A | 708 | ASP | CB-CG-OD1 | 5.44 | 123.20 | 118.30 |
| 1 | E | 348 | LEU | CA-CB-CG | 5.39 | 127.70 | 115.30 |
| 1 | E | 522 | LEU | CA-CB-CG | 5.39 | 127.69 | 115.30 |
| 3 | N | 45 | LEU | CA-CB-CG | 5.37 | 127.66 | 115.30 |
| 1 | F | 290 | LEU | CA-CB-CG | 5.37 | 127.66 | 115.30 |
| 1 | K | 244 | LEU | CA-CB-CG | 5.37 | 127.65 | 115.30 |
| 1 | K | 889 | LEU | CA-CB-CG | 5.36 | 127.64 | 115.30 |
| 1 | E | 244 | LEU | CA-CB-CG | 5.36 | 127.62 | 115.30 |
| 1 | E | 874 | LEU | CA-CB-CG | 5.34 | 127.59 | 115.30 |
| 1 | G | 740 | LEU | CA-CB-CG | 5.33 | 127.56 | 115.30 |
| 1 | H | 892 | LEU | CA-CB-CG | 5.30 | 127.50 | 115.30 |
| 1 | K | 94 | LEU | CA-CB-CG | 5.29 | 127.48 | 115.30 |
| 1 | L | 452 | LEU | CA-CB-CG | 5.27 | 127.42 | 115.30 |

*Continued on next page...*

CONFIDENTIAL VALIDATION REPORT

Continued from previous page...

| Mol | Chain | Res | Type | Atoms | Z | Observed(°) | Ideal(°) |
|---|---|---|---|---|---|---|---|
| 1 | A | 744 | LEU | CA-CB-CG | 5.26 | 127.39 | 115.30 |
| 1 | L | 312 | LEU | CA-CB-CG | 5.24 | 127.36 | 115.30 |
| 1 | G | 94 | LEU | CA-CB-CG | 5.22 | 127.32 | 115.30 |
| 1 | K | 274 | LEU | CA-CB-CG | 5.22 | 127.31 | 115.30 |
| 1 | E | 344 | LEU | CA-CB-CG | 5.21 | 127.28 | 115.30 |
| 1 | L | 629 | ILE | CG1-CB-CG2 | -5.21 | 99.94 | 111.40 |
| 1 | L | 626 | LEU | CA-CB-CG | 5.19 | 127.23 | 115.30 |
| 1 | L | 348 | LEU | CA-CB-CG | 5.17 | 127.18 | 115.30 |
| 1 | E | 740 | LEU | CA-CB-CG | 5.16 | 127.17 | 115.30 |
| 1 | L | 86 | LEU | CA-CB-CG | 5.16 | 127.17 | 115.30 |
| 1 | A | 545 | LEU | CA-CB-CG | 5.15 | 127.15 | 115.30 |
| 1 | D | 711 | LEU | CA-CB-CG | 5.15 | 127.14 | 115.30 |
| 1 | K | 874 | LEU | CA-CB-CG | 5.15 | 127.14 | 115.30 |
| 1 | E | 516 | LEU | CA-CB-CG | 5.13 | 127.09 | 115.30 |
| 1 | G | 516 | LEU | CA-CB-CG | 5.13 | 127.09 | 115.30 |
| 1 | D | 861 | LEU | CA-CB-CG | 5.10 | 127.03 | 115.30 |
| 1 | L | 874 | LEU | CA-CB-CG | 5.10 | 127.02 | 115.30 |
| 1 | E | 86 | LEU | CA-CB-CG | 5.08 | 126.97 | 115.30 |
| 1 | G | 279 | ASP | CB-CG-OD1 | 5.07 | 122.87 | 118.30 |
| 1 | F | 386 | LEU | CA-CB-CG | 5.07 | 126.96 | 115.30 |
| 1 | E | 620 | LEU | CA-CB-CG | 5.06 | 126.94 | 115.30 |
| 1 | C | 620 | LEU | CA-CB-CG | 5.05 | 126.92 | 115.30 |
| 1 | J | 711 | LEU | CA-CB-CG | 5.04 | 126.90 | 115.30 |
| 1 | H | 344 | LEU | CA-CB-CG | 5.04 | 126.89 | 115.30 |
| 1 | C | 253 | PRO | C-N-CA | 5.04 | 134.29 | 121.70 |
| 1 | L | 276 | TYR | C-N-CA | -5.04 | 109.11 | 121.70 |
| 1 | B | 892 | LEU | CA-CB-CG | 5.03 | 126.88 | 115.30 |

There are no chirality outliers.

All (63) planarity outliers are listed below:

| Mol | Chain | Res | Type | Group |
|---|---|---|---|---|
| 1 | A | 312 | LEU | Peptide |
| 1 | A | 612 | ASN | Peptide |
| 1 | A | 704 | TRP | Peptide |
| 1 | A | 756 | VAL | Peptide |
| 1 | B | 334 | VAL | Peptide |
| 1 | B | 409 | GLY | Peptide |
| 1 | B | 612 | ASN | Peptide |
| 1 | B | 704 | TRP | Peptide |
| 1 | B | 796 | HIS | Peptide |
| 1 | B | 916 | THR | Peptide |

Continued on next page...

CONFIDENTIAL VALIDATION REPORT

*Continued from previous page...*

| Mol | Chain | Res | Type | Group |
|---|---|---|---|---|
| 1 | C | 167 | THR | Peptide |
| 1 | C | 285 | ILE | Peptide |
| 1 | C | 525 | ASN | Peptide |
| 1 | C | 59 | ASP | Peptide |
| 1 | C | 612 | ASN | Peptide |
| 1 | C | 630 | PRO | Peptide |
| 1 | C | 762 | ASP | Peptide |
| 1 | C | 796 | HIS | Peptide |
| 1 | D | 630 | PRO | Peptide |
| 1 | D | 641 | PRO | Peptide |
| 1 | E | 149 | PHE | Peptide |
| 1 | E | 285 | ILE | Peptide |
| 1 | E | 314 | TYR | Peptide |
| 1 | E | 377 | ILE | Peptide |
| 1 | E | 70 | PRO | Peptide |
| 1 | E | 757 | PRO | Peptide |
| 1 | E | 916 | THR | Peptide |
| 1 | F | 397 | ALA | Peptide |
| 1 | F | 419 | ALA | Peptide |
| 1 | F | 617 | ASN | Peptide |
| 1 | G | 273 | HIS | Peptide |
| 1 | G | 286 | SER | Peptide |
| 1 | G | 536 | PRO | Peptide |
| 1 | G | 916 | THR | Peptide |
| 1 | H | 159 | GLY | Peptide |
| 1 | H | 319 | GLY | Peptide |
| 1 | H | 354 | GLY | Peptide |
| 1 | H | 401 | THR | Peptide |
| 1 | I | 384 | ASP | Peptide |
| 1 | I | 536 | PRO | Peptide |
| 1 | I | 618 | ASP | Peptide |
| 1 | I | 70 | PRO | Peptide |
| 1 | I | 796 | HIS | Peptide |
| 1 | J | 224 | GLY | Peptide |
| 1 | J | 292 | THR | Peptide |
| 1 | J | 536 | PRO | Peptide |
| 1 | J | 55 | ASP | Peptide |
| 1 | J | 704 | TRP | Peptide |
| 1 | J | 787 | TYR | Peptide |
| 1 | K | 149 | PHE | Peptide |
| 1 | K | 251 | SER | Peptide |
| 1 | K | 273 | HIS | Peptide |

*Continued on next page...*

CONFIDENTIAL VALIDATION REPORT

*Continued from previous page...*

| Mol | Chain | Res | Type | Group |
|---|---|---|---|---|
| 1 | K | 288 | ALA | Peptide |
| 1 | K | 289 | ASP | Peptide |
| 1 | K | 536 | PRO | Peptide |
| 1 | K | 704 | TRP | Peptide |
| 1 | K | 796 | HIS | Peptide |
| 1 | L | 174 | ASP | Peptide |
| 1 | L | 630 | PRO | Peptide |
| 1 | L | 641 | PRO | Peptide |
| 2 | M | 100 | THR | Peptide |
| 2 | M | 331 | LEU | Peptide |
| 2 | M | 62 | TYR | Peptide |

#### 5.2 Too-close contacts ⓘ

In the following table, the Non-H and H(model) columns list the number of non-hydrogen atoms and hydrogen atoms in the chain respectively. The H(added) column lists the number of hydrogen atoms added and optimized by MolProbity. The Clashes column lists the number of clashes within the asymmetric unit, whereas Symm-Clashes lists symmetry related clashes.

| Mol | Chain | Non-H | H(model) | H(added) | Clashes | Symm-Clashes |
|---|---|---|---|---|---|---|
| 1 | A | 7244 | 0 | 6936 | 157 | 0 |
| 1 | B | 7250 | 0 | 6942 | 138 | 0 |
| 1 | C | 7254 | 0 | 6947 | 149 | 0 |
| 1 | D | 7214 | 0 | 6905 | 159 | 0 |
| 1 | E | 7273 | 0 | 6963 | 159 | 0 |
| 1 | F | 7229 | 0 | 6923 | 149 | 0 |
| 1 | G | 7229 | 0 | 6923 | 151 | 0 |
| 1 | H | 7262 | 0 | 6959 | 144 | 0 |
| 1 | I | 7214 | 0 | 6905 | 157 | 0 |
| 1 | J | 7254 | 0 | 6947 | 134 | 0 |
| 1 | K | 7242 | 0 | 6936 | 144 | 0 |
| 1 | L | 7238 | 0 | 6930 | 144 | 0 |
| 2 | M | 3609 | 0 | 3546 | 50 | 0 |
| 3 | N | 2315 | 0 | 2312 | 29 | 0 |
| 4 | O | 1379 | 0 | 1307 | 41 | 0 |
| 4 | P | 1379 | 0 | 1307 | 24 | 0 |
| 5 | Q | 365 | 0 | 363 | 4 | 0 |
| 5 | R | 365 | 0 | 363 | 0 | 0 |
| 5 | S | 365 | 0 | 363 | 2 | 0 |
| 5 | T | 365 | 0 | 363 | 2 | 0 |
| 6 | U | 218 | 0 | 202 | 4 | 0 |
| 6 | V | 218 | 0 | 202 | 1 | 0 |

*Continued on next page...*

CONFIDENTIAL VALIDATION REPORT

*Continued from previous page...*

| Mol | Chain | Non-H | H(model) | H(added) | Clashes | Symm-Clashes |
|---|---|---|---|---|---|---|
| 6 | Y | 218 | 0 | 202 | 6 | 0 |
| 6 | u | 218 | 0 | 202 | 0 | 0 |
| 7 | W | 37 | 0 | 31 | 3 | 0 |
| 7 | w | 83 | 0 | 70 | 0 | 0 |
| All | All | 98037 | 0 | 94049 | 1624 | 0 |

The all-atom clashscore is defined as the number of clashes found per 1000 atoms (including hydrogen atoms). The all-atom clashscore for this structure is 8.

All (1624) close contacts within the same asymmetric unit are listed below, sorted by their clash magnitude.

| Atom-1 | Atom-2 | Interatomic distance (Å) | Clash overlap (Å) |
|---|---|---|---|
| 1:E:754:PHE:HB2 | 1:F:363:TRP:HE1 | 1.43 | 0.84 |
| 1:A:201:ARG:H | 1:C:813:GLN:HE22 | 1.35 | 0.75 |
| 1:D:754:PHE:HB2 | 1:E:363:TRP:HE1 | 1.51 | 0.74 |
| 1:G:74:GLU:HB3 | 1:G:81:LYS:HB2 | 1.71 | 0.73 |
| 1:D:360:PHE:HB3 | 1:D:365:GLN:HB3 | 1.72 | 0.72 |
| 1:J:731:GLN:HE21 | 1:K:533:ILE:HG22 | 1.55 | 0.72 |
| 1:G:310:ILE:HG22 | 1:G:538:LYS:HE3 | 1.72 | 0.71 |
| 1:K:356:ARG:HG3 | 1:K:367:VAL:HG12 | 1.73 | 0.71 |
| 1:B:111:PRO:HA | 1:B:527:ARG:HH12 | 1.56 | 0.70 |
| 1:D:643:ARG:NH2 | 1:I:699:ASP:O | 2.24 | 0.70 |
| 1:J:777:GLN:HG2 | 1:K:526:GLY:HA3 | 1.72 | 0.70 |
| 1:A:512:ARG:HA | 1:A:517:ARG:HH21 | 1.56 | 0.69 |
| 1:L:651:TRP:HZ3 | 1:L:696:ILE:HG12 | 1.58 | 0.69 |
| 1:G:449:ASN:HD21 | 1:G:512:ARG:HH21 | 1.38 | 0.69 |
| 2:M:250:ARG:HH21 | 2:M:372:GLN:HE21 | 1.40 | 0.69 |
| 1:E:248:ALA:HB2 | 1:E:256:PRO:HA | 1.75 | 0.69 |
| 1:D:341:ASN:OD1 | 1:D:624:ASN:ND2 | 2.26 | 0.69 |
| 1:A:483:VAL:HG22 | 1:A:485:SER:H | 1.58 | 0.69 |
| 1:E:906:ARG:NH2 | 1:L:328:ALA:O | 2.26 | 0.69 |
| 1:A:68:PHE:HB2 | 1:A:588:LEU:HB3 | 1.73 | 0.69 |
| 1:G:81:LYS:HD2 | 1:G:557:ASN:HB3 | 1.73 | 0.68 |
| 1:I:114:LYS:NZ | 1:I:270:PRO:O | 2.26 | 0.68 |
| 1:D:322:GLY:HA3 | 1:D:556:TRP:HD1 | 1.59 | 0.68 |
| 1:H:725:GLU:HB3 | 1:I:67:ARG:HH12 | 1.58 | 0.68 |
| 3:N:204:LEU:HD21 | 4:O:201:SER:HB3 | 1.76 | 0.67 |
| 1:F:101:PHE:HB2 | 1:F:535:VAL:HB | 1.76 | 0.67 |
| 1:D:405:ILE:HG22 | 1:D:414:ALA:HA | 1.77 | 0.66 |
| 1:I:643:ARG:NH2 | 1:L:699:ASP:O | 2.29 | 0.66 |
| 1:J:310:ILE:HG22 | 1:J:538:LYS:HE3 | 1.76 | 0.66 |

*Continued on next page...*

CONFIDENTIAL VALIDATION REPORT

Continued from previous page...

| Atom-1 | Atom-2 | Interatomic distance (Å) | Clash overlap (Å) |
|---|---|---|---|
| 1:B:306:ARG:HH12 | 1:B:678:VAL:H | 1.44 | 0.66 |
| 1:K:701:SER:OG | 4:O:16:GLN:NE2 | 2.29 | 0.66 |
| 1:C:217:ALA:O | 1:C:226:GLN:NE2 | 2.29 | 0.65 |
| 1:D:374:VAL:HG23 | 1:D:450:VAL:HG22 | 1.79 | 0.65 |
| 1:H:652:SER:HB2 | 1:H:892:LEU:HB3 | 1.78 | 0.65 |
| 1:G:44:LYS:HE2 | 1:I:546:LEU:HB2 | 1.78 | 0.65 |
| 1:D:13:MET:HG3 | 1:F:914:LEU:HD21 | 1.76 | 0.65 |
| 1:D:201:ARG:HG2 | 1:D:261:TYR:HB2 | 1.78 | 0.65 |
| 1:J:67:ARG:HH12 | 1:L:725:GLU:HG3 | 1.62 | 0.65 |
| 1:A:402:TYR:HB3 | 1:C:247:PHE:HB3 | 1.77 | 0.65 |
| 1:E:749:ILE:HG22 | 1:E:752:GLN:HE21 | 1.61 | 0.65 |
| 1:H:19:ASP:HA | 1:H:48:PRO:HD2 | 1.78 | 0.65 |
| 1:K:107:LEU:HD11 | 1:K:566:LEU:HD11 | 1.78 | 0.65 |
| 1:B:18:GLN:HE22 | 4:O:185:ARG:HA | 1.62 | 0.65 |
| 1:K:174:ASP:H | 1:K:178:GLN:HE21 | 1.43 | 0.65 |
| 1:E:629:ILE:HG12 | 1:E:636:VAL:HG11 | 1.79 | 0.65 |
| 1:K:624:ASN:ND2 | 1:K:890:TYR:OH | 2.30 | 0.65 |
| 1:K:94:LEU:HB3 | 1:K:547:LEU:HB3 | 1.79 | 0.65 |
| 1:K:914:LEU:HD12 | 4:O:31:MET:HG2 | 1.79 | 0.65 |
| 1:D:382:VAL:HG11 | 1:D:439:ALA:HA | 1.79 | 0.64 |
| 1:I:563:ASN:ND2 | 1:I:672:PHE:O | 2.30 | 0.64 |
| 1:I:360:PHE:HB3 | 1:I:365:GLN:HB3 | 1.79 | 0.64 |
| 1:A:423:ALA:HB1 | 1:C:143:VAL:H | 1.63 | 0.64 |
| 1:B:509:ASN:ND2 | 1:B:674:TYR:OH | 2.30 | 0.64 |
| 1:K:509:ASN:ND2 | 1:K:674:TYR:OH | 2.31 | 0.64 |
| 1:A:495:ALA:HB2 | 1:B:525:ASN:HB2 | 1.80 | 0.64 |
| 1:J:388:ASN:HB3 | 1:K:440:ASN:HD21 | 1.63 | 0.64 |
| 1:K:132:PRO:HB2 | 1:K:202:VAL:HG12 | 1.79 | 0.64 |
| 1:D:12:TYR:O | 1:F:900:ARG:NH2 | 2.31 | 0.64 |
| 1:H:732:CYS:HB2 | 1:H:837:PHE:HB3 | 1.80 | 0.64 |
| 1:E:495:ALA:HB2 | 1:F:525:ASN:HB2 | 1.79 | 0.64 |
| 1:E:786:GLU:HG2 | 1:F:218:LYS:HD3 | 1.80 | 0.63 |
| 1:J:134:GLU:HG2 | 1:J:143:VAL:HG12 | 1.80 | 0.63 |
| 1:K:360:PHE:HB3 | 1:K:365:GLN:HB3 | 1.80 | 0.63 |
| 1:G:217:ALA:HB3 | 1:G:226:GLN:HE21 | 1.61 | 0.63 |
| 1:G:374:VAL:HG11 | 1:G:510:HIS:HE1 | 1.63 | 0.63 |
| 1:G:124:LEU:H | 1:I:798:ASN:HD21 | 1.44 | 0.63 |
| 1:J:447:TYR:HA | 1:J:451:ALA:HB3 | 1.80 | 0.63 |
| 1:A:152:THR:OG1 | 1:A:161:GLN:NE2 | 2.31 | 0.63 |
| 1:F:135:TRP:HE1 | 1:F:142:LYS:HE3 | 1.64 | 0.63 |
| 1:G:545:LEU:HD13 | 1:G:901:ILE:HD12 | 1.80 | 0.63 |

Continued on next page...

CONFIDENTIAL VALIDATION REPORT

*Continued from previous page...*

| Atom-1 | Atom-2 | Interatomic distance (Å) | Clash overlap (Å) |
|---|---|---|---|
| 1:A:341:ASN:H | 1:A:624:ASN:HD22 | 1.45 | 0.63 |
| 1:D:612:ASN:HD21 | 1:E:25:SER:H | 1.46 | 0.63 |
| 1:F:459:LYS:HB3 | 1:F:480:ARG:HB3 | 1.81 | 0.63 |
| 1:J:302:TYR:OH | 1:J:517:ARG:NH1 | 2.32 | 0.63 |
| 1:K:563:ASN:ND2 | 1:K:672:PHE:O | 2.31 | 0.63 |
| 1:A:307:ASP:HA | 1:A:519:ARG:HH12 | 1.64 | 0.63 |
| 1:F:474:TYR:HB2 | 1:F:572:ASN:HD22 | 1.63 | 0.63 |
| 1:A:688:ASN:ND2 | 1:A:842:THR:O | 2.32 | 0.63 |
| 1:G:134:GLU:HG2 | 1:G:143:VAL:HG22 | 1.81 | 0.62 |
| 1:C:845:ARG:HH22 | 6:Y:32:GLY:HA3 | 1.64 | 0.62 |
| 1:H:74:GLU:HB3 | 1:H:81:LYS:HB3 | 1.80 | 0.62 |
| 1:J:306:ARG:HE | 1:J:567:GLN:HB2 | 1.65 | 0.62 |
| 1:I:643:ARG:HH22 | 1:L:699:ASP:HB3 | 1.65 | 0.62 |
| 1:D:649:ARG:NH2 | 7:W:21:TYR:O | 2.33 | 0.62 |
| 1:C:315:TYR:HE2 | 1:C:342:THR:HG23 | 1.64 | 0.62 |
| 1:E:340:ARG:HH21 | 1:E:915:ARG:HD3 | 1.64 | 0.62 |
| 1:F:218:LYS:NZ | 1:F:268:GLU:OE2 | 2.33 | 0.62 |
| 1:F:408:ASN:HB3 | 1:F:411:THR:HB | 1.81 | 0.62 |
| 1:G:506:ASN:ND2 | 1:G:509:ASN:OD1 | 2.33 | 0.62 |
| 1:J:752:GLN:NE2 | 1:L:39:PHE:O | 2.33 | 0.62 |
| 1:E:356:ARG:HB3 | 1:E:367:VAL:HG11 | 1.81 | 0.62 |
| 1:C:699:ASP:O | 1:E:643:ARG:NH2 | 2.32 | 0.62 |
| 1:J:274:LEU:HD21 | 1:J:277:LYS:HD3 | 1.81 | 0.62 |
| 1:C:459:LYS:HG2 | 1:C:482:ALA:HB2 | 1.81 | 0.61 |
| 1:D:558:PHE:HE2 | 1:D:586:ILE:HG21 | 1.62 | 0.61 |
| 1:E:911:ALA:HB3 | 4:O:109:ALA:HB3 | 1.82 | 0.61 |
| 1:A:559:ARG:NH1 | 1:A:564:MET:SD | 2.73 | 0.61 |
| 1:L:104:ARG:HG3 | 1:L:584:ASP:HB3 | 1.81 | 0.61 |
| 1:E:146:GLN:NE2 | 1:F:429:ASN:OD1 | 2.33 | 0.61 |
| 1:G:483:VAL:HG22 | 1:G:485:SER:H | 1.64 | 0.61 |
| 1:A:430:ILE:O | 1:B:810:ARG:NH2 | 2.34 | 0.61 |
| 1:G:788:GLN:HB3 | 1:H:222:GLU:HA | 1.81 | 0.61 |
| 1:J:111:PRO:HD2 | 1:J:577:ASP:HB3 | 1.83 | 0.61 |
| 1:C:546:LEU:HD21 | 1:C:603:LEU:HD21 | 1.82 | 0.61 |
| 1:G:376:ILE:HD13 | 1:G:496:ARG:HE | 1.65 | 0.61 |
| 1:I:732:CYS:SG | 1:I:733:ASN:N | 2.73 | 0.61 |
| 1:L:375:ARG:HD3 | 1:L:507:PRO:HG3 | 1.81 | 0.61 |
| 1:D:146:GLN:HE22 | 1:E:430:ILE:HG13 | 1.66 | 0.61 |
| 1:J:657:LYS:HG2 | 1:J:887:THR:HG22 | 1.83 | 0.61 |
| 1:K:58:THR:HG22 | 1:K:60:ARG:H | 1.65 | 0.61 |
| 1:A:818:ASN:OD1 | 1:B:226:GLN:NE2 | 2.34 | 0.61 |

*Continued on next page...*

CONFIDENTIAL VALIDATION REPORT

Continued from previous page...

| Atom-1 | Atom-2 | Interatomic distance (Å) | Clash overlap (Å) |
|---|---|---|---|
| 1:B:388:ASN:HB3 | 1:C:440:ASN:HD21 | 1.65 | 0.61 |
| 1:K:203:LEU:HD12 | 1:K:207:THR:HG21 | 1.83 | 0.61 |
| 4:P:7:THR:O | 4:P:25:GLN:NE2 | 2.34 | 0.61 |
| 1:D:813:GLN:HE22 | 1:E:201:ARG:H | 1.49 | 0.61 |
| 1:B:80:TYR:HB3 | 1:B:558:PHE:HB2 | 1.83 | 0.60 |
| 1:C:483:VAL:HG11 | 1:C:792:LEU:HD21 | 1.82 | 0.60 |
| 1:E:563:ASN:ND2 | 1:E:672:PHE:O | 2.34 | 0.60 |
| 1:H:459:LYS:HB3 | 1:H:480:ARG:HB3 | 1.83 | 0.60 |
| 1:L:623:ALA:HB2 | 1:L:915:ARG:HH22 | 1.64 | 0.60 |
| 1:C:509:ASN:ND2 | 1:C:674:TYR:OH | 2.33 | 0.60 |
| 1:G:306:ARG:HE | 1:G:567:GLN:HB2 | 1.66 | 0.60 |
| 1:L:565:ILE:HG23 | 1:L:566:LEU:HG | 1.82 | 0.60 |
| 4:P:7:THR:HB | 4:P:25:GLN:HE21 | 1.66 | 0.60 |
| 1:A:104:ARG:NH2 | 1:C:724:GLY:O | 2.33 | 0.60 |
| 1:D:301:ASN:HA | 1:D:569:SER:HB2 | 1.84 | 0.60 |
| 1:I:548:LEU:HD11 | 1:I:607:LEU:HB2 | 1.83 | 0.60 |
| 1:A:436:ASN:ND2 | 1:B:123:SER:O | 2.35 | 0.60 |
| 1:G:268:GLU:HG3 | 1:G:270:PRO:HD3 | 1.82 | 0.60 |
| 1:G:247:PHE:HA | 1:H:404:GLY:HA2 | 1.84 | 0.60 |
| 4:O:7:THR:O | 4:O:25:GLN:NE2 | 2.34 | 0.60 |
| 1:G:204:LYS:HD2 | 1:G:264:ASN:HB2 | 1.83 | 0.60 |
| 1:D:307:ASP:OD2 | 1:D:356:ARG:NH2 | 2.34 | 0.60 |
| 1:D:591:ASN:ND2 | 1:F:750:GLY:O | 2.35 | 0.60 |
| 1:J:3:THR:HG22 | 1:J:5:SER:H | 1.66 | 0.60 |
| 2:M:371:SER:HB3 | 2:M:405:LEU:HB3 | 1.82 | 0.60 |
| 3:N:239:ASN:N | 4:O:4:GLU:OE2 | 2.35 | 0.60 |
| 1:C:132:PRO:HB3 | 1:C:145:GLY:HA2 | 1.82 | 0.60 |
| 1:L:643:ARG:HE | 1:L:918:PHE:H | 1.50 | 0.60 |
| 1:C:130:PRO:HA | 1:C:212:CYS:H | 1.67 | 0.59 |
| 1:D:762:ASP:HB3 | 1:D:769:ARG:HB2 | 1.84 | 0.59 |
| 1:G:559:ARG:NH1 | 1:G:564:MET:SD | 2.75 | 0.59 |
| 1:I:209:MET:O | 1:I:293:GLN:NE2 | 2.34 | 0.59 |
| 1:K:401:THR:HG23 | 1:K:423:ALA:H | 1.66 | 0.59 |
| 1:L:583:PHE:HB3 | 1:L:586:ILE:HD11 | 1.83 | 0.59 |
| 1:D:210:LEU:HD12 | 1:D:267:ILE:HG22 | 1.83 | 0.59 |
| 1:D:363:TRP:HE1 | 1:F:754:PHE:HB2 | 1.66 | 0.59 |
| 1:E:459:LYS:O | 1:E:480:ARG:NH2 | 2.35 | 0.59 |
| 1:I:52:PRO:HD3 | 6:U:11:PRO:HA | 86.46 | 0.59 |
| 1:F:175:LYS:HB2 | 1:F:242:VAL:HG21 | 1.84 | 0.59 |
| 1:H:558:PHE:HE2 | 1:H:586:ILE:HG12 | 1.66 | 0.59 |
| 1:E:226:GLN:HG3 | 1:E:265:VAL:HG11 | 1.85 | 0.59 |

Continued on next page...

CONFIDENTIAL VALIDATION REPORT

Continued from previous page...

| Atom-1 | Atom-2 | Interatomic distance (Å) | Clash overlap (Å) |
|---|---|---|---|
| 1:G:380:HIS:HE1 | 1:H:517:ARG:HB2 | 1.67 | 0.59 |
| 1:A:900:ARG:NH1 | 1:B:12:TYR:O | 2.34 | 0.59 |
| 1:C:496:ARG:HH21 | 1:C:772:GLN:HE21 | 1.50 | 0.59 |
| 1:D:688:ASN:ND2 | 1:D:842:THR:O | 2.36 | 0.59 |
| 1:I:161:GLN:NE2 | 1:I:163:GLY:O | 2.35 | 0.59 |
| 1:J:335:VAL:O | 1:J:340:ARG:NH2 | 2.35 | 0.59 |
| 1:K:613:ASP:OD2 | 1:K:900:ARG:NE | 2.36 | 0.59 |
| 1:B:563:ASN:ND2 | 1:B:672:PHE:O | 2.36 | 0.59 |
| 1:D:33:ARG:NH2 | 6:V:20:GLY:O | 2.35 | 0.59 |
| 1:J:163:GLY:HA3 | 1:J:171:ILE:HB | 1.85 | 0.59 |
| 1:I:649:ARG:NH2 | 6:U:32:GLY:O | 85.58 | 0.59 |
| 1:C:52:PRO:HD3 | 6:Y:11:PRO:HA | 1.84 | 0.59 |
| 1:B:508:PHE:O | 1:B:513:ASN:ND2 | 2.35 | 0.59 |
| 1:E:73:ARG:HH12 | 1:E:585:SER:HA | 1.68 | 0.59 |
| 1:H:326:GLY:HA2 | 1:H:552:TYR:HA | 1.85 | 0.59 |
| 1:D:575:ARG:HD3 | 5:S:40:VAL:HA | 1.84 | 0.59 |
| 1:C:114:LYS:HE3 | 1:C:298:ASN:HB2 | 1.85 | 0.59 |
| 1:F:368:ASP:O | 1:F:841:ARG:NH2 | 2.36 | 0.59 |
| 1:J:174:ASP:HB3 | 1:J:177:TYR:HB3 | 1.85 | 0.59 |
| 1:K:114:LYS:NZ | 1:K:116:TYR:O | 2.34 | 0.59 |
| 4:O:7:THR:HB | 4:O:25:GLN:HE21 | 1.66 | 0.59 |
| 1:C:624:ASN:HD21 | 1:C:892:LEU:HD12 | 1.68 | 0.58 |
| 1:H:621:CYS:O | 1:H:649:ARG:NH2 | 2.36 | 0.58 |
| 1:A:725:GLU:O | 1:B:104:ARG:NH2 | 2.36 | 0.58 |
| 1:D:436:ASN:ND2 | 1:E:123:SER:O | 2.36 | 0.58 |
| 1:E:632:ASN:ND2 | 1:E:883:MET:O | 2.36 | 0.58 |
| 2:M:142:LYS:NZ | 2:M:248:GLN:OE1 | 2.35 | 0.58 |
| 1:F:496:ARG:NH1 | 1:F:773:PRO:O | 2.36 | 0.58 |
| 1:H:146:GLN:HA | 1:I:426:GLU:HB3 | 1.84 | 0.58 |
| 1:C:108:ASP:O | 1:C:527:ARG:NH1 | 2.35 | 0.58 |
| 1:C:503:ASP:OD2 | 1:C:689:HIS:NE2 | 2.36 | 0.58 |
| 1:K:736:LYS:NZ | 1:L:65:THR:OG1 | 2.36 | 0.58 |
| 1:A:629:ILE:HD11 | 1:A:889:LEU:HB2 | 1.86 | 0.58 |
| 1:I:565:ILE:HD11 | 1:I:581:VAL:HG11 | 1.85 | 0.58 |
| 1:G:86:LEU:HB2 | 1:G:552:TYR:HB2 | 1.84 | 0.58 |
| 1:H:337:LEU:HD21 | 1:H:920:ALA:HB2 | 1.86 | 0.58 |
| 1:G:188:TRP:HB3 | 1:I:809:MET:HE1 | 1.85 | 0.58 |
| 1:J:546:LEU:HB3 | 1:J:614:GLN:HE22 | 1.69 | 0.58 |
| 1:L:458:TYR:O | 1:L:480:ARG:NH2 | 2.36 | 0.58 |
| 1:B:101:PHE:HB2 | 1:B:535:VAL:HB | 1.85 | 0.58 |
| 1:B:511:HIS:O | 1:B:517:ARG:NH1 | 2.36 | 0.58 |

Continued on next page...

CONFIDENTIAL VALIDATION REPORT

*Continued from previous page...*

| Atom-1 | Atom-2 | Interatomic distance (Å) | Clash overlap (Å) |
|---|---|---|---|
| 1:B:627:TYR:HB2 | 1:B:889:LEU:HB3 | 1.86 | 0.58 |
| 1:D:476:TYR:OH | 1:D:480:ARG:NH1 | 2.36 | 0.58 |
| 1:H:305:PHE:HZ | 1:H:312:LEU:HD22 | 1.69 | 0.58 |
| 1:I:350:LEU:HD21 | 1:I:365:GLN:HG3 | 1.85 | 0.58 |
| 1:I:437:LEU:HA | 1:I:440:ASN:HD22 | 1.69 | 0.58 |
| 1:I:912:VAL:HG12 | 4:P:108:LEU:HD22 | 1.86 | 0.58 |
| 1:G:823:LEU:HB3 | 1:H:526:GLY:HA2 | 1.85 | 0.58 |
| 1:J:107:LEU:HD23 | 1:J:109:ARG:HE | 1.69 | 0.58 |
| 1:A:563:ASN:HB2 | 1:A:575:ARG:HB2 | 1.85 | 0.58 |
| 1:E:144:ARG:NH2 | 1:F:394:GLY:O | 2.37 | 0.58 |
| 1:I:607:LEU:HD23 | 1:I:612:ASN:HB3 | 1.86 | 0.58 |
| 1:K:516:LEU:HA | 1:K:519:ARG:HE | 1.68 | 0.58 |
| 1:K:74:GLU:HB3 | 1:K:81:LYS:HB2 | 1.85 | 0.58 |
| 1:C:443:ARG:NH1 | 1:C:488:ASP:OD2 | 2.35 | 0.57 |
| 1:F:376:ILE:HD11 | 1:F:772:GLN:HE22 | 1.69 | 0.57 |
| 1:H:206:THR:OG1 | 1:H:264:ASN:ND2 | 2.37 | 0.57 |
| 1:H:318:THR:HA | 1:H:321:MET:HB3 | 1.86 | 0.57 |
| 1:A:563:ASN:ND2 | 1:A:672:PHE:O | 2.36 | 0.57 |
| 1:B:900:ARG:NH1 | 1:C:12:TYR:O | 2.37 | 0.57 |
| 1:B:276:TYR:OH | 1:C:187:GLN:NE2 | 2.38 | 0.57 |
| 1:A:46:ARG:NH1 | 1:C:617:ASN:OD1 | 2.38 | 0.57 |
| 1:C:655:ARG:HE | 1:C:887:THR:HG21 | 1.69 | 0.57 |
| 1:D:437:LEU:HB2 | 1:F:437:LEU:HD11 | 1.84 | 0.57 |
| 1:H:101:PHE:HB2 | 1:H:535:VAL:HB | 1.86 | 0.57 |
| 1:H:508:PHE:O | 1:H:513:ASN:ND2 | 2.37 | 0.57 |
| 1:G:363:TRP:NE1 | 1:I:754:PHE:O | 2.36 | 0.57 |
| 1:G:649:ARG:HH21 | 1:G:894:GLU:HG2 | 1.70 | 0.57 |
| 2:M:342:SER:HB3 | 2:M:345:LEU:HB2 | 1.85 | 0.57 |
| 1:A:74:GLU:HG3 | 1:A:81:LYS:HE2 | 1.85 | 0.57 |
| 1:D:347:GLN:NE2 | 1:D:682:ASP:OD1 | 2.31 | 0.57 |
| 1:G:652:SER:HB2 | 1:G:892:LEU:HB3 | 1.86 | 0.57 |
| 1:I:269:ALA:HB1 | 1:I:272:THR:HB | 1.85 | 0.57 |
| 2:M:144:ARG:HE | 2:M:242:CYS:HA | 1.70 | 0.57 |
| 2:M:397:TYR:OH | 2:M:477:ARG:NH1 | 2.37 | 0.57 |
| 3:N:147:SER:HB3 | 3:N:180:VAL:HB | 1.85 | 0.57 |
| 1:J:709:ARG:NH2 | 4:P:233:ASP:OD2 | 2.38 | 0.57 |
| 1:C:90:ASP:H | 1:C:906:ARG:HH12 | 1.50 | 0.57 |
| 1:E:204:LYS:HD3 | 1:E:264:ASN:HB2 | 1.85 | 0.57 |
| 1:A:160:ILE:HB | 1:A:259:VAL:HG11 | 1.86 | 0.57 |
| 1:A:509:ASN:ND2 | 1:A:674:TYR:OH | 2.38 | 0.57 |
| 1:D:130:PRO:HA | 1:D:212:CYS:H | 1.69 | 0.57 |

*Continued on next page...*

CONFIDENTIAL VALIDATION REPORT

*Continued from previous page...*

| Atom-1 | Atom-2 | Interatomic distance (Å) | Clash overlap (Å) |
|---|---|---|---|
| 1:D:340:ARG:NH2 | 1:D:622:ALA:O | 2.38 | 0.57 |
| 1:D:278:PRO:O | 1:D:464:ASN:ND2 | 2.38 | 0.57 |
| 1:F:313:MET:HE1 | 1:F:537:GLN:HA | 1.86 | 0.57 |
| 1:J:306:ARG:NH2 | 1:J:564:MET:O | 2.37 | 0.57 |
| 1:A:405:ILE:HG12 | 1:C:246:PHE:HB2 | 1.87 | 0.57 |
| 1:D:222:GLU:HG3 | 1:D:223:LYS:HG2 | 1.87 | 0.57 |
| 1:F:155:ASN:ND2 | 1:F:172:TYR:OH | 2.38 | 0.57 |
| 1:F:202:VAL:HB | 1:F:260:LEU:HD13 | 1.85 | 0.57 |
| 1:I:450:VAL:HG12 | 1:I:487:LEU:HD13 | 1.87 | 0.57 |
| 1:K:697:MET:HG2 | 1:K:703:SER:HA | 1.86 | 0.57 |
| 1:L:73:ARG:NH1 | 1:L:80:TYR:OH | 2.37 | 0.57 |
| 1:B:537:GLN:NE2 | 1:B:554:TYR:OH | 2.37 | 0.57 |
| 1:C:629:ILE:HD11 | 1:C:889:LEU:HB2 | 1.87 | 0.57 |
| 1:J:413:THR:OG1 | 1:J:414:ALA:N | 2.38 | 0.57 |
| 1:E:92:ARG:NH1 | 1:E:595:MET:O | 2.38 | 0.56 |
| 1:E:62:GLN:OE1 | 1:E:92:ARG:NH2 | 2.38 | 0.56 |
| 1:H:449:ASN:OD1 | 1:H:510:HIS:NE2 | 2.38 | 0.56 |
| 1:L:278:PRO:O | 1:L:464:ASN:ND2 | 2.38 | 0.56 |
| 2:M:205:LYS:HE2 | 2:M:442:LEU:HA | 1.86 | 0.56 |
| 1:A:382:VAL:HG11 | 1:A:439:ALA:HA | 1.87 | 0.56 |
| 1:A:717:GLU:OE1 | 1:A:720:ARG:NH2 | 2.38 | 0.56 |
| 1:C:108:ASP:OD1 | 1:C:527:ARG:NH1 | 2.38 | 0.56 |
| 1:D:94:LEU:HD23 | 1:D:547:LEU:HD22 | 1.86 | 0.56 |
| 1:H:201:ARG:NH2 | 1:H:263:GLU:OE1 | 2.38 | 0.56 |
| 1:G:788:GLN:NE2 | 1:H:220:THR:O | 2.37 | 0.56 |
| 1:G:182:GLN:NE2 | 1:I:793:PRO:O | 2.37 | 0.56 |
| 1:G:146:GLN:HE21 | 1:I:814:ALA:H | 1.53 | 0.56 |
| 1:A:521:MET:HB3 | 1:C:496:ARG:HB2 | 1.86 | 0.56 |
| 1:A:617:ASN:ND2 | 7:W:20:MET:SD | 82.37 | 0.56 |
| 1:B:766:SER:O | 1:B:770:ASN:ND2 | 2.37 | 0.56 |
| 1:D:340:ARG:HH21 | 1:D:622:ALA:H | 1.52 | 0.56 |
| 1:G:208:PRO:HB3 | 1:G:284:THR:HG22 | 1.87 | 0.56 |
| 1:H:301:ASN:HA | 1:H:569:SER:HB2 | 1.88 | 0.56 |
| 1:F:12:TYR:HH | 4:O:167:SER:HG | 1.52 | 0.56 |
| 1:F:158:ASN:ND2 | 1:F:173:ALA:O | 2.37 | 0.56 |
| 1:F:154:ILE:HD11 | 1:F:256:PRO:HB2 | 1.86 | 0.56 |
| 1:G:219:PRO:O | 1:I:788:GLN:NE2 | 2.38 | 0.56 |
| 1:G:474:TYR:HB2 | 1:G:572:ASN:HD22 | 1.70 | 0.56 |
| 1:G:607:LEU:HD21 | 1:G:614:GLN:HE22 | 1.70 | 0.56 |
| 1:J:94:LEU:HD12 | 1:J:592:PHE:HE1 | 1.70 | 0.56 |
| 1:L:717:GLU:HG2 | 1:L:735:THR:HG21 | 1.87 | 0.56 |

*Continued on next page...*

CONFIDENTIAL VALIDATION REPORT

*Continued from previous page...*

| Atom-1 | Atom-2 | Interatomic distance (Å) | Clash overlap (Å) |
|---|---|---|---|
| 1:L:650:GLY:H | 1:L:848:PHE:H | 1.54 | 0.56 |
| 1:E:315:TYR:OH | 1:E:342:THR:O | 2.22 | 0.56 |
| 1:E:899:VAL:HA | 1:E:913:TYR:HB3 | 1.87 | 0.56 |
| 1:J:723:ASP:OD2 | 1:K:104:ARG:NH1 | 2.36 | 0.56 |
| 1:B:94:LEU:HD23 | 1:B:547:LEU:HD22 | 1.87 | 0.56 |
| 1:D:25:SER:H | 1:F:612:ASN:HD21 | 1.54 | 0.56 |
| 1:G:132:PRO:HB2 | 1:G:202:VAL:HG22 | 1.88 | 0.56 |
| 1:K:709:ARG:HB3 | 1:L:64:LEU:HD13 | 1.88 | 0.56 |
| 1:L:503:ASP:OD1 | 1:L:689:HIS:NE2 | 2.38 | 0.56 |
| 2:M:49:GLY:O | 2:M:114:ARG:NH1 | 2.38 | 0.56 |
| 1:B:705:PRO:HB3 | 1:B:710:LEU:HD22 | 1.86 | 0.56 |
| 1:C:161:GLN:HA | 1:C:172:TYR:HA | 1.88 | 0.56 |
| 1:C:360:PHE:HB3 | 1:C:365:GLN:HB3 | 1.88 | 0.56 |
| 1:I:136:LYS:HA | 1:I:141:ILE:HG22 | 1.87 | 0.56 |
| 1:L:150:ILE:HD12 | 1:L:185:GLN:HE21 | 1.70 | 0.56 |
| 1:B:485:SER:O | 1:B:490:TYR:OH | 2.22 | 0.56 |
| 1:F:86:LEU:HB2 | 1:F:552:TYR:HB2 | 1.86 | 0.56 |
| 1:J:742:GLN:HE21 | 1:J:746:HIS:HE1 | 1.54 | 0.56 |
| 1:K:13:MET:O | 1:K:46:ARG:NH2 | 2.38 | 0.56 |
| 4:P:14:GLN:NE2 | 4:P:16:GLN:OE1 | 2.39 | 0.56 |
| 1:C:447:TYR:HA | 1:C:451:ALA:HB3 | 1.87 | 0.56 |
| 1:C:94:LEU:HB3 | 1:C:547:LEU:HB3 | 1.88 | 0.56 |
| 1:L:109:ARG:NH1 | 1:L:303:ILE:O | 2.39 | 0.56 |
| 3:N:56:GLN:O | 3:N:61:ARG:NH1 | 2.38 | 0.56 |
| 4:O:14:GLN:NE2 | 4:O:16:GLN:OE1 | 2.39 | 0.56 |
| 1:E:230:ILE:HG22 | 1:E:239:THR:H | 1.71 | 0.56 |
| 1:G:517:ARG:HH11 | 1:I:380:HIS:CG | 2.24 | 0.56 |
| 1:G:820:PRO:HG2 | 1:H:121:TYR:HE1 | 1.71 | 0.56 |
| 1:B:629:ILE:HD11 | 1:B:889:LEU:HB2 | 1.88 | 0.55 |
| 1:F:298:ASN:ND2 | 1:F:478:ASN:OD1 | 2.39 | 0.55 |
| 1:A:279:ASP:O | 1:A:464:ASN:ND2 | 2.39 | 0.55 |
| 1:B:632:ASN:ND2 | 1:B:883:MET:O | 2.37 | 0.55 |
| 1:C:563:ASN:ND2 | 1:C:672:PHE:O | 2.39 | 0.55 |
| 1:D:114:LYS:NZ | 1:D:116:TYR:O | 2.37 | 0.55 |
| 1:D:116:TYR:OH | 1:D:271:ASP:OD2 | 2.23 | 0.55 |
| 1:D:306:ARG:NH2 | 1:D:563:ASN:O | 2.40 | 0.55 |
| 1:G:201:ARG:NH2 | 1:G:263:GLU:OE1 | 2.39 | 0.55 |
| 1:G:483:VAL:HG21 | 1:G:792:LEU:HD21 | 1.88 | 0.55 |
| 1:K:388:ASN:HB3 | 1:L:440:ASN:HD21 | 1.71 | 0.55 |
| 1:B:210:LEU:HD12 | 1:B:267:ILE:HG22 | 1.88 | 0.55 |
| 1:B:814:ALA:H | 1:C:146:GLN:HE21 | 1.53 | 0.55 |

*Continued on next page...*

CONFIDENTIAL VALIDATION REPORT

Continued from previous page...

| Atom-1 | Atom-2 | Interatomic distance (Å) | Clash overlap (Å) |
|---|---|---|---|
| 1:K:180:GLU:HG2 | 1:K:182:GLN:H | 1.71 | 0.55 |
| 1:K:462:PRO:HG2 | 1:K:465:ILE:HD12 | 1.89 | 0.55 |
| 1:K:814:ALA:H | 1:L:146:GLN:HE21 | 1.54 | 0.55 |
| 2:M:389:ARG:HD3 | 2:M:390:PRO:HD2 | 1.89 | 0.55 |
| 1:A:123:SER:O | 1:C:436:ASN:ND2 | 2.39 | 0.55 |
| 1:B:263:GLU:HB3 | 1:B:265:VAL:HG13 | 1.87 | 0.55 |
| 1:C:732:CYS:SG | 1:C:733:ASN:N | 2.79 | 0.55 |
| 1:C:763:ARG:HE | 1:C:764:MET:H | 1.53 | 0.55 |
| 1:D:898:VAL:HG22 | 1:E:46:ARG:HH12 | 1.72 | 0.55 |
| 1:A:503:ASP:OD1 | 1:A:689:HIS:NE2 | 2.36 | 0.55 |
| 1:C:776:ARG:HD3 | 1:C:834:GLN:HE21 | 1.71 | 0.55 |
| 1:E:519:ARG:NH2 | 1:E:567:GLN:OE1 | 2.40 | 0.55 |
| 1:E:85:THR:OG1 | 1:L:83:ARG:NH2 | 2.40 | 0.55 |
| 1:G:626:LEU:HG | 1:G:664:LEU:HD11 | 1.89 | 0.55 |
| 1:J:652:SER:HB2 | 1:J:892:LEU:HB3 | 1.88 | 0.55 |
| 1:A:350:LEU:HD22 | 1:A:356:ARG:HH11 | 1.71 | 0.55 |
| 1:B:155:ASN:ND2 | 1:B:172:TYR:OH | 2.40 | 0.55 |
| 1:C:349:MET:SD | 1:C:543:LYS:NZ | 2.76 | 0.55 |
| 1:F:447:TYR:HA | 1:F:451:ALA:HB3 | 1.88 | 0.55 |
| 1:H:472:ASN:HB3 | 1:H:573:ASP:HB2 | 1.87 | 0.55 |
| 1:J:356:ARG:HB3 | 1:J:367:VAL:HG11 | 1.89 | 0.55 |
| 1:L:351:ASP:HB2 | 1:L:356:ARG:HE | 1.72 | 0.55 |
| 1:A:72:ASP:HB3 | 1:A:83:ARG:HD3 | 1.89 | 0.55 |
| 1:C:176:THR:HG22 | 1:C:225:GLY:HA3 | 1.89 | 0.55 |
| 1:G:347:GLN:NE2 | 1:G:682:ASP:O | 2.40 | 0.55 |
| 1:G:537:GLN:NE2 | 1:G:542:ILE:O | 2.40 | 0.55 |
| 1:H:392:PRO:HA | 1:I:433:MET:HG2 | 1.87 | 0.55 |
| 1:H:710:LEU:HD11 | 1:H:741:ILE:HD12 | 1.89 | 0.55 |
| 1:A:399:THR:HB | 1:A:426:GLU:HA | 1.89 | 0.55 |
| 1:D:613:ASP:HB2 | 1:D:900:ARG:HH11 | 1.72 | 0.55 |
| 1:G:548:LEU:HD11 | 1:G:607:LEU:HB3 | 1.88 | 0.55 |
| 1:G:786:GLU:HB2 | 1:G:788:GLN:HE22 | 1.72 | 0.55 |
| 1:K:620:LEU:HD21 | 1:K:892:LEU:HD21 | 1.89 | 0.55 |
| 1:A:15:ILE:HA | 1:A:46:ARG:HH21 | 1.71 | 0.54 |
| 1:B:811:GLU:HB2 | 1:C:184:GLY:HA3 | 1.89 | 0.54 |
| 1:E:436:ASN:ND2 | 1:F:123:SER:O | 2.37 | 0.54 |
| 1:G:490:TYR:OH | 1:G:795:GLN:NE2 | 2.40 | 0.54 |
| 1:I:575:ARG:NH1 | 5:Q:38:SER:O | 2.40 | 0.54 |
| 1:J:642:SER:HB3 | 1:J:871:ALA:HB1 | 1.89 | 0.54 |
| 3:N:238:PRO:HG2 | 4:O:5:ILE:HD11 | 1.89 | 0.54 |
| 1:B:559:ARG:HG2 | 1:B:561:ASP:H | 1.72 | 0.54 |

Continued on next page...

CONFIDENTIAL VALIDATION REPORT

*Continued from previous page...*

| Atom-1 | Atom-2 | Interatomic distance (Å) | Clash overlap (Å) |
|---|---|---|---|
| 1:D:65:THR:HG22 | 1:D:591:ASN:HB3 | 1.89 | 0.54 |
| 1:E:402:TYR:HB2 | 1:E:423:ALA:HB3 | 1.88 | 0.54 |
| 1:G:583:PHE:HB3 | 1:G:586:ILE:HD11 | 1.89 | 0.54 |
| 1:G:630:PRO:HB2 | 1:G:633:ALA:HB3 | 1.90 | 0.54 |
| 1:G:440:ASN:HD21 | 1:I:389:TYR:H | 1.53 | 0.54 |
| 1:J:388:ASN:ND2 | 1:K:440:ASN:OD1 | 2.40 | 0.54 |
| 1:A:810:ARG:NH2 | 1:C:430:ILE:O | 2.40 | 0.54 |
| 1:H:563:ASN:ND2 | 1:H:672:PHE:O | 2.40 | 0.54 |
| 1:J:107:LEU:HD11 | 1:J:566:LEU:HD21 | 1.88 | 0.54 |
| 1:E:24:LEU:HD22 | 1:E:28:LEU:HD23 | 1.89 | 0.54 |
| 1:E:472:ASN:HB3 | 1:E:573:ASP:HB2 | 1.88 | 0.54 |
| 1:F:548:LEU:HD11 | 1:F:607:LEU:HB2 | 1.89 | 0.54 |
| 1:C:548:LEU:HD12 | 1:C:903:GLN:HB2 | 1.89 | 0.54 |
| 1:H:657:LYS:HG2 | 1:H:887:THR:HG22 | 1.88 | 0.54 |
| 1:L:568:SER:OG | 1:L:675:SER:O | 2.26 | 0.54 |
| 1:E:640:ILE:HD13 | 1:E:643:ARG:HH12 | 1.72 | 0.54 |
| 1:E:76:THR:HB | 1:E:79:SER:H | 1.72 | 0.54 |
| 1:E:624:ASN:ND2 | 1:E:680:TYR:OH | 2.41 | 0.54 |
| 1:E:762:ASP:OD2 | 1:F:358:ARG:NH1 | 2.41 | 0.54 |
| 1:G:83:ARG:HA | 1:G:555:GLU:HA | 1.90 | 0.54 |
| 1:J:382:VAL:HG11 | 1:J:439:ALA:HA | 1.88 | 0.54 |
| 1:D:631:SER:OG | 5:Q:11:SER:O | 2.25 | 0.54 |
| 1:B:382:VAL:HG23 | 1:B:384:ASP:HB2 | 1.90 | 0.54 |
| 1:C:301:ASN:HA | 1:C:569:SER:HB2 | 1.88 | 0.54 |
| 1:C:376:ILE:HD11 | 1:C:838:LEU:HG | 1.89 | 0.54 |
| 1:D:378:GLU:HG2 | 1:D:496:ARG:HA | 1.90 | 0.54 |
| 1:F:152:THR:OG1 | 1:F:161:GLN:NE2 | 2.37 | 0.54 |
| 1:F:688:ASN:ND2 | 1:F:842:THR:O | 2.40 | 0.54 |
| 1:H:546:LEU:HB2 | 1:I:44:LYS:HE2 | 1.88 | 0.54 |
| 1:J:104:ARG:NH2 | 1:J:585:SER:OG | 2.41 | 0.54 |
| 1:K:292:THR:HG21 | 1:L:187:GLN:HA | 1.90 | 0.54 |
| 1:B:523:LEU:HB3 | 1:B:529:VAL:HG11 | 1.89 | 0.54 |
| 1:D:146:GLN:HG3 | 1:E:426:GLU:HB3 | 1.90 | 0.54 |
| 1:J:94:LEU:HB3 | 1:J:547:LEU:HB2 | 1.89 | 0.54 |
| 1:L:710:LEU:HD23 | 1:L:712:THR:H | 1.72 | 0.54 |
| 1:A:229:LEU:HA | 1:A:240:SER:HA | 1.89 | 0.54 |
| 1:C:780:ASN:HD21 | 1:C:832:LEU:HB2 | 1.73 | 0.54 |
| 1:E:812:GLY:H | 1:F:184:GLY:HA3 | 1.73 | 0.54 |
| 1:L:744:LEU:HD21 | 1:L:853:MET:HG2 | 1.90 | 0.54 |
| 1:A:780:ASN:ND2 | 1:A:783:THR:OG1 | 2.41 | 0.53 |
| 1:H:313:MET:HG2 | 1:H:556:TRP:HE1 | 1.72 | 0.53 |

*Continued on next page...*

CONFIDENTIAL VALIDATION REPORT

*Continued from previous page...*

| Atom-1 | Atom-2 | Interatomic distance (Å) | Clash overlap (Å) |
|---|---|---|---|
| 1:H:512:ARG:HG3 | 1:H:517:ARG:HH11 | 1.73 | 0.53 |
| 1:L:632:ASN:HD22 | 1:L:882:PRO:HB2 | 1.73 | 0.53 |
| 1:A:226:GLN:HG3 | 1:A:265:VAL:HG11 | 1.90 | 0.53 |
| 1:C:780:ASN:HD21 | 1:C:832:LEU:H | 1.56 | 0.53 |
| 1:D:798:ASN:HD21 | 1:E:124:LEU:HB2 | 1.73 | 0.53 |
| 1:G:390:CYS:HB2 | 1:G:433:MET:HB2 | 1.90 | 0.53 |
| 1:L:435:ILE:HD13 | 1:L:437:LEU:HB3 | 1.89 | 0.53 |
| 1:E:910:GLU:OE2 | 4:O:93:ARG:NH1 | 2.40 | 0.53 |
| 1:A:210:LEU:HD12 | 1:A:267:ILE:HG22 | 1.88 | 0.53 |
| 1:A:197:LYS:NZ | 1:A:256:PRO:O | 2.40 | 0.53 |
| 1:A:275:VAL:HA | 1:A:465:ILE:HD11 | 1.89 | 0.53 |
| 1:C:150:ILE:HG13 | 1:C:198:VAL:HG23 | 1.91 | 0.53 |
| 1:C:327:GLN:O | 1:C:330:GLN:NE2 | 2.41 | 0.53 |
| 1:C:565:ILE:HG23 | 1:C:566:LEU:HG | 1.89 | 0.53 |
| 1:C:73:ARG:HH12 | 1:C:585:SER:HA | 1.71 | 0.53 |
| 1:D:341:ASN:ND2 | 1:D:680:TYR:O | 2.41 | 0.53 |
| 1:E:299:ARG:NH1 | 1:E:477:MET:O | 2.40 | 0.53 |
| 1:F:374:VAL:HG13 | 1:F:449:ASN:HB3 | 1.90 | 0.53 |
| 1:H:688:ASN:OD1 | 1:H:844:TRP:NE1 | 2.42 | 0.53 |
| 1:J:396:SER:OG | 1:L:144:ARG:NH2 | 2.42 | 0.53 |
| 1:J:688:ASN:ND2 | 1:J:842:THR:O | 2.42 | 0.53 |
| 2:M:258:ARG:HH21 | 2:M:363:THR:HG21 | 1.73 | 0.53 |
| 1:A:360:PHE:HB3 | 1:A:365:GLN:HB2 | 1.91 | 0.53 |
| 1:C:352:ALA:HA | 1:C:763:ARG:HD3 | 1.90 | 0.53 |
| 1:H:467:LEU:HD12 | 1:H:476:TYR:HA | 1.89 | 0.53 |
| 1:I:23:TYR:OH | 1:I:45:PHE:O | 2.25 | 0.53 |
| 1:B:342:THR:HG21 | 1:B:539:PHE:HA | 1.89 | 0.53 |
| 1:D:23:TYR:OH | 1:D:45:PHE:O | 2.27 | 0.53 |
| 1:G:620:LEU:HD12 | 1:G:892:LEU:HD21 | 1.90 | 0.53 |
| 1:H:456:ASP:O | 1:H:480:ARG:NH2 | 2.42 | 0.53 |
| 2:M:81:ASN:HD22 | 2:M:90:GLN:H | 1.56 | 0.53 |
| 1:C:770:ASN:OD1 | 1:C:841:ARG:NH2 | 2.41 | 0.53 |
| 1:D:764:MET:O | 1:D:770:ASN:ND2 | 2.42 | 0.53 |
| 1:G:380:HIS:CE1 | 1:H:517:ARG:HB2 | 2.44 | 0.53 |
| 1:L:175:LYS:HB3 | 1:L:242:VAL:HG11 | 1.90 | 0.53 |
| 1:L:692:LYS:HB2 | 1:L:879:GLU:HG3 | 1.91 | 0.53 |
| 1:L:742:GLN:NE2 | 1:L:845:ARG:O | 2.39 | 0.53 |
| 1:B:230:ILE:O | 1:B:238:LEU:N | 2.42 | 0.53 |
| 1:D:496:ARG:NH1 | 1:D:773:PRO:O | 2.41 | 0.53 |
| 1:D:63:ARG:NH1 | 1:F:710:LEU:O | 2.42 | 0.53 |
| 1:E:335:VAL:O | 1:E:913:TYR:OH | 2.24 | 0.53 |

*Continued on next page...*

CONFIDENTIAL VALIDATION REPORT

*Continued from previous page...*

| Atom-1 | Atom-2 | Interatomic distance (Å) | Clash overlap (Å) |
|---|---|---|---|
| 1:E:654:THR:OG1 | 1:E:688:ASN:ND2 | 2.42 | 0.53 |
| 1:H:621:CYS:SG | 1:H:649:ARG:NH2 | 2.81 | 0.53 |
| 1:A:134:GLU:HG2 | 1:A:143:VAL:HG12 | 1.89 | 0.53 |
| 1:A:220:THR:HG21 | 1:A:228:SER:HA | 1.91 | 0.53 |
| 1:F:563:ASN:ND2 | 1:F:672:PHE:O | 2.42 | 0.53 |
| 1:I:301:ASN:HA | 1:I:569:SER:HB3 | 1.91 | 0.53 |
| 1:B:910:GLU:OE1 | 1:C:2:ALA:N | 2.42 | 0.53 |
| 1:C:209:MET:O | 1:C:293:GLN:NE2 | 2.42 | 0.53 |
| 1:G:279:ASP:HA | 1:G:464:ASN:HD21 | 1.74 | 0.53 |
| 1:H:754:PHE:O | 1:I:363:TRP:NE1 | 2.37 | 0.53 |
| 2:M:324:LYS:HD3 | 2:M:326:ARG:HH12 | 1.72 | 0.53 |
| 1:J:598:ASN:ND2 | 4:P:93:ARG:O | 2.36 | 0.53 |
| 1:A:694:VAL:HB | 1:A:716:PHE:HB2 | 1.89 | 0.52 |
| 1:B:565:ILE:HG23 | 1:B:566:LEU:HG | 1.91 | 0.52 |
| 1:D:19:ASP:HA | 1:D:48:PRO:HD2 | 1.91 | 0.52 |
| 1:F:332:ASN:ND2 | 1:F:334:VAL:O | 2.41 | 0.52 |
| 1:F:58:THR:HG22 | 1:F:60:ARG:H | 1.74 | 0.52 |
| 1:J:632:ASN:ND2 | 1:J:883:MET:O | 2.42 | 0.52 |
| 1:K:310:ILE:HD12 | 1:K:681:LEU:HB3 | 1.90 | 0.52 |
| 1:C:350:LEU:HB2 | 1:C:356:ARG:HH21 | 1.74 | 0.52 |
| 1:C:485:SER:O | 1:C:490:TYR:OH | 2.26 | 0.52 |
| 1:D:386:LEU:HD12 | 1:D:387:PRO:HD2 | 1.91 | 0.52 |
| 1:F:723:ASP:OD1 | 1:F:728:ASN:ND2 | 2.42 | 0.52 |
| 1:G:57:THR:OG1 | 1:I:860:ASP:OD2 | 2.27 | 0.52 |
| 1:G:68:PHE:HB2 | 1:G:588:LEU:HB3 | 1.90 | 0.52 |
| 1:I:308:ASN:HB3 | 1:I:310:ILE:HG23 | 1.91 | 0.52 |
| 1:J:604:GLU:O | 1:J:608:ARG:NH1 | 2.42 | 0.52 |
| 1:K:448:SER:HA | 1:K:452:LEU:HD23 | 1.91 | 0.52 |
| 2:M:144:ARG:NH1 | 2:M:241:GLY:O | 2.41 | 0.52 |
| 1:A:446:LEU:HD11 | 1:A:487:LEU:HG | 1.92 | 0.52 |
| 1:C:101:PHE:HB2 | 1:C:535:VAL:HB | 1.92 | 0.52 |
| 1:C:621:CYS:SG | 1:C:649:ARG:NH1 | 2.78 | 0.52 |
| 1:I:210:LEU:HD22 | 1:I:215:SER:HB3 | 1.90 | 0.52 |
| 1:K:766:SER:O | 1:K:770:ASN:ND2 | 2.42 | 0.52 |
| 1:C:197:LYS:NZ | 1:C:255:GLU:OE2 | 2.40 | 0.52 |
| 1:D:405:ILE:HG12 | 1:F:248:ALA:HB2 | 1.92 | 0.52 |
| 1:D:807:PRO:HB3 | 1:F:386:LEU:HD23 | 1.92 | 0.52 |
| 1:H:460:ILE:HD11 | 1:H:483:VAL:HB | 1.90 | 0.52 |
| 1:H:375:ARG:HH11 | 1:H:507:PRO:HD2 | 1.74 | 0.52 |
| 1:J:103:ILE:HG12 | 1:J:586:ILE:HD12 | 1.91 | 0.52 |
| 1:B:327:GLN:O | 2:M:389:ARG:NH1 | 2.39 | 0.52 |

*Continued on next page...*

CONFIDENTIAL VALIDATION REPORT

*Continued from previous page...*

| Atom-1 | Atom-2 | Interatomic distance (Å) | Clash overlap (Å) |
|---|---|---|---|
| 1:D:402:TYR:HE1 | 1:F:249:LEU:HD23 | 1.75 | 0.52 |
| 1:H:371:ASP:HB3 | 1:H:374:VAL:HG22 | 1.92 | 0.52 |
| 1:I:550:GLY:HA2 | 1:I:906:ARG:HG2 | 1.92 | 0.52 |
| 1:A:504:ASN:ND2 | 1:A:883:MET:SD | 2.78 | 0.52 |
| 1:E:276:TYR:OH | 1:F:187:GLN:NE2 | 2.43 | 0.52 |
| 1:E:910:GLU:OE2 | 4:O:165:ARG:NH1 | 2.42 | 0.52 |
| 1:G:519:ARG:NH1 | 1:G:567:GLN:OE1 | 2.35 | 0.52 |
| 1:H:510:HIS:CD2 | 1:H:512:ARG:H | 2.28 | 0.52 |
| 1:C:454:LEU:HD22 | 1:C:501:PRO:HB2 | 1.92 | 0.52 |
| 1:D:531:PHE:HB2 | 1:D:533:ILE:HG23 | 1.91 | 0.52 |
| 1:F:196:GLN:O | 1:F:257:LYS:NZ | 2.42 | 0.52 |
| 1:G:688:ASN:ND2 | 1:G:842:THR:O | 2.42 | 0.52 |
| 1:H:313:MET:O | 1:H:559:ARG:NH2 | 2.42 | 0.52 |
| 1:H:79:SER:OG | 1:H:80:TYR:N | 2.42 | 0.52 |
| 1:H:765:TYR:HE1 | 1:H:841:ARG:HH11 | 1.58 | 0.52 |
| 1:I:247:PHE:HB2 | 1:I:258:ALA:HB3 | 1.92 | 0.52 |
| 1:E:640:ILE:HD11 | 1:E:874:LEU:HG | 1.92 | 0.52 |
| 1:E:655:ARG:NH2 | 1:E:887:THR:OG1 | 2.42 | 0.52 |
| 1:F:94:LEU:HD23 | 1:F:547:LEU:HD23 | 1.91 | 0.52 |
| 1:K:210:LEU:HD12 | 1:K:267:ILE:HG22 | 1.92 | 0.52 |
| 1:L:402:TYR:HB2 | 1:L:423:ALA:HB3 | 1.92 | 0.52 |
| 2:M:227:TYR:OH | 2:M:280:PRO:O | 2.26 | 0.52 |
| 2:M:321:LYS:HA | 2:M:327:SER:HA | 1.91 | 0.52 |
| 1:B:708:ASP:HB3 | 1:C:61:SER:HB2 | 1.92 | 0.52 |
| 1:E:150:ILE:HG12 | 1:E:198:VAL:HG23 | 1.90 | 0.52 |
| 1:G:742:GLN:HE22 | 1:G:845:ARG:H | 1.56 | 0.52 |
| 1:K:688:ASN:ND2 | 1:K:839:CYS:SG | 2.83 | 0.52 |
| 1:L:423:ALA:HB1 | 1:L:425:ILE:HG23 | 1.91 | 0.52 |
| 1:A:302:TYR:HB3 | 1:A:520:SER:HB3 | 1.91 | 0.52 |
| 1:B:693:LYS:NZ | 1:B:879:GLU:OE1 | 2.42 | 0.52 |
| 1:C:679:PRO:HA | 1:C:684:THR:HB | 1.91 | 0.52 |
| 1:E:910:GLU:HA | 4:O:110:GLY:HA3 | 1.92 | 0.52 |
| 1:F:656:LEU:HD23 | 1:F:685:PHE:HE1 | 1.74 | 0.52 |
| 1:D:102:ASP:OD2 | 1:F:736:LYS:NZ | 2.43 | 0.52 |
| 1:G:517:ARG:HB3 | 1:I:380:HIS:HE1 | 1.75 | 0.52 |
| 1:G:649:ARG:HE | 1:G:894:GLU:HG2 | 1.74 | 0.52 |
| 1:L:537:GLN:NE2 | 1:L:554:TYR:OH | 2.43 | 0.52 |
| 1:C:618:ASP:HB3 | 1:C:621:CYS:HB3 | 1.91 | 0.51 |
| 1:D:10:TRP:HB2 | 1:D:15:ILE:HB | 1.91 | 0.51 |
| 1:D:660:GLU:HB2 | 1:D:678:VAL:HG12 | 1.92 | 0.51 |
| 1:E:107:LEU:HD11 | 1:E:566:LEU:HD21 | 1.92 | 0.51 |

*Continued on next page...*

CONFIDENTIAL VALIDATION REPORT

Continued from previous page...

| Atom-1 | Atom-2 | Interatomic distance (Å) | Clash overlap (Å) |
|---|---|---|---|
| 1:H:385:GLU:HG3 | 1:I:296:ALA:HB1 | 1.93 | 0.51 |
| 1:I:346:TYR:HB3 | 1:I:538:LYS:HD3 | 1.92 | 0.51 |
| 1:L:388:ASN:OD1 | 1:L:388:ASN:N | 2.42 | 0.51 |
| 1:C:382:VAL:HG11 | 1:C:439:ALA:HA | 1.93 | 0.51 |
| 1:C:559:ARG:NH2 | 1:C:564:MET:O | 2.44 | 0.51 |
| 1:C:621:CYS:O | 1:C:649:ARG:NH2 | 2.42 | 0.51 |
| 1:F:217:ALA:HB2 | 1:F:267:ILE:HD13 | 1.92 | 0.51 |
| 1:H:14:HIS:O | 1:H:46:ARG:NH1 | 2.43 | 0.51 |
| 1:G:276:TYR:OH | 1:H:187:GLN:NE2 | 2.43 | 0.51 |
| 3:N:35:TRP:NE1 | 3:N:68:ALA:O | 2.43 | 0.51 |
| 1:B:609:ASN:ND2 | 3:N:24:SER:OG | 2.43 | 0.51 |
| 1:D:202:VAL:HB | 1:D:260:LEU:HD13 | 1.91 | 0.51 |
| 1:G:458:TYR:HE2 | 1:G:501:PRO:HB3 | 1.75 | 0.51 |
| 1:I:366:ALA:O | 1:I:519:ARG:NH1 | 2.43 | 0.51 |
| 1:J:165:ASP:HB2 | 1:J:169:GLN:HB2 | 1.91 | 0.51 |
| 1:K:562:VAL:HG23 | 1:K:566:LEU:HD12 | 1.92 | 0.51 |
| 1:L:767:PHE:HA | 1:L:842:THR:HG21 | 1.91 | 0.51 |
| 1:L:443:ARG:HH12 | 1:L:802:VAL:HG12 | 1.75 | 0.51 |
| 1:A:766:SER:O | 1:A:770:ASN:ND2 | 2.44 | 0.51 |
| 1:B:645:TRP:HD1 | 1:B:872:HIS:HB2 | 1.76 | 0.51 |
| 1:H:352:ALA:HA | 1:H:763:ARG:HD3 | 1.92 | 0.51 |
| 1:I:476:TYR:OH | 1:I:480:ARG:NH2 | 2.43 | 0.51 |
| 3:N:60:ASN:OD1 | 6:Y:12:ARG:NH2 | 2.43 | 0.51 |
| 1:D:46:ARG:HH12 | 1:F:898:VAL:HG22 | 1.74 | 0.51 |
| 1:E:728:ASN:H | 1:E:736:LYS:HE2 | 1.76 | 0.51 |
| 1:E:756:VAL:HG23 | 1:F:365:GLN:HE21 | 1.76 | 0.51 |
| 1:G:503:ASP:OD2 | 1:G:689:HIS:NE2 | 2.41 | 0.51 |
| 1:K:342:THR:HA | 1:K:345:SER:HB3 | 1.92 | 0.51 |
| 1:J:392:PRO:HA | 1:K:433:MET:HG2 | 1.91 | 0.51 |
| 1:K:546:LEU:HB2 | 1:L:44:LYS:HD2 | 1.93 | 0.51 |
| 1:K:632:ASN:ND2 | 1:K:883:MET:O | 2.43 | 0.51 |
| 1:B:116:TYR:OH | 1:B:271:ASP:OD2 | 2.29 | 0.51 |
| 1:G:425:ILE:HG22 | 1:I:145:GLY:HA3 | 1.92 | 0.51 |
| 1:I:542:ILE:HA | 1:I:545:LEU:HD22 | 1.93 | 0.51 |
| 1:J:23:TYR:OH | 1:J:45:PHE:O | 2.28 | 0.51 |
| 1:K:86:LEU:HD12 | 1:K:94:LEU:HD11 | 1.92 | 0.51 |
| 1:A:780:ASN:OD1 | 1:A:781:THR:N | 2.44 | 0.51 |
| 1:A:626:LEU:HB3 | 1:A:888:LEU:HD12 | 1.91 | 0.51 |
| 1:B:53:THR:OG1 | 1:B:54:HIS:N | 2.43 | 0.51 |
| 1:C:180:GLU:HB2 | 1:C:183:VAL:HG23 | 1.92 | 0.51 |
| 1:D:322:GLY:HA3 | 1:D:556:TRP:CD1 | 2.43 | 0.51 |

Continued on next page...

CONFIDENTIAL VALIDATION REPORT

Continued from previous page...

| Atom-1 | Atom-2 | Interatomic distance (Å) | Clash overlap (Å) |
|---|---|---|---|
| 1:D:742:GLN:HE22 | 1:D:767:PHE:HB2 | 1.76 | 0.51 |
| 1:E:726:GLY:HA2 | 1:F:104:ARG:HH22 | 1.76 | 0.51 |
| 1:F:151:GLY:HA3 | 1:F:160:ILE:HD11 | 1.92 | 0.51 |
| 1:G:112:SER:OG | 1:G:572:ASN:ND2 | 2.44 | 0.51 |
| 1:H:657:LYS:HA | 1:H:887:THR:HA | 1.91 | 0.51 |
| 1:J:146:GLN:HA | 1:K:426:GLU:H | 1.75 | 0.51 |
| 1:K:637:PRO:HA | 1:K:877:THR:HA | 1.93 | 0.51 |
| 1:B:450:VAL:HG12 | 1:B:487:LEU:HD21 | 1.93 | 0.51 |
| 1:C:652:SER:HB2 | 1:C:892:LEU:HB3 | 1.92 | 0.51 |
| 1:D:243:ASN:O | 1:D:262:ALA:N | 2.42 | 0.51 |
| 1:D:780:ASN:HD22 | 1:D:832:LEU:HD23 | 1.76 | 0.51 |
| 1:E:292:THR:HG23 | 1:F:188:TRP:HD1 | 1.76 | 0.51 |
| 1:F:68:PHE:HB2 | 1:F:588:LEU:HB3 | 1.93 | 0.51 |
| 1:G:24:LEU:HD12 | 1:G:28:LEU:HD23 | 1.91 | 0.51 |
| 1:I:692:LYS:NZ | 1:I:881:ASP:OD1 | 2.39 | 0.51 |
| 1:K:371:ASP:HB2 | 1:K:374:VAL:HG22 | 1.92 | 0.51 |
| 1:L:903:GLN:HE21 | 1:L:909:ILE:HD12 | 1.75 | 0.51 |
| 1:C:229:LEU:HB2 | 1:C:238:LEU:HD21 | 1.93 | 0.51 |
| 1:D:104:ARG:NH1 | 1:F:723:ASP:OD2 | 2.44 | 0.51 |
| 1:D:622:ALA:HA | 1:D:894:GLU:HA | 1.93 | 0.51 |
| 1:I:404:GLY:HA3 | 1:I:418:TYR:HB2 | 1.92 | 0.51 |
| 1:J:606:MET:O | 1:J:612:ASN:ND2 | 2.43 | 0.51 |
| 2:M:53:ILE:HD13 | 2:M:65:THR:HG22 | 1.92 | 0.51 |
| 1:A:83:ARG:HG3 | 1:A:555:GLU:HG2 | 1.93 | 0.50 |
| 1:E:622:ALA:HA | 1:E:895:VAL:HG22 | 1.93 | 0.50 |
| 1:E:770:ASN:OD1 | 1:E:841:ARG:NH2 | 2.44 | 0.50 |
| 1:G:446:LEU:HD12 | 1:G:487:LEU:HD11 | 1.93 | 0.50 |
| 1:H:821:TYR:HB3 | 1:H:829:VAL:HG21 | 1.92 | 0.50 |
| 4:O:8:PRO:HB3 | 4:O:202:VAL:HG22 | 1.93 | 0.50 |
| 1:C:822:PRO:HG2 | 1:C:827:THR:HG21 | 1.93 | 0.50 |
| 1:G:220:THR:N | 1:G:225:GLY:O | 2.44 | 0.50 |
| 1:H:374:VAL:HG21 | 1:H:510:HIS:HE1 | 1.77 | 0.50 |
| 1:J:343:GLU:HB2 | 1:J:681:LEU:HD22 | 1.93 | 0.50 |
| 1:K:23:TYR:OH | 1:K:45:PHE:O | 2.29 | 0.50 |
| 1:A:187:GLN:HG3 | 1:C:292:THR:HG21 | 1.91 | 0.50 |
| 1:A:546:LEU:HB3 | 1:A:614:GLN:HE22 | 1.75 | 0.50 |
| 1:A:81:LYS:HG2 | 1:A:557:ASN:HD22 | 1.76 | 0.50 |
| 1:A:815:TYR:O | 1:B:122:ASN:ND2 | 2.44 | 0.50 |
| 1:E:203:LEU:HB3 | 1:E:207:THR:HG21 | 1.92 | 0.50 |
| 1:G:302:TYR:HE2 | 1:G:517:ARG:HG2 | 1.76 | 0.50 |
| 1:J:128:THR:HG23 | 1:J:294:GLN:HB3 | 1.92 | 0.50 |

Continued on next page...

CONFIDENTIAL VALIDATION REPORT

Continued from previous page...

| Atom-1 | Atom-2 | Interatomic distance (Å) | Clash overlap (Å) |
|---|---|---|---|
| 1:A:447:TYR:HA | 1:A:451:ALA:HB3 | 1.93 | 0.50 |
| 1:E:909:ILE:O | 4:O:111:GLY:N | 2.41 | 0.50 |
| 1:G:332:ASN:ND2 | 1:G:334:VAL:O | 2.45 | 0.50 |
| 1:K:707:ASN:N | 1:K:707:ASN:OD1 | 2.45 | 0.50 |
| 1:K:791:THR:HG23 | 1:K:793:PRO:HD2 | 1.93 | 0.50 |
| 1:L:206:THR:OG1 | 1:L:264:ASN:OD1 | 2.28 | 0.50 |
| 1:D:103:ILE:HA | 1:D:586:ILE:HG12 | 1.93 | 0.50 |
| 1:I:130:PRO:HA | 1:I:211:PRO:HA | 1.93 | 0.50 |
| 1:J:294:GLN:HE22 | 1:J:808:THR:HA | 1.77 | 0.50 |
| 2:M:189:LEU:HG | 2:M:193:ARG:HH12 | 1.77 | 0.50 |
| 1:G:382:VAL:HG11 | 1:G:439:ALA:HA | 1.93 | 0.50 |
| 1:J:490:TYR:HA | 1:J:493:ILE:HD11 | 1.93 | 0.50 |
| 1:A:217:ALA:HB2 | 1:A:267:ILE:HD13 | 1.94 | 0.50 |
| 1:A:813:GLN:HE21 | 1:B:147:ALA:HA | 1.77 | 0.50 |
| 1:E:548:LEU:HD11 | 1:E:607:LEU:HB3 | 1.94 | 0.50 |
| 1:F:308:ASN:OD1 | 1:F:347:GLN:NE2 | 2.44 | 0.50 |
| 1:G:108:ASP:HB3 | 1:G:579:ALA:HA | 1.94 | 0.50 |
| 1:G:15:ILE:HG12 | 1:I:898:VAL:HG21 | 1.94 | 0.50 |
| 1:B:248:ALA:HB2 | 1:B:256:PRO:HA | 1.94 | 0.50 |
| 1:B:705:PRO:HG2 | 1:B:708:ASP:H | 1.76 | 0.50 |
| 1:D:378:GLU:OE1 | 1:E:517:ARG:NH1 | 2.45 | 0.50 |
| 1:F:542:ILE:HG22 | 1:F:545:LEU:HD13 | 1.92 | 0.50 |
| 1:H:143:VAL:H | 1:I:423:ALA:HB3 | 1.76 | 0.50 |
| 1:K:380:HIS:NE2 | 1:L:521:MET:SD | 2.85 | 0.50 |
| 3:N:215:ASN:ND2 | 3:N:237:THR:OG1 | 2.45 | 0.50 |
| 1:A:706:GLY:O | 1:A:709:ARG:NH1 | 2.45 | 0.50 |
| 1:C:339:ASP:OD2 | 1:C:915:ARG:NH2 | 2.44 | 0.50 |
| 1:D:316:ASN:ND2 | 1:D:337:LEU:O | 2.45 | 0.50 |
| 1:G:638:ILE:HD11 | 1:G:891:VAL:HG21 | 1.93 | 0.50 |
| 1:H:210:LEU:HD21 | 1:H:269:ALA:HB2 | 1.94 | 0.50 |
| 1:J:150:ILE:HG12 | 1:J:198:VAL:HG23 | 1.93 | 0.50 |
| 1:J:88:VAL:HG13 | 1:J:549:PRO:HA | 1.94 | 0.50 |
| 1:J:549:PRO:HD2 | 1:J:608:ARG:HH22 | 1.77 | 0.50 |
| 1:K:914:LEU:HD13 | 1:L:13:MET:HG3 | 1.93 | 0.50 |
| 1:K:390:CYS:HB3 | 1:L:435:ILE:HG23 | 1.93 | 0.50 |
| 3:N:99:ALA:HA | 3:N:102:ILE:HG22 | 1.94 | 0.50 |
| 1:A:78:TYR:OH | 1:K:723:ASP:OD2 | 2.29 | 0.49 |
| 1:E:208:PRO:HG2 | 1:E:267:ILE:HB | 1.93 | 0.49 |
| 1:F:116:TYR:OH | 1:F:271:ASP:OD2 | 2.27 | 0.49 |
| 1:F:278:PRO:HG3 | 1:F:293:GLN:H | 1.77 | 0.49 |
| 1:F:354:GLY:O | 1:F:763:ARG:NH2 | 2.45 | 0.49 |

Continued on next page...

CONFIDENTIAL VALIDATION REPORT

Continued from previous page...

| Atom-1 | Atom-2 | Interatomic distance (Å) | Clash overlap (Å) |
|---|---|---|---|
| 1:G:476:TYR:OH | 1:G:480:ARG:NH1 | 2.45 | 0.49 |
| 1:G:562:VAL:HG21 | 1:G:579:ALA:HB3 | 1.93 | 0.49 |
| 1:H:724:GLY:O | 1:I:104:ARG:NH1 | 2.44 | 0.49 |
| 1:I:629:ILE:HD13 | 1:I:880:VAL:HG21 | 1.94 | 0.49 |
| 1:L:310:ILE:HG22 | 1:L:538:LYS:HE2 | 1.94 | 0.49 |
| 3:N:62:PHE:HA | 3:N:65:ILE:HG22 | 1.93 | 0.49 |
| 1:A:390:CYS:HA | 1:B:435:ILE:HB | 1.94 | 0.49 |
| 1:C:302:TYR:HE2 | 1:C:511:HIS:HB2 | 1.76 | 0.49 |
| 1:D:710:LEU:HD21 | 1:D:741:ILE:HD13 | 1.93 | 0.49 |
| 1:D:797:ASN:HD21 | 1:E:121:TYR:HA | 1.75 | 0.49 |
| 1:F:24:LEU:HB2 | 1:F:29:VAL:HG13 | 1.93 | 0.49 |
| 1:H:58:THR:HB | 1:H:596:ALA:HA | 1.94 | 0.49 |
| 1:H:645:TRP:HD1 | 1:H:872:HIS:HB2 | 1.77 | 0.49 |
| 1:I:344:LEU:HD11 | 1:I:348:LEU:HD12 | 1.93 | 0.49 |
| 1:J:12:TYR:O | 1:L:900:ARG:NH1 | 2.37 | 0.49 |
| 1:K:798:ASN:HD21 | 1:L:124:LEU:H | 1.58 | 0.49 |
| 1:L:388:ASN:HD22 | 1:L:437:LEU:HD23 | 1.77 | 0.49 |
| 2:M:73:LYS:HE3 | 2:M:95:GLN:HA | 1.94 | 0.49 |
| 1:B:146:GLN:HE22 | 1:C:430:ILE:HD11 | 1.77 | 0.49 |
| 1:D:741:ILE:O | 1:D:745:SER:OG | 2.29 | 0.49 |
| 1:I:399:THR:HG21 | 1:I:426:GLU:HA | 1.94 | 0.49 |
| 1:K:405:ILE:HD11 | 1:K:413:THR:HA | 1.93 | 0.49 |
| 1:L:878:PHE:HD2 | 1:L:889:LEU:HD11 | 1.76 | 0.49 |
| 4:P:8:PRO:HB3 | 4:P:202:VAL:HG22 | 1.93 | 0.49 |
| 1:A:655:ARG:HH21 | 1:A:887:THR:HG21 | 1.77 | 0.49 |
| 1:C:323:VAL:HG22 | 1:C:555:GLU:HB2 | 1.92 | 0.49 |
| 1:I:132:PRO:HB2 | 1:I:202:VAL:HG22 | 1.95 | 0.49 |
| 1:L:134:GLU:HG2 | 1:L:205:ASP:HB3 | 1.95 | 0.49 |
| 1:E:201:ARG:HG2 | 1:E:261:TYR:HB2 | 1.94 | 0.49 |
| 1:G:517:ARG:HB3 | 1:I:380:HIS:CE1 | 2.48 | 0.49 |
| 1:G:557:ASN:N | 1:G:557:ASN:OD1 | 2.44 | 0.49 |
| 1:H:147:ALA:HB2 | 1:H:260:LEU:HD21 | 1.93 | 0.49 |
| 1:K:272:THR:HB | 1:K:295:ALA:HB1 | 1.95 | 0.49 |
| 1:G:33:ARG:NH1 | 6:U:20:GLY:O | 82.81 | 0.49 |
| 1:A:188:TRP:HE1 | 1:C:292:THR:HA | 1.77 | 0.49 |
| 1:B:652:SER:HB2 | 1:B:892:LEU:HB3 | 1.93 | 0.49 |
| 1:I:302:TYR:HE2 | 1:I:517:ARG:HG2 | 1.77 | 0.49 |
| 1:L:90:ASP:OD1 | 1:L:906:ARG:NH1 | 2.46 | 0.49 |
| 2:M:380:ASP:HB2 | 2:M:476:GLN:HE22 | 1.78 | 0.49 |
| 1:A:397:ALA:N | 1:A:426:GLU:OE2 | 2.44 | 0.49 |
| 1:B:23:TYR:OH | 1:B:45:PHE:O | 2.25 | 0.49 |

Continued on next page...

CONFIDENTIAL VALIDATION REPORT

*Continued from previous page...*

| Atom-1 | Atom-2 | Interatomic distance (Å) | Clash overlap (Å) |
|---|---|---|---|
| 1:B:474:TYR:O | 1:B:478:ASN:N | 2.46 | 0.49 |
| 1:C:310:ILE:HD13 | 1:C:347:GLN:HE21 | 1.78 | 0.49 |
| 1:C:901:ILE:HG22 | 1:C:911:ALA:HA | 1.94 | 0.49 |
| 1:E:744:LEU:HD12 | 1:E:750:GLY:HA3 | 1.94 | 0.49 |
| 1:I:121:TYR:HE2 | 1:I:270:PRO:HG2 | 1.78 | 0.49 |
| 1:K:493:ILE:HD13 | 1:L:120:ALA:HB2 | 1.94 | 0.49 |
| 1:L:104:ARG:NH1 | 1:L:585:SER:OG | 2.41 | 0.49 |
| 1:A:445:PHE:HE1 | 1:A:512:ARG:HH12 | 1.60 | 0.49 |
| 1:A:508:PHE:HD2 | 1:A:684:THR:HG23 | 1.78 | 0.49 |
| 1:D:657:LYS:HB2 | 1:D:660:GLU:HG2 | 1.95 | 0.49 |
| 1:K:900:ARG:NH1 | 1:L:12:TYR:OH | 2.45 | 0.49 |
| 1:B:323:VAL:HG12 | 1:B:332:ASN:HD22 | 1.78 | 0.49 |
| 1:E:548:LEU:H | 1:E:903:GLN:HE21 | 1.61 | 0.49 |
| 1:D:522:LEU:HD21 | 1:F:731:GLN:HE21 | 1.78 | 0.49 |
| 1:G:216:TYR:HD1 | 1:I:817:ALA:HB3 | 1.78 | 0.49 |
| 1:I:406:LYS:HB3 | 1:I:415:ASP:HB3 | 1.94 | 0.49 |
| 1:K:383:GLU:OE2 | 1:L:512:ARG:NH2 | 2.46 | 0.49 |
| 1:L:343:GLU:HG3 | 1:L:681:LEU:HD22 | 1.95 | 0.49 |
| 1:K:913:TYR:HB2 | 4:O:32:ASN:HB3 | 1.94 | 0.49 |
| 1:A:436:ASN:O | 1:A:440:ASN:ND2 | 2.46 | 0.49 |
| 1:D:434:GLU:HG3 | 1:E:126:PRO:HB3 | 1.95 | 0.49 |
| 1:F:510:HIS:O | 1:F:569:SER:OG | 2.31 | 0.49 |
| 1:J:173:ALA:HA | 1:J:178:GLN:HE21 | 1.78 | 0.49 |
| 1:K:473:THR:HG22 | 1:K:475:ALA:H | 1.76 | 0.49 |
| 1:K:629:ILE:HG13 | 1:K:655:ARG:HH12 | 1.77 | 0.49 |
| 3:N:207:VAL:HG21 | 3:N:243:LEU:HD13 | 1.94 | 0.49 |
| 1:A:602:THR:O | 1:A:606:MET:N | 2.46 | 0.48 |
| 1:B:456:ASP:OD1 | 1:B:476:TYR:OH | 2.31 | 0.48 |
| 1:B:780:ASN:HD22 | 1:B:830:PRO:HB2 | 1.78 | 0.48 |
| 1:I:155:ASN:HD22 | 1:I:158:ASN:HB2 | 1.78 | 0.48 |
| 1:K:155:ASN:HD22 | 1:K:172:TYR:HE2 | 1.61 | 0.48 |
| 1:K:717:GLU:OE2 | 1:K:720:ARG:NH2 | 2.45 | 0.48 |
| 1:K:726:GLY:HA3 | 1:L:104:ARG:HH21 | 1.77 | 0.48 |
| 1:J:533:ILE:HG22 | 1:L:731:GLN:HE21 | 1.78 | 0.48 |
| 3:N:189:TYR:OH | 4:O:208:SER:O | 2.22 | 0.48 |
| 1:B:688:ASN:ND2 | 1:B:842:THR:O | 2.46 | 0.48 |
| 1:C:134:GLU:HG2 | 1:C:143:VAL:HG12 | 1.95 | 0.48 |
| 1:C:52:PRO:HG2 | 1:C:56:VAL:HG21 | 1.95 | 0.48 |
| 1:A:181:PRO:HB3 | 1:C:813:GLN:HE21 | 1.78 | 0.48 |
| 1:D:434:GLU:HG2 | 1:F:393:LEU:HD23 | 1.95 | 0.48 |
| 1:D:493:ILE:O | 1:E:525:ASN:ND2 | 2.36 | 0.48 |

*Continued on next page...*

CONFIDENTIAL VALIDATION REPORT

*Continued from previous page...*

| Atom-1 | Atom-2 | Interatomic distance (Å) | Clash overlap (Å) |
|---|---|---|---|
| 1:G:612:ASN:N | 1:G:612:ASN:OD1 | 2.44 | 0.48 |
| 1:G:613:ASP:OD2 | 1:G:900:ARG:NH2 | 2.32 | 0.48 |
| 1:H:73:ARG:NE | 1:H:80:TYR:OH | 2.41 | 0.48 |
| 1:I:408:ASN:OD1 | 1:I:411:THR:OG1 | 2.30 | 0.48 |
| 1:B:107:LEU:HG | 1:B:109:ARG:HG3 | 1.95 | 0.48 |
| 1:D:634:THR:OG1 | 1:D:879:GLU:OE2 | 2.32 | 0.48 |
| 1:E:217:ALA:HB2 | 1:E:267:ILE:HD13 | 1.94 | 0.48 |
| 1:F:691:PHE:HB2 | 1:F:718:ILE:HD13 | 1.95 | 0.48 |
| 1:G:399:THR:HG21 | 1:G:424:GLU:HG3 | 1.95 | 0.48 |
| 1:I:342:THR:HA | 1:I:345:SER:HB3 | 1.96 | 0.48 |
| 1:I:375:ARG:HH22 | 1:I:503:ASP:HA | 1.78 | 0.48 |
| 1:I:340:ARG:HH21 | 1:I:915:ARG:HD3 | 1.78 | 0.48 |
| 1:J:898:VAL:HG11 | 1:K:13:MET:HB3 | 1.94 | 0.48 |
| 1:K:369:SER:HA | 1:K:841:ARG:HH22 | 1.78 | 0.48 |
| 2:M:235:ASP:OD2 | 2:M:342:SER:OG | 2.32 | 0.48 |
| 3:N:190:GLN:NE2 | 3:N:192:GLY:O | 2.45 | 0.48 |
| 1:A:202:VAL:HG21 | 1:A:260:LEU:HD12 | 1.95 | 0.48 |
| 1:B:736:LYS:NZ | 1:C:100:TYR:OH | 2.36 | 0.48 |
| 1:D:224:GLY:O | 1:D:226:GLN:NE2 | 2.46 | 0.48 |
| 1:G:351:ASP:O | 1:G:763:ARG:NH1 | 2.47 | 0.48 |
| 1:H:499:PRO:HG2 | 1:H:502:MET:HB3 | 1.96 | 0.48 |
| 1:I:121:TYR:HB2 | 1:I:214:GLY:HA2 | 1.96 | 0.48 |
| 1:I:72:ASP:OD2 | 1:I:83:ARG:NH1 | 2.44 | 0.48 |
| 1:J:220:THR:HG21 | 1:J:228:SER:HA | 1.95 | 0.48 |
| 1:K:132:PRO:HB3 | 1:K:145:GLY:HA2 | 1.94 | 0.48 |
| 1:L:453:TYR:OH | 1:L:511:HIS:ND1 | 2.38 | 0.48 |
| 1:L:652:SER:OG | 1:L:894:GLU:OE2 | 2.27 | 0.48 |
| 1:B:562:VAL:HA | 1:B:565:ILE:HG22 | 1.96 | 0.48 |
| 1:B:564:MET:HB3 | 1:B:672:PHE:HE2 | 1.78 | 0.48 |
| 1:B:776:ARG:HD3 | 1:B:834:GLN:HE21 | 1.77 | 0.48 |
| 1:D:375:ARG:HH12 | 1:D:840:ASP:HB3 | 1.77 | 0.48 |
| 1:J:824:ILE:HG12 | 1:K:115:PRO:HD2 | 1.95 | 0.48 |
| 1:K:616:PHE:H | 1:K:899:VAL:HG12 | 1.79 | 0.48 |
| 1:A:709:ARG:NH1 | 4:O:233:ASP:OD2 | 2.46 | 0.48 |
| 1:B:688:ASN:O | 1:B:844:TRP:NE1 | 2.41 | 0.48 |
| 1:F:637:PRO:HA | 1:F:877:THR:HA | 1.94 | 0.48 |
| 1:H:248:ALA:HB2 | 1:H:256:PRO:HA | 1.94 | 0.48 |
| 1:J:624:ASN:ND2 | 1:J:890:TYR:OH | 2.47 | 0.48 |
| 1:L:379:ASN:HD21 | 1:L:493:ILE:HA | 1.79 | 0.48 |
| 1:A:337:LEU:HD12 | 1:A:340:ARG:HH22 | 1.78 | 0.48 |
| 1:C:346:TYR:HB3 | 1:C:538:LYS:HD3 | 1.95 | 0.48 |

*Continued on next page...*

CONFIDENTIAL VALIDATION REPORT

*Continued from previous page...*

| Atom-1 | Atom-2 | Interatomic distance (Å) | Clash overlap (Å) |
|---|---|---|---|
| 1:C:375:ARG:HD3 | 1:C:507:PRO:HB3 | 1.96 | 0.48 |
| 1:C:779:VAL:HG21 | 1:C:821:TYR:HD2 | 1.79 | 0.48 |
| 1:G:146:GLN:HA | 1:H:426:GLU:H | 1.79 | 0.48 |
| 1:H:94:LEU:HD12 | 1:H:547:LEU:HD13 | 1.96 | 0.48 |
| 1:K:94:LEU:HD22 | 1:K:547:LEU:HD13 | 1.96 | 0.48 |
| 1:B:697:MET:HA | 1:B:703:SER:HA | 1.96 | 0.48 |
| 1:C:351:ASP:HA | 1:C:356:ARG:HB2 | 1.95 | 0.48 |
| 1:C:88:VAL:O | 1:C:550:GLY:N | 2.47 | 0.48 |
| 1:C:717:GLU:O | 1:C:735:THR:OG1 | 2.28 | 0.48 |
| 1:G:339:ASP:OD1 | 1:G:915:ARG:NH1 | 2.42 | 0.48 |
| 1:H:350:LEU:HD21 | 1:H:365:GLN:HG3 | 1.94 | 0.48 |
| 1:J:684:THR:HG23 | 1:J:841:ARG:HG3 | 1.96 | 0.48 |
| 1:A:60:ARG:HH22 | 4:O:111:GLY:HA3 | 1.79 | 0.48 |
| 1:A:443:ARG:HH12 | 1:A:802:VAL:HG21 | 1.78 | 0.48 |
| 1:B:176:THR:HG22 | 1:B:225:GLY:HA3 | 1.95 | 0.48 |
| 1:E:210:LEU:HD13 | 1:E:269:ALA:HB2 | 1.96 | 0.48 |
| 1:D:434:GLU:N | 1:F:391:PHE:O | 2.46 | 0.48 |
| 1:G:298:ASN:HA | 1:G:478:ASN:HD21 | 1.78 | 0.48 |
| 1:G:73:ARG:NH2 | 1:G:80:TYR:OH | 2.47 | 0.48 |
| 1:J:327:GLN:HA | 1:J:330:GLN:HE21 | 1.79 | 0.48 |
| 1:K:678:VAL:HG23 | 1:K:681:LEU:HB2 | 1.95 | 0.48 |
| 1:L:294:GLN:NE2 | 1:L:807:PRO:O | 2.47 | 0.48 |
| 1:B:613:ASP:OD2 | 1:B:900:ARG:NH2 | 2.46 | 0.48 |
| 1:C:632:ASN:HD22 | 1:C:882:PRO:HB2 | 1.79 | 0.48 |
| 1:G:406:LYS:N | 1:G:413:THR:O | 2.45 | 0.48 |
| 1:G:323:VAL:HG22 | 1:G:555:GLU:HG2 | 1.94 | 0.48 |
| 1:J:109:ARG:HH22 | 1:J:523:LEU:HB3 | 1.78 | 0.48 |
| 1:K:506:ASN:HB2 | 1:K:508:PHE:H | 1.79 | 0.48 |
| 2:M:77:ILE:HD13 | 2:M:92:THR:HG23 | 1.96 | 0.48 |
| 1:A:361:SER:OG | 1:A:518:TYR:OH | 2.26 | 0.47 |
| 1:A:717:GLU:O | 1:A:735:THR:OG1 | 2.26 | 0.47 |
| 1:C:626:LEU:HD12 | 1:C:888:LEU:HD13 | 1.96 | 0.47 |
| 1:F:356:ARG:HB3 | 1:F:367:VAL:HG11 | 1.95 | 0.47 |
| 1:G:83:ARG:NH1 | 1:G:555:GLU:OE1 | 2.46 | 0.47 |
| 1:A:145:GLY:H | 1:B:425:ILE:HG12 | 1.79 | 0.47 |
| 1:E:315:TYR:HE2 | 1:E:342:THR:HB | 1.79 | 0.47 |
| 1:D:396:SER:OG | 1:F:144:ARG:NH2 | 2.46 | 0.47 |
| 1:F:149:PHE:HB3 | 1:F:199:ALA:HB3 | 1.97 | 0.47 |
| 1:F:717:GLU:OE1 | 1:F:720:ARG:NH2 | 2.33 | 0.47 |
| 1:I:122:ASN:HB3 | 1:I:125:ALA:HB2 | 1.96 | 0.47 |
| 1:I:447:TYR:HA | 1:I:451:ALA:HB3 | 1.96 | 0.47 |

*Continued on next page...*

CONFIDENTIAL VALIDATION REPORT

*Continued from previous page...*

| Atom-1 | Atom-2 | Interatomic distance (Å) | Clash overlap (Å) |
|---|---|---|---|
| 1:L:467:LEU:HD12 | 1:L:476:TYR:HA | 1.96 | 0.47 |
| 1:L:506:ASN:HB2 | 1:L:508:PHE:HD2 | 1.79 | 0.47 |
| 1:B:539:PHE:O | 1:B:543:LYS:N | 2.48 | 0.47 |
| 1:B:698:PHE:N | 1:B:702:VAL:O | 2.46 | 0.47 |
| 1:D:144:ARG:NH2 | 1:E:396:SER:OG | 2.47 | 0.47 |
| 1:D:360:PHE:HA | 1:F:768:PHE:HZ | 1.79 | 0.47 |
| 1:D:99:THR:HB | 1:D:588:LEU:HD12 | 1.96 | 0.47 |
| 1:F:640:ILE:HD12 | 1:F:874:LEU:HD21 | 1.95 | 0.47 |
| 1:G:727:TYR:HA | 1:G:736:LYS:HE2 | 1.96 | 0.47 |
| 1:K:652:SER:HB2 | 1:K:892:LEU:HB3 | 1.95 | 0.47 |
| 1:L:18:GLN:HB3 | 1:L:22:GLU:HB2 | 1.95 | 0.47 |
| 1:A:622:ALA:HB1 | 1:A:892:LEU:HB2 | 1.96 | 0.47 |
| 1:F:187:GLN:HB2 | 1:F:190:SER:HB3 | 1.97 | 0.47 |
| 1:F:772:GLN:HE21 | 1:F:838:LEU:HD22 | 1.79 | 0.47 |
| 1:G:88:VAL:HG13 | 1:G:550:GLY:H | 1.80 | 0.47 |
| 1:H:496:ARG:H | 1:I:521:MET:HB3 | 1.80 | 0.47 |
| 1:H:875:ASP:N | 1:H:875:ASP:OD1 | 2.46 | 0.47 |
| 1:K:657:LYS:HB2 | 1:K:660:GLU:HG2 | 1.95 | 0.47 |
| 1:K:813:GLN:HB2 | 1:L:146:GLN:HG3 | 1.95 | 0.47 |
| 1:L:10:TRP:HA | 1:L:15:ILE:HD12 | 1.96 | 0.47 |
| 1:L:347:GLN:NE2 | 1:L:682:ASP:O | 2.47 | 0.47 |
| 1:L:404:GLY:HA2 | 1:L:418:TYR:HD2 | 1.78 | 0.47 |
| 1:D:692:LYS:HB3 | 1:D:879:GLU:HG3 | 1.97 | 0.47 |
| 1:E:613:ASP:HB2 | 1:E:900:ARG:HE | 1.80 | 0.47 |
| 1:F:770:ASN:ND2 | 1:F:841:ARG:O | 2.48 | 0.47 |
| 1:J:813:GLN:NE2 | 1:K:148:PRO:HD2 | 2.29 | 0.47 |
| 1:L:58:THR:HG22 | 1:L:596:ALA:HA | 1.95 | 0.47 |
| 1:A:53:THR:OG1 | 1:A:54:HIS:N | 2.47 | 0.47 |
| 1:B:443:ARG:O | 1:B:447:TYR:N | 2.48 | 0.47 |
| 1:B:143:VAL:HG22 | 1:C:423:ALA:HB1 | 1.95 | 0.47 |
| 1:D:303:ILE:HG23 | 1:D:566:LEU:HD13 | 1.96 | 0.47 |
| 1:D:376:ILE:HG22 | 1:D:378:GLU:HG3 | 1.97 | 0.47 |
| 1:D:72:ASP:OD1 | 1:D:83:ARG:N | 2.47 | 0.47 |
| 1:E:905:HIS:HB2 | 1:E:908:VAL:HB | 1.95 | 0.47 |
| 1:G:403:SER:H | 1:G:423:ALA:HB3 | 1.78 | 0.47 |
| 1:G:582:ARG:NH1 | 1:G:584:ASP:OD1 | 2.47 | 0.47 |
| 1:I:575:ARG:NH1 | 5:Q:38:SER:OG | 2.48 | 0.47 |
| 1:K:246:PHE:HB2 | 1:L:405:ILE:HG12 | 1.96 | 0.47 |
| 1:K:610:ASP:OD2 | 4:O:43:ARG:NH2 | 2.48 | 0.47 |
| 1:K:645:TRP:NE1 | 1:K:872:HIS:O | 2.46 | 0.47 |
| 1:A:526:GLY:H | 1:C:777:GLN:HG3 | 1.78 | 0.47 |

*Continued on next page...*

CONFIDENTIAL VALIDATION REPORT

Continued from previous page...

| Atom-1 | Atom-2 | Interatomic distance (Å) | Clash overlap (Å) |
|---|---|---|---|
| 1:I:727:TYR:HA | 1:I:736:LYS:NZ | 2.30 | 0.47 |
| 1:L:257:LYS:HG3 | 1:L:258:ALA:H | 1.79 | 0.47 |
| 1:B:10:TRP:CE2 | 4:O:16:GLN:HB3 | 2.50 | 0.47 |
| 1:B:318:THR:OG1 | 1:B:336:ASP:O | 2.32 | 0.47 |
| 1:B:72:ASP:HB3 | 1:B:83:ARG:HG2 | 1.97 | 0.47 |
| 1:E:652:SER:HB2 | 1:E:892:LEU:HB3 | 1.96 | 0.47 |
| 1:G:402:TYR:N | 1:G:423:ALA:O | 2.48 | 0.47 |
| 1:I:624:ASN:ND2 | 1:I:680:TYR:OH | 2.47 | 0.47 |
| 1:J:72:ASP:HB2 | 1:J:83:ARG:HH11 | 1.78 | 0.47 |
| 1:C:136:LYS:HA | 1:C:141:ILE:HA | 1.95 | 0.47 |
| 1:D:132:PRO:HB3 | 1:D:145:GLY:HA2 | 1.96 | 0.47 |
| 1:G:301:ASN:HA | 1:G:569:SER:HB3 | 1.97 | 0.47 |
| 1:G:509:ASN:OD1 | 1:G:674:TYR:OH | 2.32 | 0.47 |
| 1:J:146:GLN:HE21 | 1:L:813:GLN:HB2 | 1.79 | 0.47 |
| 4:P:95:ALA:HA | 4:P:98:GLU:HB3 | 1.97 | 0.47 |
| 1:A:391:PHE:O | 1:B:434:GLU:N | 2.47 | 0.47 |
| 1:D:254:ASN:ND2 | 1:E:412:TRP:O | 2.48 | 0.47 |
| 1:G:221:ASN:HB2 | 1:G:225:GLY:HA3 | 1.96 | 0.47 |
| 1:G:547:LEU:HB2 | 1:G:552:TYR:HE2 | 1.80 | 0.47 |
| 1:J:860:ASP:OD2 | 1:K:57:THR:OG1 | 2.32 | 0.47 |
| 1:A:341:ASN:H | 1:A:624:ASN:ND2 | 2.11 | 0.47 |
| 1:A:398:ALA:N | 1:A:426:GLU:OE2 | 2.42 | 0.47 |
| 1:A:525:ASN:HB2 | 1:C:495:ALA:HB2 | 1.96 | 0.47 |
| 1:B:601:SER:OG | 3:N:100:GLY:N | 2.48 | 0.47 |
| 1:E:596:ALA:HB3 | 1:E:599:THR:HG22 | 1.97 | 0.47 |
| 1:F:207:THR:OG1 | 1:F:265:VAL:O | 2.33 | 0.47 |
| 1:G:187:GLN:O | 1:G:190:SER:OG | 2.30 | 0.47 |
| 1:G:217:ALA:HB2 | 1:G:267:ILE:HD13 | 1.97 | 0.47 |
| 1:H:114:LYS:NZ | 1:H:116:TYR:O | 2.43 | 0.47 |
| 1:H:272:THR:HB | 1:H:295:ALA:HB1 | 1.96 | 0.47 |
| 1:H:483:VAL:HG22 | 1:H:485:SER:H | 1.79 | 0.47 |
| 1:K:869:ASN:HB2 | 4:O:200:PRO:HD3 | 1.97 | 0.47 |
| 1:L:542:ILE:HG22 | 1:L:545:LEU:HD13 | 1.96 | 0.47 |
| 1:A:655:ARG:HE | 1:A:887:THR:HG21 | 1.80 | 0.46 |
| 1:E:375:ARG:HD2 | 1:E:507:PRO:HG3 | 1.97 | 0.46 |
| 1:H:94:LEU:HD23 | 1:H:592:PHE:HD1 | 1.80 | 0.46 |
| 1:I:490:TYR:HD2 | 1:I:820:PRO:HD3 | 1.80 | 0.46 |
| 1:I:337:LEU:HD12 | 1:I:915:ARG:HE | 1.80 | 0.46 |
| 1:J:312:LEU:HD13 | 1:J:565:ILE:HG12 | 1.96 | 0.46 |
| 3:N:57:PRO:HD2 | 6:Y:12:ARG:CZ | 2.45 | 0.46 |
| 1:B:315:TYR:HE2 | 1:B:538:LYS:HB2 | 1.80 | 0.46 |

Continued on next page...

CONFIDENTIAL VALIDATION REPORT

Continued from previous page...

| Atom-1 | Atom-2 | Interatomic distance (Å) | Clash overlap (Å) |
|---|---|---|---|
| 1:H:858:LEU:HD11 | 1:I:16:ALA:HB2 | 1.96 | 0.46 |
| 1:I:163:GLY:HA3 | 1:I:171:ILE:HD11 | 1.96 | 0.46 |
| 1:I:506:ASN:HB3 | 1:I:686:TYR:CZ | 2.50 | 0.46 |
| 1:K:390:CYS:N | 1:K:433:MET:O | 2.38 | 0.46 |
| 1:C:116:TYR:OH | 1:C:271:ASP:OD2 | 2.34 | 0.46 |
| 1:C:210:LEU:HD12 | 1:C:267:ILE:HG22 | 1.97 | 0.46 |
| 1:H:65:THR:O | 1:H:65:THR:OG1 | 2.32 | 0.46 |
| 1:I:151:GLY:N | 1:I:197:LYS:O | 2.47 | 0.46 |
| 1:I:317:SER:OG | 1:I:665:GLY:O | 2.33 | 0.46 |
| 1:J:736:LYS:NZ | 1:K:591:ASN:OD1 | 2.49 | 0.46 |
| 1:K:798:ASN:ND2 | 1:L:124:LEU:H | 2.13 | 0.46 |
| 1:L:504:ASN:ND2 | 1:L:883:MET:SD | 2.88 | 0.46 |
| 3:N:89:THR:HG22 | 3:N:94:ILE:HB | 1.97 | 0.46 |
| 1:B:699:ASP:N | 1:B:873:ALA:O | 2.49 | 0.46 |
| 1:C:217:ALA:HB2 | 1:C:267:ILE:HD13 | 1.96 | 0.46 |
| 1:G:496:ARG:HH22 | 1:G:772:GLN:HB2 | 1.81 | 0.46 |
| 1:H:103:ILE:HD12 | 1:H:312:LEU:HD21 | 1.97 | 0.46 |
| 1:H:306:ARG:HG2 | 1:H:567:GLN:HB2 | 1.98 | 0.46 |
| 1:J:693:LYS:HB3 | 1:J:879:GLU:HB3 | 1.96 | 0.46 |
| 1:K:19:ASP:HA | 1:K:48:PRO:HD2 | 1.97 | 0.46 |
| 2:M:386:VAL:HG22 | 2:M:387:THR:HG23 | 1.96 | 0.46 |
| 1:B:74:GLU:OE1 | 2:M:393:GLN:NE2 | 2.49 | 0.46 |
| 1:A:19:ASP:HA | 1:A:48:PRO:HD2 | 1.97 | 0.46 |
| 1:A:79:SER:OG | 1:A:80:TYR:N | 2.48 | 0.46 |
| 1:A:864:ASN:ND2 | 4:O:230:ASP:O | 2.48 | 0.46 |
| 1:A:627:TYR:O | 1:A:889:LEU:N | 2.49 | 0.46 |
| 1:E:101:PHE:HB2 | 1:E:535:VAL:HB | 1.98 | 0.46 |
| 1:F:221:ASN:OD1 | 1:F:225:GLY:N | 2.47 | 0.46 |
| 1:I:241:ASP:OD2 | 1:I:264:ASN:ND2 | 2.47 | 0.46 |
| 1:I:370:TYR:HE1 | 1:I:841:ARG:HE | 1.64 | 0.46 |
| 1:L:167:THR:HG23 | 1:L:169:GLN:HE22 | 1.81 | 0.46 |
| 1:L:41:LEU:HB3 | 1:L:45:PHE:HE2 | 1.81 | 0.46 |
| 1:B:906:ARG:HB2 | 3:N:104:ASN:HD21 | 1.80 | 0.46 |
| 1:C:626:LEU:HD21 | 1:C:680:TYR:HE1 | 1.80 | 0.46 |
| 1:D:101:PHE:HB2 | 1:D:535:VAL:HB | 1.98 | 0.46 |
| 1:D:435:ILE:HD13 | 1:D:437:LEU:HB3 | 1.96 | 0.46 |
| 1:E:405:ILE:HG22 | 1:E:414:ALA:HA | 1.98 | 0.46 |
| 1:E:103:ILE:HG12 | 1:E:586:ILE:HG12 | 1.97 | 0.46 |
| 1:E:690:THR:O | 1:E:881:ASP:N | 2.47 | 0.46 |
| 1:E:707:ASN:HB3 | 1:F:61:SER:HA | 1.97 | 0.46 |
| 1:H:108:ASP:HB3 | 1:H:580:SER:H | 1.81 | 0.46 |

Continued on next page...

CONFIDENTIAL VALIDATION REPORT

Continued from previous page...

| Atom-1 | Atom-2 | Interatomic distance (Å) | Clash overlap (Å) |
|---|---|---|---|
| 1:I:640:ILE:HD11 | 1:I:645:TRP:HH2 | 1.81 | 0.46 |
| 1:J:699:ASP:N | 1:J:873:ALA:O | 2.46 | 0.46 |
| 1:A:18:GLN:HE21 | 1:A:22:GLU:HB3 | 1.80 | 0.46 |
| 1:B:137:ASP:HB2 | 1:B:140:LYS:HB2 | 1.97 | 0.46 |
| 1:K:375:ARG:NH1 | 1:K:840:ASP:OD2 | 2.49 | 0.46 |
| 1:E:919:SER:OG | 4:O:105:GLY:O | 2.25 | 0.46 |
| 1:B:323:VAL:HG22 | 1:B:555:GLU:HB2 | 1.97 | 0.46 |
| 1:G:623:ALA:HB3 | 1:G:893:PHE:HB2 | 1.97 | 0.46 |
| 1:G:693:LYS:HA | 1:G:717:GLU:HA | 1.98 | 0.46 |
| 1:H:302:TYR:HB3 | 1:H:520:SER:HB3 | 1.98 | 0.46 |
| 1:J:19:ASP:OD1 | 1:J:20:ALA:N | 2.48 | 0.46 |
| 1:J:720:ARG:HH12 | 1:J:727:TYR:HD2 | 1.64 | 0.46 |
| 1:A:313:MET:HG2 | 1:A:556:TRP:HE1 | 1.80 | 0.46 |
| 1:C:766:SER:OG | 1:C:767:PHE:N | 2.47 | 0.46 |
| 1:D:325:ALA:HB2 | 1:D:332:ASN:HA | 1.96 | 0.46 |
| 1:D:546:LEU:HB3 | 1:D:614:GLN:HE22 | 1.81 | 0.46 |
| 1:F:302:TYR:HE2 | 1:F:517:ARG:HD2 | 1.80 | 0.46 |
| 1:I:375:ARG:HD2 | 1:I:507:PRO:HB3 | 1.97 | 0.46 |
| 1:K:795:GLN:HG3 | 1:K:819:TYR:HB2 | 1.97 | 0.46 |
| 2:M:430:THR:HB | 2:M:432:VAL:HG23 | 1.98 | 0.46 |
| 1:D:103:ILE:HG22 | 1:D:531:PHE:HE2 | 1.81 | 0.46 |
| 1:F:132:PRO:HB3 | 1:F:145:GLY:HA3 | 1.98 | 0.46 |
| 1:G:113:PHE:CD1 | 1:G:303:ILE:HG22 | 2.51 | 0.46 |
| 1:J:462:PRO:HD2 | 1:J:465:ILE:HD11 | 1.98 | 0.46 |
| 1:J:623:ALA:HB3 | 1:J:893:PHE:HB2 | 1.98 | 0.46 |
| 1:K:731:GLN:HG3 | 1:L:522:LEU:HD21 | 1.98 | 0.46 |
| 2:M:126:ARG:HB3 | 2:M:458:PRO:HB3 | 1.97 | 0.46 |
| 3:N:57:PRO:HD2 | 6:Y:12:ARG:NH2 | 2.30 | 0.46 |
| 1:A:120:ALA:HB2 | 1:C:493:ILE:HD13 | 1.98 | 0.45 |
| 1:B:709:ARG:HH22 | 1:B:850:SER:HB3 | 1.81 | 0.45 |
| 1:C:80:TYR:HD2 | 1:C:558:PHE:HB2 | 1.80 | 0.45 |
| 1:E:131:ASN:ND2 | 1:E:210:LEU:O | 2.49 | 0.45 |
| 1:D:389:TYR:HE2 | 1:E:810:ARG:HB3 | 1.81 | 0.45 |
| 1:G:256:PRO:HG3 | 1:H:412:TRP:CG | 2.51 | 0.45 |
| 1:G:606:MET:O | 1:G:612:ASN:ND2 | 2.46 | 0.45 |
| 1:G:663:SER:N | 1:G:669:ASP:OD2 | 2.43 | 0.45 |
| 1:G:868:ALA:HB3 | 1:H:3:THR:HG21 | 1.97 | 0.45 |
| 1:H:389:TYR:HB3 | 1:H:434:GLU:HB3 | 1.97 | 0.45 |
| 1:I:339:ASP:HB2 | 1:I:623:ALA:HB1 | 1.97 | 0.45 |
| 1:J:548:LEU:HB2 | 1:J:903:GLN:HE21 | 1.81 | 0.45 |
| 1:L:367:VAL:HG23 | 1:L:369:SER:H | 1.81 | 0.45 |

Continued on next page...

CONFIDENTIAL VALIDATION REPORT

*Continued from previous page...*

| Atom-1 | Atom-2 | Interatomic distance (Å) | Clash overlap (Å) |
|---|---|---|---|
| 1:L:381:GLY:HA2 | 1:L:493:ILE:HD11 | 1.98 | 0.45 |
| 1:A:718:ILE:HD13 | 1:A:738:TRP:CD1 | 2.51 | 0.45 |
| 1:D:356:ARG:HG3 | 1:D:367:VAL:HG12 | 1.99 | 0.45 |
| 1:D:361:SER:HB2 | 1:D:522:LEU:HD11 | 1.98 | 0.45 |
| 1:E:314:TYR:HD1 | 1:E:559:ARG:HB2 | 1.81 | 0.45 |
| 1:E:649:ARG:NH2 | 1:E:895:VAL:O | 2.44 | 0.45 |
| 1:F:377:ILE:HD11 | 1:F:450:VAL:HG11 | 1.98 | 0.45 |
| 1:F:62:GLN:HG3 | 1:F:594:PRO:HA | 1.97 | 0.45 |
| 1:F:638:ILE:HD12 | 1:F:638:ILE:HA | 1.83 | 0.45 |
| 1:G:123:SER:O | 1:I:436:ASN:ND2 | 2.49 | 0.45 |
| 1:G:150:ILE:HG12 | 1:G:198:VAL:HG23 | 1.99 | 0.45 |
| 1:G:603:LEU:HD12 | 1:H:31:PHE:HE2 | 1.81 | 0.45 |
| 1:L:688:ASN:ND2 | 1:L:842:THR:O | 2.49 | 0.45 |
| 1:A:405:ILE:HD13 | 1:A:412:TRP:CD1 | 2.52 | 0.45 |
| 1:B:292:THR:HG23 | 1:C:188:TRP:HD1 | 1.80 | 0.45 |
| 1:C:336:ASP:OD1 | 1:C:340:ARG:NH2 | 2.49 | 0.45 |
| 1:D:312:LEU:HD13 | 1:D:565:ILE:HD11 | 1.96 | 0.45 |
| 1:I:548:LEU:HB2 | 1:I:903:GLN:HE21 | 1.81 | 0.45 |
| 1:L:229:LEU:HA | 1:L:240:SER:HA | 1.97 | 0.45 |
| 1:A:558:PHE:HE2 | 1:A:586:ILE:HD13 | 1.81 | 0.45 |
| 1:B:399:THR:HG22 | 1:B:426:GLU:HG3 | 1.99 | 0.45 |
| 1:C:155:ASN:ND2 | 1:C:172:TYR:OH | 2.49 | 0.45 |
| 1:C:878:PHE:HD2 | 1:C:889:LEU:HD11 | 1.81 | 0.45 |
| 1:D:749:ILE:HD12 | 1:D:749:ILE:HA | 1.85 | 0.45 |
| 1:J:263:GLU:HB3 | 1:J:265:VAL:HG13 | 1.97 | 0.45 |
| 1:K:150:ILE:HD13 | 1:K:198:VAL:HG12 | 1.98 | 0.45 |
| 1:K:709:ARG:NH2 | 1:L:58:THR:O | 2.49 | 0.45 |
| 3:N:30:THR:OG1 | 3:N:33:ASP:OD1 | 2.32 | 0.45 |
| 1:K:10:TRP:CE2 | 4:P:16:GLN:HB3 | 2.51 | 0.45 |
| 1:F:315:TYR:HB2 | 1:F:343:GLU:HG2 | 1.97 | 0.45 |
| 1:I:201:ARG:NH2 | 1:I:263:GLU:OE2 | 2.36 | 0.45 |
| 1:K:506:ASN:HB3 | 1:K:686:TYR:CG | 2.52 | 0.45 |
| 1:L:497:TRP:CH2 | 1:L:836:LYS:HG2 | 2.51 | 0.45 |
| 3:N:232:ILE:HA | 3:N:235:LEU:HB3 | 1.99 | 0.45 |
| 1:A:10:TRP:O | 1:A:14:HIS:N | 2.49 | 0.45 |
| 1:D:87:ALA:HA | 1:D:551:SER:HA | 1.98 | 0.45 |
| 1:F:656:LEU:HD13 | 1:F:661:THR:HG21 | 1.99 | 0.45 |
| 1:I:374:VAL:HG11 | 1:I:510:HIS:CE1 | 2.52 | 0.45 |
| 1:I:573:ASP:OD1 | 1:I:573:ASP:N | 2.49 | 0.45 |
| 1:I:626:LEU:HD21 | 1:I:680:TYR:HE1 | 1.82 | 0.45 |
| 1:I:655:ARG:NH1 | 1:I:883:MET:SD | 2.87 | 0.45 |

*Continued on next page...*

CONFIDENTIAL VALIDATION REPORT

Continued from previous page...

| Atom-1 | Atom-2 | Interatomic distance (Å) | Clash overlap (Å) |
|---|---|---|---|
| 1:J:350:LEU:HD22 | 1:J:356:ARG:HH11 | 1.81 | 0.45 |
| 1:J:732:CYS:HA | 1:J:835:LYS:HB2 | 1.97 | 0.45 |
| 1:J:810:ARG:NH1 | 1:L:430:ILE:O | 2.50 | 0.45 |
| 3:N:262:LEU:HD23 | 3:N:265:LEU:HD12 | 1.98 | 0.45 |
| 3:N:60:ASN:HB2 | 6:Y:12:ARG:HH22 | 1.80 | 0.45 |
| 1:A:598:ASN:ND2 | 4:O:93:ARG:O | 2.44 | 0.45 |
| 1:A:375:ARG:HD3 | 1:A:507:PRO:HG2 | 1.99 | 0.45 |
| 1:A:593:PHE:HD2 | 1:A:595:MET:HG2 | 1.80 | 0.45 |
| 1:A:740:LEU:HD12 | 1:B:64:LEU:HD21 | 1.99 | 0.45 |
| 1:A:483:VAL:HG11 | 1:A:792:LEU:HD21 | 1.98 | 0.45 |
| 1:C:323:VAL:HG12 | 1:C:332:ASN:HD22 | 1.82 | 0.45 |
| 1:E:103:ILE:HD12 | 1:E:533:ILE:HD11 | 1.99 | 0.45 |
| 1:F:351:ASP:HB2 | 1:F:763:ARG:HE | 1.81 | 0.45 |
| 1:G:58:THR:HG22 | 1:G:60:ARG:H | 1.80 | 0.45 |
| 1:H:710:LEU:HD12 | 1:H:737:ASP:HB3 | 1.98 | 0.45 |
| 1:I:378:GLU:HB3 | 1:I:380:HIS:CD2 | 2.51 | 0.45 |
| 1:I:870:SER:OG | 1:I:871:ALA:N | 2.49 | 0.45 |
| 1:K:459:LYS:O | 1:K:480:ARG:NH2 | 2.41 | 0.45 |
| 1:K:896:PHE:HE2 | 1:L:15:ILE:HG12 | 1.82 | 0.45 |
| 1:K:612:ASN:HD21 | 1:L:25:SER:H | 1.65 | 0.45 |
| 2:M:81:ASN:HB2 | 2:M:84:ASN:HB2 | 1.98 | 0.45 |
| 4:O:56:LEU:HD11 | 4:O:204:PHE:HE2 | 1.82 | 0.45 |
| 1:D:697:MET:HA | 1:D:703:SER:HA | 1.98 | 0.45 |
| 1:E:210:LEU:HD12 | 1:E:267:ILE:HG22 | 1.99 | 0.45 |
| 1:D:752:GLN:NE2 | 1:F:40:SER:O | 2.40 | 0.45 |
| 1:F:903:GLN:HE21 | 1:F:909:ILE:HD12 | 1.82 | 0.45 |
| 1:G:762:ASP:OD2 | 1:H:358:ARG:NH2 | 2.50 | 0.45 |
| 1:H:858:LEU:HD12 | 1:I:50:VAL:HA | 1.98 | 0.45 |
| 1:G:222:GLU:HA | 1:I:788:GLN:HB3 | 1.99 | 0.45 |
| 1:J:210:LEU:HD21 | 1:J:269:ALA:HB2 | 1.97 | 0.45 |
| 1:J:90:ASP:H | 1:J:549:PRO:HB3 | 1.81 | 0.45 |
| 2:M:424:ARG:HH21 | 2:M:444:ARG:HH21 | 1.65 | 0.45 |
| 4:O:95:ALA:HA | 4:O:98:GLU:HB3 | 1.97 | 0.45 |
| 1:B:388:ASN:ND2 | 1:C:440:ASN:OD1 | 2.50 | 0.45 |
| 1:F:561:ASP:OD1 | 1:F:575:ARG:NH2 | 2.50 | 0.45 |
| 1:G:552:TYR:HE1 | 1:G:907:GLY:HA2 | 1.81 | 0.45 |
| 1:G:624:ASN:ND2 | 1:G:890:TYR:OH | 2.50 | 0.45 |
| 1:G:141:ILE:N | 1:H:418:TYR:O | 2.49 | 0.45 |
| 1:H:458:TYR:O | 1:H:480:ARG:NH2 | 2.50 | 0.45 |
| 1:J:568:SER:OG | 1:J:569:SER:N | 2.50 | 0.45 |
| 1:J:779:VAL:HG21 | 1:J:821:TYR:HD2 | 1.81 | 0.45 |

Continued on next page...

CONFIDENTIAL VALIDATION REPORT

Continued from previous page...

| Atom-1 | Atom-2 | Interatomic distance (Å) | Clash overlap (Å) |
|---|---|---|---|
| 4:P:66:ARG:HE | 4:P:69:LEU:HA | 1.81 | 0.45 |
| 1:A:292:THR:HG21 | 1:B:187:GLN:HG3 | 1.99 | 0.45 |
| 1:C:374:VAL:HG11 | 1:C:510:HIS:CE1 | 2.52 | 0.45 |
| 1:C:357:SER:OG | 1:C:764:MET:SD | 2.74 | 0.45 |
| 1:E:273:HIS:CE1 | 1:E:481:VAL:HG21 | 2.53 | 0.45 |
| 1:D:731:GLN:HE21 | 1:E:533:ILE:HG22 | 1.81 | 0.45 |
| 1:F:356:ARG:HD3 | 1:F:764:MET:HB3 | 1.98 | 0.45 |
| 1:F:89:GLY:O | 1:F:92:ARG:NH1 | 2.50 | 0.45 |
| 1:G:286:SER:OG | 1:G:286:SER:O | 2.34 | 0.45 |
| 1:H:629:ILE:HD11 | 1:H:655:ARG:HH21 | 1.82 | 0.45 |
| 1:J:10:TRP:HE1 | 1:L:916:THR:HG21 | 1.82 | 0.45 |
| 1:J:229:LEU:HA | 1:J:240:SER:HA | 1.99 | 0.45 |
| 2:M:133:ASN:HA | 2:M:168:GLU:HG3 | 1.98 | 0.45 |
| 1:B:319:GLY:HA3 | 2:M:90:GLN:HE22 | 1.82 | 0.45 |
| 4:O:66:ARG:HE | 4:O:69:LEU:HA | 1.81 | 0.45 |
| 1:B:496:ARG:HH21 | 1:B:772:GLN:HE21 | 1.65 | 0.44 |
| 1:C:78:TYR:CG | 1:C:668:PHE:HB2 | 2.52 | 0.44 |
| 1:D:301:ASN:HB2 | 1:D:569:SER:H | 1.82 | 0.44 |
| 1:F:375:ARG:HH12 | 1:F:507:PRO:HG3 | 1.82 | 0.44 |
| 1:F:390:CYS:HB3 | 1:F:433:MET:HB2 | 2.00 | 0.44 |
| 1:G:558:PHE:HE2 | 1:G:586:ILE:HG12 | 1.81 | 0.44 |
| 1:K:483:VAL:HG22 | 1:K:485:SER:H | 1.82 | 0.44 |
| 1:L:557:ASN:OD1 | 1:L:557:ASN:N | 2.50 | 0.44 |
| 1:L:624:ASN:ND2 | 1:L:680:TYR:OH | 2.50 | 0.44 |
| 1:B:301:ASN:HA | 1:B:569:SER:HB2 | 1.99 | 0.44 |
| 1:G:352:ALA:HA | 1:G:763:ARG:HH12 | 1.81 | 0.44 |
| 1:H:83:ARG:HG2 | 1:H:555:GLU:HB3 | 1.98 | 0.44 |
| 1:H:568:SER:N | 1:H:675:SER:O | 2.44 | 0.44 |
| 1:J:271:ASP:OD1 | 1:J:271:ASP:N | 2.50 | 0.44 |
| 1:J:701:SER:OG | 1:J:702:VAL:N | 2.47 | 0.44 |
| 1:L:23:TYR:OH | 1:L:45:PHE:O | 2.35 | 0.44 |
| 1:L:693:LYS:HB2 | 1:L:879:GLU:HB3 | 1.99 | 0.44 |
| 1:E:151:GLY:N | 1:E:197:LYS:O | 2.49 | 0.44 |
| 1:E:515:GLY:O | 1:E:519:ARG:NH1 | 2.50 | 0.44 |
| 1:E:804:TYR:N | 1:E:811:GLU:OE2 | 2.50 | 0.44 |
| 1:E:140:LYS:HZ1 | 1:F:416:ASP:HB3 | 1.82 | 0.44 |
| 1:H:447:TYR:HA | 1:H:451:ALA:HB3 | 1.99 | 0.44 |
| 1:H:717:GLU:HB2 | 1:H:735:THR:HG21 | 2.00 | 0.44 |
| 1:A:845:ARG:HH22 | 7:W:22:GLY:HA3 | 88.84 | 0.44 |
| 1:A:107:LEU:HG | 1:A:109:ARG:HG3 | 1.99 | 0.44 |
| 1:A:130:PRO:HA | 1:A:211:PRO:HA | 1.99 | 0.44 |

Continued on next page...

CONFIDENTIAL VALIDATION REPORT

Continued from previous page...

| Atom-1 | Atom-2 | Interatomic distance (Å) | Clash overlap (Å) |
|---|---|---|---|
| 1:B:451:ALA:HA | 1:B:487:LEU:HD23 | 1.99 | 0.44 |
| 1:B:564:MET:HE1 | 1:B:664:LEU:HD21 | 1.98 | 0.44 |
| 1:B:294:GLN:OE1 | 1:B:809:MET:N | 2.50 | 0.44 |
| 1:C:629:ILE:N | 1:C:887:THR:O | 2.47 | 0.44 |
| 1:E:292:THR:HA | 1:F:188:TRP:HE1 | 1.82 | 0.44 |
| 1:I:109:ARG:HH22 | 1:I:523:LEU:HB3 | 1.82 | 0.44 |
| 2:M:416:GLN:O | 2:M:420:SER:N | 2.44 | 0.44 |
| 4:P:7:THR:HG22 | 4:P:26:ASP:HB3 | 2.00 | 0.44 |
| 1:B:43:ASN:HB3 | 1:B:44:LYS:HG3 | 1.99 | 0.44 |
| 1:D:748:ASN:HD21 | 1:D:854:SER:H | 1.64 | 0.44 |
| 1:E:132:PRO:HB2 | 1:E:202:VAL:HG23 | 1.99 | 0.44 |
| 1:E:402:TYR:CE2 | 1:E:425:ILE:HG12 | 2.53 | 0.44 |
| 1:H:377:ILE:HG12 | 1:H:499:PRO:HD3 | 2.00 | 0.44 |
| 1:I:149:PHE:H | 1:I:184:GLY:HA2 | 1.82 | 0.44 |
| 1:I:406:LYS:HD2 | 1:I:415:ASP:HA | 1.99 | 0.44 |
| 1:H:380:HIS:CE1 | 1:I:517:ARG:HD2 | 2.52 | 0.44 |
| 1:J:179:PRO:HG3 | 1:J:199:ALA:HB1 | 2.00 | 0.44 |
| 1:J:629:ILE:HD11 | 1:J:889:LEU:HB2 | 2.00 | 0.44 |
| 1:K:462:PRO:HD3 | 1:K:481:VAL:HG12 | 2.00 | 0.44 |
| 1:L:497:TRP:HH2 | 1:L:836:LYS:HG2 | 1.82 | 0.44 |
| 2:M:67:VAL:HG23 | 2:M:498:VAL:HB | 1.99 | 0.44 |
| 4:O:48:ARG:HA | 4:O:51:ARG:HG2 | 2.00 | 0.44 |
| 1:B:824:ILE:HG23 | 1:C:114:LYS:H | 1.83 | 0.44 |
| 1:D:506:ASN:HB3 | 1:D:509:ASN:H | 1.82 | 0.44 |
| 1:E:516:LEU:HD23 | 1:E:519:ARG:HH21 | 1.83 | 0.44 |
| 1:E:52:PRO:HD3 | 6:U:11:PRO:HA | 1.99 | 0.44 |
| 1:F:443:ARG:HH12 | 1:F:802:VAL:HG12 | 1.82 | 0.44 |
| 1:F:340:ARG:HD3 | 1:F:540:PHE:HE1 | 1.83 | 0.44 |
| 1:G:314:TYR:HE1 | 1:G:559:ARG:HE | 1.66 | 0.44 |
| 1:G:63:ARG:HD2 | 1:G:66:LEU:HB3 | 1.99 | 0.44 |
| 1:H:363:TRP:HZ3 | 1:H:536:PRO:HA | 1.82 | 0.44 |
| 1:I:228:SER:HB2 | 1:I:265:VAL:HA | 1.99 | 0.44 |
| 1:J:278:PRO:HG3 | 1:J:290:LEU:HA | 1.99 | 0.44 |
| 1:B:445:PHE:HE1 | 1:B:512:ARG:HH12 | 1.66 | 0.44 |
| 1:C:130:PRO:O | 1:C:144:ARG:NH2 | 2.50 | 0.44 |
| 1:E:174:ASP:H | 1:E:178:GLN:HB2 | 1.82 | 0.44 |
| 1:E:776:ARG:HD2 | 1:E:834:GLN:HE21 | 1.83 | 0.44 |
| 1:F:374:VAL:HG11 | 1:F:510:HIS:CE1 | 2.52 | 0.44 |
| 1:H:512:ARG:HG3 | 1:H:517:ARG:NH1 | 2.32 | 0.44 |
| 1:G:434:GLU:N | 1:I:391:PHE:O | 2.48 | 0.44 |
| 1:I:779:VAL:HG21 | 1:I:821:TYR:HD2 | 1.83 | 0.44 |

Continued on next page...

CONFIDENTIAL VALIDATION REPORT

Continued from previous page...

| Atom-1 | Atom-2 | Interatomic distance (Å) | Clash overlap (Å) |
|---|---|---|---|
| 1:J:901:ILE:HD12 | 1:J:911:ALA:HB2 | 2.00 | 0.44 |
| 1:K:109:ARG:HH22 | 1:K:523:LEU:HB2 | 1.83 | 0.44 |
| 1:L:506:ASN:ND2 | 1:L:677:SER:OG | 2.51 | 0.44 |
| 4:O:7:THR:HG22 | 4:O:26:ASP:HB3 | 2.00 | 0.44 |
| 1:B:679:PRO:HA | 1:B:684:THR:HB | 1.99 | 0.44 |
| 1:E:326:GLY:HA2 | 1:E:552:TYR:HD1 | 1.82 | 0.44 |
| 1:E:617:ASN:ND2 | 1:E:897:ASP:O | 2.50 | 0.44 |
| 1:F:360:PHE:HE2 | 1:F:363:TRP:H | 1.65 | 0.44 |
| 1:H:390:CYS:N | 1:H:433:MET:O | 2.45 | 0.44 |
| 1:H:736:LYS:HE2 | 1:I:589:TYR:CD2 | 2.52 | 0.44 |
| 1:J:241:ASP:N | 1:J:241:ASP:OD1 | 2.51 | 0.44 |
| 1:J:599:THR:HA | 1:J:602:THR:HG22 | 1.99 | 0.44 |
| 1:J:629:ILE:H | 1:J:888:LEU:HA | 1.83 | 0.44 |
| 1:K:376:ILE:HD13 | 1:K:498:SER:HB2 | 2.00 | 0.44 |
| 2:M:90:GLN:HE21 | 2:M:477:ARG:HH21 | 1.65 | 0.44 |
| 1:D:509:ASN:ND2 | 1:D:674:TYR:OH | 2.51 | 0.44 |
| 1:F:325:ALA:HB3 | 1:F:553:THR:HB | 2.00 | 0.44 |
| 1:G:130:PRO:HA | 1:G:211:PRO:HA | 1.99 | 0.44 |
| 1:H:483:VAL:HG11 | 1:H:792:LEU:HD21 | 2.00 | 0.44 |
| 1:I:280:VAL:HG22 | 1:I:282:GLN:H | 1.83 | 0.44 |
| 1:K:101:PHE:HD1 | 1:K:588:LEU:HD23 | 1.83 | 0.44 |
| 1:K:27:GLY:O | 1:K:31:PHE:N | 2.51 | 0.44 |
| 1:K:654:THR:OG1 | 1:K:655:ARG:N | 2.51 | 0.44 |
| 1:K:146:GLN:HA | 1:L:426:GLU:H | 1.83 | 0.44 |
| 1:L:626:LEU:HD12 | 1:L:888:LEU:HD11 | 2.00 | 0.44 |
| 1:J:61:SER:HA | 1:L:707:ASN:HB3 | 1.98 | 0.44 |
| 4:P:178:GLN:NE2 | 4:P:180:SER:OG | 2.48 | 0.44 |
| 1:B:28:LEU:O | 1:B:32:ALA:N | 2.49 | 0.43 |
| 1:D:109:ARG:HD3 | 1:D:113:PHE:CD1 | 2.53 | 0.43 |
| 1:D:587:ASN:N | 1:D:587:ASN:OD1 | 2.50 | 0.43 |
| 1:E:474:TYR:HE1 | 1:E:570:LEU:HD21 | 1.82 | 0.43 |
| 1:E:901:ILE:HG22 | 1:E:911:ALA:HA | 1.99 | 0.43 |
| 1:J:94:LEU:O | 1:J:547:LEU:N | 2.46 | 0.43 |
| 1:L:150:ILE:HA | 1:L:198:VAL:HG23 | 2.00 | 0.43 |
| 1:L:88:VAL:HB | 1:L:92:ARG:HB2 | 2.00 | 0.43 |
| 4:P:56:LEU:HD11 | 4:P:204:PHE:HE2 | 1.82 | 0.43 |
| 1:A:485:SER:O | 1:A:490:TYR:OH | 2.25 | 0.43 |
| 1:C:311:GLY:O | 1:C:559:ARG:NH1 | 2.51 | 0.43 |
| 1:D:374:VAL:HG11 | 1:D:510:HIS:CE1 | 2.53 | 0.43 |
| 1:E:730:ALA:HB2 | 1:F:362:MET:HB2 | 2.01 | 0.43 |
| 1:G:559:ARG:HB2 | 1:G:565:ILE:HD11 | 2.00 | 0.43 |

Continued on next page...

CONFIDENTIAL VALIDATION REPORT

*Continued from previous page...*

| Atom-1 | Atom-2 | Interatomic distance (Å) | Clash overlap (Å) |
|---|---|---|---|
| 1:H:506:ASN:HB3 | 1:H:686:TYR:CG | 2.53 | 0.43 |
| 1:J:818:ASN:HD22 | 1:K:219:PRO:HG3 | 1.83 | 0.43 |
| 1:L:134:GLU:HB3 | 1:L:143:VAL:HG12 | 2.00 | 0.43 |
| 1:A:276:TYR:OH | 1:B:187:GLN:OE1 | 2.35 | 0.43 |
| 1:A:347:GLN:HE22 | 1:A:356:ARG:NH2 | 2.16 | 0.43 |
| 1:B:895:VAL:HB | 1:B:917:PRO:HD2 | 2.00 | 0.43 |
| 1:C:375:ARG:HA | 1:C:502:MET:HE1 | 2.00 | 0.43 |
| 1:C:299:ARG:HH12 | 1:C:570:LEU:HB2 | 1.84 | 0.43 |
| 1:D:405:ILE:HD11 | 1:F:256:PRO:HB3 | 1.99 | 0.43 |
| 1:F:108:ASP:HB3 | 1:F:580:SER:H | 1.83 | 0.43 |
| 1:H:339:ASP:O | 1:H:624:ASN:N | 2.47 | 0.43 |
| 1:I:107:LEU:HD23 | 1:I:109:ARG:HH21 | 1.83 | 0.43 |
| 1:H:813:GLN:HE22 | 1:I:201:ARG:H | 1.64 | 0.43 |
| 1:I:658:THR:HA | 1:I:888:LEU:HD22 | 1.99 | 0.43 |
| 1:J:499:PRO:HG2 | 1:J:502:MET:HB3 | 2.00 | 0.43 |
| 1:J:601:SER:HA | 1:J:604:GLU:HG3 | 2.00 | 0.43 |
| 1:K:497:TRP:CD1 | 1:K:776:ARG:HD2 | 2.52 | 0.43 |
| 1:A:58:THR:HG22 | 1:A:596:ALA:HA | 2.01 | 0.43 |
| 1:B:204:LYS:HB2 | 1:B:263:GLU:H | 1.84 | 0.43 |
| 1:E:459:LYS:HG2 | 1:E:482:ALA:HB2 | 2.00 | 0.43 |
| 1:H:624:ASN:ND2 | 1:H:890:TYR:OH | 2.52 | 0.43 |
| 1:I:103:ILE:HG12 | 1:I:586:ILE:HG12 | 2.00 | 0.43 |
| 1:J:23:TYR:O | 1:L:611:THR:OG1 | 2.37 | 0.43 |
| 1:J:382:VAL:HG21 | 1:J:439:ALA:HB2 | 2.00 | 0.43 |
| 1:K:742:GLN:HE22 | 1:K:845:ARG:H | 1.66 | 0.43 |
| 1:J:433:MET:HG2 | 1:L:392:PRO:HA | 2.00 | 0.43 |
| 1:L:443:ARG:NE | 1:L:488:ASP:OD1 | 2.49 | 0.43 |
| 1:L:640:ILE:HD11 | 1:L:645:TRP:CZ2 | 2.54 | 0.43 |
| 1:A:637:PRO:HA | 1:A:877:THR:HA | 2.00 | 0.43 |
| 1:E:625:MET:HB2 | 1:E:891:VAL:HB | 1.99 | 0.43 |
| 1:E:98:SER:OG | 1:E:591:ASN:ND2 | 2.40 | 0.43 |
| 1:I:10:TRP:O | 1:I:14:HIS:N | 2.51 | 0.43 |
| 1:I:149:PHE:HD2 | 1:I:199:ALA:HB3 | 1.83 | 0.43 |
| 1:I:210:LEU:HD11 | 1:I:269:ALA:HB2 | 2.00 | 0.43 |
| 1:I:364:ASN:HD21 | 1:I:519:ARG:HD3 | 1.83 | 0.43 |
| 1:I:784:TYR:HD2 | 1:I:787:TYR:HB2 | 1.83 | 0.43 |
| 1:I:692:LYS:HE2 | 1:I:879:GLU:HG2 | 2.00 | 0.43 |
| 1:J:866:LEU:HD22 | 4:P:231:GLY:HA2 | 2.01 | 0.43 |
| 1:L:165:ASP:HB3 | 1:L:169:GLN:H | 1.83 | 0.43 |
| 1:J:115:PRO:HG2 | 1:L:494:GLY:HA3 | 2.01 | 0.43 |
| 3:N:139:PHE:HE2 | 3:N:145:LEU:HD23 | 1.83 | 0.43 |

*Continued on next page...*

CONFIDENTIAL VALIDATION REPORT

*Continued from previous page...*

| Atom-1 | Atom-2 | Interatomic distance (Å) | Clash overlap (Å) |
|---|---|---|---|
| 1:B:559:ARG:NH2 | 1:B:564:MET:O | 2.52 | 0.43 |
| 1:D:16:ALA:HB2 | 1:F:858:LEU:HD21 | 2.01 | 0.43 |
| 1:D:341:ASN:H | 1:D:624:ASN:HD22 | 1.67 | 0.43 |
| 1:D:509:ASN:ND2 | 1:D:676:GLY:O | 2.40 | 0.43 |
| 1:F:490:TYR:HD1 | 1:F:820:PRO:HD3 | 1.84 | 0.43 |
| 1:G:404:GLY:HA2 | 1:I:247:PHE:HA | 2.01 | 0.43 |
| 1:I:299:ARG:O | 1:I:474:TYR:OH | 2.32 | 0.43 |
| 1:J:677:SER:OG | 1:J:682:ASP:OD2 | 2.35 | 0.43 |
| 1:L:861:LEU:HA | 1:L:864:ASN:HB2 | 2.01 | 0.43 |
| 2:M:145:LEU:HD21 | 2:M:184:ILE:HD11 | 2.00 | 0.43 |
| 2:M:87:SER:HB3 | 2:M:491:VAL:HB | 2.01 | 0.43 |
| 4:P:44:VAL:HA | 4:P:47:ILE:HD12 | 2.01 | 0.43 |
| 1:C:784:TYR:HD2 | 1:C:787:TYR:HB2 | 1.83 | 0.43 |
| 1:D:503:ASP:OD2 | 1:D:836:LYS:NZ | 2.49 | 0.43 |
| 1:D:626:LEU:HD13 | 1:D:890:TYR:HD1 | 1.83 | 0.43 |
| 1:E:241:ASP:N | 1:E:241:ASP:OD1 | 2.51 | 0.43 |
| 1:F:497:TRP:CD1 | 1:F:776:ARG:HD2 | 2.53 | 0.43 |
| 1:H:314:TYR:HD1 | 1:H:559:ARG:HG3 | 1.83 | 0.43 |
| 1:H:784:TYR:HD2 | 1:H:787:TYR:HB2 | 1.84 | 0.43 |
| 1:I:909:ILE:O | 4:P:111:GLY:N | 2.38 | 0.43 |
| 1:K:376:ILE:HG13 | 1:K:496:ARG:HH21 | 1.83 | 0.43 |
| 1:J:823:LEU:HB3 | 1:K:526:GLY:HA2 | 2.01 | 0.43 |
| 4:O:178:GLN:NE2 | 4:O:180:SER:OG | 2.48 | 0.43 |
| 1:A:375:ARG:HD2 | 1:A:502:MET:HE3 | 2.00 | 0.43 |
| 1:B:146:GLN:HA | 1:C:426:GLU:H | 1.83 | 0.43 |
| 1:C:53:THR:OG1 | 1:C:54:HIS:N | 2.52 | 0.43 |
| 1:D:114:LYS:HE3 | 1:D:298:ASN:HB2 | 2.00 | 0.43 |
| 1:D:542:ILE:HG22 | 1:D:545:LEU:HD13 | 2.01 | 0.43 |
| 1:E:401:THR:HG21 | 1:E:421:ARG:HE | 1.84 | 0.43 |
| 1:E:490:TYR:HD1 | 1:E:820:PRO:HD3 | 1.84 | 0.43 |
| 1:E:559:ARG:HD3 | 1:E:561:ASP:HB3 | 1.99 | 0.43 |
| 1:E:750:GLY:HA2 | 1:E:754:PHE:HE1 | 1.83 | 0.43 |
| 1:F:686:TYR:HA | 1:F:840:ASP:HA | 2.00 | 0.43 |
| 1:H:389:TYR:HA | 1:H:434:GLU:HA | 2.00 | 0.43 |
| 1:H:454:LEU:HD22 | 1:H:501:PRO:HB2 | 2.01 | 0.43 |
| 1:I:114:LYS:HD2 | 1:I:298:ASN:HB2 | 1.99 | 0.43 |
| 1:J:131:ASN:ND2 | 1:J:210:LEU:O | 2.51 | 0.43 |
| 1:J:388:ASN:HB3 | 1:K:440:ASN:ND2 | 2.32 | 0.43 |
| 1:J:604:GLU:OE1 | 1:J:608:ARG:NH1 | 2.44 | 0.43 |
| 1:J:711:LEU:O | 1:K:63:ARG:NH1 | 2.50 | 0.43 |
| 1:A:402:TYR:O | 1:C:248:ALA:N | 2.52 | 0.43 |

*Continued on next page...*

CONFIDENTIAL VALIDATION REPORT

Continued from previous page...

| Atom-1 | Atom-2 | Interatomic distance (Å) | Clash overlap (Å) |
|---|---|---|---|
| 1:A:815:TYR:HE2 | 1:A:818:ASN:HD21 | 1.67 | 0.43 |
| 1:D:126:PRO:HB3 | 1:F:434:GLU:HG2 | 2.01 | 0.43 |
| 1:D:900:ARG:HH22 | 1:E:14:HIS:CE1 | 2.36 | 0.43 |
| 1:E:813:GLN:HA | 1:F:146:GLN:HE21 | 1.83 | 0.43 |
| 1:E:91:ASN:ND2 | 1:E:597:HIS:HD2 | 2.16 | 0.43 |
| 1:G:9:GLN:NE2 | 4:P:104:SER:OG | 2.52 | 0.43 |
| 1:H:467:LEU:HA | 1:H:468:PRO:HD3 | 1.90 | 0.43 |
| 1:I:175:LYS:HB3 | 1:I:242:VAL:HG21 | 2.01 | 0.43 |
| 1:I:823:LEU:HA | 1:I:829:VAL:HG12 | 2.01 | 0.43 |
| 1:J:483:VAL:HG21 | 1:J:792:LEU:HD11 | 2.00 | 0.43 |
| 2:M:55:TYR:OH | 2:M:107:GLN:OE1 | 2.30 | 0.43 |
| 1:A:649:ARG:HH21 | 1:A:894:GLU:HB2 | 1.83 | 0.43 |
| 1:B:626:LEU:HD21 | 1:B:680:TYR:HE1 | 1.83 | 0.43 |
| 1:B:780:ASN:HD21 | 1:B:783:THR:HG22 | 1.84 | 0.43 |
| 1:D:779:VAL:HG21 | 1:D:821:TYR:HD2 | 1.84 | 0.43 |
| 1:E:390:CYS:N | 1:E:433:MET:O | 2.51 | 0.43 |
| 1:F:845:ARG:NH2 | 1:F:894:GLU:OE2 | 2.52 | 0.43 |
| 1:G:474:TYR:HE1 | 1:G:570:LEU:HD21 | 1.84 | 0.43 |
| 1:G:660:GLU:HG2 | 1:G:674:TYR:CE2 | 2.54 | 0.43 |
| 1:H:375:ARG:HB2 | 1:H:507:PRO:HG3 | 2.00 | 0.43 |
| 1:G:429:ASN:ND2 | 1:H:812:GLY:H | 2.17 | 0.43 |
| 2:M:445:PRO:HA | 2:M:446:PRO:HD3 | 1.85 | 0.43 |
| 1:A:316:ASN:N | 1:A:336:ASP:OD2 | 2.51 | 0.42 |
| 1:A:792:LEU:HA | 1:A:795:GLN:HB3 | 2.01 | 0.42 |
| 1:B:315:TYR:OH | 1:B:538:LYS:N | 2.42 | 0.42 |
| 1:B:552:TYR:HE1 | 1:B:907:GLY:HA2 | 1.83 | 0.42 |
| 1:C:655:ARG:HH21 | 1:C:887:THR:HG21 | 1.84 | 0.42 |
| 1:D:274:LEU:HD22 | 1:D:293:GLN:HE21 | 1.84 | 0.42 |
| 1:D:362:MET:HG2 | 1:D:534:GLN:HG2 | 2.01 | 0.42 |
| 1:D:94:LEU:HD11 | 1:D:590:ALA:HB1 | 2.00 | 0.42 |
| 1:H:753:GLY:HA2 | 1:I:98:SER:HA | 2.00 | 0.42 |
| 1:G:533:ILE:HA | 1:I:731:GLN:HE21 | 1.84 | 0.42 |
| 1:K:823:LEU:HD23 | 1:L:526:GLY:HA3 | 2.00 | 0.42 |
| 1:D:521:MET:HB3 | 1:F:496:ARG:H | 1.84 | 0.42 |
| 1:G:427:SER:HB3 | 1:I:192:VAL:HG21 | 2.00 | 0.42 |
| 1:H:277:LYS:HA | 1:H:278:PRO:HD3 | 1.89 | 0.42 |
| 1:H:513:ASN:HB3 | 1:H:516:LEU:HB2 | 2.00 | 0.42 |
| 1:H:85:THR:HA | 1:H:553:THR:HG22 | 2.00 | 0.42 |
| 1:I:402:TYR:HD2 | 1:I:425:ILE:HD11 | 1.84 | 0.42 |
| 1:I:558:PHE:HE2 | 1:I:586:ILE:HG21 | 1.84 | 0.42 |
| 1:J:59:ASP:OD1 | 1:L:709:ARG:NH2 | 2.48 | 0.42 |

Continued on next page...

CONFIDENTIAL VALIDATION REPORT

Continued from previous page...

| Atom-1 | Atom-2 | Interatomic distance (Å) | Clash overlap (Å) |
|---|---|---|---|
| 1:L:243:ASN:O | 1:L:262:ALA:N | 2.51 | 0.42 |
| 1:L:380:HIS:CD2 | 1:L:383:GLU:HG2 | 2.54 | 0.42 |
| 1:L:453:TYR:HH | 1:L:511:HIS:HD1 | 1.62 | 0.42 |
| 4:O:91:LEU:HD11 | 4:O:175:LEU:HD23 | 2.01 | 0.42 |
| 1:A:324:LEU:HD23 | 1:A:554:TYR:HD1 | 1.84 | 0.42 |
| 1:C:113:PHE:HE1 | 1:C:303:ILE:HB | 1.84 | 0.42 |
| 1:C:722:VAL:HG11 | 1:E:668:PHE:HE2 | 1.84 | 0.42 |
| 1:D:379:ASN:ND2 | 1:D:489:THR:O | 2.49 | 0.42 |
| 1:D:58:THR:HG21 | 1:D:62:GLN:HE22 | 1.83 | 0.42 |
| 1:D:72:ASP:O | 1:D:82:VAL:HG12 | 2.20 | 0.42 |
| 1:E:324:LEU:HD13 | 1:E:554:TYR:HD1 | 1.84 | 0.42 |
| 1:E:607:LEU:HD21 | 1:E:614:GLN:HE22 | 1.83 | 0.42 |
| 1:H:621:CYS:HB2 | 1:H:897:ASP:H | 1.84 | 0.42 |
| 1:J:453:TYR:CZ | 1:J:510:HIS:HA | 2.55 | 0.42 |
| 1:J:696:ILE:HG12 | 1:J:876:MET:HG2 | 2.00 | 0.42 |
| 1:K:53:THR:HG23 | 1:K:54:HIS:CD2 | 2.54 | 0.42 |
| 1:K:767:PHE:HA | 1:K:842:THR:HG21 | 2.01 | 0.42 |
| 1:B:766:SER:OG | 1:B:767:PHE:N | 2.52 | 0.42 |
| 1:D:678:VAL:HA | 1:D:679:PRO:HD3 | 1.91 | 0.42 |
| 1:E:310:ILE:HG22 | 1:E:538:LYS:HE3 | 2.01 | 0.42 |
| 1:E:657:LYS:HB2 | 1:E:660:GLU:HB2 | 2.01 | 0.42 |
| 1:E:815:TYR:CZ | 1:F:181:PRO:HG2 | 2.54 | 0.42 |
| 1:F:784:TYR:HD2 | 1:F:787:TYR:HB2 | 1.84 | 0.42 |
| 1:I:245:GLN:HE22 | 1:I:262:ALA:HB2 | 1.83 | 0.42 |
| 1:I:729:VAL:HG11 | 1:I:739:PHE:CG | 2.54 | 0.42 |
| 1:J:405:ILE:HB | 1:J:413:THR:HG23 | 2.00 | 0.42 |
| 3:N:206:THR:HB | 4:O:205:ASN:HB3 | 2.00 | 0.42 |
| 4:O:44:VAL:HA | 4:O:47:ILE:HD12 | 2.01 | 0.42 |
| 1:A:141:ILE:HG13 | 1:B:418:TYR:HA | 2.01 | 0.42 |
| 1:C:149:PHE:O | 1:C:199:ALA:N | 2.41 | 0.42 |
| 1:C:678:VAL:HA | 1:C:679:PRO:HD3 | 1.90 | 0.42 |
| 1:D:96:MET:HA | 1:D:99:THR:HG23 | 2.00 | 0.42 |
| 1:E:758:GLU:HB2 | 1:E:761:LYS:HE2 | 2.00 | 0.42 |
| 1:E:898:VAL:HG11 | 1:F:13:MET:HB3 | 2.00 | 0.42 |
| 1:F:342:THR:HG21 | 1:F:539:PHE:HA | 2.01 | 0.42 |
| 1:G:340:ARG:HH11 | 1:G:915:ARG:HD3 | 1.85 | 0.42 |
| 1:G:526:GLY:HA3 | 1:I:777:GLN:HG3 | 2.00 | 0.42 |
| 1:H:204:LYS:HG2 | 1:H:205:ASP:H | 1.84 | 0.42 |
| 1:H:799:SER:OG | 1:I:181:PRO:O | 2.37 | 0.42 |
| 1:K:204:LYS:HB2 | 1:K:263:GLU:H | 1.84 | 0.42 |
| 1:L:661:THR:HG21 | 1:L:888:LEU:HD21 | 2.01 | 0.42 |

Continued on next page...

CONFIDENTIAL VALIDATION REPORT

Continued from previous page...

| Atom-1 | Atom-2 | Interatomic distance (Å) | Clash overlap (Å) |
|---|---|---|---|
| 4:P:48:ARG:HA | 4:P:51:ARG:HG2 | 2.00 | 0.42 |
| 1:A:243:ASN:H | 1:A:262:ALA:HB3 | 1.83 | 0.42 |
| 1:A:263:GLU:OE2 | 1:C:815:TYR:OH | 2.33 | 0.42 |
| 1:D:327:GLN:OE1 | 1:D:553:THR:OG1 | 2.25 | 0.42 |
| 1:D:497:TRP:CG | 1:D:776:ARG:HD3 | 2.54 | 0.42 |
| 1:D:509:ASN:OD1 | 1:D:567:GLN:NE2 | 2.52 | 0.42 |
| 1:D:510:HIS:CE1 | 1:D:512:ARG:HB2 | 2.53 | 0.42 |
| 1:F:445:PHE:O | 1:F:449:ASN:ND2 | 2.50 | 0.42 |
| 1:F:791:THR:HG22 | 1:F:793:PRO:HD2 | 2.01 | 0.42 |
| 1:H:210:LEU:HD12 | 1:H:211:PRO:HD2 | 2.00 | 0.42 |
| 1:G:248:ALA:HB3 | 1:H:403:SER:HB2 | 2.01 | 0.42 |
| 1:H:687:LEU:HD21 | 1:H:883:MET:HE1 | 2.01 | 0.42 |
| 1:I:134:GLU:HG2 | 1:I:143:VAL:HG12 | 1.99 | 0.42 |
| 1:E:910:GLU:CD | 4:O:93:ARG:HH11 | 2.22 | 0.42 |
| 1:A:24:LEU:HD23 | 1:A:24:LEU:HA | 1.85 | 0.42 |
| 1:B:443:ARG:HH21 | 1:B:484:PRO:HB2 | 1.84 | 0.42 |
| 1:B:305:PHE:HE2 | 1:B:523:LEU:HD21 | 1.85 | 0.42 |
| 1:B:788:GLN:HG3 | 1:C:222:GLU:HA | 2.02 | 0.42 |
| 1:F:315:TYR:HE2 | 1:F:538:LYS:HG3 | 1.84 | 0.42 |
| 1:H:151:GLY:HA3 | 1:H:160:ILE:HG22 | 2.02 | 0.42 |
| 1:H:503:ASP:OD2 | 1:H:836:LYS:NZ | 2.39 | 0.42 |
| 1:I:342:THR:HG21 | 1:I:539:PHE:HA | 2.01 | 0.42 |
| 1:J:814:ALA:H | 1:K:146:GLN:HE21 | 1.67 | 0.42 |
| 1:K:766:SER:OG | 1:K:767:PHE:N | 2.53 | 0.42 |
| 1:K:910:GLU:HG3 | 4:O:37:GLY:H | 1.85 | 0.42 |
| 2:M:274:LEU:O | 2:M:341:ARG:NH1 | 2.52 | 0.42 |
| 1:H:658:THR:O | 5:T:32:ARG:NH2 | 2.52 | 0.42 |
| 1:A:452:LEU:HA | 1:A:452:LEU:HD13 | 1.88 | 0.42 |
| 1:A:511:HIS:O | 1:A:517:ARG:NH2 | 2.53 | 0.42 |
| 1:A:808:THR:OG1 | 1:A:809:MET:N | 2.53 | 0.42 |
| 1:C:263:GLU:HB3 | 1:C:265:VAL:HG13 | 2.02 | 0.42 |
| 1:D:242:VAL:HG22 | 1:D:263:GLU:HG2 | 2.02 | 0.42 |
| 1:E:147:ALA:HB2 | 1:F:425:ILE:HB | 2.02 | 0.42 |
| 1:E:645:TRP:CD1 | 1:E:872:HIS:HB2 | 2.55 | 0.42 |
| 1:E:766:SER:OG | 1:E:767:PHE:N | 2.52 | 0.42 |
| 1:F:685:PHE:O | 1:F:688:ASN:ND2 | 2.53 | 0.42 |
| 2:M:420:SER:HA | 2:M:423:ILE:HB | 2.02 | 0.42 |
| 2:M:119:ALA:HB2 | 2:M:468:LEU:HD21 | 2.02 | 0.42 |
| 1:A:340:ARG:HH21 | 1:A:915:ARG:HD3 | 1.84 | 0.42 |
| 1:B:248:ALA:HB3 | 1:C:405:ILE:HD13 | 2.02 | 0.42 |
| 1:B:624:ASN:HD21 | 1:B:892:LEU:HD13 | 1.84 | 0.42 |

Continued on next page...

CONFIDENTIAL VALIDATION REPORT

Continued from previous page...

| Atom-1 | Atom-2 | Interatomic distance (Å) | Clash overlap (Å) |
|---|---|---|---|
| 1:E:498:SER:HA | 1:E:499:PRO:HD3 | 1.93 | 0.42 |
| 1:F:516:LEU:HA | 1:F:519:ARG:HB2 | 2.01 | 0.42 |
| 1:F:688:ASN:O | 1:F:844:TRP:NE1 | 2.37 | 0.42 |
| 1:G:356:ARG:HG3 | 1:G:367:VAL:HG12 | 2.02 | 0.42 |
| 1:G:752:GLN:NE2 | 1:H:95:ASP:OD1 | 2.53 | 0.42 |
| 1:H:510:HIS:CE1 | 1:H:512:ARG:HB3 | 2.55 | 0.42 |
| 1:K:53:THR:OG1 | 1:K:54:HIS:N | 2.51 | 0.42 |
| 2:M:181:ASN:HA | 2:M:184:ILE:HG22 | 2.01 | 0.42 |
| 2:M:321:LYS:HG2 | 2:M:327:SER:HB3 | 2.02 | 0.42 |
| 1:A:405:ILE:HD13 | 1:A:412:TRP:HD1 | 1.85 | 0.42 |
| 1:C:858:LEU:HD13 | 1:C:863:GLN:HB3 | 2.01 | 0.42 |
| 1:E:20:ALA:HA | 1:E:23:TYR:HE1 | 1.84 | 0.42 |
| 1:E:617:ASN:HD22 | 1:E:618:ASP:N | 2.18 | 0.42 |
| 1:E:712:THR:HA | 1:E:713:PRO:HD3 | 1.91 | 0.42 |
| 1:F:310:ILE:HD12 | 1:F:681:LEU:HB3 | 2.01 | 0.42 |
| 1:G:419:ALA:HB2 | 1:I:140:LYS:HB3 | 2.01 | 0.42 |
| 1:H:790:VAL:HG12 | 1:I:224:GLY:HA2 | 2.01 | 0.42 |
| 1:H:73:ARG:HG2 | 1:H:82:VAL:HG12 | 2.01 | 0.42 |
| 1:H:857:ALA:HB1 | 1:H:896:PHE:HE1 | 1.84 | 0.42 |
| 1:H:794:PHE:HB3 | 1:I:224:GLY:HA3 | 2.02 | 0.42 |
| 1:J:545:LEU:HD22 | 1:J:547:LEU:HD22 | 2.02 | 0.42 |
| 1:J:691:PHE:HB2 | 1:J:718:ILE:HG21 | 2.01 | 0.42 |
| 1:L:144:ARG:HD3 | 1:L:291:LEU:HD11 | 2.02 | 0.42 |
| 1:A:249:LEU:HD12 | 1:A:250:PRO:HD2 | 2.01 | 0.41 |
| 1:B:220:THR:OG1 | 1:B:227:ALA:O | 2.35 | 0.41 |
| 1:B:360:PHE:CE2 | 1:B:362:MET:HB3 | 2.55 | 0.41 |
| 1:B:515:GLY:O | 1:B:519:ARG:NE | 2.53 | 0.41 |
| 1:B:304:GLY:HA3 | 1:B:519:ARG:HG3 | 2.01 | 0.41 |
| 1:D:350:LEU:HD21 | 1:D:365:GLN:HE21 | 1.85 | 0.41 |
| 1:D:612:ASN:ND2 | 1:E:25:SER:H | 2.15 | 0.41 |
| 1:E:454:LEU:HD11 | 1:E:487:LEU:HD23 | 2.02 | 0.41 |
| 1:F:387:PRO:HB2 | 1:F:389:TYR:CZ | 2.54 | 0.41 |
| 1:G:159:GLY:HA2 | 1:G:244:LEU:HD21 | 2.01 | 0.41 |
| 1:G:517:ARG:HH11 | 1:I:380:HIS:CD2 | 2.38 | 0.41 |
| 1:H:101:PHE:O | 1:H:535:VAL:N | 2.52 | 0.41 |
| 1:L:784:TYR:HB2 | 1:L:830:PRO:HD2 | 2.02 | 0.41 |
| 2:M:277:GLY:HA3 | 2:M:346:ALA:HB2 | 2.01 | 0.41 |
| 3:N:196:PHE:CE2 | 4:O:208:SER:HB3 | 2.55 | 0.41 |
| 5:Q:21:LEU:HD13 | 5:S:21:LEU:HD22 | 2.02 | 0.41 |
| 1:A:861:LEU:HD12 | 1:A:867:TYR:HE2 | 1.86 | 0.41 |
| 1:B:348:LEU:HD12 | 1:B:619:TYR:HE2 | 1.85 | 0.41 |

Continued on next page...

CONFIDENTIAL VALIDATION REPORT

*Continued from previous page...*

| Atom-1 | Atom-2 | Interatomic distance (Å) | Clash overlap (Å) |
|---|---|---|---|
| 1:B:914:LEU:HD13 | 1:C:13:MET:HG3 | 2.02 | 0.41 |
| 1:A:427:SER:HB2 | 1:C:198:VAL:HG21 | 2.02 | 0.41 |
| 1:F:106:VAL:H | 1:F:582:ARG:HB3 | 1.85 | 0.41 |
| 1:G:764:MET:N | 1:G:764:MET:SD | 2.93 | 0.41 |
| 1:H:113:PHE:CE2 | 1:H:115:PRO:HG3 | 2.55 | 0.41 |
| 1:J:24:LEU:HD13 | 1:J:28:LEU:HD23 | 2.01 | 0.41 |
| 1:J:729:VAL:HG11 | 1:J:739:PHE:CG | 2.55 | 0.41 |
| 1:K:253:PRO:HB2 | 1:K:255:GLU:HG2 | 2.02 | 0.41 |
| 1:K:342:THR:OG1 | 1:K:538:LYS:O | 2.38 | 0.41 |
| 1:K:754:PHE:HB2 | 1:L:363:TRP:CZ3 | 2.55 | 0.41 |
| 1:C:305:PHE:HE2 | 1:C:523:LEU:HD21 | 1.85 | 0.41 |
| 1:B:777:GLN:HG3 | 1:C:526:GLY:H | 1.85 | 0.41 |
| 1:C:736:LYS:HA | 1:C:739:PHE:HB3 | 2.02 | 0.41 |
| 1:C:741:ILE:HG13 | 1:C:852:PHE:CD2 | 2.56 | 0.41 |
| 1:D:632:ASN:HB2 | 1:D:882:PRO:HB2 | 2.03 | 0.41 |
| 1:E:80:TYR:CE2 | 1:E:586:ILE:HD12 | 2.55 | 0.41 |
| 1:F:883:MET:HE1 | 1:F:887:THR:HG21 | 2.02 | 0.41 |
| 1:H:780:ASN:ND2 | 1:H:832:LEU:HB2 | 2.36 | 0.41 |
| 1:G:432:ALA:HB3 | 1:H:810:ARG:HE | 1.86 | 0.41 |
| 1:I:729:VAL:N | 1:I:734:MET:O | 2.54 | 0.41 |
| 1:K:508:PHE:CE2 | 1:K:684:THR:HG22 | 2.55 | 0.41 |
| 2:M:200:SER:O | 2:M:340:TYR:OH | 2.38 | 0.41 |
| 4:P:91:LEU:HD11 | 4:P:175:LEU:HD23 | 2.01 | 0.41 |
| 1:B:648:PHE:HB2 | 1:B:895:VAL:HG12 | 2.02 | 0.41 |
| 1:D:690:THR:HB | 1:D:881:ASP:HB3 | 2.02 | 0.41 |
| 1:E:562:VAL:HA | 1:E:565:ILE:HG22 | 2.02 | 0.41 |
| 1:F:340:ARG:HH21 | 1:F:915:ARG:HD3 | 1.86 | 0.41 |
| 1:J:210:LEU:HD12 | 1:J:211:PRO:HD2 | 2.02 | 0.41 |
| 1:J:79:SER:OG | 1:J:80:TYR:N | 2.52 | 0.41 |
| 1:K:23:TYR:HA | 4:P:181:SER:HB2 | 2.03 | 0.41 |
| 1:K:253:PRO:HB2 | 1:K:255:GLU:H | 1.85 | 0.41 |
| 1:K:795:GLN:H | 1:L:182:GLN:HE22 | 1.69 | 0.41 |
| 1:L:101:PHE:CD1 | 1:L:588:LEU:HD23 | 2.55 | 0.41 |
| 1:A:44:LYS:HE2 | 1:C:546:LEU:HB2 | 2.03 | 0.41 |
| 1:D:25:SER:HA | 1:D:26:PRO:HD3 | 1.93 | 0.41 |
| 1:D:94:LEU:HB3 | 1:D:547:LEU:HB2 | 2.02 | 0.41 |
| 1:H:508:PHE:CE2 | 1:H:684:THR:HG22 | 2.56 | 0.41 |
| 1:K:377:ILE:HG22 | 1:K:450:VAL:HG21 | 2.02 | 0.41 |
| 4:P:72:PRO:HG3 | 4:P:225:VAL:HG13 | 2.03 | 0.41 |
| 1:A:53:THR:HG23 | 1:A:54:HIS:CD2 | 2.56 | 0.41 |
| 1:A:813:GLN:HE22 | 1:B:200:GLY:HA3 | 1.85 | 0.41 |

*Continued on next page...*

CONFIDENTIAL VALIDATION REPORT

*Continued from previous page...*

| Atom-1 | Atom-2 | Interatomic distance (Å) | Clash overlap (Å) |
|---|---|---|---|
| 1:B:654:THR:OG1 | 1:B:655:ARG:N | 2.53 | 0.41 |
| 1:B:694:VAL:HB | 1:B:716:PHE:HB2 | 2.01 | 0.41 |
| 1:B:822:PRO:HG2 | 1:B:827:THR:HG21 | 2.03 | 0.41 |
| 1:D:574:LEU:HA | 1:D:574:LEU:HD23 | 1.89 | 0.41 |
| 1:E:711:LEU:HD12 | 1:E:727:TYR:HE2 | 1.85 | 0.41 |
| 1:F:14:HIS:HB3 | 1:F:48:PRO:HD3 | 2.03 | 0.41 |
| 1:L:650:GLY:HA2 | 1:L:847:PRO:HA | 2.02 | 0.41 |
| 1:K:644:ASN:HB3 | 4:O:28:SER:HB2 | 2.02 | 0.41 |
| 1:A:618:ASP:OD1 | 1:A:619:TYR:N | 2.54 | 0.41 |
| 1:A:736:LYS:HA | 1:A:739:PHE:HB3 | 2.03 | 0.41 |
| 1:B:717:GLU:OE1 | 1:B:720:ARG:NH2 | 2.53 | 0.41 |
| 1:E:316:ASN:ND2 | 1:E:342:THR:H | 2.18 | 0.41 |
| 1:H:317:SER:OG | 1:H:665:GLY:N | 2.53 | 0.41 |
| 1:H:254:ASN:ND2 | 1:I:411:THR:O | 2.48 | 0.41 |
| 1:I:340:ARG:NE | 1:I:897:ASP:OD2 | 2.54 | 0.41 |
| 1:J:44:LYS:HE3 | 1:L:546:LEU:HD22 | 2.03 | 0.41 |
| 1:L:393:LEU:HA | 1:L:393:LEU:HD22 | 1.97 | 0.41 |
| 1:L:80:TYR:CE2 | 1:L:586:ILE:HD13 | 2.56 | 0.41 |
| 1:J:46:ARG:HB2 | 1:L:616:PHE:HA | 2.01 | 0.41 |
| 1:E:85:THR:HG1 | 1:L:83:ARG:NH2 | 2.18 | 0.41 |
| 1:A:379:ASN:HD21 | 1:A:493:ILE:HD13 | 1.85 | 0.41 |
| 1:A:474:TYR:O | 1:A:478:ASN:N | 2.54 | 0.41 |
| 1:C:497:TRP:CD2 | 1:C:776:ARG:HD2 | 2.55 | 0.41 |
| 1:D:134:GLU:HB3 | 1:D:143:VAL:HG12 | 2.02 | 0.41 |
| 1:E:493:ILE:HD13 | 1:F:120:ALA:HB2 | 2.03 | 0.41 |
| 1:F:229:LEU:HA | 1:F:240:SER:HA | 2.03 | 0.41 |
| 1:F:382:VAL:HG11 | 1:F:439:ALA:HA | 2.03 | 0.41 |
| 1:F:324:LEU:HD23 | 1:F:554:TYR:HD1 | 1.85 | 0.41 |
| 1:G:900:ARG:NH1 | 1:H:12:TYR:O | 2.51 | 0.41 |
| 1:H:514:ALA:HA | 1:H:517:ARG:HG2 | 2.03 | 0.41 |
| 1:I:323:VAL:HG22 | 1:I:555:GLU:HB3 | 2.03 | 0.41 |
| 1:I:351:ASP:HB3 | 1:I:763:ARG:HB3 | 2.02 | 0.41 |
| 1:I:506:ASN:HB3 | 1:I:686:TYR:CE1 | 2.56 | 0.41 |
| 1:L:435:ILE:HG13 | 1:L:435:ILE:H | 1.77 | 0.41 |
| 1:A:122:ASN:HB3 | 1:A:125:ALA:HB2 | 2.03 | 0.41 |
| 1:A:629:ILE:HG13 | 1:A:636:VAL:HG11 | 2.02 | 0.41 |
| 1:B:246:PHE:HA | 1:B:259:VAL:HG12 | 2.02 | 0.41 |
| 1:C:275:VAL:HG23 | 1:C:462:PRO:HG2 | 2.03 | 0.41 |
| 1:D:25:SER:H | 1:F:612:ASN:ND2 | 2.17 | 0.41 |
| 1:D:72:ASP:N | 1:D:72:ASP:OD1 | 2.54 | 0.41 |
| 1:E:406:LYS:N | 1:E:414:ALA:O | 2.54 | 0.41 |

*Continued on next page...*

CONFIDENTIAL VALIDATION REPORT

*Continued from previous page...*

| Atom-1 | Atom-2 | Interatomic distance (Å) | Clash overlap (Å) |
|---|---|---|---|
| 1:E:650:GLY:HA3 | 1:E:847:PRO:HA | 2.02 | 0.41 |
| 1:F:485:SER:OG | 1:F:490:TYR:OH | 2.32 | 0.41 |
| 1:G:374:VAL:HG11 | 1:G:510:HIS:CE1 | 2.50 | 0.41 |
| 1:G:424:GLU:O | 1:I:145:GLY:N | 2.48 | 0.41 |
| 1:I:211:PRO:HG2 | 1:I:295:ALA:HB2 | 2.03 | 0.41 |
| 1:J:497:TRP:CD2 | 1:J:776:ARG:HG3 | 2.56 | 0.41 |
| 1:J:868:ALA:O | 4:P:14:GLN:NE2 | 2.44 | 0.41 |
| 1:J:825:GLY:O | 1:K:527:ARG:NH1 | 2.53 | 0.41 |
| 1:K:558:PHE:HE2 | 1:K:586:ILE:HG21 | 1.86 | 0.41 |
| 1:K:86:LEU:HD23 | 1:K:554:TYR:HB3 | 2.02 | 0.41 |
| 1:L:19:ASP:HA | 1:L:48:PRO:HD2 | 2.03 | 0.41 |
| 1:J:435:ILE:HG13 | 1:L:390:CYS:HB3 | 2.03 | 0.41 |
| 1:L:629:ILE:HD13 | 1:L:889:LEU:HD12 | 2.02 | 0.41 |
| 2:M:126:ARG:HG3 | 2:M:493:LYS:HB3 | 2.01 | 0.41 |
| 2:M:143:ALA:HA | 2:M:244:VAL:HA | 2.03 | 0.41 |
| 1:A:388:ASN:HD21 | 1:A:437:LEU:HB2 | 1.85 | 0.41 |
| 1:B:378:GLU:HG2 | 1:B:380:HIS:CE1 | 2.55 | 0.41 |
| 1:B:490:TYR:HD1 | 1:B:820:PRO:HD3 | 1.86 | 0.41 |
| 1:B:779:VAL:HG21 | 1:B:821:TYR:HD2 | 1.86 | 0.41 |
| 1:D:705:PRO:HB3 | 1:D:852:PHE:HZ | 1.86 | 0.41 |
| 1:D:440:ASN:HD21 | 1:F:388:ASN:HA | 1.85 | 0.41 |
| 1:H:109:ARG:HH22 | 1:H:523:LEU:HB2 | 1.86 | 0.41 |
| 1:I:220:THR:H | 1:I:226:GLN:HA | 1.86 | 0.41 |
| 1:J:204:LYS:HB2 | 1:J:262:ALA:HB1 | 2.02 | 0.41 |
| 1:J:874:LEU:HD13 | 1:J:876:MET:HG3 | 2.03 | 0.41 |
| 1:K:81:LYS:HE3 | 1:K:557:ASN:HD21 | 1.85 | 0.41 |
| 1:L:490:TYR:HE2 | 1:L:819:TYR:HD2 | 1.68 | 0.41 |
| 3:N:238:PRO:HA | 3:N:241:ARG:HB2 | 2.03 | 0.41 |
| 4:P:88:THR:HA | 4:P:176:THR:HG22 | 2.03 | 0.41 |
| 1:H:628:PRO:HB3 | 5:T:25:LEU:HD22 | 2.02 | 0.41 |
| 1:A:301:ASN:HB2 | 1:A:569:SER:H | 1.86 | 0.41 |
| 1:A:377:ILE:HD11 | 1:A:450:VAL:HG11 | 2.03 | 0.41 |
| 1:A:573:ASP:HB3 | 1:A:576:VAL:HG12 | 2.03 | 0.41 |
| 1:A:730:ALA:HB2 | 1:B:362:MET:HB2 | 2.03 | 0.41 |
| 1:A:865:MET:O | 1:A:869:ASN:ND2 | 2.53 | 0.41 |
| 1:C:332:ASN:ND2 | 1:C:334:VAL:O | 2.54 | 0.41 |
| 1:E:453:TYR:HD2 | 1:E:510:HIS:HD1 | 1.68 | 0.41 |
| 1:G:343:GLU:HG3 | 1:G:681:LEU:HD22 | 2.02 | 0.41 |
| 1:J:617:ASN:ND2 | 1:K:46:ARG:HE | 2.19 | 0.41 |
| 2:M:118:GLY:O | 2:M:501:LYS:N | 2.54 | 0.41 |
| 1:A:375:ARG:HB2 | 1:A:507:PRO:HG2 | 2.04 | 0.40 |

*Continued on next page...*

CONFIDENTIAL VALIDATION REPORT

*Continued from previous page...*

| Atom-1 | Atom-2 | Interatomic distance (Å) | Clash overlap (Å) |
|---|---|---|---|
| 1:A:399:THR:H | 1:A:426:GLU:HG2 | 1.86 | 0.40 |
| 1:A:914:LEU:HD13 | 1:B:13:MET:HG3 | 2.03 | 0.40 |
| 1:B:32:ALA:HA | 1:B:41:LEU:HD23 | 2.03 | 0.40 |
| 1:D:192:VAL:HG23 | 1:D:196:GLN:HB2 | 2.02 | 0.40 |
| 1:F:631:SER:HB2 | 1:F:886:PRO:HG3 | 2.03 | 0.40 |
| 1:F:795:GLN:HA | 1:F:818:ASN:HD21 | 1.86 | 0.40 |
| 1:G:604:GLU:OE2 | 1:G:608:ARG:NH2 | 2.54 | 0.40 |
| 1:I:559:ARG:HH22 | 1:I:664:LEU:HD21 | 1.86 | 0.40 |
| 1:K:210:LEU:HD13 | 1:K:269:ALA:HB2 | 2.02 | 0.40 |
| 1:L:255:GLU:HA | 1:L:256:PRO:HD3 | 1.91 | 0.40 |
| 1:L:487:LEU:HB2 | 1:L:491:VAL:HG11 | 2.03 | 0.40 |
| 1:L:84:PHE:N | 1:L:554:TYR:O | 2.45 | 0.40 |
| 4:O:72:PRO:HG3 | 4:O:225:VAL:HG13 | 2.03 | 0.40 |
| 1:A:119:THR:OG1 | 1:A:120:ALA:N | 2.55 | 0.40 |
| 1:A:287:SER:HB2 | 1:A:291:LEU:HG | 2.04 | 0.40 |
| 1:A:302:TYR:HB2 | 1:A:516:LEU:HD13 | 2.02 | 0.40 |
| 1:A:322:GLY:HA3 | 1:A:556:TRP:CE3 | 2.57 | 0.40 |
| 1:E:271:ASP:HA | 1:E:298:ASN:ND2 | 2.36 | 0.40 |
| 1:E:860:ASP:OD2 | 1:F:57:THR:OG1 | 2.36 | 0.40 |
| 1:F:521:MET:HB3 | 1:F:521:MET:HE2 | 1.91 | 0.40 |
| 1:G:732:CYS:HA | 1:G:835:LYS:HB3 | 2.03 | 0.40 |
| 1:H:380:HIS:HE1 | 1:I:517:ARG:HD2 | 1.85 | 0.40 |
| 1:H:94:LEU:HD23 | 1:H:592:PHE:CD1 | 2.56 | 0.40 |
| 1:I:88:VAL:HG23 | 1:I:549:PRO:HA | 2.02 | 0.40 |
| 1:K:246:PHE:HB2 | 1:L:405:ILE:HG21 | 2.03 | 0.40 |
| 1:K:621:CYS:HB2 | 1:K:649:ARG:NH2 | 2.36 | 0.40 |
| 1:K:88:VAL:HG13 | 1:K:549:PRO:HA | 2.04 | 0.40 |
| 1:L:21:SER:OG | 1:L:33:ARG:NH2 | 2.54 | 0.40 |
| 1:K:857:ALA:N | 1:L:49:THR:OG1 | 2.51 | 0.40 |
| 1:L:652:SER:HB2 | 1:L:892:LEU:HB2 | 2.02 | 0.40 |
| 1:C:348:LEU:HD12 | 1:C:619:TYR:HE2 | 1.86 | 0.40 |
| 1:D:350:LEU:HD11 | 1:D:365:GLN:HG3 | 2.03 | 0.40 |
| 1:D:360:PHE:CE2 | 1:D:362:MET:HB3 | 2.57 | 0.40 |
| 1:D:343:GLU:HG3 | 1:D:681:LEU:HD22 | 2.02 | 0.40 |
| 1:D:736:LYS:HZ2 | 1:E:589:TYR:HB3 | 1.87 | 0.40 |
| 1:E:258:ALA:HB1 | 1:F:425:ILE:HD11 | 2.02 | 0.40 |
| 1:E:380:HIS:CE1 | 1:F:521:MET:HG3 | 2.56 | 0.40 |
| 1:E:796:HIS:CD2 | 1:E:817:ALA:HA | 2.57 | 0.40 |
| 1:F:161:GLN:HA | 1:F:172:TYR:HA | 2.03 | 0.40 |
| 1:F:448:SER:HA | 1:F:452:LEU:HD12 | 2.03 | 0.40 |
| 1:H:201:ARG:HD2 | 1:H:261:TYR:HB2 | 2.03 | 0.40 |

*Continued on next page...*

CONFIDENTIAL VALIDATION REPORT

*Continued from previous page...*

| Atom-1 | Atom-2 | Interatomic distance (Å) | Clash overlap (Å) |
|---|---|---|---|
| 1:H:813:GLN:HB3 | 1:I:148:PRO:HD3 | 2.03 | 0.40 |
| 1:J:151:GLY:N | 1:J:197:LYS:O | 2.55 | 0.40 |
| 1:J:317:SER:N | 1:J:336:ASP:OD2 | 2.46 | 0.40 |
| 1:K:389:TYR:HA | 1:K:434:GLU:HA | 2.03 | 0.40 |
| 2:M:149:LYS:HB3 | 2:M:157:PRO:HB3 | 2.02 | 0.40 |
| 2:M:279:ILE:HD11 | 2:M:342:SER:HB2 | 2.02 | 0.40 |
| 1:A:201:ARG:HD2 | 1:A:261:TYR:HB2 | 2.03 | 0.40 |
| 1:C:625:MET:HB3 | 1:C:627:TYR:CE2 | 2.56 | 0.40 |
| 1:C:62:GLN:HB3 | 1:C:92:ARG:HH12 | 1.86 | 0.40 |
| 1:D:474:TYR:HB2 | 1:D:572:ASN:HD22 | 1.86 | 0.40 |
| 1:E:202:VAL:HG11 | 1:E:260:LEU:HD13 | 2.02 | 0.40 |
| 1:D:145:GLY:N | 1:E:424:GLU:O | 2.36 | 0.40 |
| 1:G:627:TYR:CE2 | 1:G:638:ILE:HG13 | 2.56 | 0.40 |
| 1:I:678:VAL:HA | 1:I:679:PRO:HD3 | 1.98 | 0.40 |
| 1:J:440:ASN:HD21 | 1:L:388:ASN:HB2 | 1.85 | 0.40 |
| 1:L:447:TYR:HA | 1:L:451:ALA:HB3 | 2.03 | 0.40 |
| 1:L:743:MET:HG3 | 1:L:749:ILE:HB | 2.03 | 0.40 |
| 3:N:194:ASP:N | 3:N:194:ASP:OD1 | 2.54 | 0.40 |
| 1:C:8:PRO:HD3 | 3:N:67:GLU:OE2 | 2.22 | 0.40 |
| 4:O:47:ILE:O | 4:O:50:THR:OG1 | 2.34 | 0.40 |
| 1:A:115:PRO:HD2 | 1:C:824:ILE:HG12 | 2.04 | 0.40 |
| 1:B:134:GLU:HA | 1:B:143:VAL:HG12 | 2.03 | 0.40 |
| 1:D:559:ARG:HD3 | 1:D:561:ASP:HB3 | 2.03 | 0.40 |
| 1:E:640:ILE:HD13 | 1:E:643:ARG:NH1 | 2.37 | 0.40 |
| 1:F:355:ASP:HA | 1:F:763:ARG:HH22 | 1.86 | 0.40 |
| 1:F:426:GLU:HG2 | 1:F:427:SER:H | 1.86 | 0.40 |
| 1:G:114:LYS:HA | 1:G:115:PRO:HD3 | 1.82 | 0.40 |
| 1:H:741:ILE:HG21 | 1:H:741:ILE:HD13 | 1.86 | 0.40 |
| 1:I:109:ARG:HD3 | 1:I:113:PHE:CE1 | 2.57 | 0.40 |
| 1:I:503:ASP:OD1 | 1:I:689:HIS:NE2 | 2.45 | 0.40 |
| 1:I:632:ASN:HB2 | 1:I:882:PRO:HB2 | 2.04 | 0.40 |
| 1:J:664:LEU:HD12 | 1:J:680:TYR:HE2 | 1.86 | 0.40 |
| 1:K:103:ILE:HA | 1:K:586:ILE:HG12 | 2.02 | 0.40 |
| 1:K:18:GLN:NE2 | 4:P:184:PRO:O | 2.54 | 0.40 |
| 1:K:712:THR:HA | 1:K:713:PRO:HD3 | 1.90 | 0.40 |
| 1:K:691:PHE:HB2 | 1:K:718:ILE:HG21 | 2.03 | 0.40 |
| 2:M:175:MET:O | 2:M:179:LEU:N | 2.55 | 0.40 |

There are no symmetry-related clashes.

CONFIDENTIAL VALIDATION REPORT

#### 5.3 Torsion angles (i)

##### 5.3.1 Protein backbone (i)

In the following table, the Percentiles column shows the percent Ramachandran outliers of the chain as a percentile score with respect to all PDB entries followed by that with respect to all EM entries.

The Analysed column shows the number of residues for which the backbone conformation was analysed, and the total number of residues.

| Mol | Chain | Analysed | Favoured | Allowed | Outliers | Percentiles | |
|---|---|---|---|---|---|---|---|
| 1 | A | 908/925 (98%) | 776 (86%) | 132 (14%) | 0 | 100 | 100 |
| 1 | B | 909/925 (98%) | 780 (86%) | 129 (14%) | 0 | 100 | 100 |
| 1 | C | 910/925 (98%) | 792 (87%) | 118 (13%) | 0 | 100 | 100 |
| 1 | D | 904/925 (98%) | 759 (84%) | 145 (16%) | 0 | 100 | 100 |
| 1 | E | 913/925 (99%) | 779 (85%) | 132 (14%) | 2 (0%) | 49 | 84 |
| 1 | F | 906/925 (98%) | 764 (84%) | 139 (15%) | 3 (0%) | 43 | 80 |
| 1 | G | 906/925 (98%) | 760 (84%) | 144 (16%) | 2 (0%) | 49 | 84 |
| 1 | H | 911/925 (98%) | 761 (84%) | 145 (16%) | 5 (0%) | 31 | 72 |
| 1 | I | 904/925 (98%) | 751 (83%) | 151 (17%) | 2 (0%) | 49 | 84 |
| 1 | J | 910/925 (98%) | 761 (84%) | 148 (16%) | 1 (0%) | 53 | 87 |
| 1 | K | 908/925 (98%) | 760 (84%) | 146 (16%) | 2 (0%) | 49 | 84 |
| 1 | L | 905/925 (98%) | 769 (85%) | 136 (15%) | 0 | 100 | 100 |
| 2 | M | 448/508 (88%) | 406 (91%) | 42 (9%) | 0 | 100 | 100 |
| 3 | N | 296/579 (51%) | 272 (92%) | 24 (8%) | 0 | 100 | 100 |
| 4 | O | 176/233 (76%) | 160 (91%) | 16 (9%) | 0 | 100 | 100 |
| 4 | P | 176/233 (76%) | 160 (91%) | 16 (9%) | 0 | 100 | 100 |
| 5 | Q | 48/133 (36%) | 39 (81%) | 9 (19%) | 0 | 100 | 100 |
| 5 | R | 48/133 (36%) | 39 (81%) | 9 (19%) | 0 | 100 | 100 |
| 5 | S | 48/133 (36%) | 39 (81%) | 9 (19%) | 0 | 100 | 100 |
| 5 | T | 48/133 (36%) | 39 (81%) | 9 (19%) | 0 | 100 | 100 |
| 6 | U | 27/266 (10%) | 26 (96%) | 1 (4%) | 0 | 100 | 100 |
| 6 | V | 27/266 (10%) | 26 (96%) | 1 (4%) | 0 | 100 | 100 |
| 6 | Y | 27/266 (10%) | 26 (96%) | 1 (4%) | 0 | 100 | 100 |
| 6 | u | 27/266 (10%) | 26 (96%) | 1 (4%) | 0 | 100 | 100 |
| 7 | W | 4/183 (2%) | 2 (50%) | 2 (50%) | 0 | 100 | 100 |

*Continued on next page...*

CONFIDENTIAL VALIDATION REPORT

*Continued from previous page...*

| Mol | Chain | Analysed | Favoured | Allowed | Outliers | Percentiles | |
|---|---|---|---|---|---|---|---|
| 7 | w | 10/183 (6%) | 6 (60%) | 4 (40%) | 0 | 100 | 100 |
| All | All | 12304/14615 (84%) | 10478 (85%) | 1809 (15%) | 17 (0%) | 56 | 87 |

All (17) Ramachandran outliers are listed below:

| Mol | Chain | Res | Type |
|---|---|---|---|
| 1 | G | 917 | PRO |
| 1 | H | 320 | ASN |
| 1 | F | 630 | PRO |
| 1 | H | 816 | PRO |
| 1 | E | 630 | PRO |
| 1 | F | 385 | GLU |
| 1 | F | 816 | PRO |
| 1 | K | 150 | ILE |
| 1 | H | 665 | GLY |
| 1 | J | 757 | PRO |
| 1 | E | 917 | PRO |
| 1 | G | 287 | SER |
| 1 | H | 148 | PRO |
| 1 | H | 764 | MET |
| 1 | I | 278 | PRO |
| 1 | I | 757 | PRO |
| 1 | K | 757 | PRO |

##### 5.3.2 Protein sidechains ⓘ

In the following table, the Percentiles column shows the percent sidechain outliers of the chain as a percentile score with respect to all PDB entries followed by that with respect to all EM entries.

The Analysed column shows the number of residues for which the sidechain conformation was analysed, and the total number of residues.

| Mol | Chain | Analysed | Rotameric | Outliers | Percentiles | |
|---|---|---|---|---|---|---|
| 1 | A | 787/797 (99%) | 776 (99%) | 11 (1%) | 69 | 85 |
| 1 | B | 788/797 (99%) | 779 (99%) | 9 (1%) | 76 | 88 |
| 1 | C | 788/797 (99%) | 777 (99%) | 11 (1%) | 69 | 85 |
| 1 | D | 783/797 (98%) | 771 (98%) | 12 (2%) | 67 | 84 |
| 1 | E | 790/797 (99%) | 780 (99%) | 10 (1%) | 71 | 86 |

*Continued on next page...*

CONFIDENTIAL VALIDATION REPORT

*Continued from previous page...*

| Mol | Chain | Analysed | Rotameric | Outliers | Percentiles | |
|---|---|---|---|---|---|---|
| 1 | F | 785/797 (98%) | 771 (98%) | 14 (2%) | 62 | 83 |
| 1 | G | 785/797 (98%) | 773 (98%) | 12 (2%) | 67 | 84 |
| 1 | H | 789/797 (99%) | 776 (98%) | 13 (2%) | 65 | 84 |
| 1 | I | 783/797 (98%) | 777 (99%) | 6 (1%) | 83 | 92 |
| 1 | J | 788/797 (99%) | 772 (98%) | 16 (2%) | 58 | 80 |
| 1 | K | 787/797 (99%) | 776 (99%) | 11 (1%) | 69 | 85 |
| 1 | L | 787/797 (99%) | 779 (99%) | 8 (1%) | 78 | 89 |
| 2 | M | 403/447 (90%) | 398 (99%) | 5 (1%) | 74 | 87 |
| 3 | N | 253/501 (50%) | 251 (99%) | 2 (1%) | 83 | 92 |
| 4 | O | 149/193 (77%) | 147 (99%) | 2 (1%) | 71 | 86 |
| 4 | P | 149/193 (77%) | 147 (99%) | 2 (1%) | 71 | 86 |
| 5 | Q | 38/97 (39%) | 36 (95%) | 2 (5%) | 25 | 58 |
| 5 | R | 38/97 (39%) | 36 (95%) | 2 (5%) | 25 | 58 |
| 5 | S | 38/97 (39%) | 36 (95%) | 2 (5%) | 25 | 58 |
| 5 | T | 38/97 (39%) | 36 (95%) | 2 (5%) | 25 | 58 |
| 6 | U | 22/227 (10%) | 22 (100%) | 0 | 100 | 100 |
| 6 | V | 22/227 (10%) | 22 (100%) | 0 | 100 | 100 |
| 6 | Y | 22/227 (10%) | 22 (100%) | 0 | 100 | 100 |
| 6 | u | 22/227 (10%) | 22 (100%) | 0 | 100 | 100 |
| 7 | W | 2/139 (1%) | 2 (100%) | 0 | 100 | 100 |
| 7 | w | 6/139 (4%) | 6 (100%) | 0 | 100 | 100 |
| All | All | 10642/12472 (85%) | 10490 (99%) | 152 (1%) | 71 | 85 |

All (152) residues with a non-rotameric sidechain are listed below:

| Mol | Chain | Res | Type |
|---|---|---|---|
| 1 | A | 47 | ASN |
| 1 | A | 138 | ASN |
| 1 | A | 155 | ASN |
| 1 | A | 156 | LYS |
| 1 | A | 218 | LYS |
| 1 | A | 308 | ASN |
| 1 | A | 408 | ASN |
| 1 | A | 417 | ASN |
| 1 | A | 538 | LYS |

*Continued on next page...*

CONFIDENTIAL VALIDATION REPORT

*Continued from previous page...*

| Mol | Chain | Res | Type |
|---|---|---|---|
| 1 | A | 776 | ARG |
| 1 | A | 810 | ARG |
| 1 | B | 153 | ASN |
| 1 | B | 218 | LYS |
| 1 | B | 308 | ASN |
| 1 | B | 367 | VAL |
| 1 | B | 408 | ASN |
| 1 | B | 417 | ASN |
| 1 | B | 625 | MET |
| 1 | B | 810 | ARG |
| 1 | B | 818 | ASN |
| 1 | C | 6 | MET |
| 1 | C | 47 | ASN |
| 1 | C | 83 | ARG |
| 1 | C | 158 | ASN |
| 1 | C | 218 | LYS |
| 1 | C | 308 | ASN |
| 1 | C | 335 | VAL |
| 1 | C | 625 | MET |
| 1 | C | 774 | MET |
| 1 | C | 810 | ARG |
| 1 | C | 835 | LYS |
| 1 | D | 7 | MET |
| 1 | D | 92 | ARG |
| 1 | D | 122 | ASN |
| 1 | D | 138 | ASN |
| 1 | D | 153 | ASN |
| 1 | D | 243 | ASN |
| 1 | D | 308 | ASN |
| 1 | D | 388 | ASN |
| 1 | D | 644 | ASN |
| 1 | D | 797 | ASN |
| 1 | D | 810 | ARG |
| 1 | D | 869 | ASN |
| 1 | E | 88 | VAL |
| 1 | E | 153 | ASN |
| 1 | E | 232 | ASN |
| 1 | E | 388 | ASN |
| 1 | E | 478 | ASN |
| 1 | E | 572 | ASN |
| 1 | E | 617 | ASN |
| 1 | E | 798 | ASN |

*Continued on next page...*

CONFIDENTIAL VALIDATION REPORT

*Continued from previous page...*

| Mol | Chain | Res | Type |
|---|---|---|---|
| 1 | E | 810 | ARG |
| 1 | E | 864 | ASN |
| 1 | F | 6 | MET |
| 1 | F | 67 | ARG |
| 1 | F | 168 | ASN |
| 1 | F | 323 | VAL |
| 1 | F | 335 | VAL |
| 1 | F | 341 | ASN |
| 1 | F | 408 | ASN |
| 1 | F | 417 | ASN |
| 1 | F | 436 | ASN |
| 1 | F | 478 | ASN |
| 1 | F | 609 | ASN |
| 1 | F | 632 | ASN |
| 1 | F | 774 | MET |
| 1 | F | 810 | ARG |
| 1 | G | 47 | ASN |
| 1 | G | 142 | LYS |
| 1 | G | 144 | ARG |
| 1 | G | 308 | ASN |
| 1 | G | 341 | ASN |
| 1 | G | 388 | ASN |
| 1 | G | 408 | ASN |
| 1 | G | 470 | ASN |
| 1 | G | 478 | ASN |
| 1 | G | 598 | ASN |
| 1 | G | 774 | MET |
| 1 | G | 810 | ARG |
| 1 | H | 136 | LYS |
| 1 | H | 153 | ASN |
| 1 | H | 158 | ASN |
| 1 | H | 308 | ASN |
| 1 | H | 341 | ASN |
| 1 | H | 388 | ASN |
| 1 | H | 399 | THR |
| 1 | H | 408 | ASN |
| 1 | H | 421 | ARG |
| 1 | H | 470 | ASN |
| 1 | H | 797 | ASN |
| 1 | H | 798 | ASN |
| 1 | H | 810 | ARG |
| 1 | I | 144 | ARG |

*Continued on next page...*

CONFIDENTIAL VALIDATION REPORT

*Continued from previous page...*

| Mol | Chain | Res | Type |
|---|---|---|---|
| 1 | I | 324 | LEU |
| 1 | I | 341 | ASN |
| 1 | I | 417 | ASN |
| 1 | I | 810 | ARG |
| 1 | I | 864 | ASN |
| 1 | J | 6 | MET |
| 1 | J | 144 | ARG |
| 1 | J | 153 | ASN |
| 1 | J | 155 | ASN |
| 1 | J | 158 | ASN |
| 1 | J | 168 | ASN |
| 1 | J | 308 | ASN |
| 1 | J | 341 | ASN |
| 1 | J | 379 | ASN |
| 1 | J | 470 | ASN |
| 1 | J | 563 | ASN |
| 1 | J | 572 | ASN |
| 1 | J | 636 | VAL |
| 1 | J | 774 | MET |
| 1 | J | 810 | ARG |
| 1 | J | 861 | LEU |
| 1 | K | 43 | ASN |
| 1 | K | 67 | ARG |
| 1 | K | 144 | ARG |
| 1 | K | 242 | VAL |
| 1 | K | 308 | ASN |
| 1 | K | 341 | ASN |
| 1 | K | 436 | ASN |
| 1 | K | 470 | ASN |
| 1 | K | 478 | ASN |
| 1 | K | 810 | ARG |
| 1 | K | 864 | ASN |
| 1 | L | 122 | ASN |
| 1 | L | 221 | ASN |
| 1 | L | 341 | ASN |
| 1 | L | 644 | ASN |
| 1 | L | 649 | ARG |
| 1 | L | 714 | ASN |
| 1 | L | 780 | ASN |
| 1 | L | 810 | ARG |
| 2 | M | 126 | ARG |
| 2 | M | 131 | ASN |

*Continued on next page...*

CONFIDENTIAL VALIDATION REPORT

*Continued from previous page...*

| Mol | Chain | Res | Type |
|---|---|---|---|
| 2 | M | 223 | MET |
| 2 | M | 238 | LEU |
| 2 | M | 486 | ARG |
| 3 | N | 49 | ASN |
| 3 | N | 201 | ARG |
| 4 | O | 32 | ASN |
| 4 | O | 171 | ARG |
| 4 | P | 32 | ASN |
| 4 | P | 171 | ARG |
| 5 | Q | 39 | ASN |
| 5 | Q | 43 | ARG |
| 5 | R | 39 | ASN |
| 5 | R | 43 | ARG |
| 5 | S | 39 | ASN |
| 5 | S | 43 | ARG |
| 5 | T | 39 | ASN |
| 5 | T | 43 | ARG |

Some sidechains can be flipped to improve hydrogen bonding and reduce clashes. All (273) such sidechains are listed below:

| Mol | Chain | Res | Type |
|---|---|---|---|
| 1 | A | 18 | GLN |
| 1 | A | 47 | ASN |
| 1 | A | 54 | HIS |
| 1 | A | 91 | ASN |
| 1 | A | 138 | ASN |
| 1 | A | 155 | ASN |
| 1 | A | 161 | GLN |
| 1 | A | 178 | GLN |
| 1 | A | 308 | ASN |
| 1 | A | 330 | GLN |
| 1 | A | 347 | GLN |
| 1 | A | 408 | ASN |
| 1 | A | 417 | ASN |
| 1 | A | 440 | ASN |
| 1 | A | 509 | ASN |
| 1 | A | 557 | ASN |
| 1 | A | 591 | ASN |
| 1 | A | 612 | ASN |
| 1 | A | 624 | ASN |
| 1 | A | 688 | ASN |
| 1 | A | 746 | HIS |

*Continued on next page...*

CONFIDENTIAL VALIDATION REPORT

Continued from previous page...

| Mol | Chain | Res | Type |
|---|---|---|---|
| 1 | A | 752 | GLN |
| 1 | A | 813 | GLN |
| 1 | A | 872 | HIS |
| 1 | B | 18 | GLN |
| 1 | B | 54 | HIS |
| 1 | B | 146 | GLN |
| 1 | B | 153 | ASN |
| 1 | B | 155 | ASN |
| 1 | B | 308 | ASN |
| 1 | B | 380 | HIS |
| 1 | B | 388 | ASN |
| 1 | B | 408 | ASN |
| 1 | B | 417 | ASN |
| 1 | B | 506 | ASN |
| 1 | B | 509 | ASN |
| 1 | B | 537 | GLN |
| 1 | B | 567 | GLN |
| 1 | B | 609 | ASN |
| 1 | B | 612 | ASN |
| 1 | B | 624 | ASN |
| 1 | B | 746 | HIS |
| 1 | B | 772 | GLN |
| 1 | B | 777 | GLN |
| 1 | B | 834 | GLN |
| 1 | C | 47 | ASN |
| 1 | C | 155 | ASN |
| 1 | C | 158 | ASN |
| 1 | C | 185 | GLN |
| 1 | C | 187 | GLN |
| 1 | C | 308 | ASN |
| 1 | C | 347 | GLN |
| 1 | C | 388 | ASN |
| 1 | C | 429 | ASN |
| 1 | C | 440 | ASN |
| 1 | C | 510 | HIS |
| 1 | C | 532 | HIS |
| 1 | C | 624 | ASN |
| 1 | C | 632 | ASN |
| 1 | C | 742 | GLN |
| 1 | C | 772 | GLN |
| 1 | C | 780 | ASN |
| 1 | C | 813 | GLN |

Continued on next page...

CONFIDENTIAL VALIDATION REPORT

*Continued from previous page...*

| Mol | Chain | Res | Type |
|---|---|---|---|
| 1 | C | 834 | GLN |
| 1 | C | 872 | HIS |
| 1 | D | 54 | HIS |
| 1 | D | 122 | ASN |
| 1 | D | 138 | ASN |
| 1 | D | 153 | ASN |
| 1 | D | 155 | ASN |
| 1 | D | 178 | GLN |
| 1 | D | 243 | ASN |
| 1 | D | 282 | GLN |
| 1 | D | 293 | GLN |
| 1 | D | 308 | ASN |
| 1 | D | 365 | GLN |
| 1 | D | 388 | ASN |
| 1 | D | 537 | GLN |
| 1 | D | 612 | ASN |
| 1 | D | 624 | ASN |
| 1 | D | 644 | ASN |
| 1 | D | 714 | ASN |
| 1 | D | 742 | GLN |
| 1 | D | 746 | HIS |
| 1 | D | 748 | ASN |
| 1 | D | 777 | GLN |
| 1 | D | 780 | ASN |
| 1 | D | 797 | ASN |
| 1 | D | 813 | GLN |
| 1 | D | 869 | ASN |
| 1 | E | 14 | HIS |
| 1 | E | 122 | ASN |
| 1 | E | 153 | ASN |
| 1 | E | 155 | ASN |
| 1 | E | 232 | ASN |
| 1 | E | 320 | ASN |
| 1 | E | 330 | GLN |
| 1 | E | 341 | ASN |
| 1 | E | 388 | ASN |
| 1 | E | 440 | ASN |
| 1 | E | 537 | GLN |
| 1 | E | 587 | ASN |
| 1 | E | 597 | HIS |
| 1 | E | 624 | ASN |
| 1 | E | 632 | ASN |

*Continued on next page...*

CONFIDENTIAL VALIDATION REPORT

Continued from previous page...

| Mol | Chain | Res | Type |
|---|---|---|---|
| 1 | E | 688 | ASN |
| 1 | E | 746 | HIS |
| 1 | E | 752 | GLN |
| 1 | E | 780 | ASN |
| 1 | E | 834 | GLN |
| 1 | E | 863 | GLN |
| 1 | E | 864 | ASN |
| 1 | F | 54 | HIS |
| 1 | F | 62 | GLN |
| 1 | F | 146 | GLN |
| 1 | F | 155 | ASN |
| 1 | F | 168 | ASN |
| 1 | F | 178 | GLN |
| 1 | F | 226 | GLN |
| 1 | F | 298 | ASN |
| 1 | F | 308 | ASN |
| 1 | F | 341 | ASN |
| 1 | F | 347 | GLN |
| 1 | F | 417 | ASN |
| 1 | F | 478 | ASN |
| 1 | F | 506 | ASN |
| 1 | F | 509 | ASN |
| 1 | F | 609 | ASN |
| 1 | F | 612 | ASN |
| 1 | F | 632 | ASN |
| 1 | F | 731 | GLN |
| 1 | F | 772 | GLN |
| 1 | F | 777 | GLN |
| 1 | F | 834 | GLN |
| 1 | G | 9 | GLN |
| 1 | G | 47 | ASN |
| 1 | G | 146 | GLN |
| 1 | G | 178 | GLN |
| 1 | G | 226 | GLN |
| 1 | G | 298 | ASN |
| 1 | G | 308 | ASN |
| 1 | G | 341 | ASN |
| 1 | G | 388 | ASN |
| 1 | G | 408 | ASN |
| 1 | G | 449 | ASN |
| 1 | G | 464 | ASN |
| 1 | G | 470 | ASN |

Continued on next page...

CONFIDENTIAL VALIDATION REPORT

*Continued from previous page...*

| Mol | Chain | Res | Type |
|---|---|---|---|
| 1 | G | 478 | ASN |
| 1 | G | 537 | GLN |
| 1 | G | 572 | ASN |
| 1 | G | 587 | ASN |
| 1 | G | 598 | ASN |
| 1 | G | 742 | GLN |
| 1 | G | 746 | HIS |
| 1 | G | 770 | ASN |
| 1 | G | 788 | GLN |
| 1 | G | 795 | GLN |
| 1 | G | 796 | HIS |
| 1 | H | 153 | ASN |
| 1 | H | 158 | ASN |
| 1 | H | 185 | GLN |
| 1 | H | 226 | GLN |
| 1 | H | 264 | ASN |
| 1 | H | 341 | ASN |
| 1 | H | 408 | ASN |
| 1 | H | 440 | ASN |
| 1 | H | 591 | ASN |
| 1 | H | 742 | GLN |
| 1 | H | 770 | ASN |
| 1 | H | 777 | GLN |
| 1 | H | 797 | ASN |
| 1 | H | 813 | GLN |
| 1 | H | 872 | HIS |
| 1 | I | 54 | HIS |
| 1 | I | 182 | GLN |
| 1 | I | 185 | GLN |
| 1 | I | 189 | ASN |
| 1 | I | 245 | GLN |
| 1 | I | 316 | ASN |
| 1 | I | 341 | ASN |
| 1 | I | 380 | HIS |
| 1 | I | 417 | ASN |
| 1 | I | 510 | HIS |
| 1 | I | 557 | ASN |
| 1 | I | 624 | ASN |
| 1 | I | 746 | HIS |
| 1 | I | 777 | GLN |
| 1 | I | 798 | ASN |
| 1 | I | 864 | ASN |

*Continued on next page...*

CONFIDENTIAL VALIDATION REPORT

*Continued from previous page...*

| Mol | Chain | Res | Type |
|---|---|---|---|
| 1 | J | 54 | HIS |
| 1 | J | 122 | ASN |
| 1 | J | 153 | ASN |
| 1 | J | 155 | ASN |
| 1 | J | 158 | ASN |
| 1 | J | 161 | GLN |
| 1 | J | 168 | ASN |
| 1 | J | 178 | GLN |
| 1 | J | 245 | GLN |
| 1 | J | 254 | ASN |
| 1 | J | 294 | GLN |
| 1 | J | 298 | ASN |
| 1 | J | 308 | ASN |
| 1 | J | 330 | GLN |
| 1 | J | 341 | ASN |
| 1 | J | 379 | ASN |
| 1 | J | 470 | ASN |
| 1 | J | 478 | ASN |
| 1 | J | 510 | HIS |
| 1 | J | 614 | GLN |
| 1 | J | 624 | ASN |
| 1 | J | 632 | ASN |
| 1 | J | 742 | GLN |
| 1 | K | 9 | GLN |
| 1 | K | 54 | HIS |
| 1 | K | 178 | GLN |
| 1 | K | 298 | ASN |
| 1 | K | 341 | ASN |
| 1 | K | 470 | ASN |
| 1 | K | 472 | ASN |
| 1 | K | 478 | ASN |
| 1 | K | 509 | ASN |
| 1 | K | 557 | ASN |
| 1 | K | 612 | ASN |
| 1 | K | 624 | ASN |
| 1 | K | 632 | ASN |
| 1 | K | 688 | ASN |
| 1 | K | 798 | ASN |
| 1 | K | 864 | ASN |
| 1 | K | 902 | HIS |
| 1 | L | 146 | GLN |
| 1 | L | 169 | GLN |

*Continued on next page...*

CONFIDENTIAL VALIDATION REPORT

*Continued from previous page...*

| Mol | Chain | Res | Type |
|---|---|---|---|
| 1 | L | 185 | GLN |
| 1 | L | 196 | GLN |
| 1 | L | 221 | ASN |
| 1 | L | 327 | GLN |
| 1 | L | 332 | ASN |
| 1 | L | 338 | GLN |
| 1 | L | 341 | ASN |
| 1 | L | 347 | GLN |
| 1 | L | 365 | GLN |
| 1 | L | 440 | ASN |
| 1 | L | 464 | ASN |
| 1 | L | 537 | GLN |
| 1 | L | 591 | ASN |
| 1 | L | 624 | ASN |
| 1 | L | 632 | ASN |
| 1 | L | 644 | ASN |
| 1 | L | 707 | ASN |
| 1 | L | 780 | ASN |
| 1 | L | 797 | ASN |
| 1 | L | 798 | ASN |
| 1 | L | 813 | GLN |
| 2 | M | 86 | HIS |
| 2 | M | 90 | GLN |
| 2 | M | 181 | ASN |
| 2 | M | 187 | ASN |
| 2 | M | 372 | GLN |
| 2 | M | 463 | HIS |
| 2 | M | 476 | GLN |
| 3 | N | 49 | ASN |
| 3 | N | 215 | ASN |
| 4 | O | 14 | GLN |
| 4 | O | 16 | GLN |
| 4 | O | 25 | GLN |
| 4 | O | 32 | ASN |
| 4 | O | 53 | GLN |
| 4 | O | 58 | GLN |
| 4 | P | 25 | GLN |
| 4 | P | 32 | ASN |
| 5 | Q | 39 | ASN |
| 5 | R | 39 | ASN |
| 5 | S | 39 | ASN |
| 5 | T | 39 | ASN |

CONFIDENTIAL VALIDATION REPORT

##### 5.3.3 RNA (i)

There are no RNA molecules in this entry.

#### 5.4 Non-standard residues in protein, DNA, RNA chains (i)

There are no non-standard protein/DNA/RNA residues in this entry.

#### 5.5 Carbohydrates (i)

There are no carbohydrates in this entry.

#### 5.6 Ligand geometry (i)

There are no ligands in this entry.

#### 5.7 Other polymers (i)

There are no such residues in this entry.

#### 5.8 Polymer linkage issues (i)

There are no chain breaks in this entry.

CONFIDENTIAL VALIDATION REPORT

### 6 Map visualisation (i)

This section contains visualisations of the EMDB entry EMD-10768. These are intended to permit visual inspection of the internal detail of the map and identification of artifacts.

#### 6.1 Orthogonal projections (i)

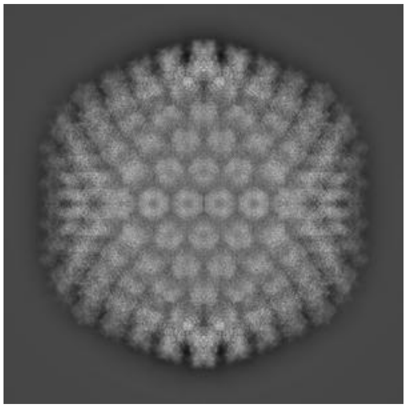
X

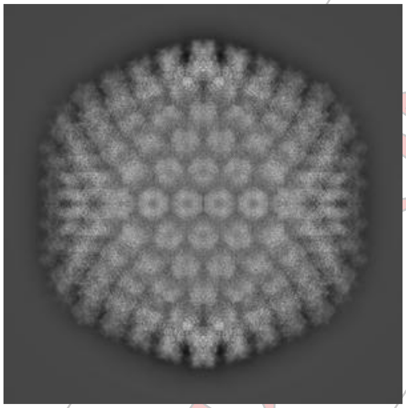
Y

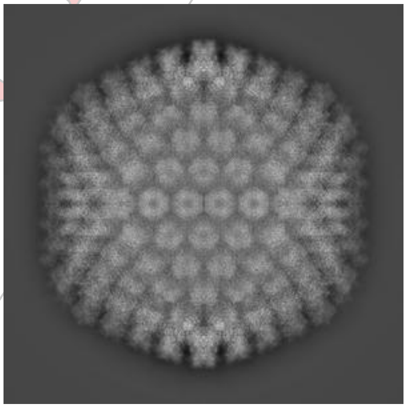
Z

The images above show the map projected in three orthogonal projections, in greyscale.

#### 6.2 Central slices (i)

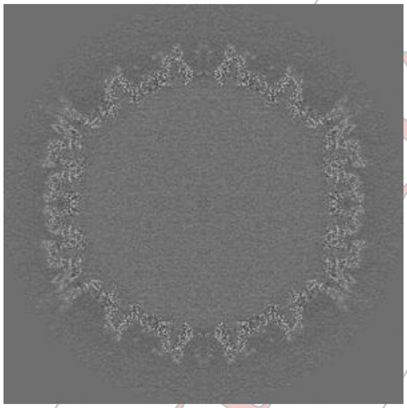
X

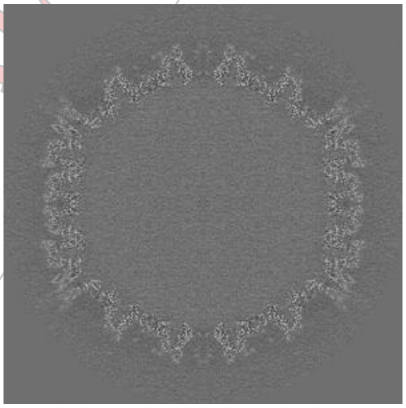
Y

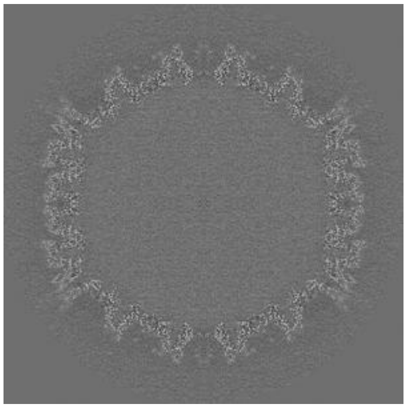
Z

The images above show central slices of the map in three orthogonal directions, in greyscale.

CONFIDENTIAL VALIDATION REPORT

#### 6.3 Largest variance slices (i)

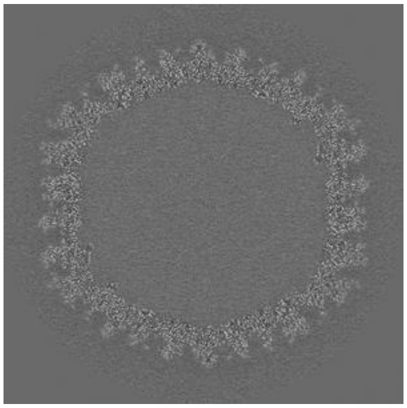

X Index: 405

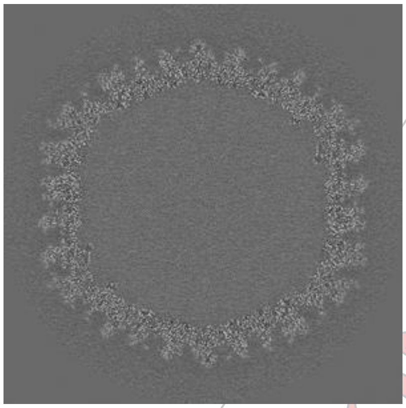

Y Index: 405

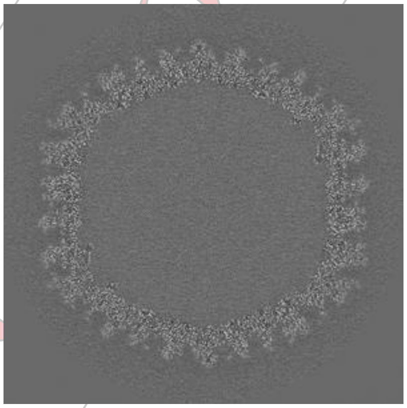

Z Index: 405

The images above show the highest variance slices of the map in three orthogonal directions, in greyscale.

#### 6.4 Orthogonal surface views (i)

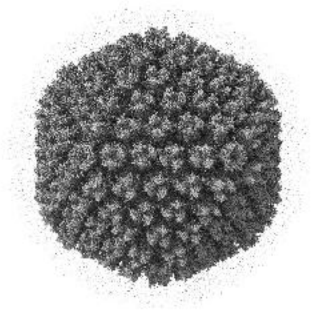

X

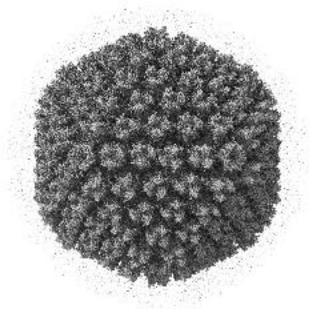

Y

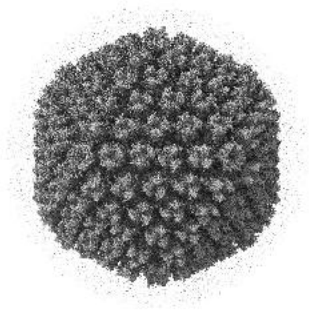

Z

The images above show the 3D surface view of the map at the recommended contour level 0.07. This in conjunction with the slice images can indicate whether an appropriate contour level has been selected.

#### 6.5 Mask visualisation (i)

This section was not generated. No masks were provided.

CONFIDENTIAL - VALIDATION REPORT

### 7 Map analysis (i)

This section contains the results of statistical analysis of the map.

#### 7.1 Map-value distribution (i)

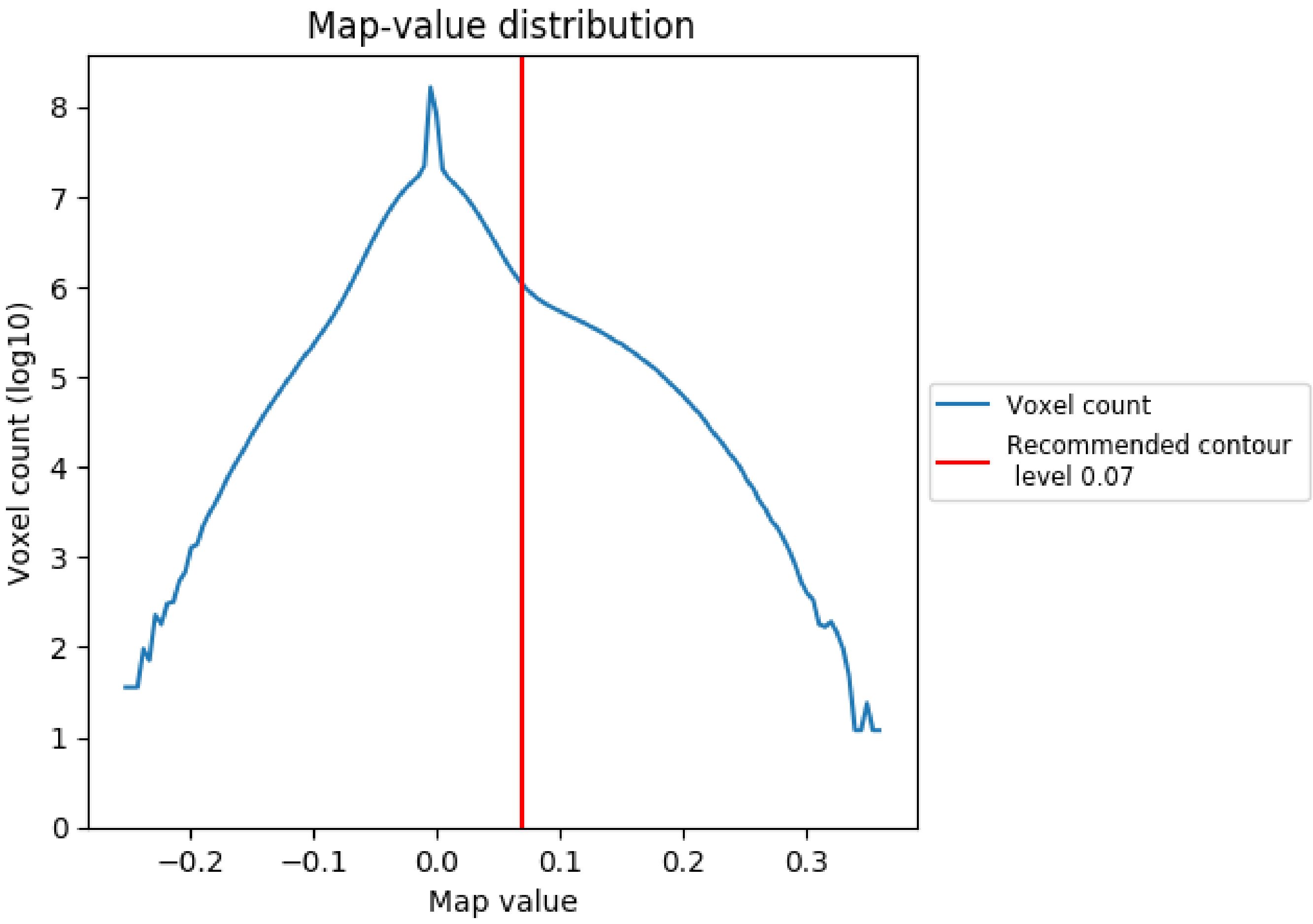


The map-value distribution is plotted in 128 intervals along the x-axis. The y-axis is logarithmic. A spike in this graph at zero usually indicates that the volume has been masked.

CONFIDENTIAL REPORT

#### 7.2 Volume estimate (i)

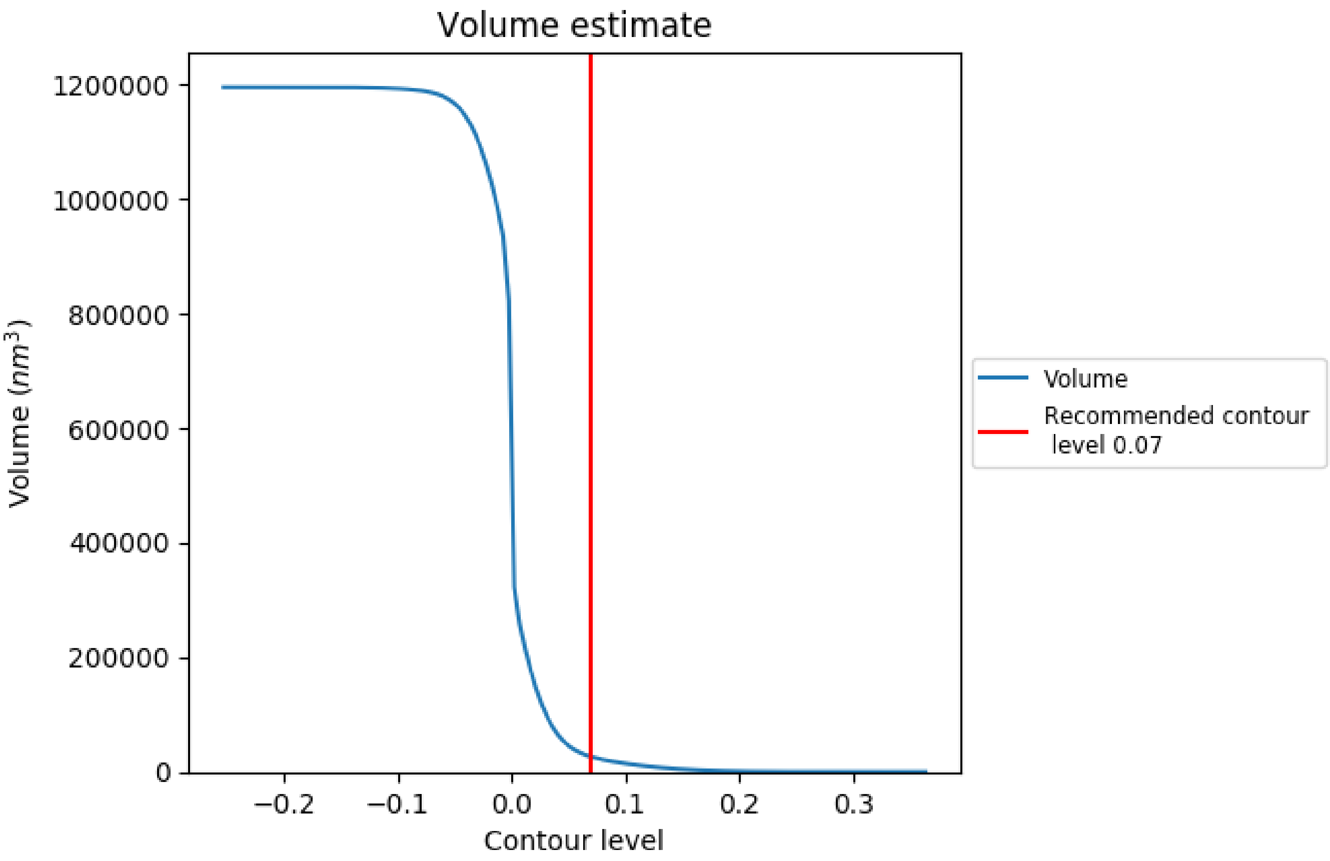


The volume at the recommended contour level is 26562 nm$^3$; this corresponds to an approximate mass of 23994 kDa.

The volume estimate graph shows how the enclosed volume varies with the contour level. The recommended contour level is shown as a vertical line and the intersection between the line and the curve gives the volume of the enclosed surface at the given level.

CONFIDENTIAL REPORT

#### 7.3 Rotationally averaged power spectrum ⓘ

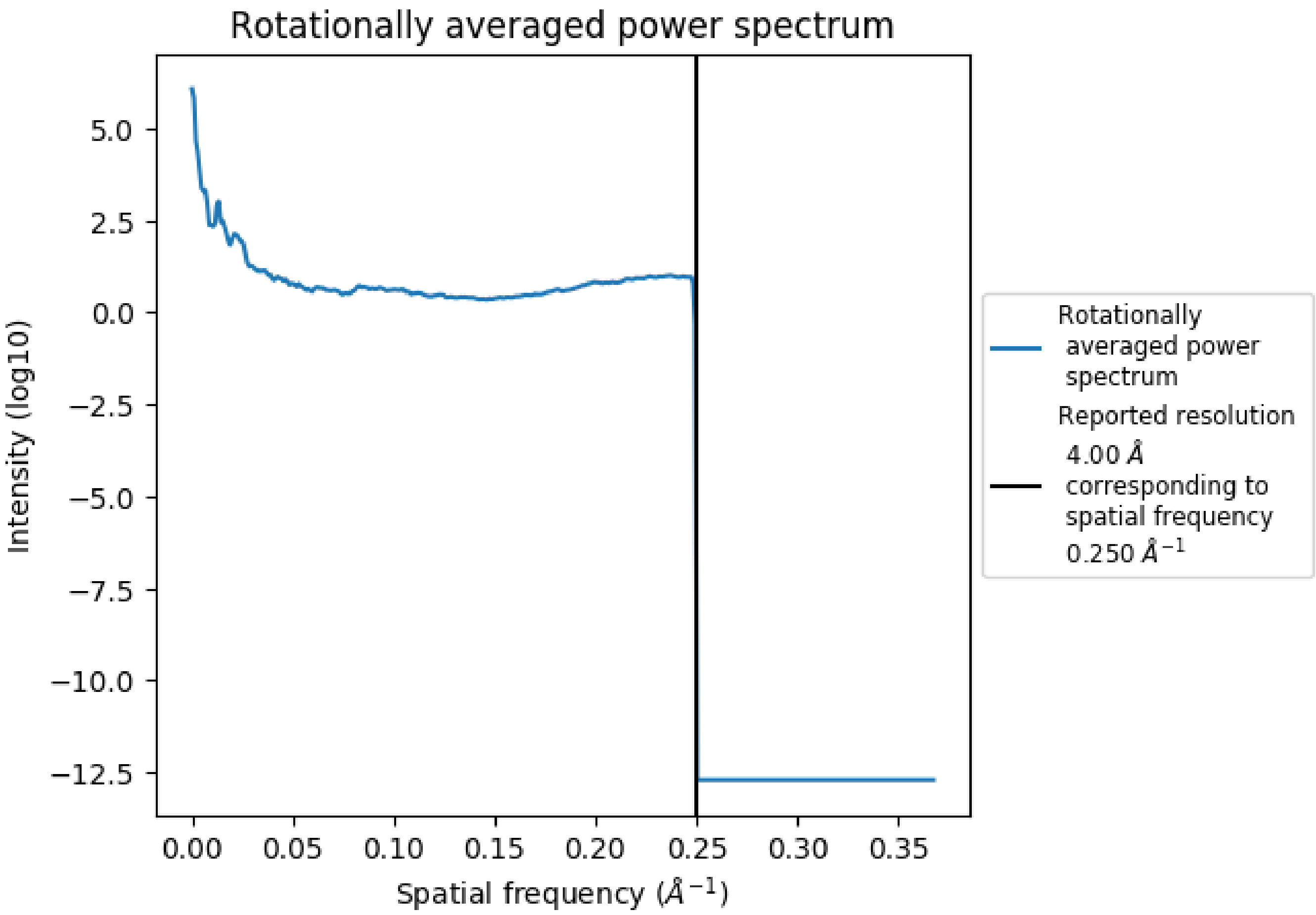


CONFIDENTIAL REPORT

### 8 Fourier-Shell correlation (i)

This section was not generated. No FSC curve or half maps provided.

CONFIDENTIAL VALIDATION REPORT

### 9 Map-model fit (i)

This section contains information regarding the fit between EMDB map EMD-10768 and PDB model 6YBA. Per-residue inclusion information can be found in section 3 on page 6.

#### 9.1 Map-model overlay (i)

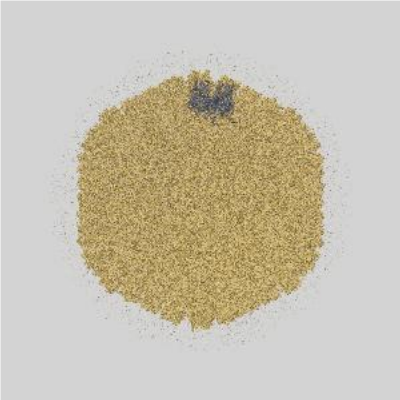

X

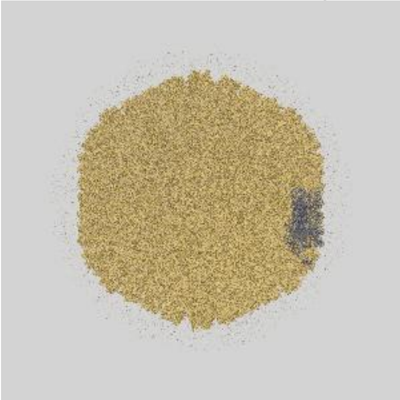

Y

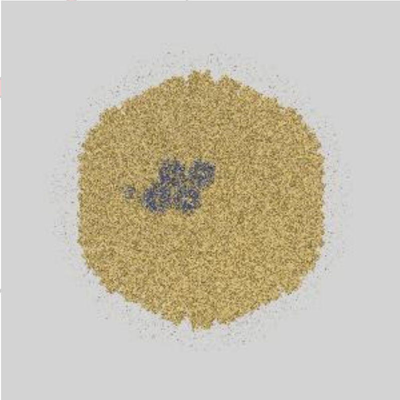

Z

The images above show the 3D surface view of the map at the recommended contour level 0.07 at 50% transparency in yellow overlaid with a ribbon representation of the model coloured in blue. These images allow for the visual assessment of the quality of fit between the atomic model and the map.

CONFIDENTIAL VALIDATION REPORT

#### 9.2 Atom inclusion ⓘ

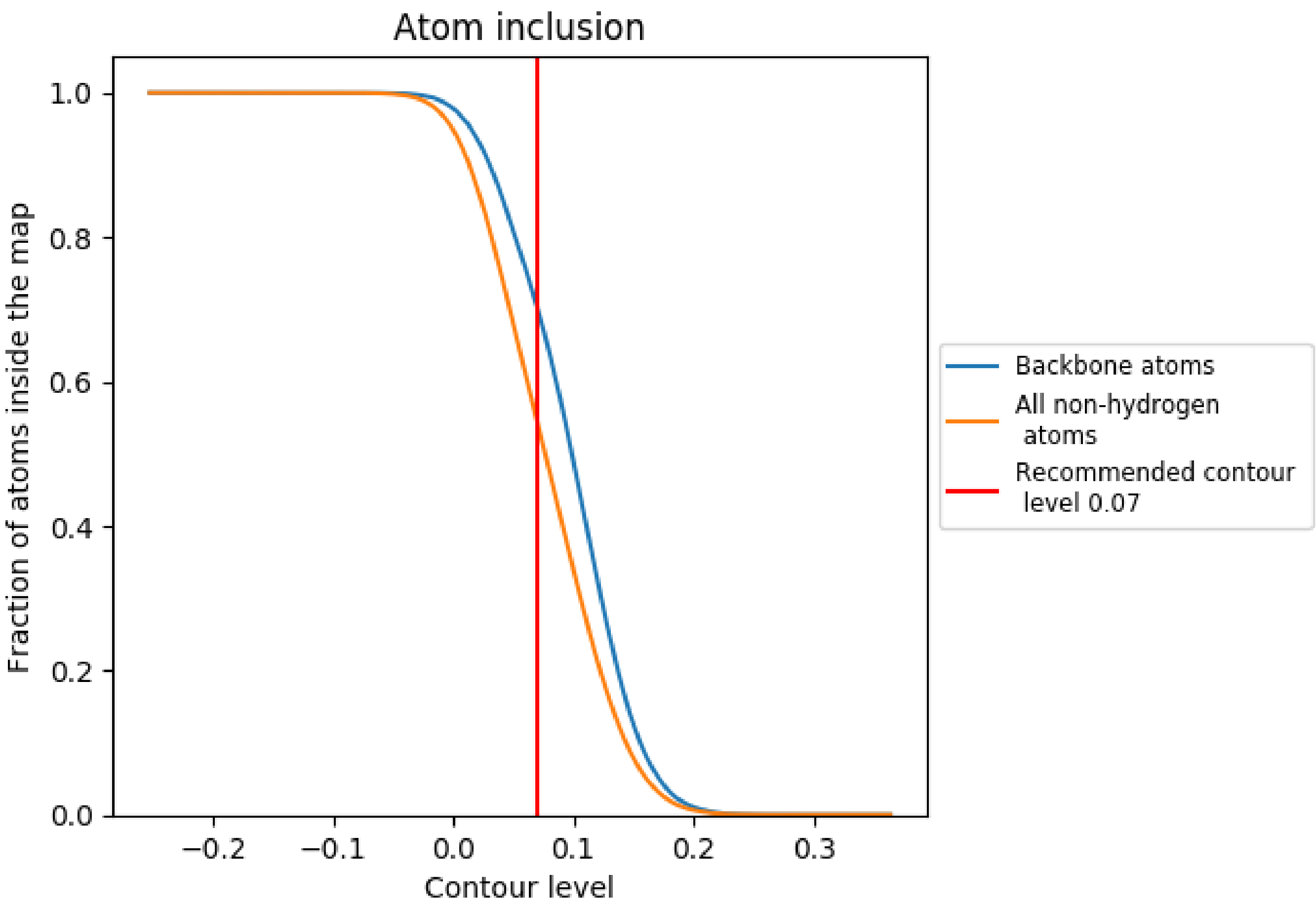


At the recommended contour level, 70% of all backbone atoms, 55% of all non-hydrogen atoms, are inside the map.

CONFIDENTIAL
